# A mismatch between striatal cholinergic pauses and dopaminergic reward prediction errors

**DOI:** 10.1101/2024.05.09.593336

**Authors:** Mariana Duhne, Ali Mohebi, Kyoungjun Kim, Lilian Pelattini, Joshua Berke

## Abstract

Movement, motivation and reward-related learning depend strongly on striatal dopamine and acetylcholine. These neuromodulators each regulate the other, and disturbances to their coordinated signals contribute to human disorders ranging from Parkinson’s Disease to depression and addiction. Pauses in the firing of cholinergic interneurons (CINs) are thought to coincide with pulses in dopamine release that encode reward prediction errors (RPEs), together shaping synaptic plasticity and thereby learning. However, such models are based upon recordings from unidentified neurons, and do not incorporate the distinct characteristics of striatal subregions. Here we compare the firing of identified, individual CINs to dopamine release as unrestrained rats performed a probabilistic decision-making task. The relationships between CIN spiking, dopamine release, and behavior varied strongly by subregion. In dorsal-lateral striatum a *Go!* cue evoked burst-pause CIN spiking, quickly followed by a very brief (~150ms) dopamine pulse that was unrelated to RPE. In dorsal-medial striatum the same cue evoked only a CIN pause; this pause was curtailed by a movement-selective rebound in firing. Finally in ventral striatum a reward cue evoked slower, RPE-coding increases in both dopamine and CIN firing, without any distinct pause. Our results demonstrate a spatial and temporal dissociation between CIN pauses and dopamine RPE signals, and will inform new models of striatal microcircuits and their contributions to behavior.

## Introduction

A striking feature of vertebrate brains is the very dense network of intertwined dopaminergic (DA) and cholinergic (ACh) axons within the striatum (Descarries & Mechawar, 2000; Zhou et al., 2001; Nosaka & Wickens, 2022). While DA axons originate from cell bodies in the midbrain, striatal ACh is provided predominantly by local, large aspiny cholinergic interneurons (CINs). Both modulators have powerful, complex, effects on striatal circuitry (Abudukeyoumu et al., 2018), and influence each other’s release, via nicotinic ACh receptors on DA axons (Kramer et al., 2022) and D2 DA receptors on CINs (Maurice et al., 2004; Kharkwal et al., 2016).

Investigation of movement disorders lead long ago to the idea that striatal function requires a “balance” between ACh and DA (Barbeau 1962; Aosaki et al., 2010; Sanchez-Catasus et al., 2022). Loss of striatal DA causes movement slowing in Parkinson’s Disease, which can be alleviated by ACh antagonists (Pisani et al., 2007) or (in animal models) by artificial suppression of CINs (Ztaou et al., 2016). Conversely, loss of normal CIN function in dorsal striatum is implicated in a range of uncontrolled movements including dyskinesias, tics and dystonias (Xu et al., 2015; Pappas et al., 2018; Eskow Jaunarajs et al., 2019). In ventral striatum, perturbed CIN function has been associated with depression and addiction (Warner-Schmidt et al., 2012, Cheng et al., 2019, Hikida et al. 2003).

However, the specific contributions of CINs, and striatal ACh-DA interactions, to normal behavior are poorly understood. Current thinking is based largely on classic electrophysiological recordings of unidentified neurons in head-fixed monkeys (Kimura et al., 1984; Apicella, 2017). In that context it was found that a minority of striatal cells fire continuously at relatively low rates (~5Hz; “tonically active neurons”, TANs). These TANs have been presumed to correspond to CINs, which are tonically active in brain slices and anesthetized rats (Bennett & Wilson, 1999; Inokawa et al., 2010; Doig et al., 2014). TANs typically show a brief (~150-250ms) pause in firing after salient cues, especially cues that evoke behavioral responses. The mechanisms shaping the onset and offset of this TAN pause have been studied extensively (albeit in slices rather than behaving animals). While the TAN pause is thought to be at least partly dependent on DA release (Aosaki et al., 1994; Ding et al., 2010; Kharkwal et al., 2016; Zhang et al., 2018; Gallo et al., 2022), the specific underlying mechanisms, and relationships to DA, continue to be actively debated (Schulz & Reynolds, 2012; Goldberg & Wilson, 2016; Tubert & Murer, 2020).

In elegant work, Morris et al. (2004) found that striatal TAN pauses coincide with burst firing of midbrain DA cells encoding reward prediction errors (RPEs). In computational models of reward-guided behavior (Sutton & Barto, 1981) RPEs are learning signals: they indicate that reward predictions need to be updated. Morris et al. proposed that the TAN pause defines a temporal window for plasticity - “*when*ဝto learn - that coincides with an RPE-coding DA pulse determining the extent of plasticity - “*how*” to learn (Yagishita et al., 2014; Yamanaka et al., 2017; Reynolds et al., 2022). The TAN pause, and its apparent close relationship to DA RPE signals, have inspired a wide range of computational models, centered especially upon the adaptive control of learning (Ashby & Crossley, 2011; Stocco, 2012; Franklin & Frank, 2015; Kim et al., 2019).

However there are multiple grounds for caution. The evidence that TANs correspond to CINs is indirect, and it is now known that several distinct classes of striatal cells can be tonically active in awake animals (Gage et al., 2010; Sharott et al., 2012). In particular GABAergic, somatostatin-expressing interneurons also possess intrinsic currents that drive continuous firing *ex vivo* (Beatty et al., 2012), and so would be expected to be tonically-active *in vivo*. The lack of positive identification in behaving animals could have contributed to observations that TAN firing patterns can vary across different areas of striatum (Yarom & Cohen, 2011; Marche et al., 2017). We also now know that DA release can differ between striatal subregions (Parker et al., 2016; Mohebi et al., 2024), rather than providing a homogeneous global RPE signal as originally proposed (Schultz, 1998). It is thus vital to observe the firing patterns of *identified* CINs and determine under what circumstances, and in which locations, cholinergic pauses actually coincide with DA RPE signals. Several recent studies have investigated CIN activity using optical approaches including photometry (e.g. Howe et al., 2019). However optical methods have generally lacked the ideal temporal resolution to study the brief pauses and other rapid dynamics of CIN firing (though see Shroff et al., 2023).

We therefore used electrophysiology with optogenetic tagging (Cardin et al., 2010) to positively identify CINs in freely behaving rats. We examined three distinct striatal subregions, often described as “sensorimotor” (dorsolateral, DLS), “associative” (dorsomedial, DMS) and “limbic” (ventral, VS; we targeted nucleus accumbens Core). We took advantage of a probabilistic reward (“bandit”) task in which a varying reward rate helps assess reward predictions and RPEs (Hamid et al., 2016). We compared CIN firing to DA release in the same striatal subregions, measured using a genetically encoded optical DA sensor with high temporal resolution (Mohebi et al., 2024). Our primary finding is that identified CINs can indeed show tonic firing with cue-evoked pauses, but that these pauses occur at different times, and in different locations, to RPE-coding DA increases. Furthermore, in ventral striatum CIN firing shows a matching, rather than opposite, pattern to DA release: both show an RPE-encoding increase in response to the bandit task reward cue. These findings necessitate a major revision of our understanding of striatal learning mechanisms, and provide a solid foundation for the development of new models.

## Results

### Identified cholinergic interneurons show a distinctive firing pattern, throughout striatum

To distinguish CINs in extracellular recordings we infused a virus for Cre-dependent expression of the red-light-sensitive cation channel Chrimson (Klapoetke et al., 2014) (AAV5-Syn-FLEX-rc[ChrimsonR-tdTomato]) into the striatum of ChAT-Cre rats (Witten et al., 2011). Post-mortem histology confirmed high selectivity of Chrimson expression (Fig. 1A; 96.4 %, 240/249 of sampled Chrimson-expressing cells also expressed choline acetyltransferase (ChAT+; see Methods).

**Figure 1.**
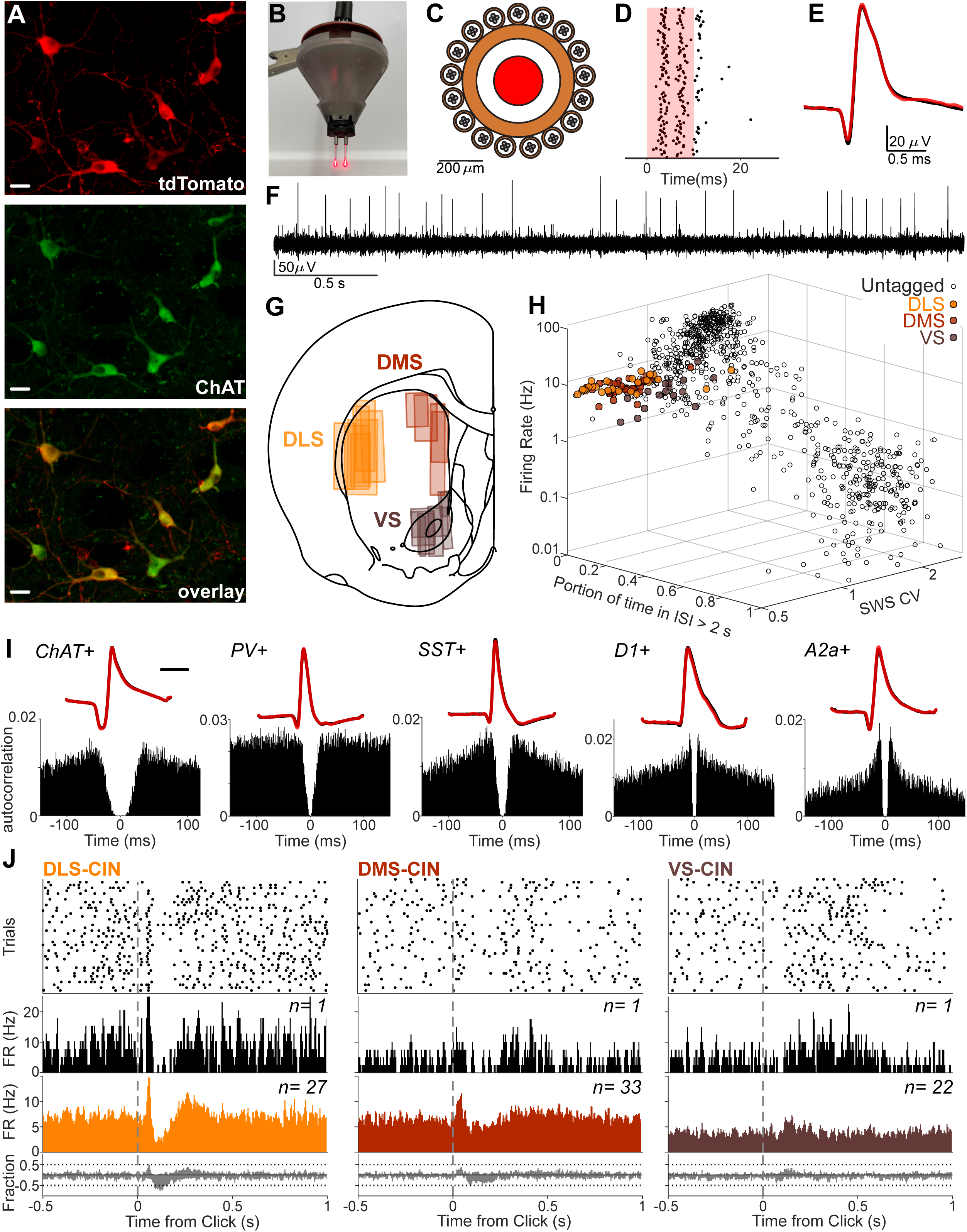
Distinct characteristics of optogenetically identified CINs. **A.** Colocalization of tdTomato expression with ChAT staining (scale bars, 20 µm). **B.** Picture of assembly before bilateral implantation. **C.** Schematic of the cage of tetrodes surrounding each optic fiber. **D.** Example responses of a Chrimson-expressing CIN to red light pulses (10ms duration, 100 presentations). **E.** Mean waveforms of the same example CIN as D, during light delivery (red) and spontaneous firing (black). **F.** Example snippet of (wavelet-filtered) signal from one tetrode wire; the larger visible spikes are produced by one identified CIN. **G.** Diagram of recorded striatal subregions. Each rectangle corresponds to the reconstructed recording zone for one implant (see Supp. Fig. 1 for histology). **H.** Scatter plot of each isolated neuron’s firing rate, CV during slow wave sleep (SWS) and portion of time in ISI> 2s. Filled colored dots are ChAT+ interneurons from different subregions. Empty circles are unidentified DLS units. **I.** Example waveforms (top) and autocorrelograms (bottom) of optogenetically identified striatal neurons in DLS. CIN example is from this study, remainder are from optotagging studies of projection neurons (expressing dopamine D1 receptors, D1+, or adenosine 2a receptors, A2a+; Gonzalez Montalvo et al. 2023) and GABAergic interneurons (parvalbumin, PV+, and somatostatin, SST+; Duhne & Berke, 2024). Scale bars, 20 µm. **J.** Responses of CINs from different subregions (DLS, DMS and VS) to unexpected reward delivery. From top: activity rasters for example single neurons, aligned to reward delivery (‘Click’); corresponding perievent time histograms (PETHs); averaged PETHs for all CINs tested; fractions of CINs significantly up- or down-modulated at each time point (10 ms, 50% overlapping bins, shuffle test p < 0.01, 10,000 shuffles).

Into each brain hemisphere we also implanted an optic fiber surrounded by a driveable “cage” of 16 tetrodes (each 4 x 12.5µm diameter wires; 128 total channels/rat; Fig. 1B-C). We recorded neural activity for 1-2 hours during behavioral task performance (see below), then as rats rested quietly (including bouts of sleep). Towards the end of each recording session we gave a series of brief red light pulses through the fiber to evoke spiking in Chrimson-expressing cells (Fig. 1D). Cells with reliable spiking within 10ms of light onset were considered to be Chrimson-expressing and therefore ChAT+ (see Methods for full criteria).

We tagged 101 distinct CINs across our three target regions (Fig. 1 G-H; Supp. Fig. 1; n=33 DLS, 45 DMS, 23 VS), from 56 recording sessions in 12 rats. In the same sessions we recorded and isolated an additional 2,384 unidentified neurons. The majority of these had only sporadic activity and were presumed to be projection neurons, but as in prior studies we also observed cells with high mean firing rates that are presumed to be other classes of interneurons (Fig. 1H; Gage et al., 2010).

All tagged CINs were tonically-active (Figs. 1F, 1H, Supp Fig. 2A). This tonic quality was quantified as a low proportion of time spent not spiking (i.e. within inter-spike intervals > 2s; mean 1.99 %, range 0.73 % - 17.3%, Fig. 1H, Supp. Fig. 2A; Schmitzer-Torbert & Redish, 2008). Average CIN firing rates ranged from ~2-10Hz, with a significant effect of subregion (mean +-SEM: DLS 6.90 +-0.05; DMS 6.18 +-0.026; VS 4.2 +-0.0652; 1-way ANOVA, F = 24.67, p = 2.02 x 10^−9^). Identified CINs also fired more regularly compared to the striatal population as a whole, as quantified by coefficient of variation (CV; mean 1.01, range 0.60 - 1.84, Supp. Fig. 3F). As a result, tagged CINs clustered together when firing rate, proportion-of-time-in-long-ISIs, and CV were jointly plotted (Fig. 1H). This CIN cluster was apparent within each of the three subregions individually (Supp. Fig. 2C).

Compared to other classes of identified striatal neurons, CINs also typically showed longer spike waveforms and greater “post-spike suppression” (absence of short ISIs; examples, Fig. 1I, for population see Supp. Fig. 2B). This is consistent with prior results for presumed CINs (Schmitzer-Torbert & Redish, 2008; Thorn & Graybiel, 2014; Peters et al., 2021; Krok et al., 2023). However, some identified CINs had either short waveform duration or little post-spike suppression, and so neither measure was reliable for CIN identification (Supp. Fig. 2D-E). We also examined whether CINs show distinct firing patterns across the sleep-wake cycle (which can be useful for establishing cell identity without optogenetic tagging; e.g. Mallet et al., 2016). Identified CINs consistently showed slightly higher firing when rats were awake, compared to slow-wave sleep (SWS; 1-way ANOVA, F = 64.39, p = 2.17 x 10^−13^; KS test p = 2.15 x 10^−10^; Supp. Fig. 3C). Overall however CIN firing was not very different between awake and sleep states (Supp. Fig. 3B-F), as expected from prior work (Sharott et al. 2012).

To begin assessment of the functional correlates of identified CINs, we examined their responses to unexpected presentation of a reward cue, within an operant box. The brief, familiar “click” sound of a food hopper (sucrose pellet delivery) evoked firing changes in many CINs (Fig. 1J). In dorsal striatum this response closely resembled the classic TAN pause. A significant decrease in firing 100-250ms after the click was observed in 21/27 DLS CINs (78%), and 20/33 DMS CINs (61%) (p < 0.01, two-way shuffle test using 10,000 random time points, correcting for multiple comparisons). About 50% of DLS CINs also showed a brief significant increase in firing preceding this pause (Fig. 1J). VS CIN responses were less common and less consistent: similar minorities of cells increased and decreased firing after click onset, resulting in little change on average (Fig. 1J).

### CINs and dopamine release show subregion-specific response profiles during operant performance

Before implantation rats had received substantial (>3 months) prior training in the bandit task (Fig. 2A; Hamid et al., 2016; Mohebi et al., 2019). In this task the illumination of a nose-poke port (Light On) encourages approach and entry into that port (Center In). The rat maintains this position over a variable delay (500-1500ms, uniform distribution) until presentation of an auditory *Go!* cue (white noise burst), then quickly withdraws (Center Out) and pokes another port immediately to the left or right (Side In). At that time - on a subset of trials - the rat hears the click of the food hopper and moves to the food port to collect reward (Food In). Reward probabilities (10, 50, or 90%) were set independently for left and right choices, and changed every 35-45 trials without warning.

**Figure 2.**
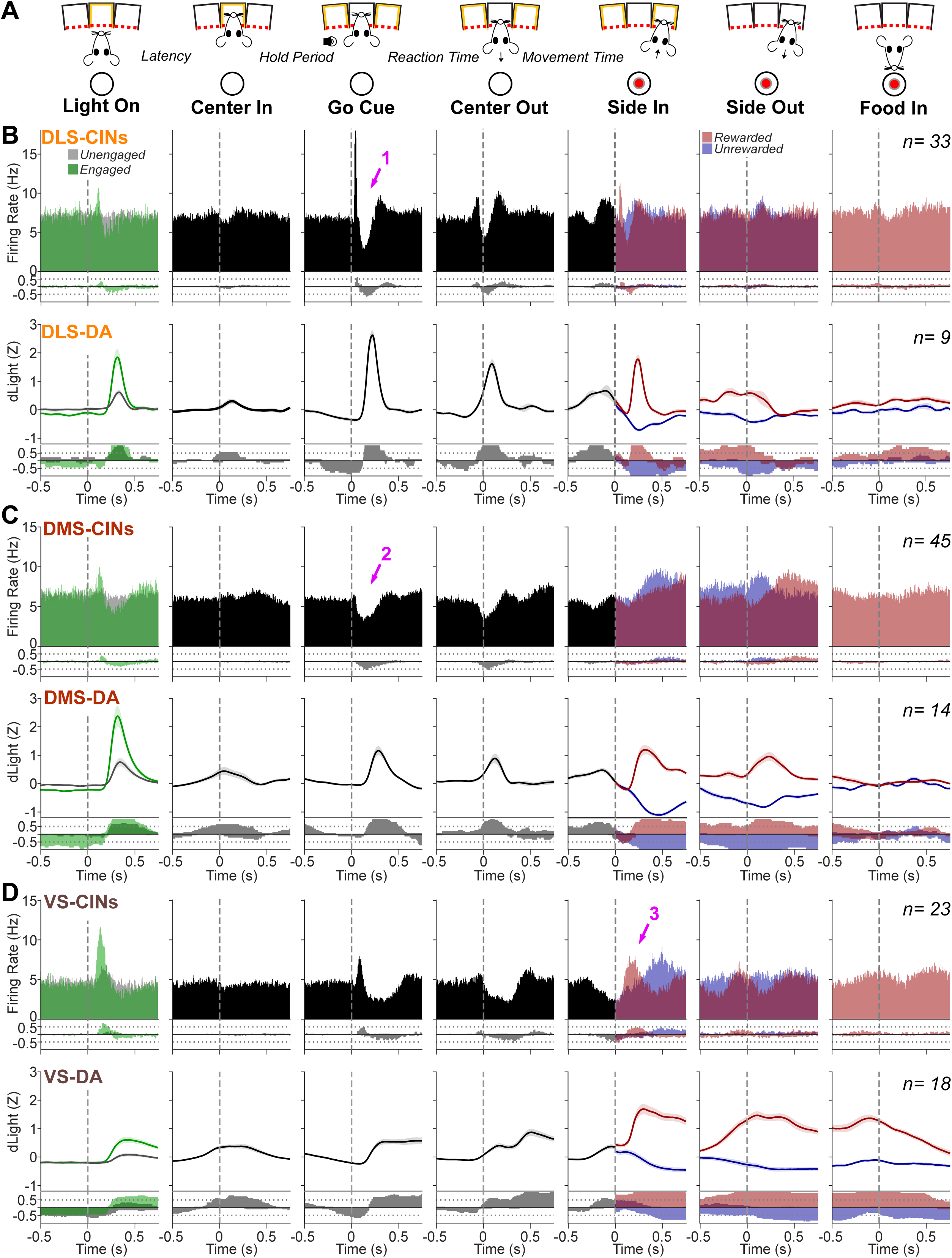
Behavior-related CIN firing and DA release are consistent within-, but not between-, subregions. **A.** Events within each trial of the bandit task. **B.** Top: Peri-event firing averaged across all identified DLS CINs. 10 ms bins, sliding in 5ms steps. For the Light On event, trials are separated by “engaged” (latencies < 1s; green) and “unengaged” (latencies > 1s; gray). For the Side In event, trials are separated by rewarded (with food hopper click; red) and unrewarded (blue). For the Food In event, only rewarded trials are shown since on unrewarded trials rats typically did not enter the food port (<10% of unrewarded trials). Arrow “1” points to the burst-pause sequence after the *Go!* cue. Next row: fraction of significantly modulated units per time bin. Firing increases are upward and decreases downward (shuffle test, p<0.005, 10,000 shuffles, corrected for multiple comparisons). Next row, perievent DA release in DLS, measured using dLight1.3b fiber photometry. Signals were Z-scored before averaging across fiber placements. Data are from 13 rats over 15 recording sessions (Mohebi et al., 2024). Bottom, fraction of fiber placements significantly up- or down-modulated at each time point (shuffle test, p<0.005, 10,000 shuffles, corrected for multiple comparisons). **C,** Same as B, but for DMS. Arrow “2” points to the CIN pause after the Go Cue. **D,** Same as B, but for VS. Arrow “3” points to the CIN firing increase after reward delivery.

We examined the activity of identified CINs around each task event, in each striatal subregion (Fig. 2B-D; firing patterns for each individual CIN are shown in the Supplementary File). Average CIN activity differed clearly between subregions (Fig. 2B-D, upper portions). These differences arose from distinct activity patterns that were quite consistent within each subregion (i.e. were shared by a majority of cells; Fig. 2B-D, fraction plots). Regional differences in CIN firing were particularly apparent following cue onsets. After the *Go!* cue most DLS CINs showed the classic three-component TAN sequence of “burst-pause-rebound” (arrow 1) while in DMS, only the pause was prominent and common (arrow 2). In VS the *Go!* cue provoked a slower increase in CIN firing (compared to DLS), with a less well-defined pause. The reward delivery cue at Side-In increased firing of most CINs in VS (arrow 3, n=16/23), but only a minority in DLS and virtually none in DMS.

Many DA neurons also show fast responses to salient events (Schultz & Romo, 1990; Mohebi et al., 2019). and often such DA responses can be more readily interpreted as “alerting” or “detection”, rather than RPE (Bromberg-Martin & Hikosaka, 2010; Pasquereau & Turner, 2015; Schultz, 2016). We expected that the relative strength of responding to each cue, in each location, would show a close correspondence between DA and CINs. To test this we compared DA release around the same events, in the same subregions (in a separate set of rats; Mohebi et al. 2019, 2024). DLS DA increased sharply after the *Go!* cue, matching the strong DLS CIN response at this time. However, at Light On the VS response was relatively strong for CINs and weak for DA, while the opposite was observed for DMS. Thus, while striatal DA and ACh both respond to behaviorally relevant cues, they show distinct response profiles across subregions.

### Fast CIN firing and dopamine release dynamics during cue-evoked movement initiation

We next looked in more detail at key event-related features of CIN firing and their temporal relationships to DA release. We began with the *Go!* cue which - at least in DLS CINs - evokes the “burst-pause-rebound” compound response most associated with TANs (Fig. 3A, B). We examined the timing of *Go!* cue-evoked increases and decreases in CIN firing (Fig. 3C, D). Further, we determined if each activity component is more closely related to cue onset, or to the subsequent movement (Fig. 3E) and its specific left/right direction (assessed as contraversive or ipsiversive, relative to the brain recording hemisphere).

**Figure 3.**
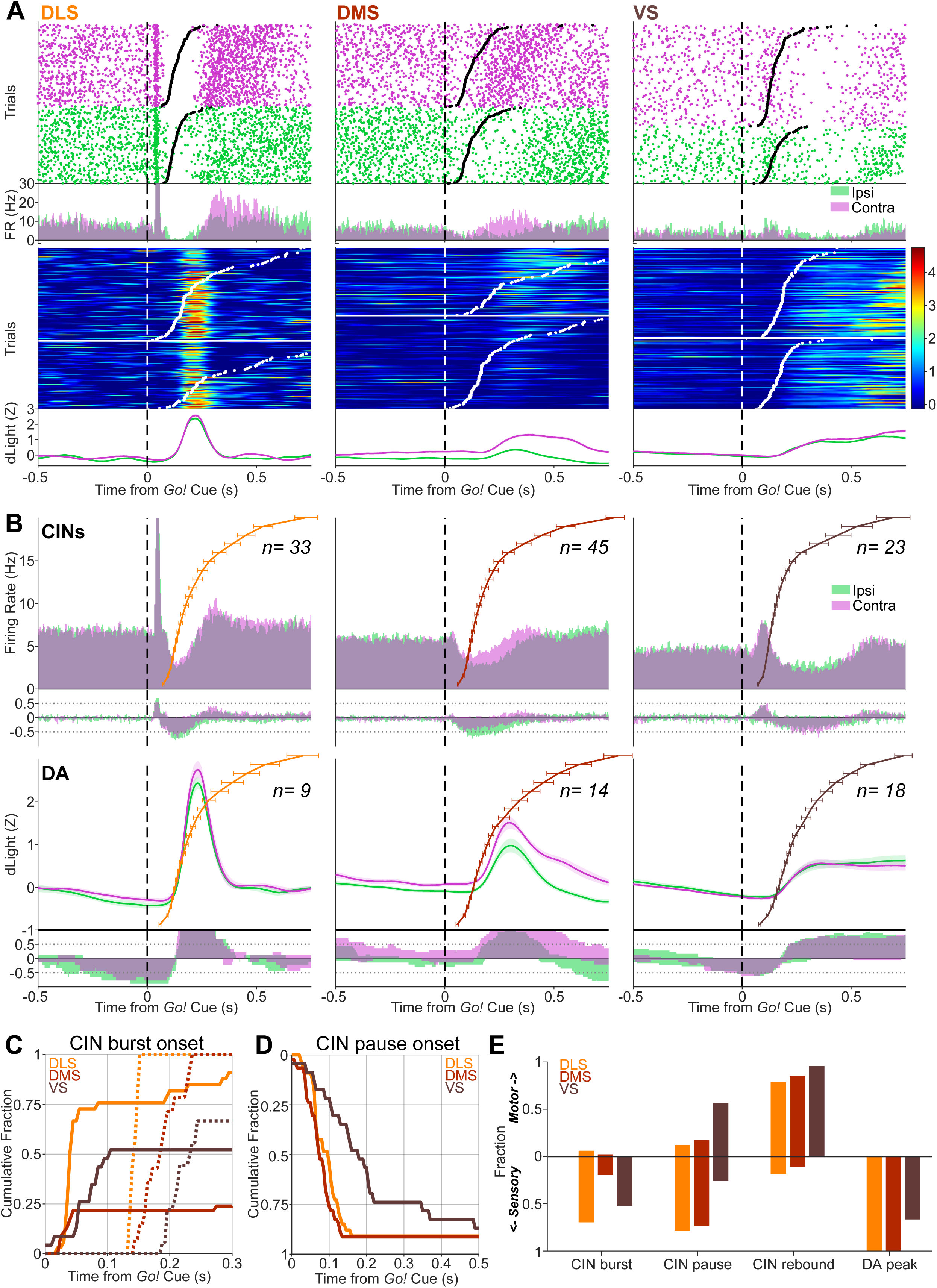
Sensory and motor correlates of CIN firing and DA release. **A.** Individual examples of *Go!* cue-aligned CIN firing (top panels) and DA release (bottom panels) in each subregion. CIN firing is shown as a raster plot (above) and peri-event histogram (below), in each case separated into trials with contraversive (magenta) and ipsiversive (green) choices. Trials in the raster plot are sorted by reaction time (RT), with black ticks indicating time of detected movement onset (Center Out). Color plots show Z-scored DA release in each trial (from different example sessions), with contraversive choice trials in the upper panel and ipsiversive choice trials in the lower panel. Trials are again sorted by RT, with white tick marks indicating movement onset. Bottom, averaged DA responses for those sessions. **B.** Top, *Go!-*aligned firing rate histograms, averaged across all identified CINs (contraversive, magenta; ipsiversive, green). Colored lines show overall RT distributions, averaged across the corresponding recording sessions (shown as group quantiles; error bars show SEM for each quantile; Ratcliff 1979). Next row shows fractions of CINs with significant firing rate increases (up) and decreases (down) at each time point (shuffle test, p<0.005, 10,000 shuffles, multiple comparisons corrected). For the same data aligned on movement onset, see Supp. Fig. 4A. Bottom two rows show DA release data, in the same format. **C.** Timing of *Go!-*evoked CIN bursts and DA transients. Cumulative distributions of the first time bin with a significant increase (shuffle test, 10,000 shuffles, p < 0.005, corrected for multiple comparisons) following the *Go!* cue (CIN firing, solid lines; DA release, dashed lines). **D.** As C, but showing timing of pause onset (first significant decrease). DA is not shown, as no DA decreases were observed in this time range. For CIN rebound timing, see Supp. Fig. 4B. **E.** Assessment of whether neural signals are more closely related to the *Go!* cue or the subsequent movement. For each CIN and each DA recording we aligned the signal to the *Go!* cue, and to Center Out, and compared the relative amplitude of changes from baseline (Methods). Bars indicate fractions of signals that were stronger when cue-aligned (“Sensory”, downwards) or movement-aligned (“Motor”, upwards), respectively.

In DLS the initial CIN burst was consistent and fast (onsets in the 30-50ms range; Fig 3C). This burst was locked to Go cue onset (Fig. 3E) - it did not differ depending on the subsequent movement timing or direction (Supp. Fig. 4A). The DLS pause was similarly Go cue-locked and direction-insensitive (Supp Fig. 4A). This pause slightly preceded a pulse in DLS DA release (Fig. 3A, B, lower panels), which was also rapid, brief, cue-locked, and direction-insensitive. The highly trained rats typically responded to the *Go!* cue with short reaction times (RTs; from *Go!* cue onset until Center-Out; median RTs: 169 ms for sessions with DLS tagged CINs; 139 ms for DMS; 155 ms for VS; Supp. Fig. 5). As a result, movement onset was typically detected during, or even before, the DLS CIN pause and DA pulse (Fig. 3A,B). Following the pause CINs typically increased firing. However, this was not a true “rebound” response; unlike the cue-locked pause component, the timing of the increase was instead locked to the movement (Fig. 3E) and was often dependent on movement direction (Supp. Fig. 4C).

Activity in DMS around the *Go!* cue was strongly dependent on movement direction. As noted above, DMS CINs generally lacked an initial burst phase, and possibly as a result the DMS pause onset was detectable slightly though not significantly earlier than in DLS (Fig. 3B; median onset times, DMS 80 ms, DLS 105 ms; KS test of distributions, p=0.217). This pause was more prominent on trials when rats made ipsiversive, compared to contraversive, movements. However, this does not appear to reflect a particular relevance of pauses to ipsiversive trials. Rather, this difference arises from the a more robust DMS CIN “rebound” increase on contra trials, that begins earlier (median onset times, DMS contra: 185 ms, ipsi: 322.5 ms, KS test of distributions, p=0.0011) and curtails the CIN pause (Supp. Fig. 4A). This contra-preferring rebound in DMS CIN firing roughly coincided with a contra-preferring increase in DMS DA release, although this DA increase was instead cue-locked (Fig. 3E). DMS DA (and to a lesser extent DLS DA) distinguished movement direction even before the *Go!* cue, potentially reflecting a state of movement bias or preparation (Supp. Fig. 4C); this movement-selectivity was not apparent for VS DA or CINs.

In VS both CIN firing and DA release dynamics were visibly slower (Fig. 3A-B). As in DLS, VS CINs showed burst firing after *Go!* cue, but with delayed onset (medians for responsive CINs, DLS 45 ms, VS 110 ms; KS test p = 2.02 x 10^−8^) and longer duration. Unlike DMS there was little indication of direction-specificity in either VS signal (Supp. Fig. 4A). As in the other regions, VS DA increases were cue-locked, rather than movement-locked (Fig. 4E).

**Figure 4.**
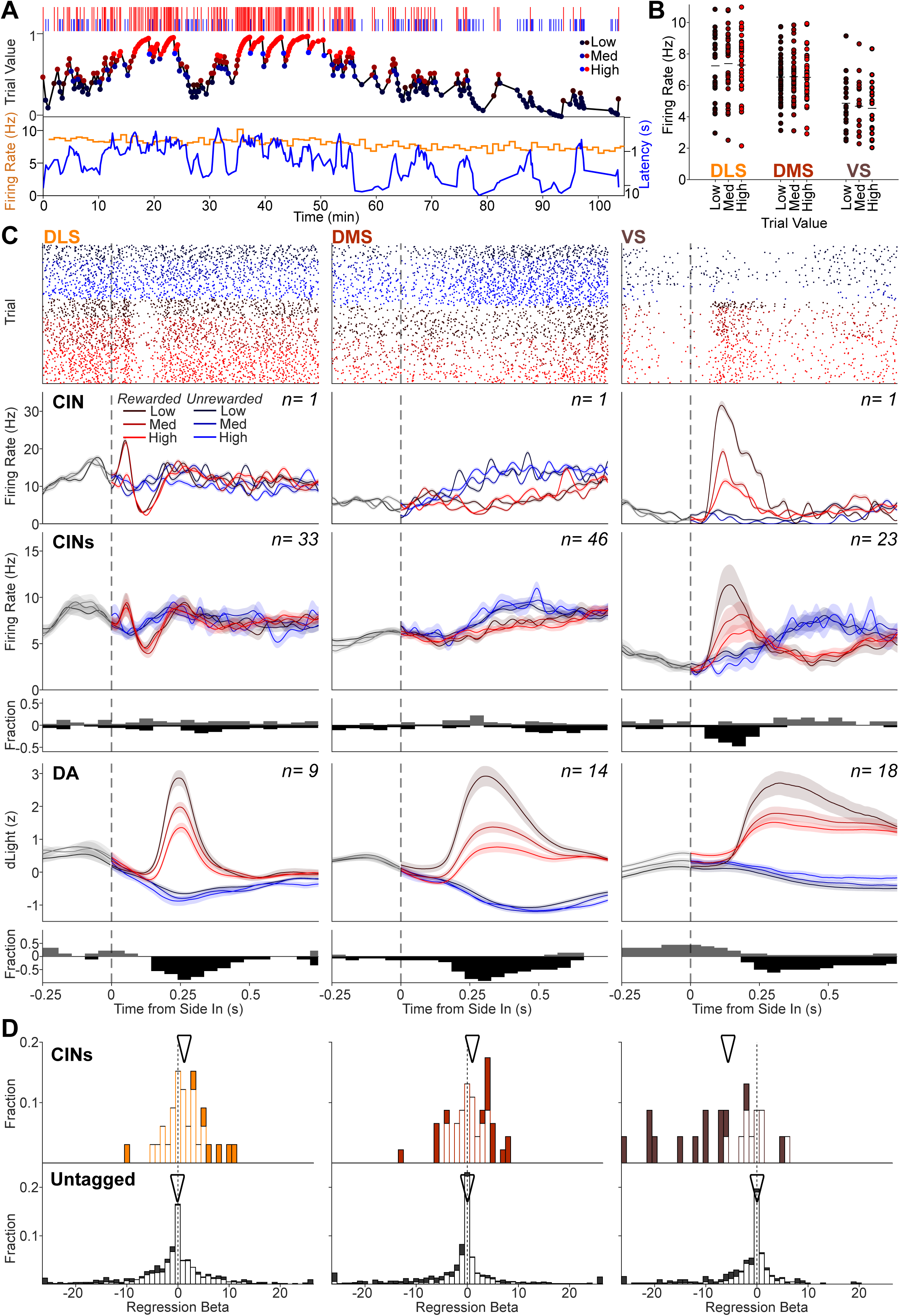
Reward prediction error encoding specifically in VS CINs. **A.** Top, Outcome sequence in a single example session. Tall red ticks represent rewarded trials, short blue ticks unrewarded trials. Middle, “trial value” estimated with a Bayesian model; see Methods). Values for all trials were divided into terciles, illustrated by color. Bottom, average firing rate (in one minute bins) of an example CIN recorded in the same session. Blue: latency to Center Nose In for each trial. **B.** Average firing rates for each CIN during the inter-trial interval (−3 to 0s before Light On) for trials with low, medium (med) and high trial value. **C.** Relationships of reward-cue-evoked CIN firing and DA release to reward expectation. Top row, raster plots of single example CINs in each subregion. Trials are divided according by rewarded (red) and unrewarded (blue), and further subdivided by trial value (low, medium and high), shown by the color shade, the band shows SEM. Next row, corresponding peri-event histograms. Next row, averaged histograms for all identified CINs. Next row, fractions of CINs with significant value encoding at each time point (p < 0.05, simple regression, multiple comparisons corrected; positive regression coefficients are shown upwards in gray and negative coefficients downward in black). Next row, averaged Z-scored DA release, by outcome and trial value. Below, the fraction of recorded fiber placements with significant value coding. Format for DA data is the same as for CIN data. **D.** Distributions of trial value regression coefficients for CIN interneurons (top) and unidentified neurons (bottom) in each striatal subregion. Analyzed epoch was 100-250 ms after Side In on rewarded trials; for other time epochs see Supp. Fig. 6C. Significant coefficients (p < 0.05, two-tailed, F-test) are shown in filled bars, non-significant in empty bars. Arrowheads indicate medians for each population. For CINs, mean regression coefficients and proportion significant (positive or negative): DLS: 1.38 n=7/33, DMS=0.68 n=16/45 and VS: −7.14 n=13/23; for unidentified neurons, DLS: −0.81, n=173/897; DMS: −1.35, n=237/965; VS: −1.93, n=94/448.

Overall, both CIN firing and DA release respond to a salient sensory cue, which prompts movement but does not provide information about whether the trial will be rewarded. CINs responses are multiphasic; the initial burst and pause responses are sensory locked and precede DA release, while a subsequent increase activity is movement-related and follows DA release.

### VS CIN firing and DA release both show RPE-scaled increases after a reward cue

We next assessed how CIN firing is affected by reward expectation. Rats show higher motivation to perform the bandit task when more recent trials have been rewarded, as seen in their latency to initiate trials (Fig. 4A). Using microdialysis we previously observed that this higher reward expectation is associated with greater DA release in VS (nucleus accumbens Core), and not in some other subregions including DMS and accumbens Shell (Hamid et al., 2016; Mohebi et al., 2019). This aspect of DA release was not mirrored in the tonic firing of identified lateral VTA DA neurons (Mohebi et al. 2019). However, prolonged states of reward expectation can be reflected in the tonic firing of other cell populations, including in lateral habenula (Post et al., 2022) and dorsal raphe (Cohen et al., 2015). We therefore examined whether reward expectation changes the tonic firing of CINs, as this could conceivably be responsible for the relationship between VS DA and reward rate. On slower timescales (1 min bins) the firing rate of CINs within sessions appeared stable (Fig. 4A bottom; Supplementary File), with no significant difference between blocks with higher vs lower reward expectation (Supp. Fig. 6A; Wilcoxon rank test, p = 0.79). In a further analysis we measured the average firing rate of each CIN during the 3 s interval immediately before Light On in each trial. No effect of reward history (assessed as “trial value”, V; see Methods) was apparent in any subregion (Fig. 4B). Two-way ANOVA showed no effect of trial value on average CIN firing during this epoch (factors of REGION and VALUE yielded a main effect of region, F = 50.31, p = 1.62×10^−19^, but no main effect of VALUE, F = 0.05, p = 0.955, and no interaction, F = 0.995).

We previously found that the audible delivery of food pellets at Side In evokes RPE-scaled increases in VTA DA cell firing (Mohebi *et al*. 2019) and RPE-scaled DA release across DLS, DMS and VS (Mohebi *et al*. 2024). In monkeys Morris et al. (2004) found TANs pausing after the same cues that evoked RPE-coding DA cell firing, without the TANs themselves encoding RPE (Morris et al., 2004). However, in other studies greater RPE has been associated with longer TAN pauses (Joshua et al., 2008; Apicella et al., 2009) or larger rebounds (Apicella et al., 2011).

We therefore assessed whether, and how, CINs encode RPE in the bandit task. After the *Go!* cue neither changes in CIN firing nor the pulse of DA release showed modulation by reward history (Supp. Fig. 6B), consistent with a lack of RPE coding at this time. After the reward cue at Side In, we found that VS CINs responded with a strong increase in firing (Fig. 4C, right), rather than a pause. This increase scaled with RPE, in a similar manner to VS DA release (albeit faster). No comparable CIN firing increase was seen in DLS or DMS (Fig. 4C, left and center), even though the reward cue evoked RPE-scaled DA increases in all striatal subregions tested (Fig. 4C bottom; Mohebi et al. 2024). Rather, on average DLS CINs responded with a burst-pause sequence, similar to their *Go!* cue response but much less pronounced. DMS CINs instead showed a moderate, prolonged increase, selectively on trials in which the reward cue was omitted.

Regression analysis of individual neurons confirmed that a majority of CINs in VS, but not in DLS or DMS, encoded RPE after the reward cue (Fig. 4D, right; negative coefficients for value indicate greater activity when the reward is less expected, i.e. RPE). This selective RPE coding by VS CINs was not dependent on the specific choice of analysis window (Supp. Fig. 6C) or model of reward expectation (Supp. Fig. 7), and was much stronger and more consistent for CINs than for unidentified VS neurons (Fig. 4D; bottom). Overall, these results suggest a particular role for VS CIN firing in RPE processing.

## Discussion

In the original report on monkey TANs, Kimura et al. (1984) concluded: *“It is clear that a high priority in further work…will be the definitive neurochemical and anatomical identification of the tonic putamen neurons…”.* Such a definitive identification has been elusive (Apicella, 2017), despite the vast increase in knowledge about striatal cells and circuits during the last 40 years. From brain slices and anesthetized animals there is considerable indirect evidence that TANs correspond to CINs. However the present work is the first, to our knowledge, to conclusively demonstrate this relationship. In particular, in behaving rats identified DLS CINs show the tonic firing and the burst-pause-rebound response to cues characteristic of TANs.

Recordings from three distinct subregions allow us to assess which properties of CIN activity are shared across circuits, and which reflect subregion-specific inputs and local informational processing. In each subregion CINs fired tonically at relatively low rates, but this rate was highest in DLS and lowest in VS. This is consistent with a spatial gradient in the overall “tempo” of striatal activity, as observed in the pace of DA fluctuations (Mohebi et al., 2024) and the firing of presumed projection neurons (Ito & Doya 2015). In all subregions CINs fired more regularly than other neuronal populations and maintained their tonic firing during slow-wave sleep.

However, event-related activity of identified CINs clearly differed between subregions, so CINs do not provide a global cholinergic signal throughout striatum. Notably, the classic TAN burst-pause-rebound was preferentially seen in DLS (corresponding to the putamen in primates), in response to the *Go!* cue. Our results help resolve prior uncertainties about why patterns of TAN activity appeared distinct and more variable in rodents compared to monkey TANs. Rather than species differences, this appears to reflect the specific targeted subregions, together with prior challenges with cell identification.

Comparing CIN firing to DA fluctuations in the same subregions and behavioral task sheds light on their mutual interactions and relative timing. Critically, our results do not provide support for the most widely held current theory of CIN function: that pauses in tonic CIN firing coincide with RPE-coding DA release to jointly enable synaptic plasticity and thereby learning (Morris et al., 2004; Reynolds et al., 2022). There was no event time and striatal subregion in which we observed a clear coincidence of a CIN pause and an RPE-scaled DA pulse. Instead we found that in VS a task-critical reward cue was associated with joint, RPE-coding *increases* in both CIN firing and DA release, as discussed further below.

### Mechanisms shaping CIN responses to behaviorally significant cues

CIN firing patterns reflect a complex interplay between intrinsic properties and afferent inputs. While pacemaker currents produce autonomous spiking (Goldberg & Wilson, 2016) at relatively low rates, CIN membrane potentials remain close to spike threshold (Wilson et al., 1990). As a result, CINs can spike very quickly in response to excitatory inputs (Fino et al., 2008; Zhang et al., 2018), such as those associated with salient sensory cues. Among the wide range of inputs to CINs (Guo et al., 2015; Klug et al., 2018), several projections are known to be sensitive to cues and may drive the fast Go-cue-evoked CIN burst we observed in DLS. The intralaminar thalamic complex (ILT; notably the parafasicular nucleus) contains neurons with rapid responses to surprising cues, and their projection to CINs may promote attentive responses to those cues (Yamanaka et al., 2018). However, the ILT areas with fast cue responses predominantly project to associative striatum (DMS / caudate) - where we rarely saw a fast CIN burst - rather than sensorimotor striatum (DLS / putamen; Sadikot & Rymar, 2009). Furthermore, inactivation of ILT interferes with CIN pauses and rebound, but the initial burst response often remains (Matsumoto et al., 2001). A more likely driver of DLS CIN bursting after the *Go!* cue is the pedunculopontine nucleus (PPN). PPN neurons can respond very rapidly to salient auditory stimuli (Pan & Hyland, 2005), and PPN glutamatergic projections to striatum preferentially contact interneurons (Assous et al., 2019). PPN also predominantly targets dorsal and lateral striatal subregions (Dautan et al., 2014), potentially accounting for the greater appearance of Go-evoked bursting in DLS CINs compared to DMS.

A brief pulse of excitatory input to CINs can be sufficient to trigger a triphasic response - burst, pause, and rebound (Doig et al., 2014). Excitation evokes a range of slowly-activating potassium conductances (Tubert & Murer, 2020), which result in a pause in CIN spiking as the excitatory input recedes (Zhang et al., 2018). This mechanism can produce pauses even if the initial excitation is insufficient to cause spikes (Goldberg & Wilson, 2016), as may be the case in DMS after the *Go!* cue. Pauses can themselves produce rebounds in CIN firing, at least when pauses are artificially generated using optogenetics (English et al., 2011; Zucca et al., 2018). Nonetheless, our results indicate that in behaving animals the multiphasic CIN response to a salient cue results from multiple inputs with distinct timing. In particular, while the burst and pause are time-locked to cue onset, the subsequent “rebound” is instead time-locked to movements. Furthermore, especially in DMS the CIN rebound component is stronger for contraversive movements, and occurs early enough to truncate the pause. These observations support prior thinking that the rebound reflects excitatory input distinct from that producing the initial burst and/or pause components (Schulz & Reynolds, 2013). As the rebound occurs in conjunction with the left/right movement, it may be driven by corticostriatal input involved in processes such as action selection (Klaus et al., 2019), efference copy (Redgrave & Gurney, 2006) and performance monitoring (Choi et al., 2023).

An influential early observation was that TAN pauses are abolished when DA axons are lesioned (Aosaki et al., 1994). This has been interpreted as a “permissive” role for DA (Schulz & Reynolds, 2013) - that is, DA simply needs to be present - but it has also been argued that cue-evoked DA release drives the CIN pauses (Chuhma et al., 2014; Straub, et al., 2014). In dorsal striatal brain slices, CIN pauses can be produced by local electrical stimulation (Kharkwal at al., 2016), or by stimulation of incoming thalamic axons (Ding et al., 2010) or nigral DA axons (Straub et al., 2014; Chuhma et al., 2018; Cai & Ford, 2018). These artificially evoked CIN pauses are dependent on DA, and D2 receptors on CINs (Kharkwal et al., 2016). Furthermore, co-release of glutamate from DA axons can produce fast excitation of CINs, preferentially in DLS compared to DMS (Cai & Ford, 2018) - an apparent match to our observation of fast Go-evoked CIN bursts in DLS. However, the idea that DA transients drive CIN pauses (or burst-pause sequences in DLS) is not supported by the time course of DA release we observed after the Go cue. In both DLS and DMS, the pause is maximal at a time (~125ms) before any detected increase in DA (Fig. 3). By the time of peak detected DA (~240ms) the pause is already over.

We cannot exclude the possibility that some DA transmission occurs before it is detectable with dLight1.3b photometry. However, our finding that cues evoke faster responses in CINs than in DA release is consistent with recordings of presumed SNc DA cells, which were found to respond to conditioned auditory cues with latencies starting around 50-65ms in rodents (Pan & Hyland, 2005; Pan et al., 2013), by which time the DLS CIN burst is completed and the DLS/DMS CIN pause has already begun. Cue-evoked DA, and co-released glutamate, is thus more likely to contribute to shaping CIN activity during the subsequent “rebound” phase (Wieland et al., 2014; Chuhma et al., 2018). Even at that later time, the impact on CIN firing is likely to be modulatory rather than a strong glutamate-driven excitation, since the DA Go response is time-locked to cues while the CIN rebound is instead movement-related.

### Functional impact of *Go!* cue-evoked CIN activity

Many components of striatal microcircuitry express cholinergic receptors, and changes in CIN firing will therefore have a broad, coordinated impact on striatal information processing. Before addressing potential consequences for synaptic plasticity and learning, we first consider how CINs may influence immediate performance of actions. Of note, our rats were highly trained in the bandit task and responded to the *Go!* cue very quickly (the median of session median RTs was 161 ms). As a result, of the various features of cue-evoked CIN firing only the DLS burst consistently preceded action initiation. This CIN burst likely diminishes the impact of excitatory inputs on DLS output, in at least two ways. First, muscarinic AChRs on afferent terminals reduce glutamate release onto SPNs (Dodt & Misgeld, 1986; Akaike et al., 1988; Sugita et al., 1991; Calabresi et al., 1998; Barral et al., 1999), on a time scale of 10s of ms (Pakhotin & Bracci, 2007). Second, increased CIN firing activates a variety of nearby GABAergic interneurons via nicotinic receptors (Dorst et al., 2020; Kocaturk et al., 2022), in turn inhibiting SPNs (De Rover et al., 2002; Sullivan et al., 2008; English et al., 2012; Luo et al., 2012). This combined suppressive action in response to unexpected cues may help interrupt or delay any ongoing action sequences (Schmidt & Berke, 2017), and also transiently shift behavioral control away from DLS towards circuits mediating more flexible behavior, including DMS (Balleine et al., 2009; Gremel & Costa 2013; Vandaele et al., 2019; Badreddine et al., 2022; Parrini et al., 2024).

Conversely, the subsequent CIN pause throughout dorsal striatum - even if brief - is thought to rapidly enhance excitatory inputs, and diminish GABAergic interneuron activity (Pakhotin & Bracci, 2007; Matityahu et al., 2022). The CIN pause may thereby make SPNs transiently more receptive to input, and - together with the DA increase - contribute to invigorating performance of actions that would otherwise be executed slowly (Mazzoni et al., 2007). Finally, the “rebound” in CIN firing that accompanies performance of actions has been suggested to help maintain action choices (Schulz & Reynolds, 2013), potentially through suppression of SPNs involved in competing behaviors. However, assigning functions to specific components of complex CIN firing patterns remains a challenge, given the many sites and mechanisms of cholinergic action in striatum (Tanimura et al., 2018; Morgenstern et al., 2022).

Also uncertain at the present time is how these CIN firing patterns contribute to nearby DA dynamics. Synchronous optogenetic activation of CINs can enhance DA release by acting at nicotinic receptors on DA axons (Threlfell et al., 2012; Cachope & Cheer, 2014), locally initiating action potentials (Liu et al., 2022). However, spontaneous DA dynamics in the dorsal striatum of head-fixed mice are not obviously diminished by genetic removal of DA axon nicotinic receptors (Krok et al., 2023). Furthermore, in our bandit task the external cues evoke spike bursts at midbrain DA neuron cell bodies (Mohebi et al., 2019), accounting for cue-evoked striatal DA release without requiring local striatal control. There is some recent evidence that CIN firing maintains nearby DA axons in a state of diminished responsiveness (Zhang et al., 2024). This could serve to enhance the temporal precision of DLS DA release transients. In one speculative scenario, the initial CIN burst would ensure that all DA axons are briefly release-incompetent, and desensitize nAChRs; the CIN pause would then allow DA axons to fully respond to the subsequent burst of action potentials arriving from the midbrain with enhanced DA release.

The fast DA response to the *Go!* cue is consistent with prior reports of short-latency DA firing to salient, movement-triggering stimuli with unpredictable timing (e.g. Schultz & Romo 1990; Pasquereau & Turner, 2015). This appears to reflect a fast “alerting” aspect of DA signals that is insensitive to reward predictions (Redgrave & Gurney 2006; Bromberg-Martin et al., 2010; Schultz, 2016), as we also found here, and thus distinct to the RPE-coding DA aspect. It is noteworthy that we found that strong CIN pauses coincide with this alerting - rather than the RPE-coding - DA signal. This fits well with prior observations that CINs pause following unexpected, movement-triggering stimuli, but lose this response if the stimuli become temporally predictable (Sardo at al., 2000; Ravel et al., 2001). Predictability may also explain why CIN pauses were small or absent in response to the reward cue at Side-In. The timing of this event is determined by the rat’s own movement: even though the probabilistic reward cue may or may not occur on a given trial, if it does its timing can be fully predicted. By pausing in conjunction with some DA transients, but not others, CINs might conceivably help striatal circuits appropriately distinguish DA signals that convey different meaning (Berke 2018).

### CINs and the control of striatal plasticity

The influence of CINs over nearby SPNs is also thought to be essential for normal corticostriatal plasticity (Calabresi et al., 1999; Wang et al., 2006; Deffains & Bergman 2015). In particular, long-term potentiation of synapses onto SPNs is believed to require near-simultaneous depolarization of the SPN, an increase in DA, and a pause in CIN firing (Reynolds et al. 2022). In our bandit task these conditions may be met in the dorsal striatum after the *Go!* cue. What type of learning would be supported by this plasticity is currently less clear. One relevant proposal is that (dorsal) striatal circuits combine fast event-evoked DA and efference copy from cortex to discover which actions may have caused unpredicted events (Redgrave & Gurney 2006). Dorsal striatal circuits have also been suggested to learn behavioral policies directly, without requiring explicit rewards or value-related signals (e.g. Miller et al., 2019; Bennett at al., 2021).

Nonetheless, both dorsal and ventral striatal subregions showed a RPE-scaled DA pulse in response to the reward cue. At this time VS CINs do not typically pause, but rather also show an RPE-scaled increase in firing. This surprising similarity to DA release may reflect a common source of input to VS CINs and VTA DA cells. As one notable example, individual cholinergic neurons in the LDT send branching projections to both VTA DA neurons, where they drive burst firing (Lodge & Grace, 2006), and to VS (Dautan et al., 2014), where they excite CINs (Dautan et al. 2020). LDT-VS projections are closely involved in both positive reinforcement and motivation (Coimbra et al., 2019), and it would be useful to confirm whether RPE information is already present in this branching projection from LDT. Regardless, the observation of increased VS CIN activity after rewards has precedents in the literature (Benhamou et al., 2014; Howe et al., 2019). For example, presumed CINs in rat VS were found to have prominent increases after reaching the reward port in a maze task (Atallah et al, 2014). These responses faded over extended training then re-emerged when the task contingencies changed, potentially reflecting RPE. RPE-scaled increased firing has also been reported in the “rebound” phase of TAN responses to reward cues (Apicella et al., 2011). In other contexts VS CINs appear to pause, similarly to dorsal striatal TANs (Marche et al., 2017). It will be important to further explore the factors, including but not limited to temporal predictability, that shape VS CIN responses.

Inevitably, our study has several limitations. We focused on the activity dynamics within a single behavioral task, we did not sample from all striatal subregions, and our comparison of CIN firing and DA release is based on separate rather than simultaneous recordings. Furthermore, our use of electrophysiology and fiber photometry provides an incomplete view of the potentially rich spatiotemporal dynamics of ACh release (Matityahu et al., 2023) and ACh:DA interactions (Liu et al. 2022). Nonetheless, our account of the subregion-specific firing of identified CINs in behaving animals, and their unexpected relationships to DA release, should serve as a strong foundation for future experimental and modeling studies.

## Supporting information

Supplementary File: Each identified neuron

## Acknowledgements

We thank Howard Fields, Dennis Burke, Elyssa Margolis, Alex Legaria and members of the Berke Lab for constructive comments on the manuscript, and Isabelle Gonzalez Montalvo for contributing data on striatal projection neurons. Mariana Duhne, is a Latin American Fellow in the Biomedical Sciences, supported by The Pew Charitable Trusts. This work was funded by the National Institute of Neurological Disorders and Stroke (R01NS123516), the National Institute on Drug Abuse (R01DA045783), the National Institute of Mental Health (K01MH126223), and the State of California.

## Author Contributions

Conceptualization: J.B. and M.D.; Investigation, Methodology, Software: M.D. and A.M; Data Curation and Analysis: M.D., A.M., L.P., K.K.; Visualization: M.D.; Writing – original draft: M.D. and J.B.; Writing - Review and Editing: J.B.; Funding Acquisition: J.B., M.D., A.M.; Supervision: J.B.

## Declaration of Interests

The authors declare no competing interests.

## Methods

### Animals

All animal procedures were approved by the University of California San Francisco Institutional Committee on use and care of Animals. Male rats (400-550 g, ChAT-Cre+ on a Long Evans background, 6-12 months old) were maintained on a reverse 12:12 light:dark cycle and tested during the dark phase. Rats were mildly food deprived, receiving 4g / 100g of body weight of standard laboratory chow in addition to food pellets received during task performance. Body weight was constantly monitored to stay between 85-90% of baseline. No sample size pre-calculations were performed.

### Behavior

Pretraining and testing were performed in computer-controlled Med Associates operant chambers (25 cm × 30 cm at widest point) each with a five-hole nose-poke wall, as previously described (Hamid et al., 2016: Mohebi et al., 2019, 2024). Bandit task sessions used the following parameters: block lengths were 35–45 trials, randomly selected for each block; hold period before Go cue was 500–1500 ms (uniform distribution); left/right reward probabilities were 10, 50, 90%. Rats were trained to complete 75% of trials without procedural errors before implant surgery.

### Electrophysiology

Rats (n=20) were bilaterally infused with 1μl of AAV5-Syn-FLEX-rc[ChrimsonR-tdTomato] into each striatal subregion (DLS: AP 0, ML ±4.0, DV 4.0mm; DMS: AP 1.6, ML ±2.0, DV 4.0mm; VS: AP 1.6, ML ±1.8, DV 6.5mm). During the same surgery we implanted custom designed assemblies, consisting of 2 sets of 16 tetrodes each (constructed from 12.5 μm nichrome wire, Sandvik, Palm Coast, FL) inserted into a polyimide tube which slides around a 200 μm tapered optic fiber (Doric Lenses). The tetrodes were initially placed 500 μm above the fiber tip. Additionally, 5 bone screws (Fine Science Tools, catalog # 19010) were placed in contact with the brain surface. Two of them recorded frontal ECoG from each hemisphere (AP 5, ML ±2), one was placed 1 mm posterior to bregma to serve as reference, and two were placed in the posterior lateral skull to serve as ground. During recording sessions, wideband (1–9000Hz) brain signals were sampled (30,000 samples/s) using a custom headstage with 2 x 64-channel Intan RHD2164 digital amplifier chips (Farries et al. 2023).

After bandit task performance a subset of the implanted rats also performed a Pavlovian approach task (Mohebi et al. 2019), in the same operant chamber with the house light on throughout the session. Three auditory cues (2, 5, 9kHz) were associated with different probabilities of sucrose pellet delivery (0, 25, 75%, counterbalanced across rats). Cues were played as a train of tone pips (100ms on / 50ms off) for a total duration of 2.6s followed by a delay period of 500ms. Cues, and unpredicted reward deliveries, were delivered in pseudorandom order with a variable inter-trial interval (15 - 30s, uniform distribution; results for each CIN are shown in the Supplementary File).

Rats were then left for 40-60 min in the recording chamber to record a sleep session, with white noise constantly playing at 45 dB. Finally, optic fibers were connected for light delivery and the laser stimulation session was recorded. Tetrodes were lowered by at least 80 μm between sessions to avoid repeated recording from the same units, up to 500 μm below the fiber tip or until no light responsive units were detected over multiple sessions. Eight rats in which we did not record any identified CINs were excluded from further analysis.

### Histology

To confirm expression of Chrimson in ChAT+ interneurons we performed immunohistochemical staining. After recordings were finished animals were anesthetized with isoflurane, then perfused with PBS 1X (Sigma-Aldrich P4417) solution followed by formaldehyde solution 4% (Sigma-Aldrich, F8775). 50-100 µm slices were blocked using a PBS solution containing 5% normal donkey serum and 0.4% Triton x-100 (Sigma-Aldrich, 93443), then incubated with primary goat anti-ChAT antibody (ab254118, abcam 1:1000), and mouse anti CD11b (MCA618R, BioRad 1:1000) antibodies followed by secondary donkey anti-goat antibody conjugated with Alexa fluor 647 (ab150135, Thermo Fisher Scientific, 1:500) and donkey anti-mouse antibody conjugated with Alexa Fluor 488 (catalog# A-212-2, RRID AB_141607, Thermo Fisher Scientific 1:500). To assess co-expression of ChAT and Chrimson, a total of 29 brain slices from 9 animals (6 of the recorded animals and 3 additional rats) were quantified. Each field was imaged using a Keyence BZ-X800 microscope with a 10x objective, selecting a field fully in striatum with obvious tdTomato expression. In each field the number of Chrimson+, ChAT+ and cells with colocalized expression was counted. Within the selected fields, overall infection efficiency was 70%: 240/343 ChAT+ neurons expressed tdTomato as well. This number includes areas of lower viral expression, further from the recording sites; nonetheless, it is likely that our untagged cell recordings include at least some CINs.

### Classification

To determine whether an isolated striatal unit was cholinergic (ChAT+), we evaluated response to light pulse trains delivered at the end of the session (Kvitsiani et al., 2013). Trains of different widths: 2, 5 and 10 ms and frequencies: 1, 2, 5 and 10 Hz were used. For a unit to be identified as light responsive it needed to fire with a spike latency after laser stimulation significantly shorter (Wilcoxon rank sum test, *p* < 0.01) than the spike latency following randomly selected times within the same session. We also required that spikes appear within 15 ms of laser onset, in at least 25% of trials for at least one stimulation condition. Finally, to address the possibility that laser-evoked spikes were fired by a different neuron and included with the analyzed cell due to a spike sorting error, we required that the waveforms of spikes occurring <10 ms following laser stimulation have a Pearson correlation coefficient >0.9 compared to the average prestimulus waveform (Farries et al., 2023). Peak width was defined as the full-width-at-half-maximum of the most prominent negative-voltage component of the averaged spike waveform. Peak-to-valley time was the interval between the time of this peak and the time of the most positive voltage after this peak, within the 2 ms total duration of the spike analysis window.

### Analysis

All data analyses were performed in MATLAB (Mathworks, Inc.; Natick, MA). Individual units were isolated offline using the MountainSort algorithm (Chung et al. 2017) followed by careful manual inspection. For slow wave sleep (SWS) detection we measured ECoG power in several frequency bands. To establish epochs of predominantly lower frequencies, we calculated the ratio of 0.5 – 8 Hz (r1) to 20 – 60 Hz (r2) power. To correct for occasional low frequency artifacts we subtracted from this the ratio of the power in the 0.5 – 4 Hz (r3) and 8 – 12 Hz (r4) bands. The resulting measure k = power(r1) / power(r2) – power(r3) / power(r4) was used to detect SWS. Whenever k exceeded the threshold 3, we labeled this as a putative SWS epoch. We excluded putative SWS epochs that were < 30 s in duration (Mallet et al., 2016).

Post-spike suppression was calculated as the time when the value in the autocorrelogram (1 ms bins) first exceeds half of its mean value (Schmitzer-Tobert & Redish, 2008; Thorn & Graybiel, 2014).

We characterized the response of units to events by building peri-event time histograms (PETHs). For all events we analyzed a 3 s window in 10 ms overlapping (5 ms) time bins. To determine significant modulation in all PETHs we used shuffle tests (Mohebi et al., 2019; Faust et al., 2023). Briefly we compared the firing rate of the neuron’s PETH in each time point to a shuffled distribution of 10,000 samples selected randomly from the same session. We used a threshold of p<0.005, after multiple comparisons correction. To assess movement direction selectivity (contraversive vs. ipsiversive) we calculated a Selectivity Index, defined as the difference of the mean PETH magnitude of contralateral and the ipsilateral choice trials divided by the sum of both (Gage et al., 2010).

To determine whether activity was more sensory-related or movement-related, we calculated the ratio of the maximum in each unit’s average response when aligned to Center Out vs. Go Cue events (for pauses, we used the minimum instead). We included only CINs with significant (shuffle test as described above) positive (burst and rebound) or negative (pause) modulation. The time windows used for significance testing were: Go cue aligned, burst 0-120 ms, pause 0-400 ms, rebound 200-500 ms; Center out aligned: burst: −400—150 ms, pause: 200-200 ms and 0-400 ms rebound. These windows were chosen based on the onset times for modulation on each event (Fig 3B-D, Supp. Fig. 4B-C). When the ratio was bigger than one activity was considered movement-related, and when it was smaller than one it was considered sensory-related.

### Photometry

DA fiber photometry procedures and recording locations have been previously described in detail (Mohebi et al. 2024). We used a viral approach to express the genetically encoded optical sensor dLight 1.3b. Under isoflurane anesthesia, 1µL of AAV9-CAG-dLight (1×10^12^ vg/mL - UC Davis vector core) was slowly (100nL/min) injected (Nanoject III, Drummond, Broomall, PA) through a 30μm glass micropipette in the different striatal subregions (DLS: AP 0, ML 4.0, DV 4.0, DMS: AP 1.6, ML 1.8, DV 4.0, VS: AP 1.6, ML 1.6, DV 6.5). During the same surgery optical fibers (200μm core, 250μm total diameter) attached to a metal ferrule (Doric) were inserted (target depth 200μm higher than virus) and cemented in place. Data were collected >3 weeks later, to allow for dLight expression.

For dLight excitation blue (470nm) and violet (405nm; control) light emitting diodes were switched in 10 ms frames (4 ms on and 6 ms off). Both excitation and emission signals passed through minicube filters (Doric) and bulk fluorescence was measured with a femtowatt detector (Newport, Model 2151) sampling at 10KHz. Time-division multiplexing produced separate 470nm (dopamine) and 405nm (control) signals, which were then rescaled to each other via a least-square fit. Fractional fluorescence signal (dF/F) was then defined as (470–405_fit)/405_fit. For all analyses this signal was downsampled to 250 Hz, smoothed with a 5-point median filter and z-scored.

### Modeling

A Bayesian model (Funamizu et al., 2012) estimated value in each trial (V) as a beta distribution (Bishop,2006). The distribution has two parameters: α which increases with each rewarded outcome and β which increases with each unrewarded outcome. There is one free parameter gamma that determines the decay rate of both α and β. For all analyses, gamma was selected based on the rat’s behavior, maximizing the (negative) correlation between reward rate and log (latency to Center In) in each session. The correlations between DA release or CIN firing with trial value were not highly sensitive to this choice of gamma (Supp. Fig. 6 D). Additionally, estimating reward expectation with other models gave similar results (Supp. Fig. 7).

## Notes

### Competing Interest Statement

The authors have declared no competing interest.

