## Supplementary Figures 1-7 for "A mismatch between striatal cholinergic pauses and dopaminergic reward prediction errors"

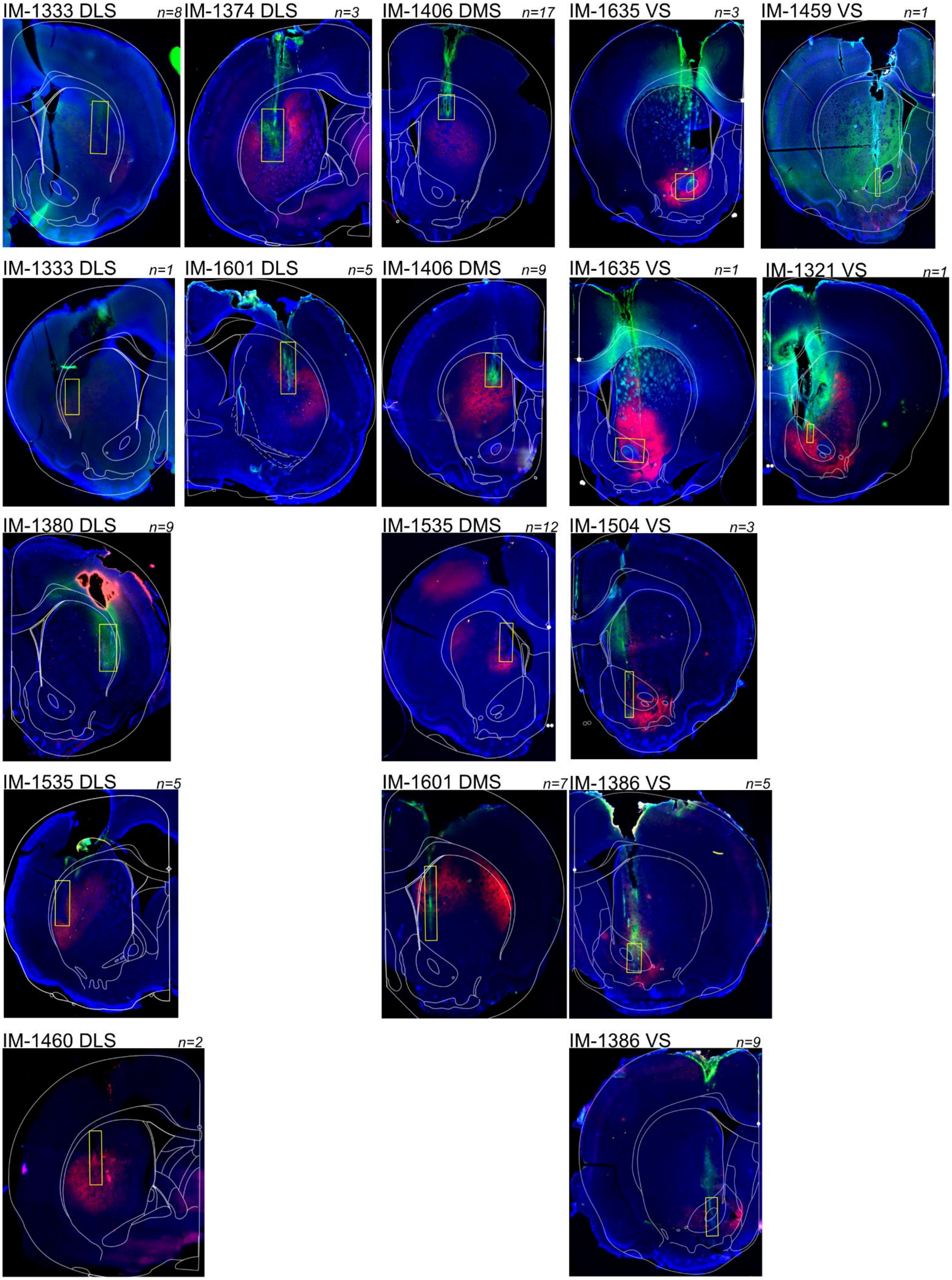
IM-1333 DLS n=8
IM-1374 DLS n=3
IM-1406 DMS n=17
IM-1635 VS n=3
IM-1459 VS n=1
IM-1333 DLS n=1
IM-1601 DLS n=5
IM-1406 DMS n=9
IM-1635 VS n=1
IM-1321 VS n=1
IM-1380 DLS n=9
IM-1535 DMS n=12
IM-1504 VS n=3
IM-1535 DLS n=5
IM-1601 DMS n=7
IM-1386 VS n=5
IM-1460 DLS n=2
IM-1386 VS n=9

**Supplementary Figure 1. Recording locations.**

Photos show histology images for each ChAT-Cre rat ("IM-XXXX") with tagged neurons. n = the number of tagged CINs isolated from each implant (in some rats we obtained CINs from both hemispheres, shown separately here). The ‘cage’ made with recording tetrodes has a 700 µm diameter before implant. The range of recording sites was estimated from tissue damage and expression of CD11b (green), a marker for microglial activation. For rat IM-1459 instead of CD11b we stained for calbindin, to better visualize the boundary between accumbens Core and Shell.

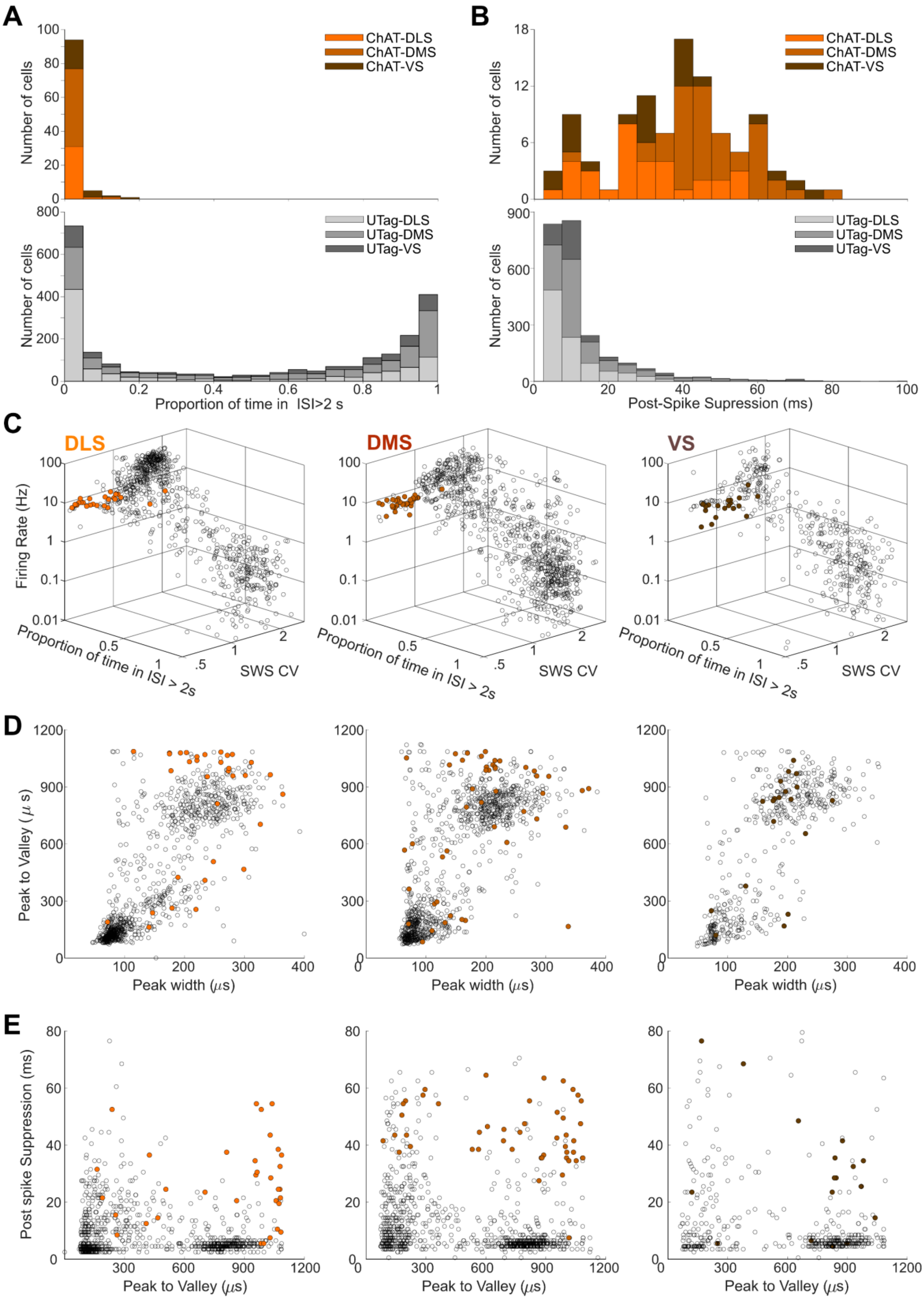

A
Number of cells
ChAT-DLS
ChAT-DMS
ChAT-VS
UTag-DLS
UTag-DMS
UTag-VS
Proportion of time in ISI>2 s
B
Post-Spike Supression (ms)
C
DLS
DMS
VS
Firing Rate (Hz)
Proportion of time in ISI > 2s
SWS CV
D
Peak to Valley (μs)
Peak width (μs)
E
Post spike Suppression (ms)
Peak to Valley (μs)

**Supplementary Figure 2. Firing pattern and waveform features are insufficient to identify ChAT+ interneurons.**

**A.** Proportions of time spent not firing (interspike interval > 2s) for CINs (top) and unidentified units ("UTag", bottom) in each subregion. All CINs were tonically active.

**B.** Post-spike suppression for CINs (top) and unidentified units (bottom).

**C.** Scatter plots of: CV during slow wave sleep (SWS) vs. mean firing rate vs proportion of time in ISI> 2s. Filled colored dots are tagged CINs, empty circles are unidentified units recorded during the same sessions.

**D.** Waveform feature durations: peak-width-at-half-max vs. peak-to-valley time.

**E.** Scatter plots of peak-to-valley time and post-spike suppression for each subregion.

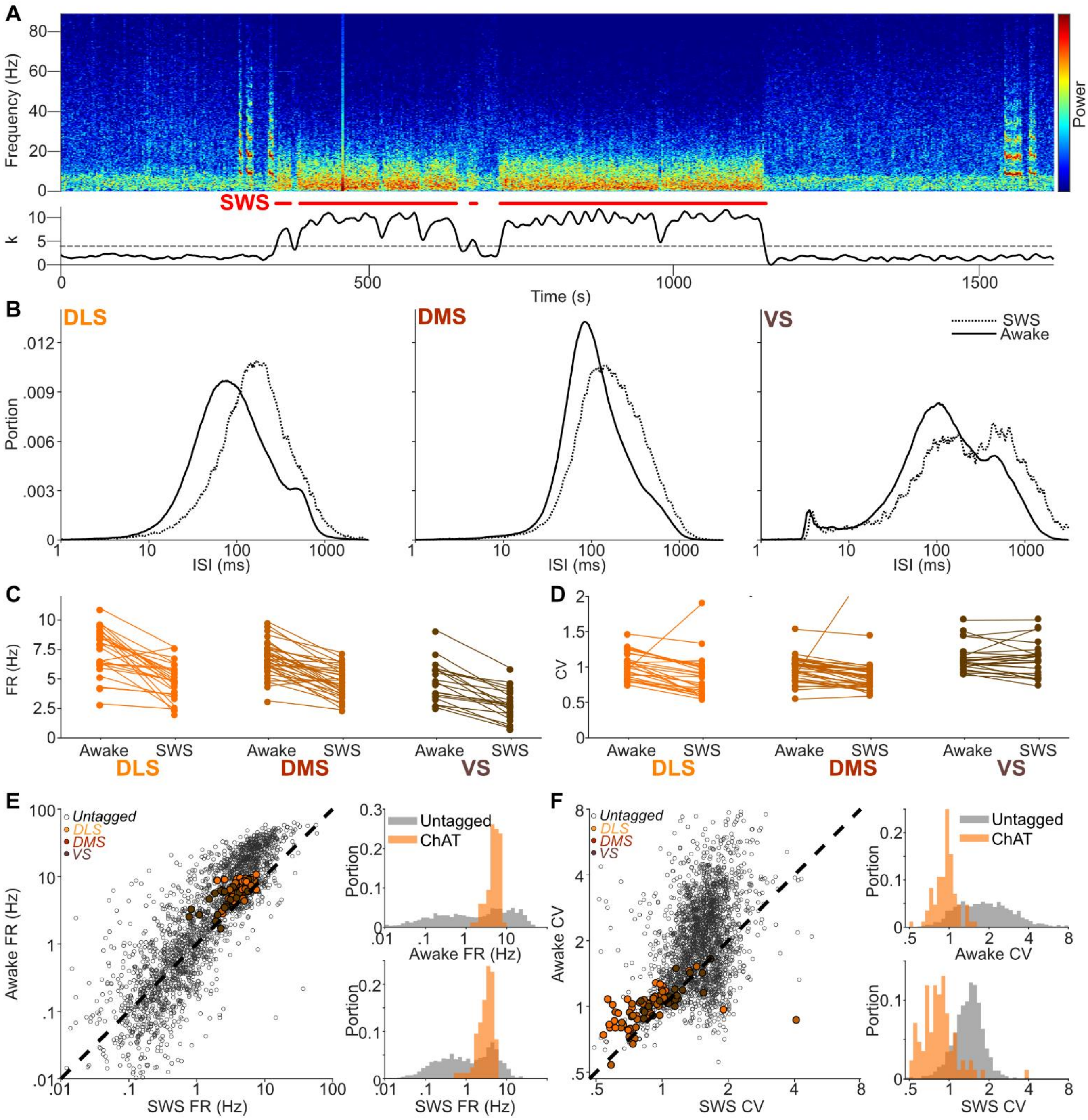


**Supplementary Figure 3. CIN activity during wake and slow-wave-sleep states.**

**A.** Top: Example spectrogram showing how ECoG power is used to distinguish states. Bottom shows the “k” power-ratio metric (see Methods).

**B.** Inter-spike-interval (ISI) histograms averaged across for all tagged CINs in each subregion, comparing wake (bandit task performance) to SWS.

**C.** Firing rates (FR) for individual CINs during RL task performance and SWS. Two-way ANOVA, with factors SUBREGION and STATE gives significant effects for each separately (SUBREGION: F=30.98, p=4.06 x $10^{-12}$; STATE: F=79.66, p=9.19 x $10^{-16}$) but no interaction (F=1.03).

**D.** CIN ISI coefficient of variation (CV) varies by subregion, but is unaffected by wakefulness.Two-way ANOVA, SUBREGION: F=4.28, p=0.015, STATE: F=2.17, p=0.14; no interaction (F=0.91).

**E.** Left, Scatter plot of FR during awake and SWS epochs, with CINs in colored circles and unidentified units in empty circles. Right, same data in histogram form. There was a significant change in FR distribution for both CINs (kstest, $p=2.15 \times 10^{-10}$) and unidentified units (KS test, $p=8.61 \times 10^{-46}$).

**F.** As E, but for CV. CIN CV was reduced moderately during SWS (CV mean 0.96, range 0.543-4.09; 1-way ANOVA, $F=1.96$, $p=0.16$; KS test p=2.35x10-4; Sharott et al. 2012). There is a significant change in CV distribution for both CINs (KS test, $p=2.35 \times 10^{-4}$) and unidentified units (KS test, $p=2.34 \times 10^{-143}$).

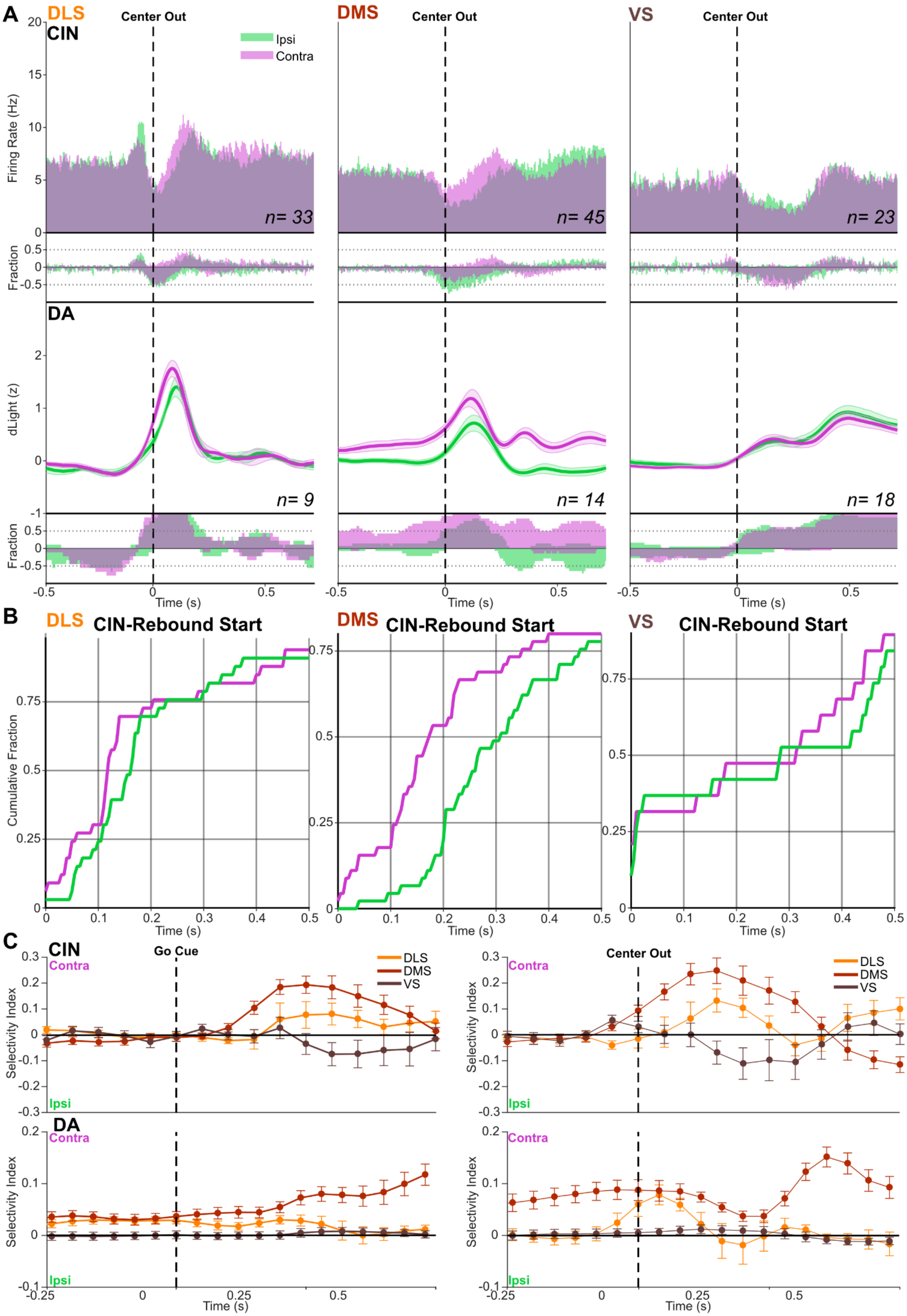

A
DLS
CIN
Center Out
Ipsi
Contra
Firing Rate (Hz)
n= 33
DMS
n= 45
VS
n= 23
Fraction
DA
dLight (z)
n= 9
n= 14
n= 18
Time (s)
B
DLS CIN-Rebound Start
DMS CIN-Rebound Start
VS CIN-Rebound Start
Cumulative Fraction
Time (s)
C
CIN
Go Cue
Contra
Ipsi
DLS
DMS
VS
Selectivity Index
Center Out
DA
Time (s)

**Supplementary Figure 4. Further analysis of movement-related activity in CIN and DA signals.**

**A.** Same as Fig. 3B, but aligned on Center Out (detected movement onset).

**B.** Timing of CIN "rebounds". CDFs of onset of significant increases in firing (shuffle test, $p<0.005$, 10,000 shuffles, corrected for multiple comparisons) after Center Out, separated by movement direction. For DMS the onset for contraversive choice trials begins significantly before ipsiversive trials (median onset times, DMS contraversive: 185 ms, ipsiversive: 322.5 ms, KS test of distributions, $p=0.0011$. For DLS this pattern did not reach significance (contraversive: 125 ms, ipsiversive: 160 ms; KS test of distributions, $p=0.0773$).

**C.** Selectivity Index for CIN activity and DA release aligned to Go Cue (left) and Center Out (right) for each subregion. Dots show mean ± S.E.M. for all units and sessions (CINs: DLS n=33; DMS n=45; VS n=23; DA: DLS n=9; DMS n=14; VS n=18). DLS and DMS CIN firing is stronger on contraversive trials, after Center Out. DLS and DMS DA release is stronger on contraversive trials even before the *Go!* Cue.

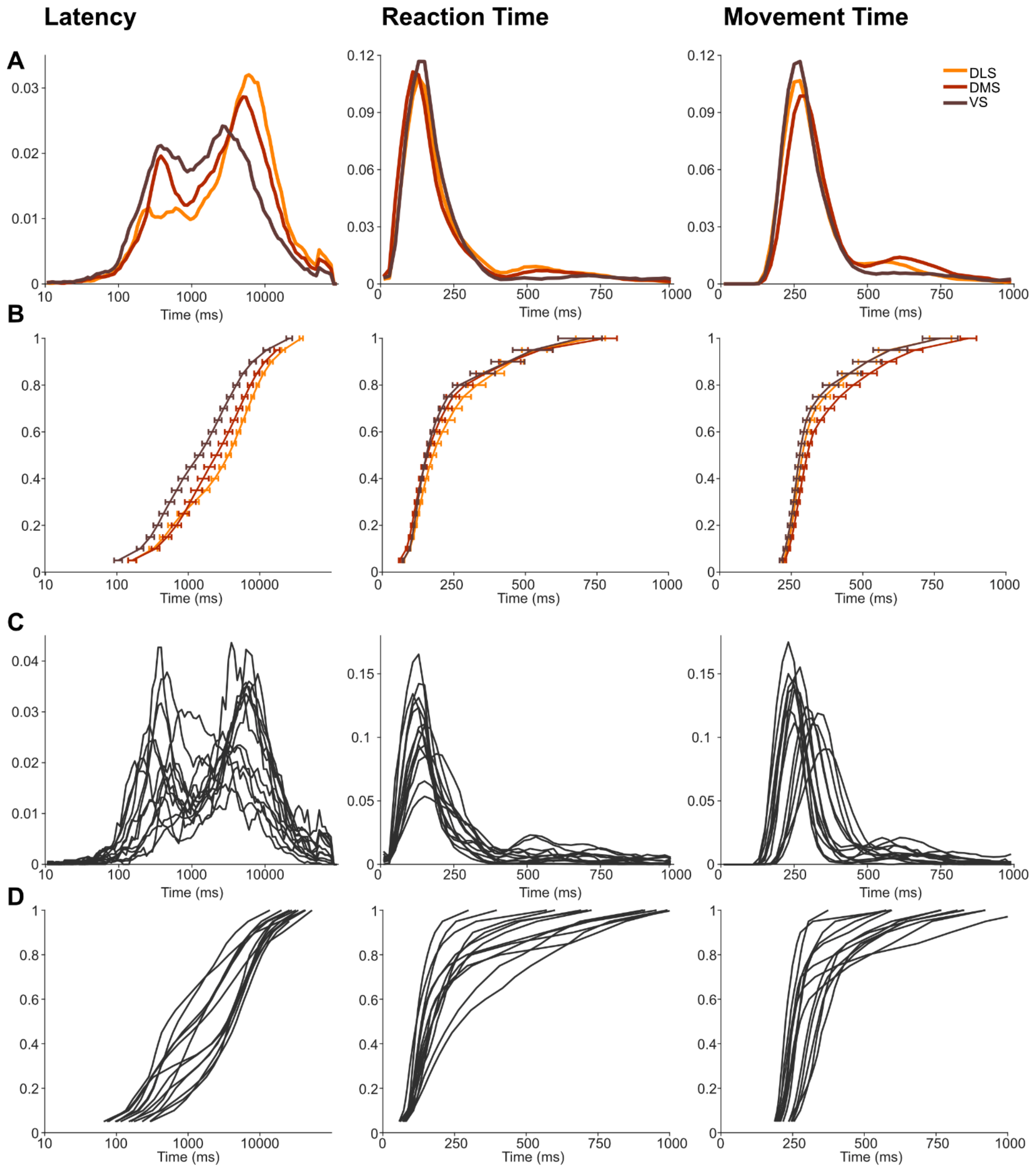


**Supplementary Figure 5. Distributions of behavioral variables.**

**A**. Latency, reaction time (RT) and movement time (MT) distributions aggregating all trials across all sessions for each subregion.

**B**. Same data shown as cumulative RT distributions (Vincentized; Ratcliff, 1979). Error bars = S.E.M.

**C, D.** As A,B but for each individual rat (averaging across sessions in which tagged CINs were recorded).

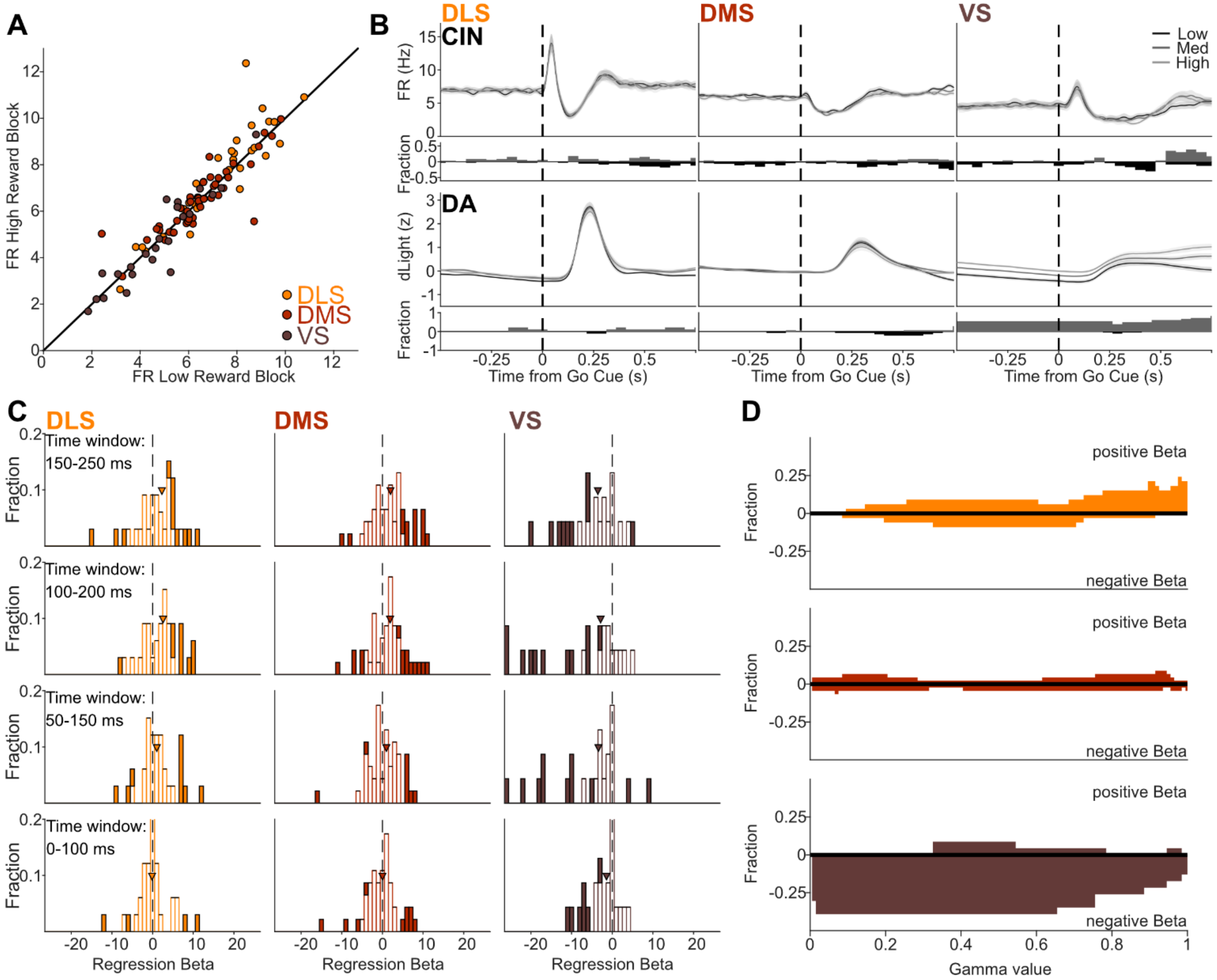


**Supplementary Figure 6. Further analyses of the relationships between reward expectation, CIN firing, and DA release.**

**A.** Trial blocks were median-split into higher and lower average trial value. These showed no significant difference in overall mean CIN firing rate (1 min bins; Wilcoxon rank test p=0.79).

**B.** Average CIN firing and DA dLight signal around the *Go!* cue, subdivided by trial value terciles (low, medium and high); error band shows SEM. The lower panels show the fraction of sessions or units with significant positive or negative regression coefficients to trial value per time bin ($p < 0.05$, simple regression, multiple comparisons corrected; positive regression coefficients are shown upwards in gray and negative coefficients downward in black).

**C**. Effect of choice of window on the distribution of regression betas of CINs per subregion. Significant coefficients ($p<0.05$, F-test) are shown in filled bars and non-significant in empty bars. Arrowheads indicate medians for each population.

**D**. Effect of choice of Bayesian model decay parameter gamma upon the fraction of units per subregion with significant positive or negative regression relationships to trial value ($p < 0.05$, simple regression, multiple comparisons corrected).

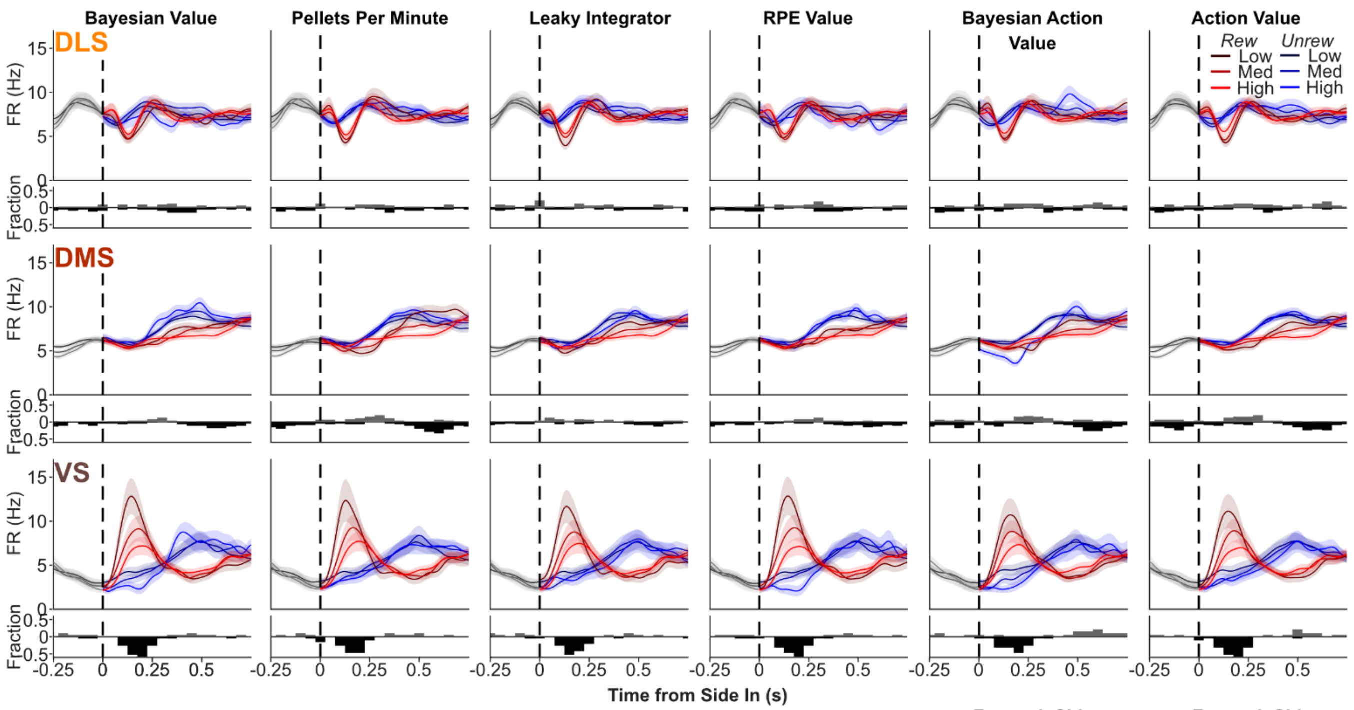


$$a_i = \gamma a_{(i-1)} + r_i$$
$$b_i = \gamma b_{(i-1)} + 1 - r_i$$
$$V = \frac{a_i}{a_i + b_i}$$
$$V_{STD} = \frac{a_i B_i}{(a_i + b_i)^2 + (a_i + b_i + 1)}$$

$$RR_i = RR_{(i-1)} e^{-\frac{1}{\tau}} + r_i$$

$$RPE = r_i - V_{(i-1)}$$
$$V_i = V_{(i-1)} + \alpha RPE$$

**For each Side:**

$$a_i = \gamma a_{(i-1)} + r_i$$
$$b_i = \gamma b_{(i-1)} + 1 - r_i$$
$$Q = \frac{a_i}{a_i + b_i}$$
$$Q_{STD} = \frac{a_i b_i}{(a_i + b_i)^2 + (a_i + b_i + 1)}$$
$$\beta_i = \frac{k}{Q_{STD-L(i-1)} + Q_{STD-R(i-1)}}$$
$$p_{L(i)} = \sqrt{\frac{1}{1 + e^{\beta_i (Q_{R(i-1)} - Q_{L(i-1)})}}}$$

**For each Side:**

$$RPE_i = r_i - (Q_{chosen(i-1)})$$
$$Q_{chosen(i)} = Q_{chosen(i-1)} + \alpha_a RPE$$
$$Q_{unchosen(i)} = Q_{unchosen(i-1)} \alpha_f$$
$$p_{L(i)} = \frac{1}{1 + e^{(Q_{R(i-1)} - Q_{L(i-1)})}}$$

**Supplementary Figure 7. VS CIN RPE coding does not depend on choice of model.**

Average CIN firing rates and fractions of value-modulated CINs, using different models to estimate values. Data and format is the same as Fig. 4B, in each case using terciles of value. Rows from top: Bayesian model value (as Fig. 4B; see Methods), calculated as the mean of a beta distribution; number of pellets received in the previous minute; leaky integrator for value, with decay parameter best-fit to latencies in each session; delta-rule / Rescorla-Wagner model, with learning rate alpha best-fit to latencies in each session; last two rows extend the previous two models by calculating separate action values for each side, and fitting parameters based on rat choices, with CIN firing compared to the action value for the chosen option).
