## Supplementary File: Each identified neuron for "A mismatch between striatal cholinergic pauses and dopaminergic reward prediction errors"

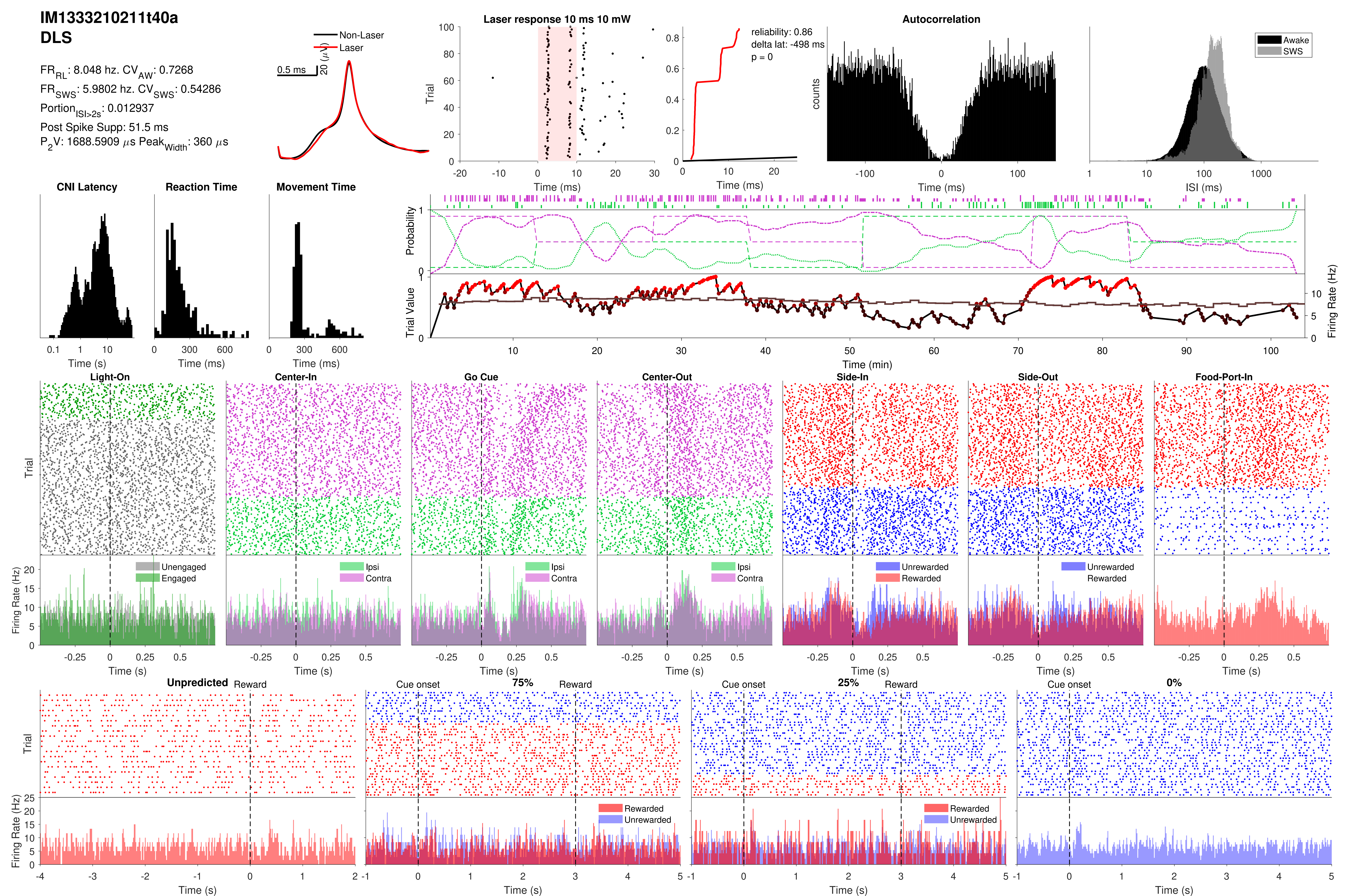

IM1333210211t40a
DLS
FR_RL: 8.048 hz. CV_AW: 0.7268
FR_SWS: 5.9802 hz. CV_SWS: 0.54286
Portion_ISI>2s: 0.012937
Post Spike Supp: 51.5 ms
P_2V: 1688.5909 μs Peak_Width: 360 μs
Non-Laser
Laser
0.5 ms
20 (μV)
Laser response 10 ms 10 mW
Trial
Time (ms)
reliability: 0.86
delta lat: -498 ms
p = 0
Time (ms)
Autocorrelation
counts
Time (ms)
Awake
SWS
ISI (ms)
CNI Latency
Reaction Time
Movement Time
Time (s)
Time (ms)
Time (ms)
Probability
Trial Value
Firing Rate (Hz)
Time (min)
Light-On
Center-In
Go Cue
Center-Out
Side-In
Side-Out
Food-Port-In
Trial
Firing Rate (Hz)
Unengaged
Engaged
Ipsi
Contra
Unrewarded
Rewarded
Time (s)
Unpredicted
Reward
Cue onset
75%
25%
0%
Rewarded
Unrewarded
Time (s)

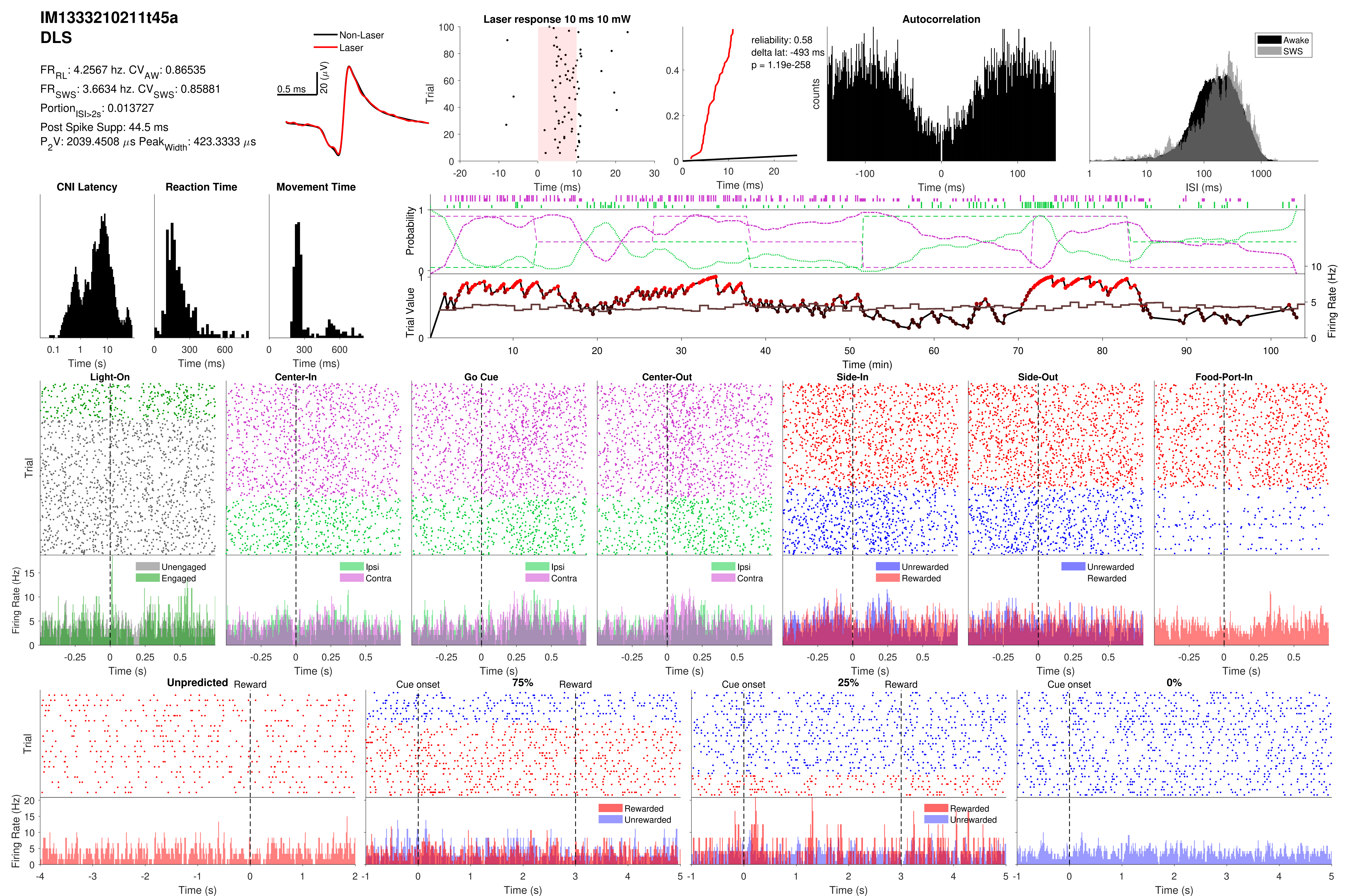
IM1333210211t45a
DLS
$FR_{RL}$: 4.2567 hz. $CV_{AW}$: 0.86535
$FR_{SWS}$: 3.6634 hz. $CV_{SWS}$: 0.85881
$Portion_{ISI>2s}$: 0.013727
Post Spike Supp: 44.5 ms
$P_2V$: 2039.4508 $\mu s$ $Peak_{Width}$: 423.3333 $\mu s$
Non-Laser
Laser
0.5 ms
20 ($\mu$V)
Laser response 10 ms 10 mW
Trial
Time (ms)
reliability: 0.58
delta lat: -493 ms
p = 1.19e-258
Time (ms)
Autocorrelation
counts
Time (ms)
Awake
SWS
ISI (ms)
CNI Latency
Time (s)
Reaction Time
Time (ms)
Movement Time
Time (ms)
Probability
Trial Value
Firing Rate (Hz)
Time (min)
Light-On
Center-In
Go Cue
Center-Out
Side-In
Side-Out
Food-Port-In
Trial
Firing Rate (Hz)
Unengaged
Engaged
Ipsi
Contra
Unrewarded
Rewarded
Time (s)
Unpredicted
Reward
Cue onset
75%
25%
0%
Rewarded
Unrewarded
Time (s)

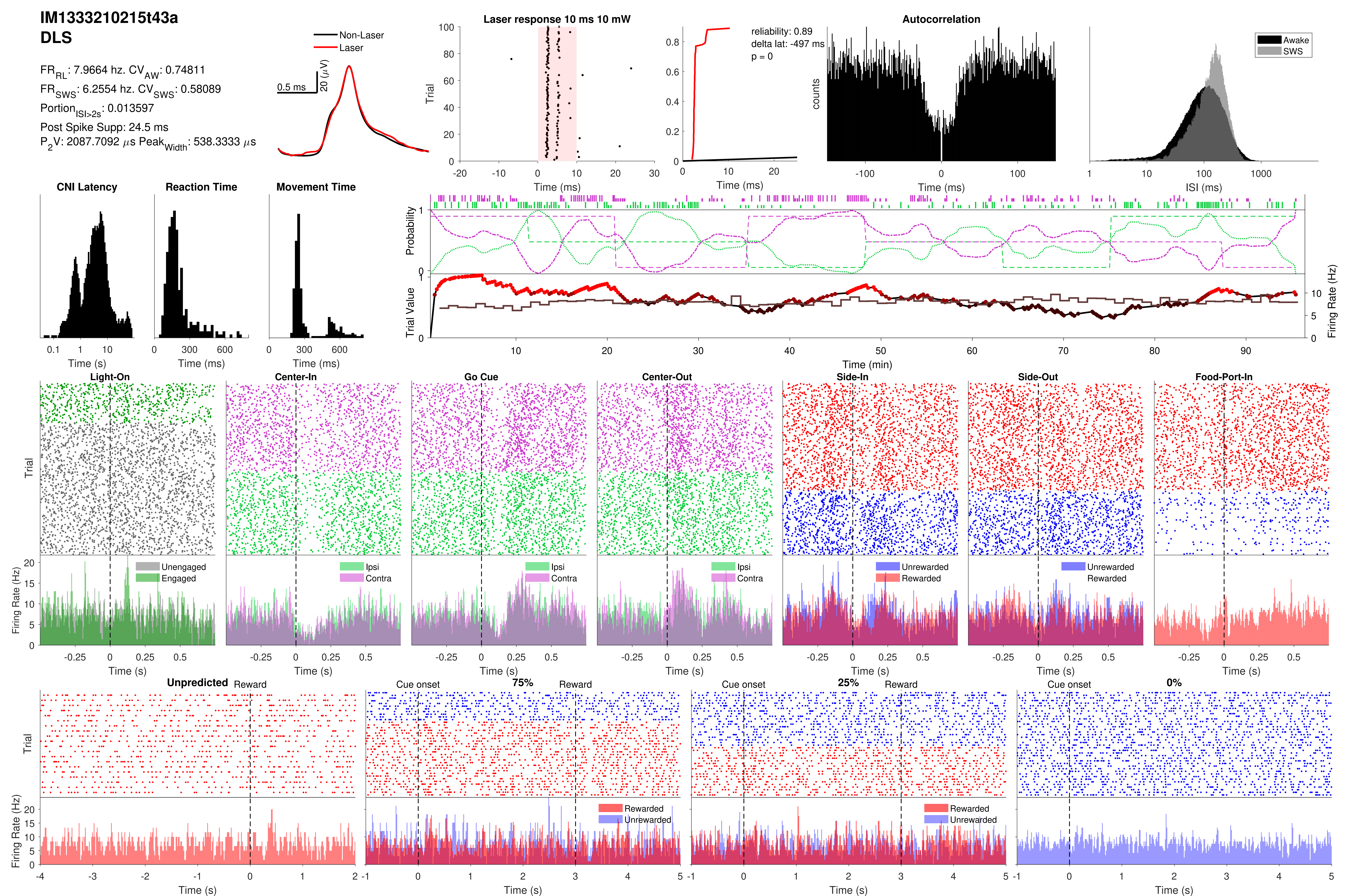

IM1333210215t43a
DLS
FR_RL: 7.9664 hz. CV_AW: 0.74811
FR_SWS: 6.2554 hz. CV_SWS: 0.58089
Portion_ISI>2s: 0.013597
Post Spike Supp: 24.5 ms
P_2V: 2087.7092 µs Peak_Width: 538.3333 µs
Non-Laser
Laser
0.5 ms
20 (µV)
Laser response 10 ms 10 mW
Trial
Time (ms)
reliability: 0.89
delta lat: -497 ms
p = 0
Autocorrelation
counts
Awake
SWS
ISI (ms)
CNI Latency
Reaction Time
Movement Time
Time (s)
Time (ms)
Probability
Trial Value
Firing Rate (Hz)
Time (min)
Light-On
Center-In
Go Cue
Center-Out
Side-In
Side-Out
Food-Port-In
Unengaged
Engaged
Ipsi
Contra
Unrewarded
Rewarded
Unpredicted
Reward
Cue onset
75%
25%
0%

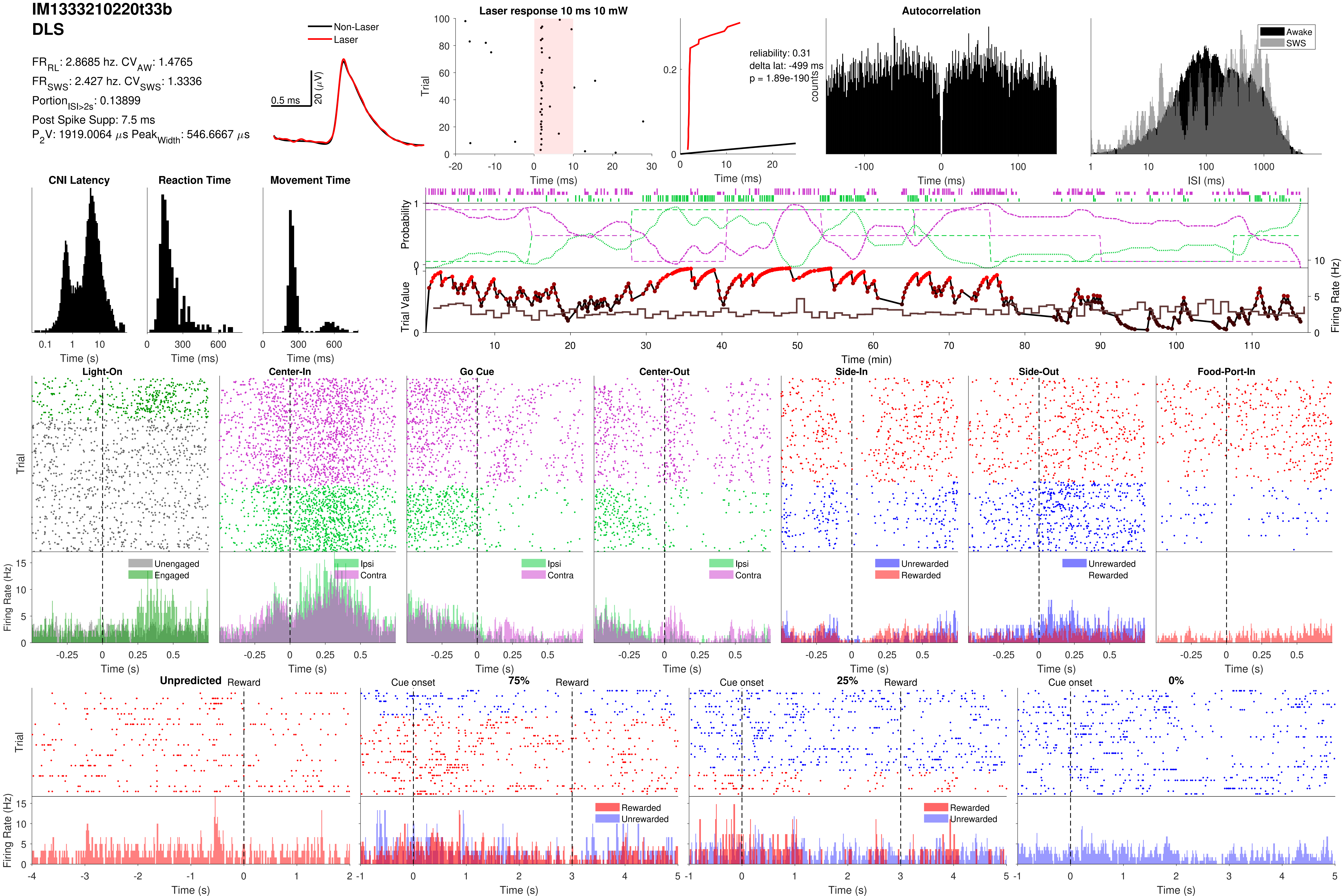
IM1333210220t33b
DLS
FR_RL: 2.8685 hz. CV_AW: 1.4765
FR_SWS: 2.427 hz. CV_SWS: 1.3336
Portion_ISI>2s: 0.13899
Post Spike Supp: 7.5 ms
P2V: 1919.0064 µs Peak_Width: 546.6667 µs
Non-Laser
Laser
0.5 ms
20 (µV)
Laser response 10 ms 10 mW
Trial
Time (ms)
reliability: 0.31
delta lat: -499 ms
p = 1.89e-190
Time (ms)
Autocorrelation
counts
Time (ms)
Awake
SWS
ISI (ms)
CNI Latency
Time (s)
Reaction Time
Time (ms)
Movement Time
Time (ms)
Probability
Trial Value
Firing Rate (Hz)
Time (min)
Light-On
Center-In
Go Cue
Center-Out
Side-In
Side-Out
Food-Port-In
Trial
Firing Rate (Hz)
Unengaged
Engaged
Ipsi
Contra
Unrewarded
Rewarded
Time (s)
Unpredicted
Reward
Cue onset
75%
25%
0%
Rewarded
Unrewarded
Time (s)

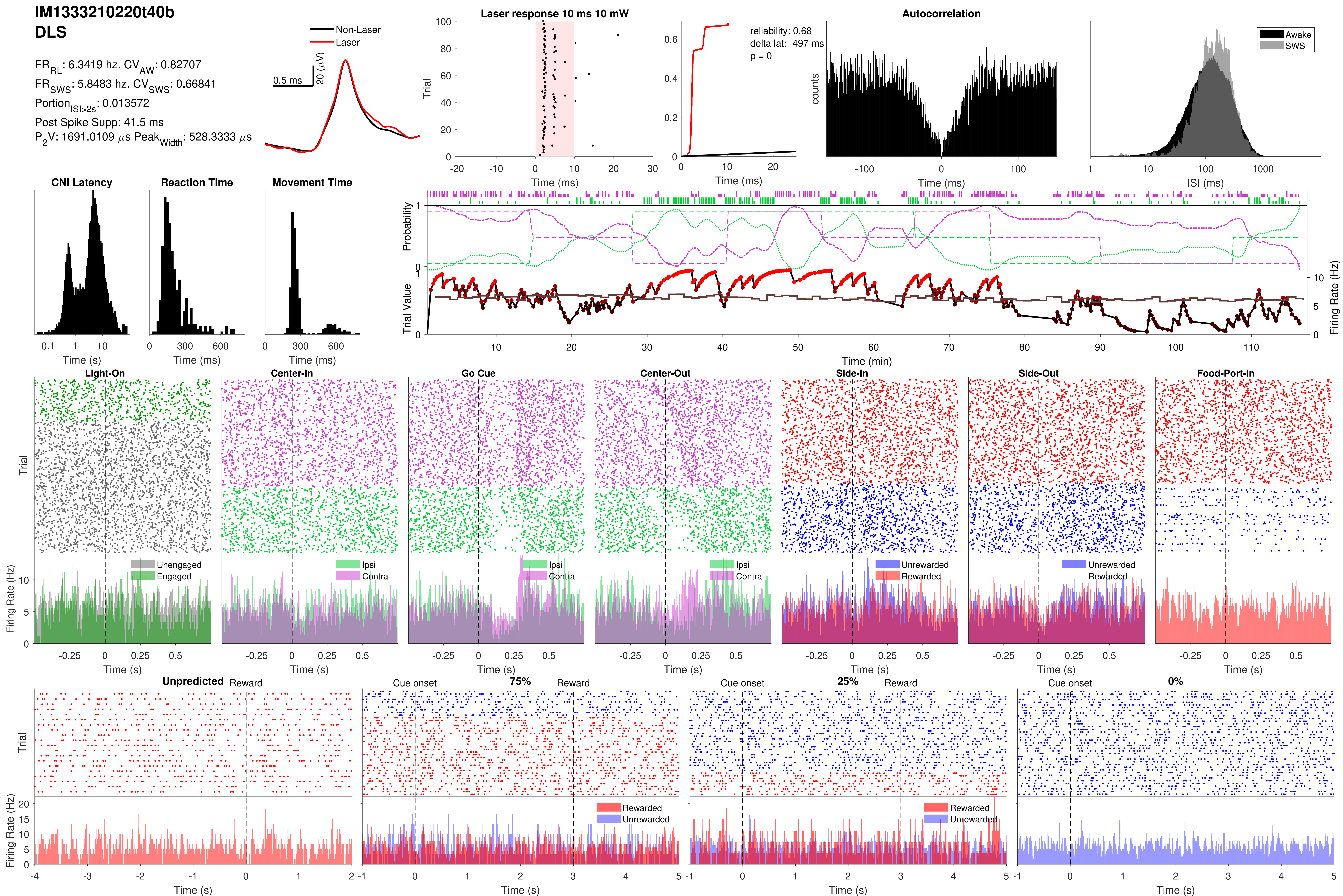
IM1333210220t40b
DLS
FR_RL: 6.3419 hz. CV_AW: 0.82707
FR_SWS: 5.8483 hz. CV_SWS: 0.66841
Portion_ISI>2s: 0.013572
Post Spike Supp: 41.5 ms
P_2V: 1691.0109 µs Peak_Width: 528.3333 µs
Non-Laser
Laser
0.5 ms
20 (µV)
Laser response 10 ms 10 mW
Trial
Time (ms)
reliability: 0.68
delta lat: -497 ms
p = 0
Autocorrelation
counts
Awake
SWS
ISI (ms)
CNI Latency
Reaction Time
Movement Time
Time (s)
Time (ms)
Probability
Trial Value
Firing Rate (Hz)
Time (min)
Light-On
Center-In
Go Cue
Center-Out
Side-In
Side-Out
Food-Port-In
Unengaged
Engaged
Ipsi
Contra
Unrewarded
Rewarded
Unpredicted
Reward
Cue onset
75%
25%
0%

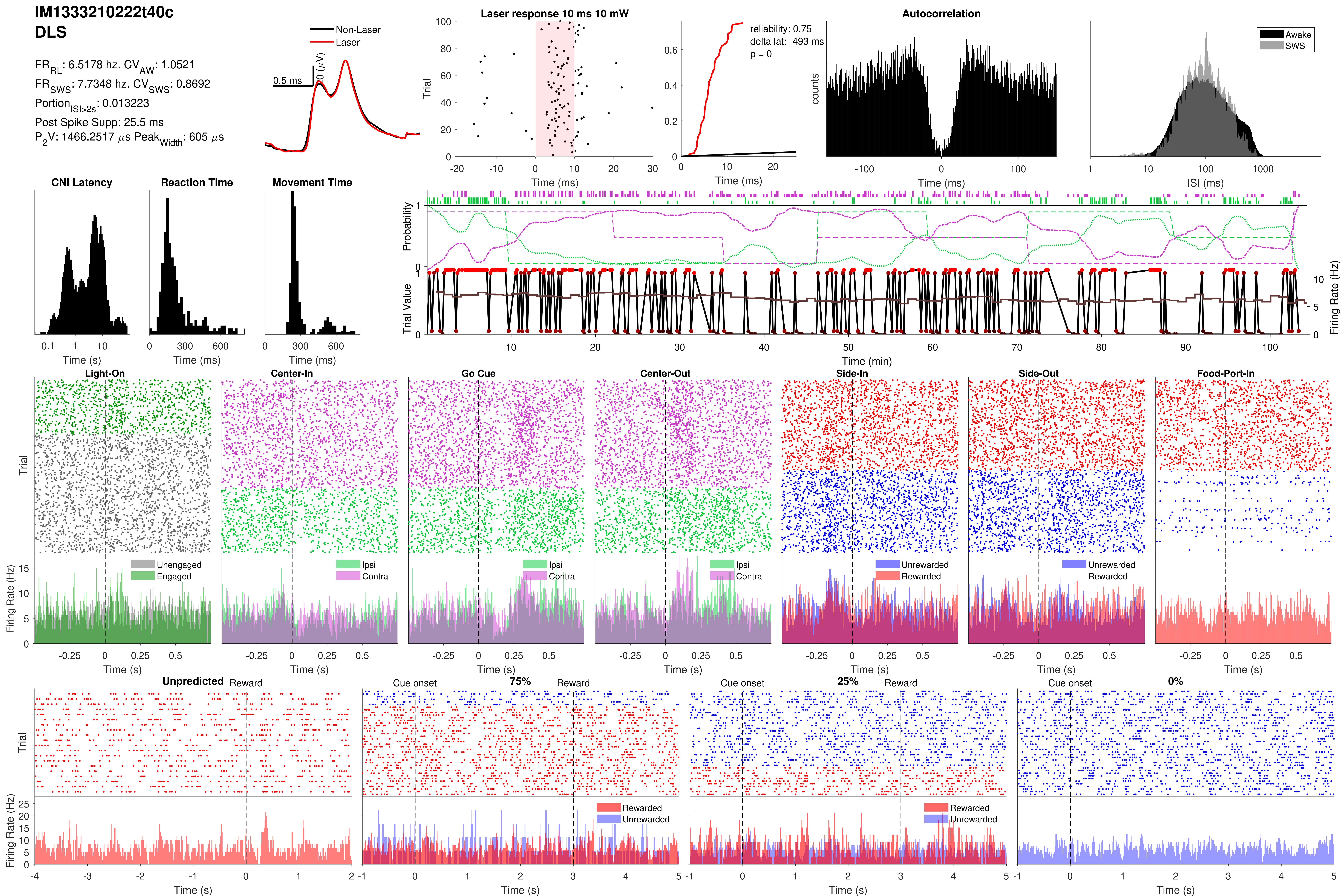

IM1333210222t40c
DLS
FR_RL: 6.5178 hz. CV_AW: 1.0521
FR_SWS: 7.7348 hz. CV_SWS: 0.8692
Portion_ISI>2s: 0.013223
Post Spike Supp: 25.5 ms
P_2V: 1466.2517 μs Peak_Width: 605 μs
Non-Laser
Laser
0.5 ms
20 (μV)
Laser response 10 ms 10 mW
reliability: 0.75
delta lat: -493 ms
p = 0
Autocorrelation
Awake
SWS
counts
ISI (ms)
CNI Latency
Reaction Time
Movement Time
Probability
Trial Value
Firing Rate (Hz)
Time (min)
Light-On
Center-In
Go Cue
Center-Out
Side-In
Side-Out
Food-Port-In
Unengaged
Engaged
Ipsi
Contra
Unrewarded
Rewarded
Trial
Time (ms)
Time (s)
Unpredicted
Reward
Cue onset
75%
25%
0%

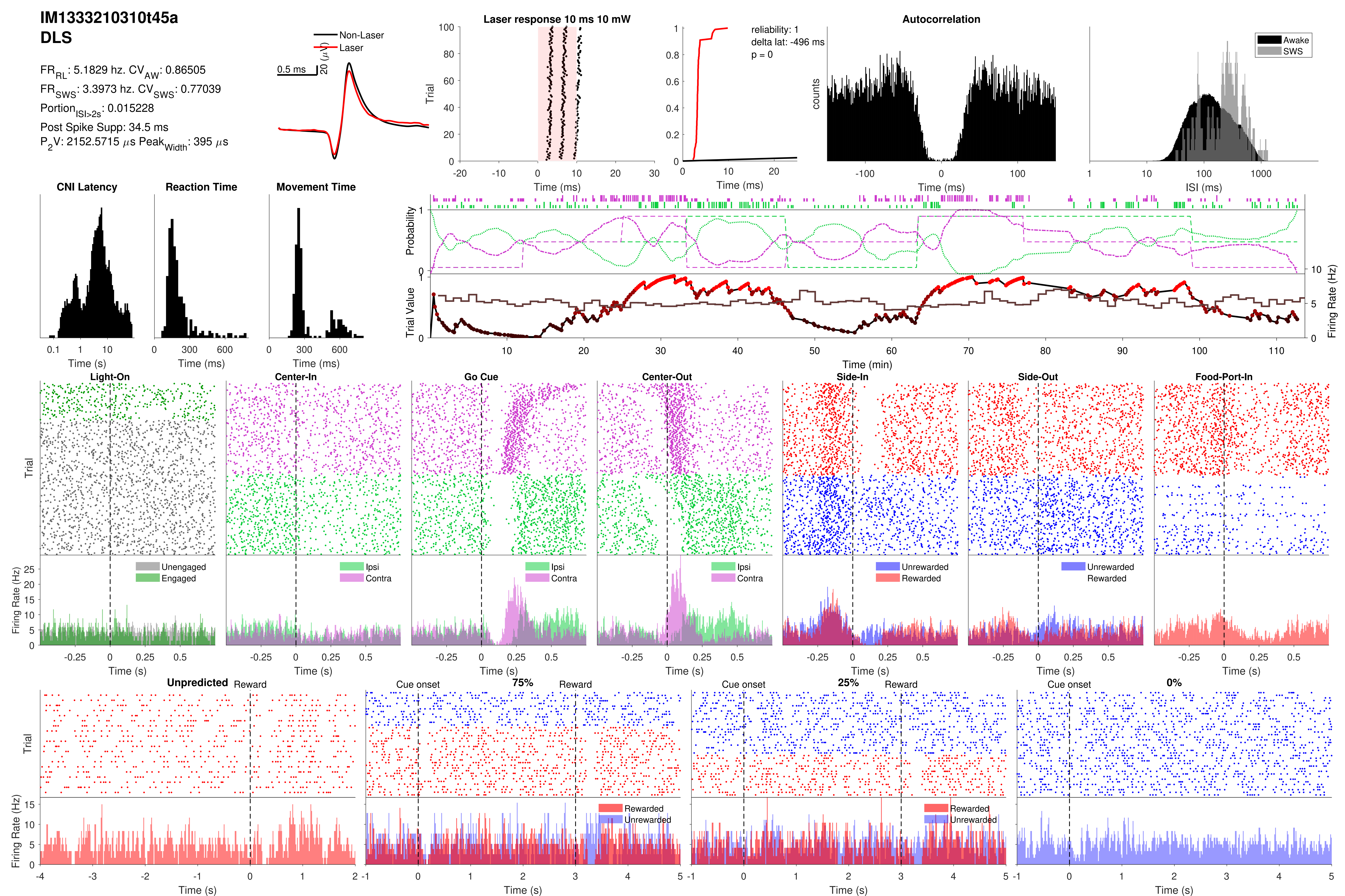

IM1333210310t45a
DLS
FR_RL: 5.1829 hz. CV_AW: 0.86505
FR_SWS: 3.3973 hz. CV_SWS: 0.77039
Portion_ISI>2s: 0.015228
Post Spike Supp: 34.5 ms
P_2V: 2152.5715 µs Peak_Width: 395 µs
Non-Laser
Laser
0.5 ms
20 (µV)
Laser response 10 ms 10 mW
Trial
Time (ms)
reliability: 1
delta lat: -496 ms
p = 0
Autocorrelation
counts
Awake
SWS
ISI (ms)
CNI Latency
Reaction Time
Movement Time
Time (s)
Time (ms)
Probability
Trial Value
Firing Rate (Hz)
Time (min)
Light-On
Center-In
Go Cue
Center-Out
Side-In
Side-Out
Food-Port-In
Unengaged
Engaged
Ipsi
Contra
Unrewarded
Rewarded
Unpredicted
Reward
Cue onset
75%
25%
0%

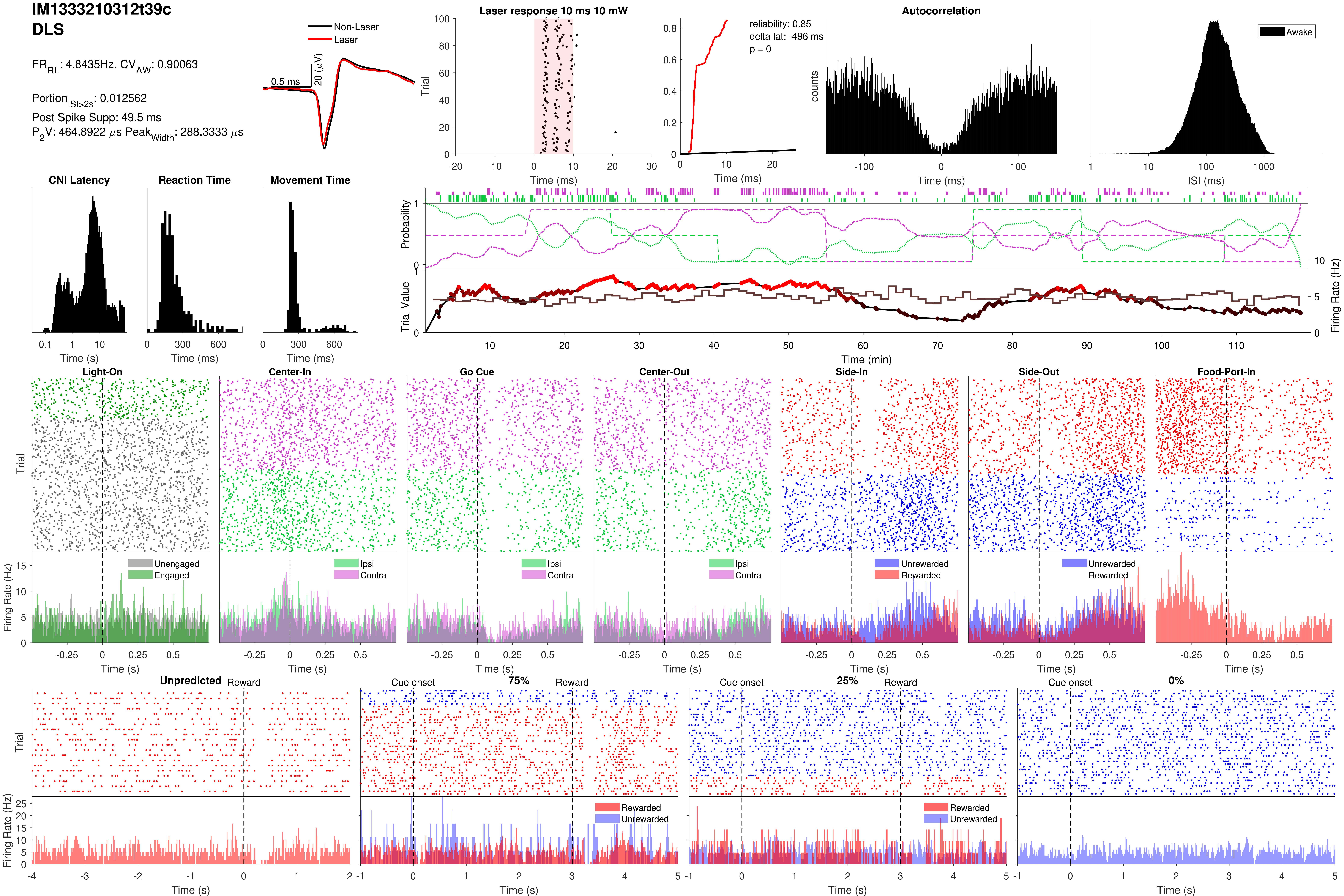

IM1333210312t39c
DLS
FR$_{RL}$: 4.8435Hz. CV$_{AW}$: 0.90063
Portion$_{ISI>2s}$: 0.012562
Post Spike Supp: 49.5 ms
P$_2$V: 464.8922 $\mu$s Peak$_{Width}$: 288.3333 $\mu$s
Non-Laser
Laser
0.5 ms
20 ($\mu$V)
Laser response 10 ms 10 mW
Trial
Time (ms)
reliability: 0.85
delta lat: -496 ms
p = 0
Time (ms)
Autocorrelation
counts
Time (ms)
Awake
ISI (ms)
CNI Latency
Time (s)
Reaction Time
Time (ms)
Movement Time
Time (ms)
Probability
Trial Value
Firing Rate (Hz)
Time (min)
Light-On
Center-In
Go Cue
Center-Out
Side-In
Side-Out
Food-Port-In
Trial
Firing Rate (Hz)
Unengaged
Engaged
Ipsi
Contra
Unrewarded
Rewarded
Time (s)
Unpredicted
Reward
Cue onset
75%
25%
0%
Rewarded
Unrewarded
Time (s)

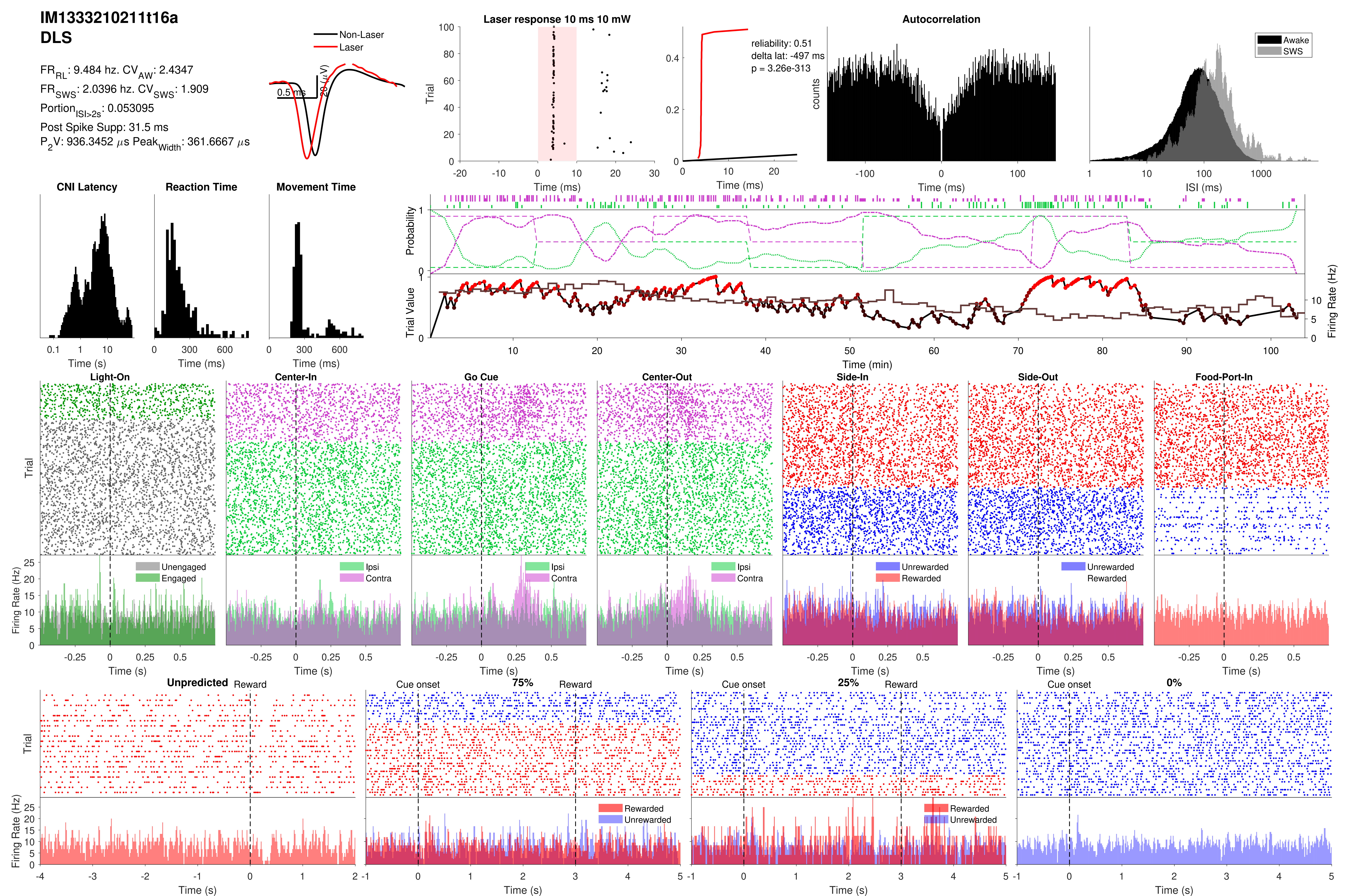

IM1333210211t16a
DLS
FR_RL: 9.484 hz. CV_AW: 2.4347
FR_SWS: 2.0396 hz. CV_SWS: 1.909
Portion_ISI>2s: 0.053095
Post Spike Supp: 31.5 ms
P_2V: 936.3452 μs Peak_Width: 361.6667 μs
Non-Laser
Laser
0.5 ms
20 (μV)
Laser response 10 ms 10 mW
Trial
Time (ms)
reliability: 0.51
delta lat: -497 ms
p = 3.26e-313
Time (ms)
Autocorrelation
counts
Time (ms)
Awake
SWS
ISI (ms)
CNI Latency
Time (s)
Reaction Time
Time (ms)
Movement Time
Time (ms)
Probability
Trial Value
Firing Rate (Hz)
Time (min)
Light-On
Center-In
Go Cue
Center-Out
Side-In
Side-Out
Food-Port-In
Trial
Firing Rate (Hz)
Unengaged
Engaged
Ipsi
Contra
Unrewarded
Rewarded
Time (s)
Unpredicted
Reward
Cue onset
75%
25%
0%
Rewarded
Unrewarded
Time (s)

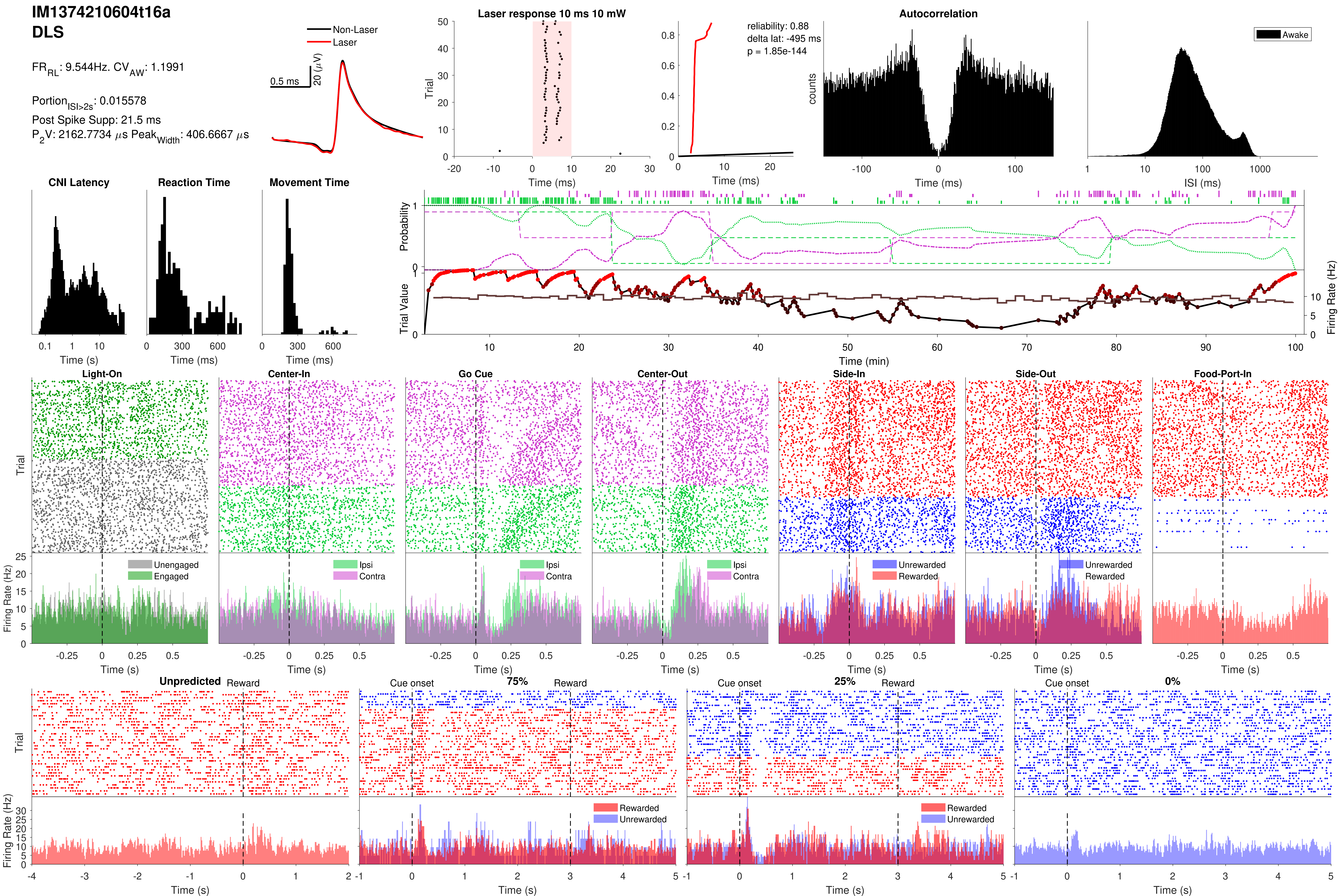

IM1374210604t16a
DLS
FR_RL: 9.544Hz. CV_AW: 1.1991
Portion_ISI>2s: 0.015578
Post Spike Supp: 21.5 ms
P_2V: 2162.7734 µs Peak_Width: 406.6667 µs
Non-Laser
Laser
0.5 ms
20 (µV)
Laser response 10 ms 10 mW
Trial
Time (ms)
reliability: 0.88
delta lat: -495 ms
p = 1.85e-144
Time (ms)
Autocorrelation
counts
Time (ms)
Awake
ISI (ms)
CNI Latency
Time (s)
Reaction Time
Time (ms)
Movement Time
Time (ms)
Probability
Trial Value
Firing Rate (Hz)
Time (min)
Light-On
Center-In
Go Cue
Center-Out
Side-In
Side-Out
Food-Port-In
Trial
Firing Rate (Hz)
Unengaged
Engaged
Ipsi
Contra
Unrewarded
Rewarded
Time (s)
Unpredicted
Reward
Cue onset
75%
25%
0%
Rewarded
Unrewarded
Trial
Firing Rate (Hz)
Time (s)

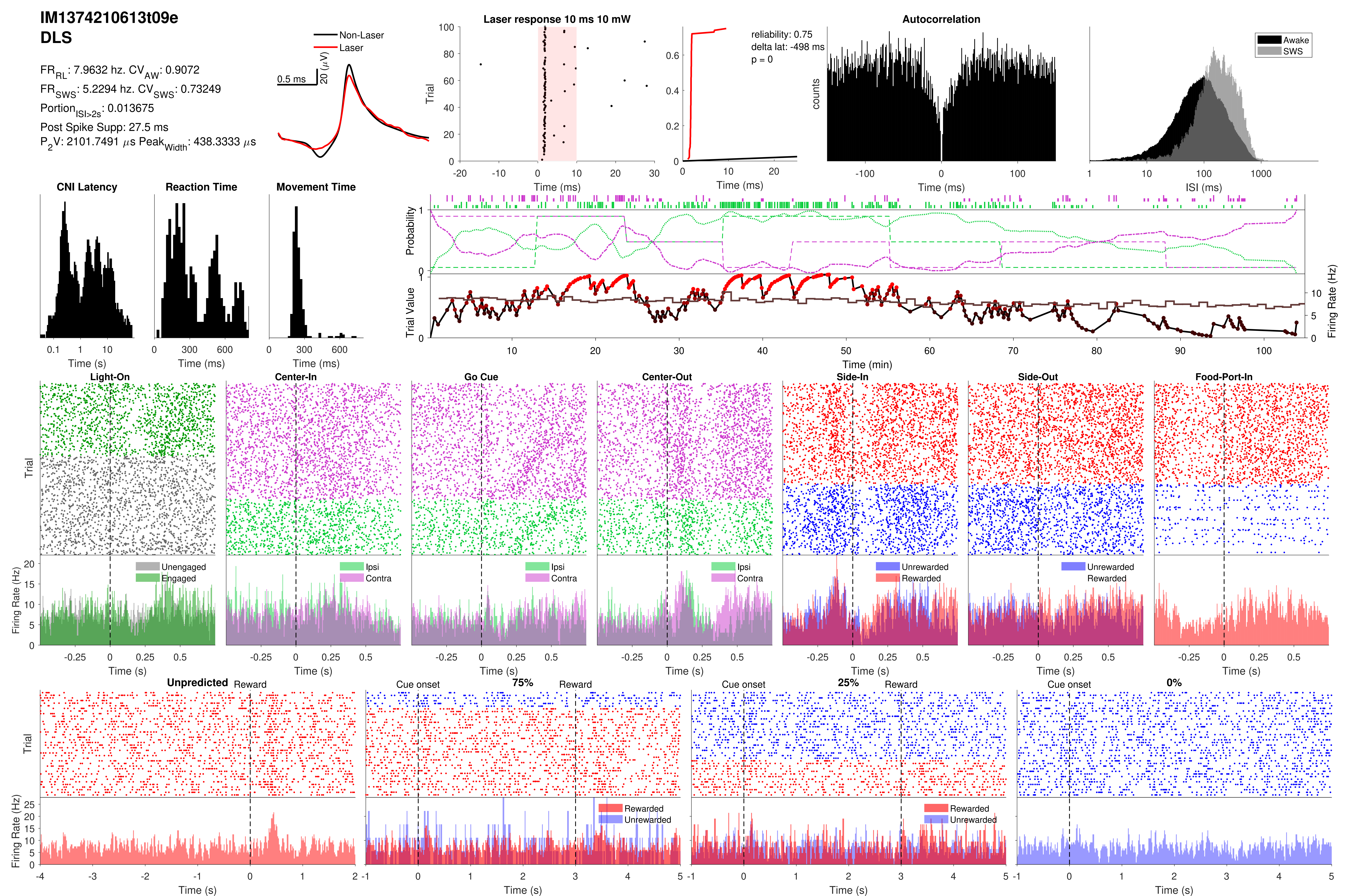

IM1374210613t09e
DLS
FR_RL: 7.9632 hz. CV_AW: 0.9072
FR_SWS: 5.2294 hz. CV_SWS: 0.73249
Portion_ISI>2s: 0.013675
Post Spike Supp: 27.5 ms
P_2V: 2101.7491 μs Peak_Width: 438.3333 μs
Non-Laser
Laser
0.5 ms
20 (μV)
Laser response 10 ms 10 mW
Trial
Time (ms)
reliability: 0.75
delta lat: -498 ms
p = 0
Autocorrelation
counts
Awake
SWS
ISI (ms)
CNI Latency
Reaction Time
Movement Time
Time (s)
Probability
Trial Value
Firing Rate (Hz)
Time (min)
Light-On
Center-In
Go Cue
Center-Out
Side-In
Side-Out
Food-Port-In
Unengaged
Engaged
Ipsi
Contra
Unrewarded
Rewarded
Unpredicted
Reward
Cue onset
75%
25%
0%

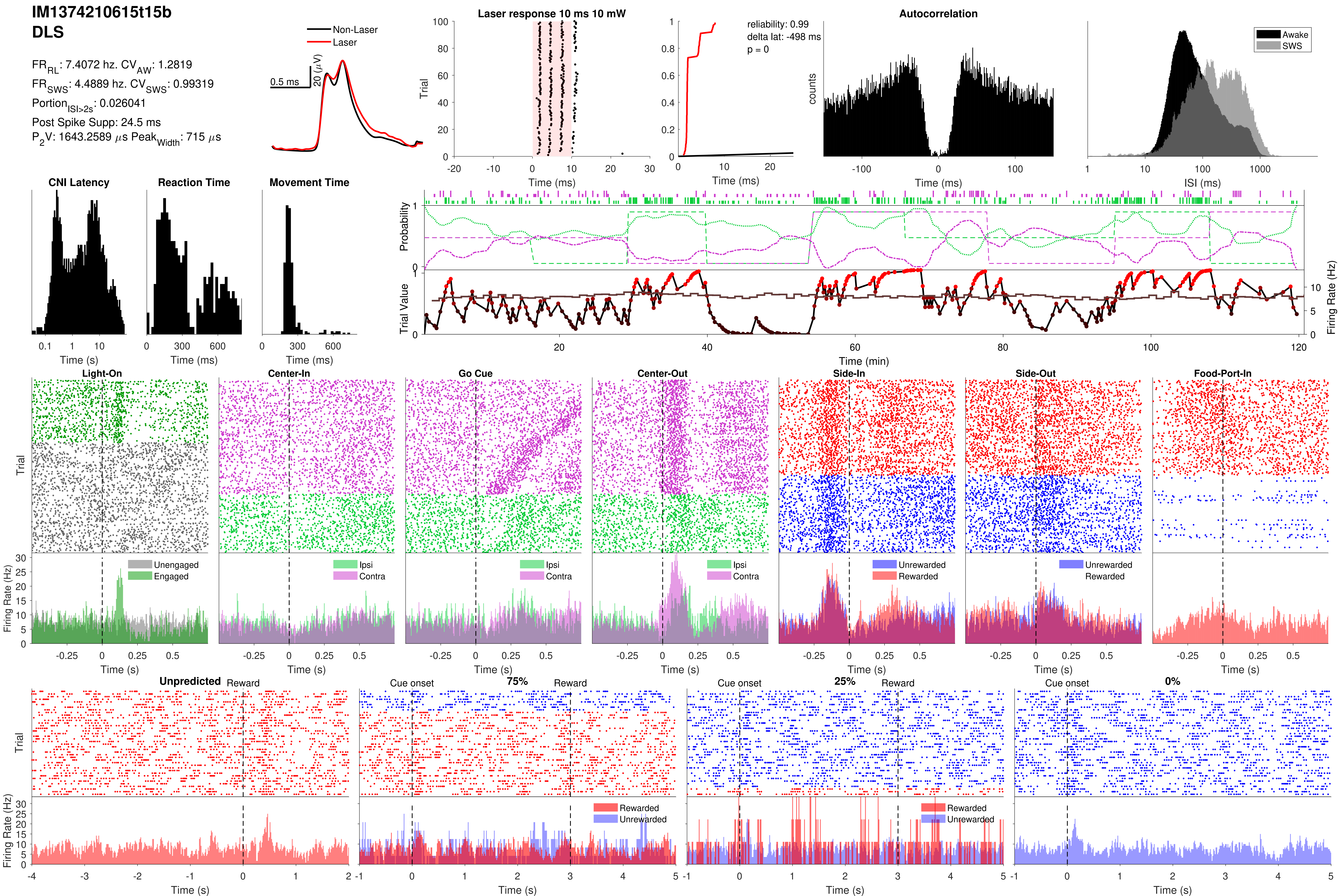

IM1374210615t15b
DLS
FR_RL: 7.4072 hz. CV_AW: 1.2819
FR_SWS: 4.4889 hz. CV_SWS: 0.99319
Portion_ISI>2s: 0.026041
Post Spike Supp: 24.5 ms
P_2V: 1643.2589 µs Peak_Width: 715 µs
Non-Laser
Laser
0.5 ms
20 (µV)
Laser response 10 ms 10 mW
Trial
Time (ms)
reliability: 0.99
delta lat: -498 ms
p = 0
Time (ms)
Autocorrelation
counts
Time (ms)
Awake
SWS
ISI (ms)
CNI Latency
Time (s)
Reaction Time
Time (ms)
Movement Time
Time (ms)
Probability
Trial Value
Firing Rate (Hz)
Time (min)
Light-On
Center-In
Go Cue
Center-Out
Side-In
Side-Out
Food-Port-In
Trial
Firing Rate (Hz)
Unengaged
Engaged
Ipsi
Contra
Unrewarded
Rewarded
Time (s)
Unpredicted
Reward
Cue onset
75%
25%
0%
Trial
Firing Rate (Hz)
Rewarded
Unrewarded
Time (s)

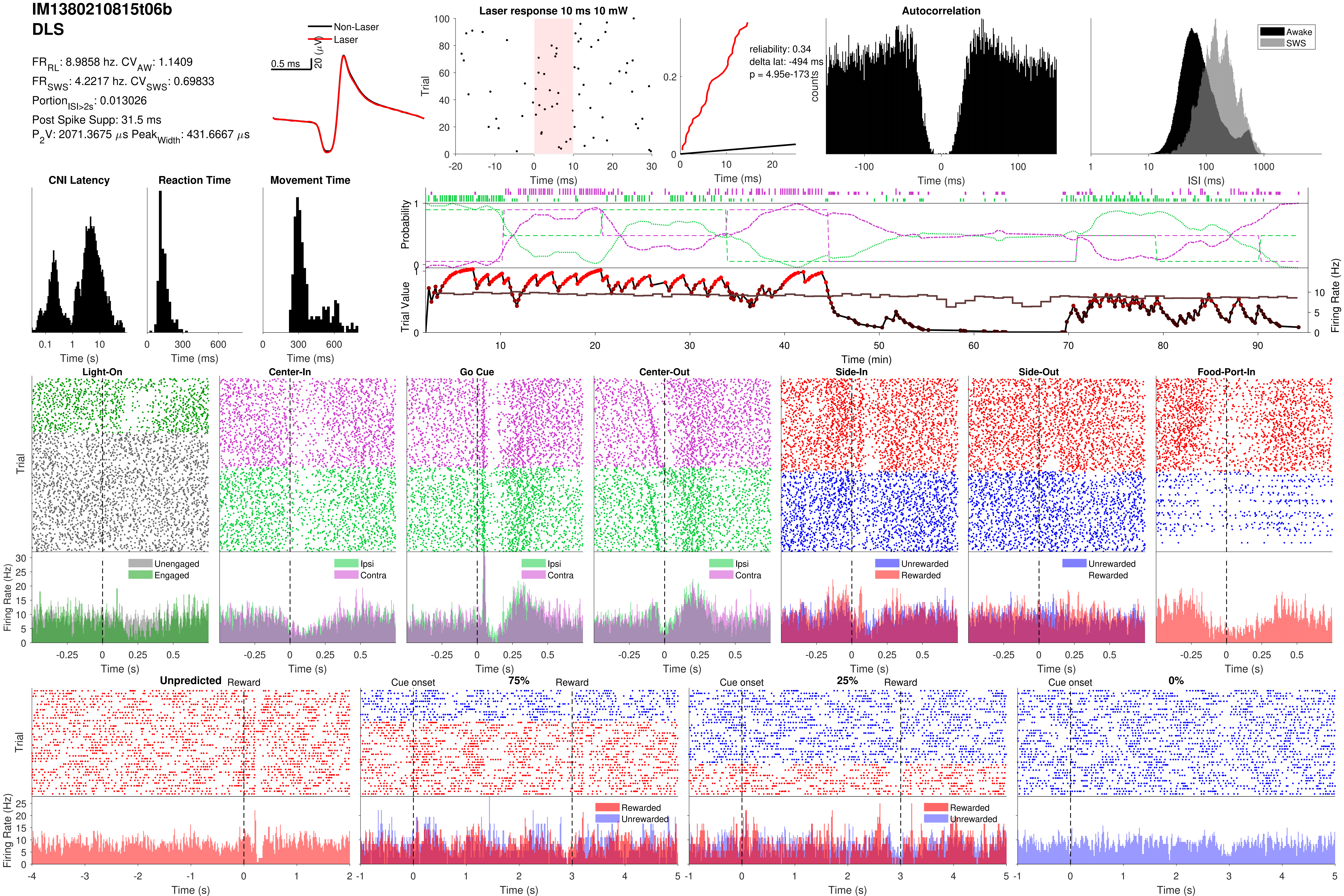

IM1380210815t06b
DLS
FR_RL: 8.9858 hz. CV_AW: 1.1409
FR_SWS: 4.2217 hz. CV_SWS: 0.69833
Portion_ISI>2s: 0.013026
Post Spike Supp: 31.5 ms
P_2V: 2071.3675 μs Peak_Width: 431.6667 μs
Non-Laser
Laser
0.5 ms
20 (μV)
Laser response 10 ms 10 mW
Trial
Time (ms)
reliability: 0.34
delta lat: -494 ms
p = 4.95e-173
Autocorrelation
counts
Awake
SWS
ISI (ms)
CNI Latency
Reaction Time
Movement Time
Time (s)
Time (ms)
Probability
Trial Value
Firing Rate (Hz)
Time (min)
Light-On
Center-In
Go Cue
Center-Out
Side-In
Side-Out
Food-Port-In
Unengaged
Engaged
Ipsi
Contra
Unrewarded
Rewarded
Unpredicted
Reward
Cue onset
75%
25%
0%

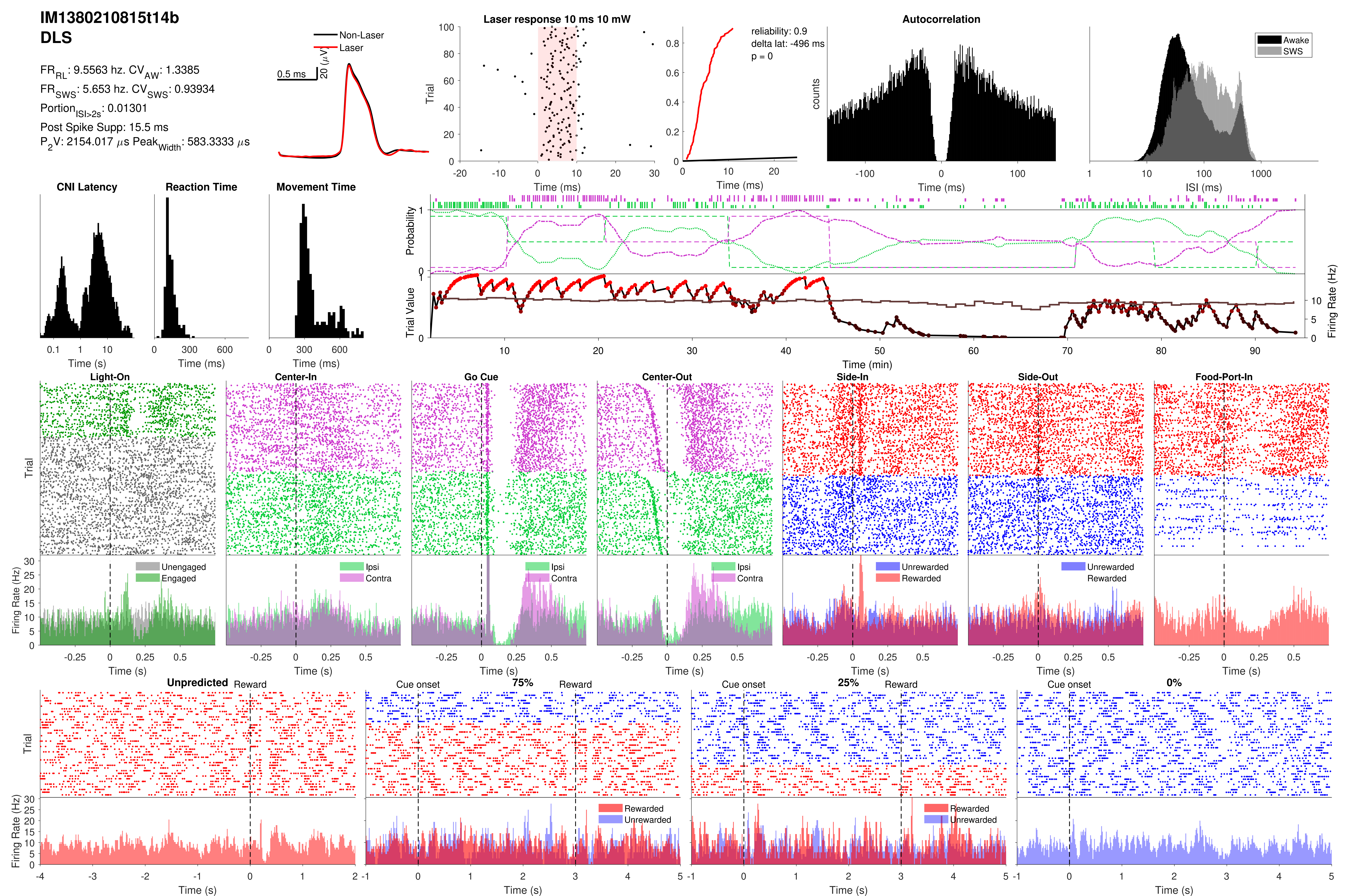
IM1380210815t14b
DLS
FR_RL: 9.5563 hz. CV_AW: 1.3385
FR_SWS: 5.653 hz. CV_SWS: 0.93934
Portion_ISI>2s: 0.01301
Post Spike Supp: 15.5 ms
P_2V: 2154.017 μs Peak_Width: 583.3333 μs
Non-Laser
Laser
0.5 ms
20 (μV)
Laser response 10 ms 10 mW
Trial
Time (ms)
reliability: 0.9
delta lat: -496 ms
p = 0
Time (ms)
Autocorrelation
counts
Time (ms)
Awake
SWS
ISI (ms)
CNI Latency
Time (s)
Reaction Time
Time (ms)
Movement Time
Time (ms)
Probability
Trial Value
Firing Rate (Hz)
Time (min)
Light-On
Center-In
Go Cue
Center-Out
Side-In
Side-Out
Food-Port-In
Trial
Firing Rate (Hz)
Unengaged
Engaged
Ipsi
Contra
Unrewarded
Rewarded
Time (s)
Unpredicted
Reward
Cue onset
75%
25%
0%
Rewarded
Unrewarded
Trial
Firing Rate (Hz)
Time (s)

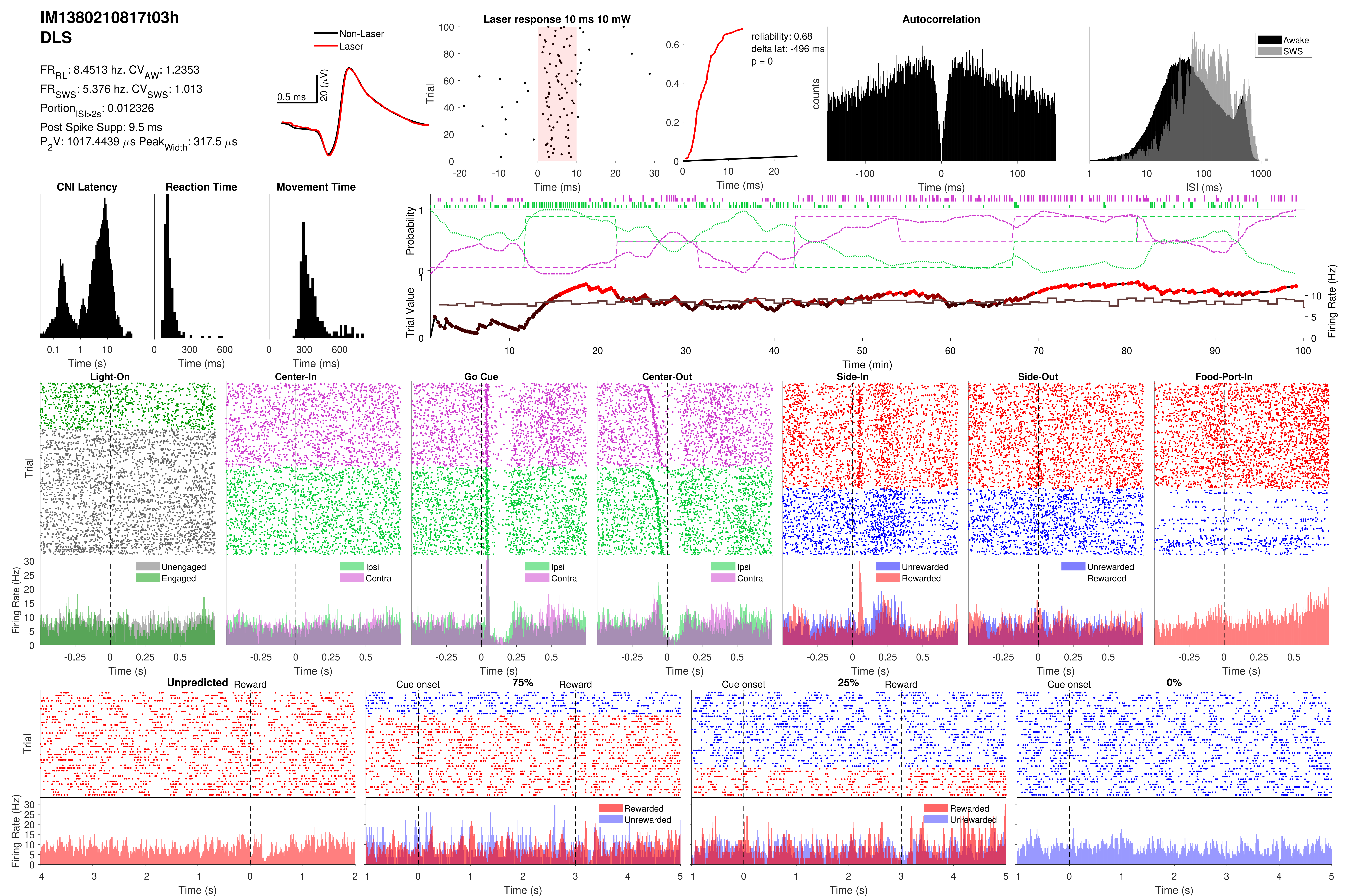

IM1380210817t03h
DLS
FR_RL: 8.4513 hz. CV_AW: 1.2353
FR_SWS: 5.376 hz. CV_SWS: 1.013
Portion_ISI>2s: 0.012326
Post Spike Supp: 9.5 ms
P_2V: 1017.4439 μs Peak_Width: 317.5 μs
Non-Laser
Laser
0.5 ms
20 (μV)
Laser response 10 ms 10 mW
reliability: 0.68
delta lat: -496 ms
p = 0
Autocorrelation
Awake
SWS
CNI Latency
Reaction Time
Movement Time
Light-On
Center-In
Go Cue
Center-Out
Side-In
Side-Out
Food-Port-In
Unengaged
Engaged
Ipsi
Contra
Unrewarded
Rewarded
Unpredicted
75%
25%
0%
Cue onset
Reward

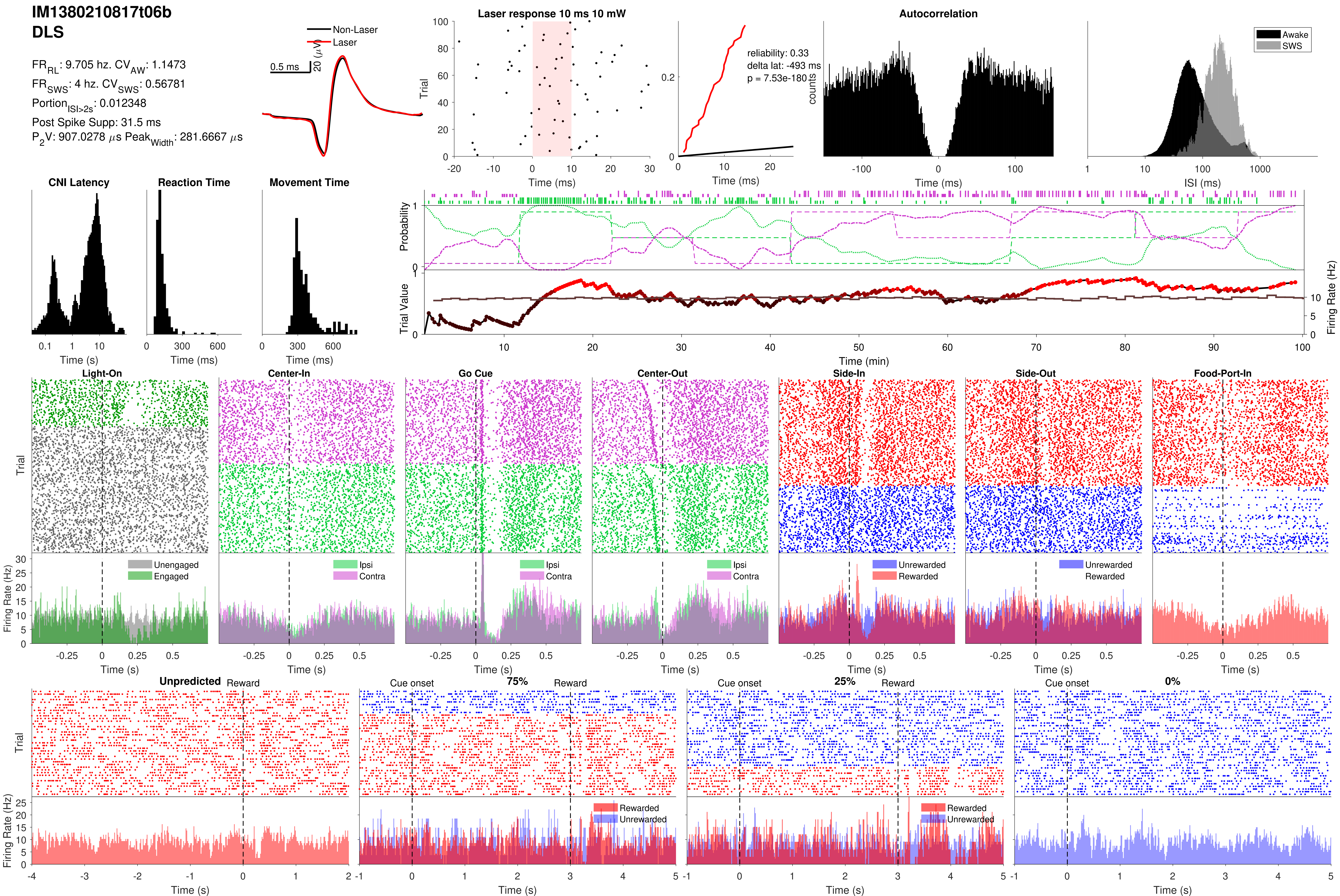
IM1380210817t06b
DLS
FR_RL: 9.705 hz. CV_AW: 1.1473
FR_SWS: 4 hz. CV_SWS: 0.56781
Portion_ISI>2s: 0.012348
Post Spike Supp: 31.5 ms
P_2V: 907.0278 µs Peak_Width: 281.6667 µs
Non-Laser
Laser
0.5 ms
20 (µV)
Laser response 10 ms 10 mW
reliability: 0.33
delta lat: -493 ms
p = 7.53e-180
Autocorrelation
Awake
SWS
CNI Latency
Reaction Time
Movement Time
Probability
Trial Value
Firing Rate (Hz)
Time (min)
Light-On
Center-In
Go Cue
Center-Out
Side-In
Side-Out
Food-Port-In
Unengaged
Engaged
Ipsi
Contra
Unrewarded
Rewarded
Unpredicted
Reward
Cue onset
75%
25%
0%
Trial
Time (ms)
Time (s)
ISI (ms)
counts

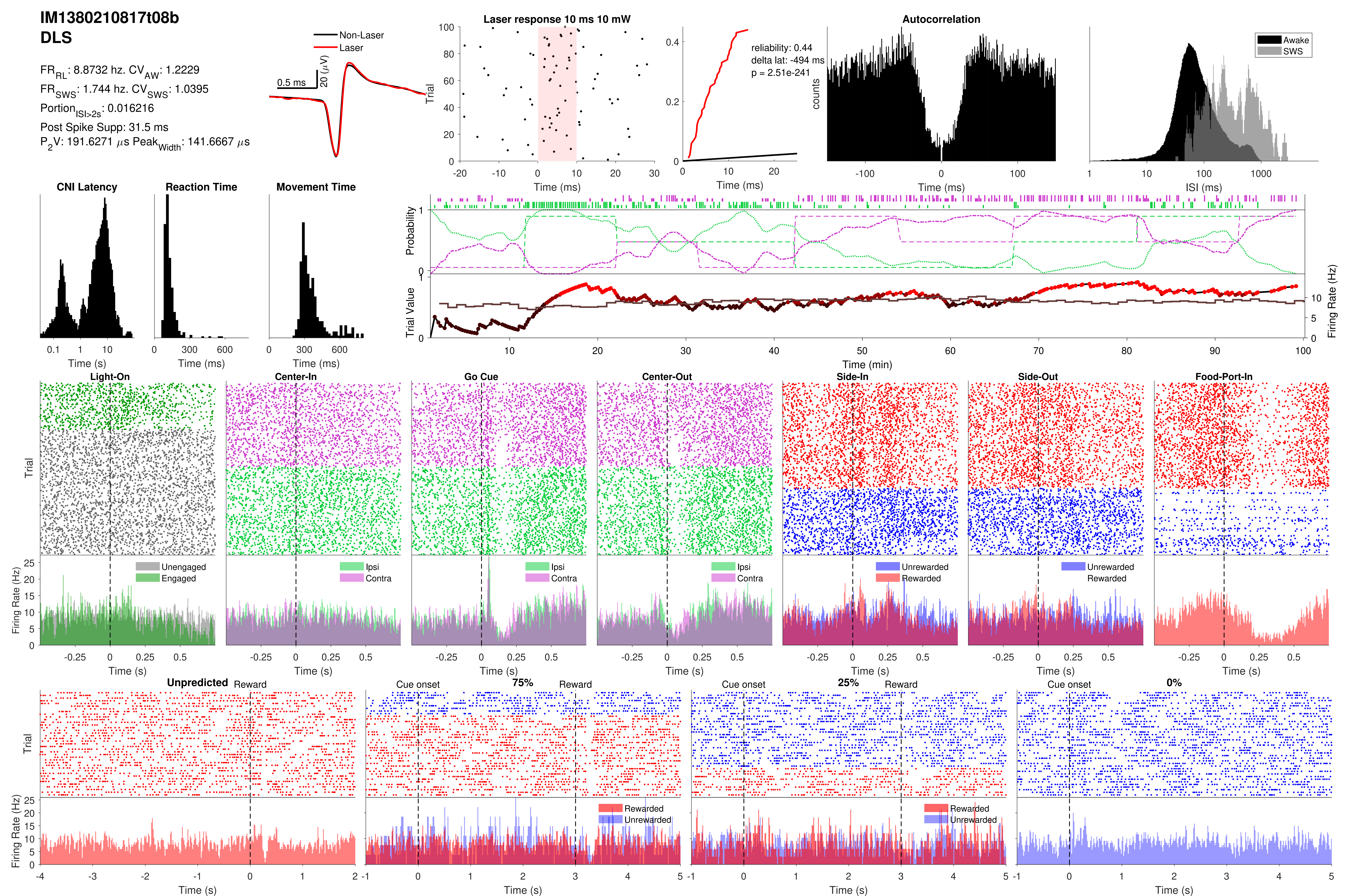
IM1380210817t08b
DLS
FR_RL: 8.8732 hz. CV_AW: 1.2229
FR_SWS: 1.744 hz. CV_SWS: 1.0395
Portion_ISI>2s: 0.016216
Post Spike Supp: 31.5 ms
P_2V: 191.6271 µs Peak_Width: 141.6667 µs
Non-Laser
Laser
0.5 ms
20 (µV)
Laser response 10 ms 10 mW
Trial
Time (ms)
reliability: 0.44
delta lat: -494 ms
p = 2.51e-241
Time (ms)
Autocorrelation
counts
Time (ms)
Awake
SWS
ISI (ms)
CNI Latency
Time (s)
Reaction Time
Time (ms)
Movement Time
Time (ms)
Probability
Trial Value
Firing Rate (Hz)
Time (min)
Light-On
Center-In
Go Cue
Center-Out
Side-In
Side-Out
Food-Port-In
Trial
Firing Rate (Hz)
Unengaged
Engaged
Ipsi
Contra
Unrewarded
Rewarded
Time (s)
Unpredicted
Reward
Cue onset
75%
25%
0%
Rewarded
Unrewarded
Time (s)

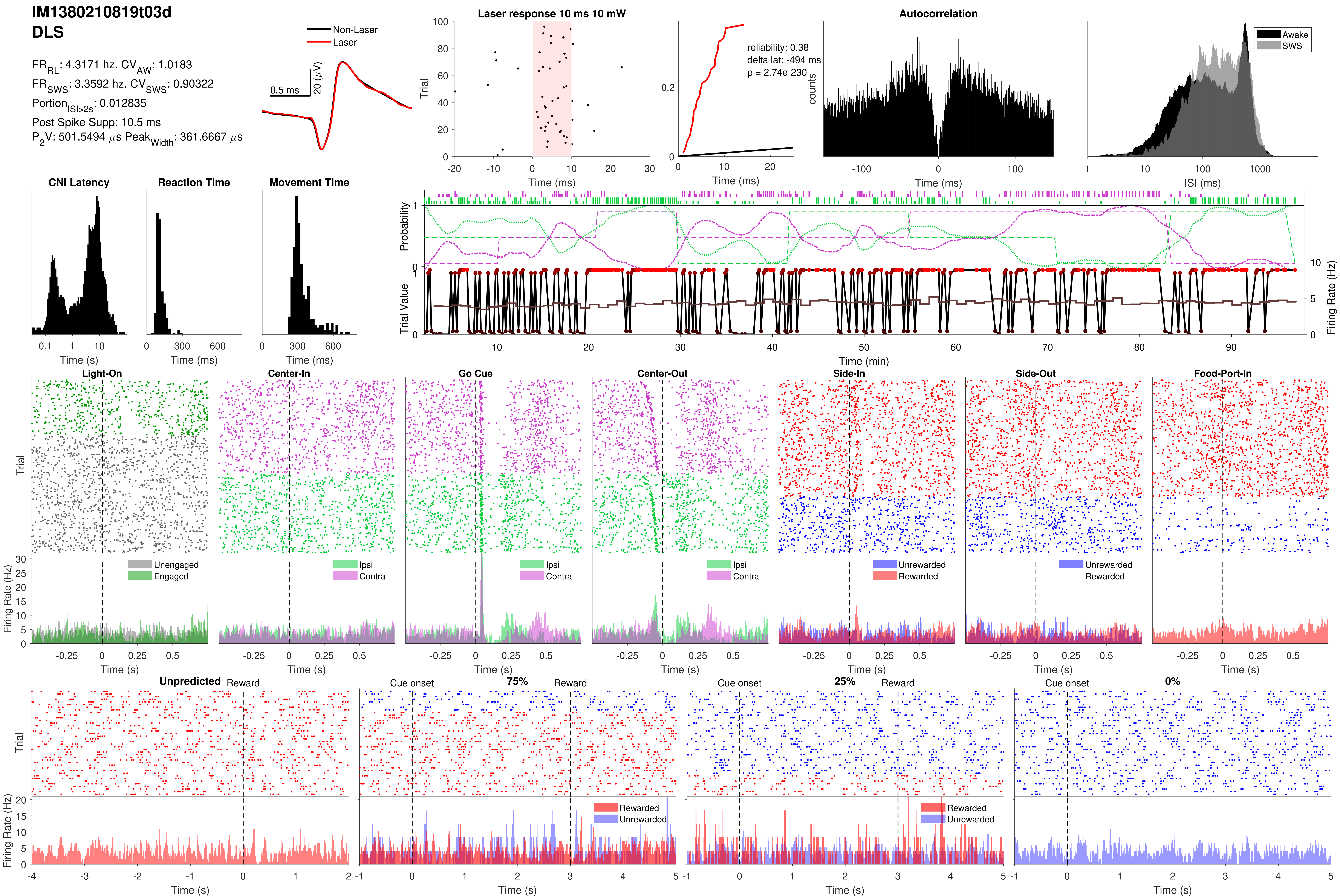
IM1380210819t03d
DLS
FR_RL: 4.3171 hz. CV_AW: 1.0183
FR_SWS: 3.3592 hz. CV_SWS: 0.90322
Portion_ISI>2s: 0.012835
Post Spike Supp: 10.5 ms
P_2V: 501.5494 μs Peak_Width: 361.6667 μs
Non-Laser
Laser
0.5 ms
20 (μV)
Laser response 10 ms 10 mW
Trial
Time (ms)
reliability: 0.38
delta lat: -494 ms
p = 2.74e-230
Autocorrelation
counts
Awake
SWS
ISI (ms)
CNI Latency
Reaction Time
Movement Time
Time (s)
Probability
Trial Value
Firing Rate (Hz)
Time (min)
Light-On
Center-In
Go Cue
Center-Out
Side-In
Side-Out
Food-Port-In
Unengaged
Engaged
Ipsi
Contra
Unrewarded
Rewarded
Unpredicted
Reward
Cue onset
75%
25%
0%

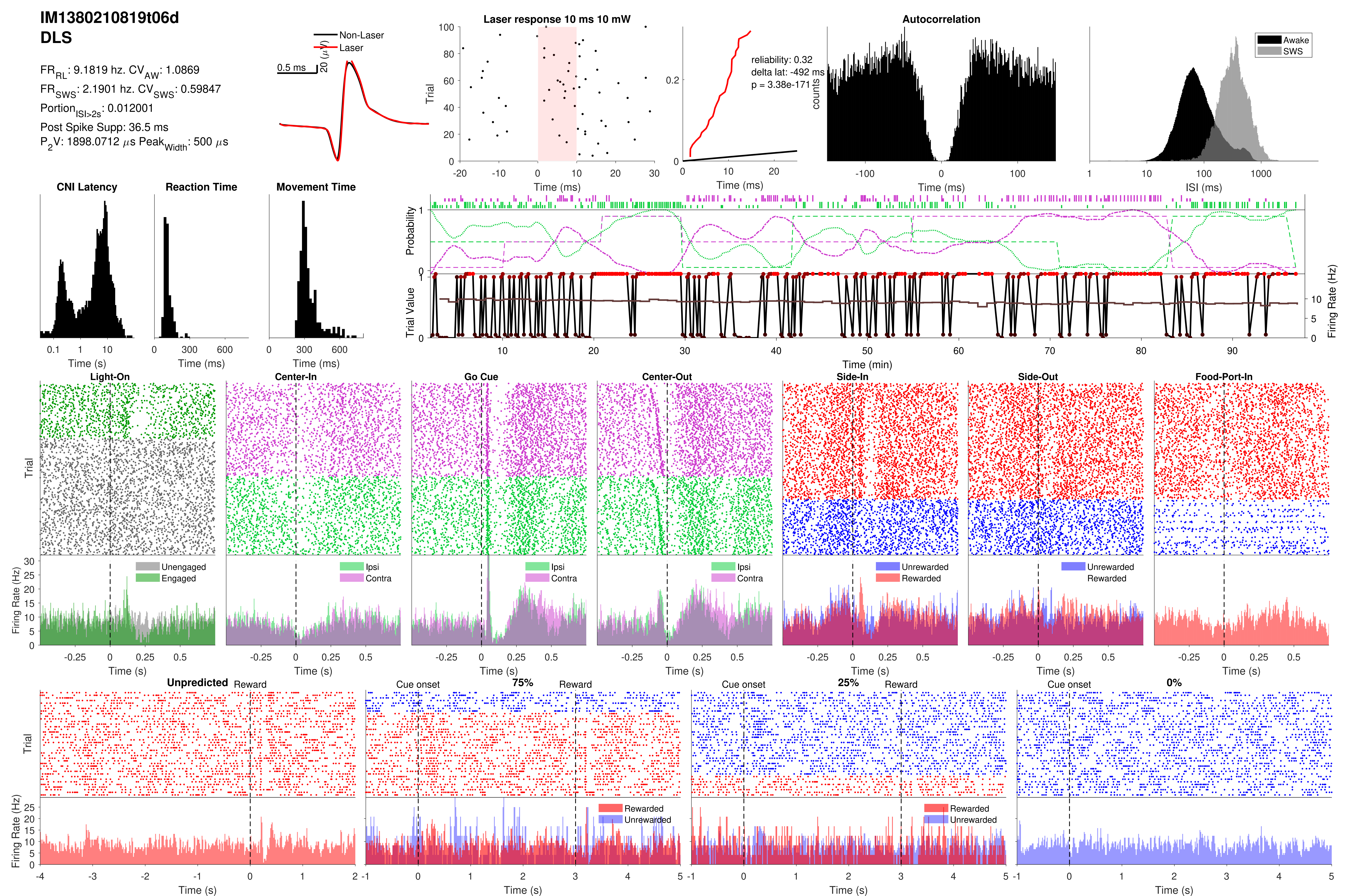
IM1380210819t06d
DLS
$FR_{RL}$: 9.1819 hz. $CV_{AW}$: 1.0869
$FR_{SWS}$: 2.1901 hz. $CV_{SWS}$: 0.59847
$Portion_{ISI>2s}$: 0.012001
Post Spike Supp: 36.5 ms
$P_2V$: 1898.0712 $\mu$s $Peak_{Width}$: 500 $\mu$s
Non-Laser
Laser
0.5 ms
20 ($\mu$V)
Laser response 10 ms 10 mW
Trial
Time (ms)
reliability: 0.32
delta lat: -492 ms
p = 3.38e-171
Time (ms)
Autocorrelation
counts
Time (ms)
Awake
SWS
ISI (ms)
CNI Latency
Time (s)
Reaction Time
Time (ms)
Movement Time
Time (ms)
Probability
Trial Value
Firing Rate (Hz)
Time (min)
Light-On
Center-In
Go Cue
Center-Out
Side-In
Side-Out
Food-Port-In
Trial
Firing Rate (Hz)
Unengaged
Engaged
Ipsi
Contra
Unrewarded
Rewarded
Time (s)
Unpredicted
Reward
Cue onset
75%
25%
0%
Rewarded
Unrewarded
Time (s)

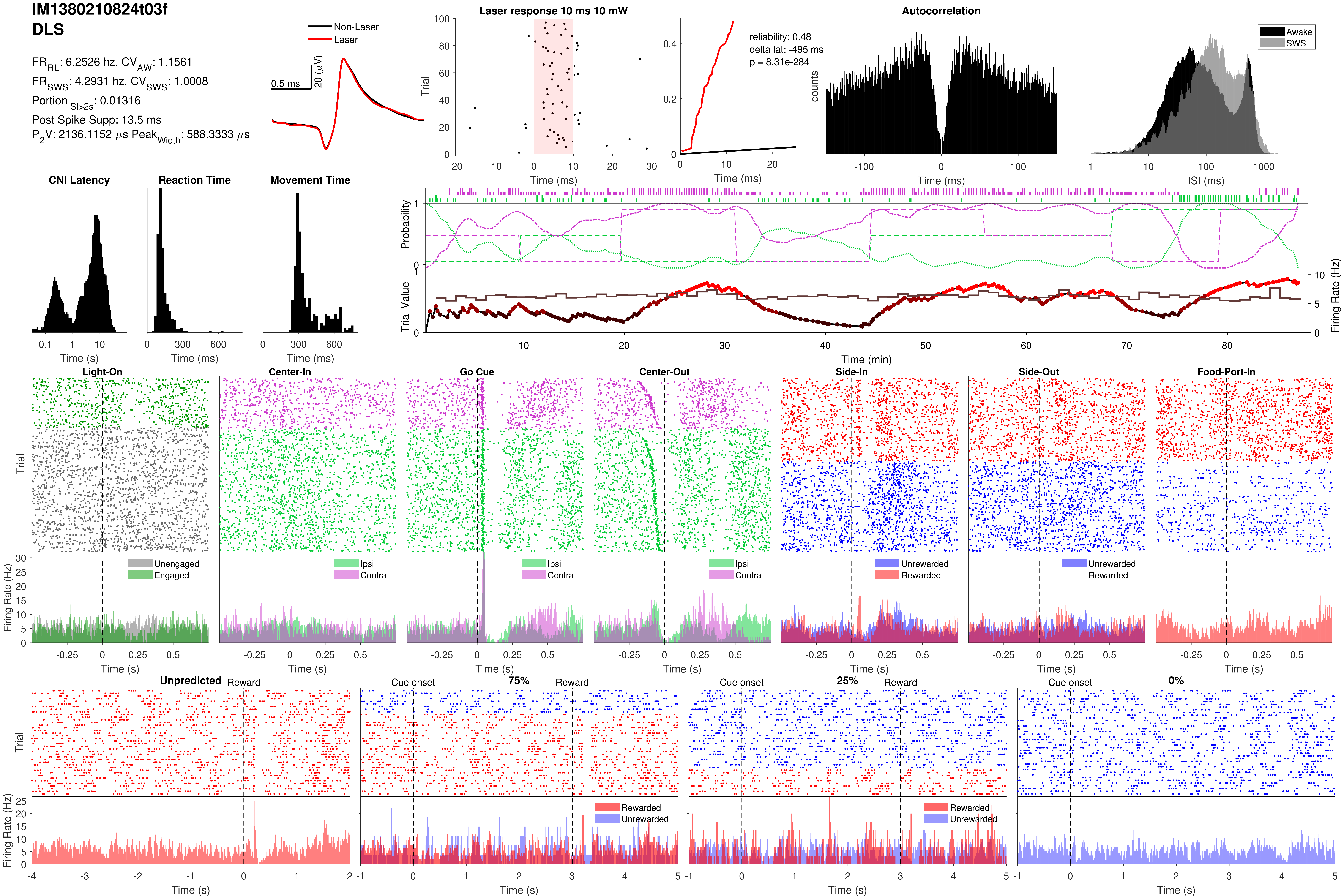
IM1380210824t03f
DLS
$FR_{RL}$: 6.2526 hz. $CV_{AW}$: 1.1561
$FR_{SWS}$: 4.2931 hz. $CV_{SWS}$: 1.0008
$Portion_{ISI>2s}$: 0.01316
Post Spike Supp: 13.5 ms
$P_2V$: 2136.1152 $\mu$s $Peak_{Width}$: 588.3333 $\mu$s
Non-Laser
Laser
0.5 ms
20 ($\mu$V)
Laser response 10 ms 10 mW
Trial
Time (ms)
reliability: 0.48
delta lat: -495 ms
p = 8.31e-284
Time (ms)
Autocorrelation
counts
Time (ms)
Awake
SWS
ISI (ms)
CNI Latency
Time (s)
Reaction Time
Time (ms)
Movement Time
Time (ms)
Probability
Trial Value
Firing Rate (Hz)
Time (min)
Light-On
Center-In
Go Cue
Center-Out
Side-In
Side-Out
Food-Port-In
Trial
Firing Rate (Hz)
Unengaged
Engaged
Ipsi
Contra
Unrewarded
Rewarded
Time (s)
Unpredicted
Reward
Cue onset
75%
25%
0%
Rewarded
Unrewarded
Trial
Firing Rate (Hz)
Time (s)

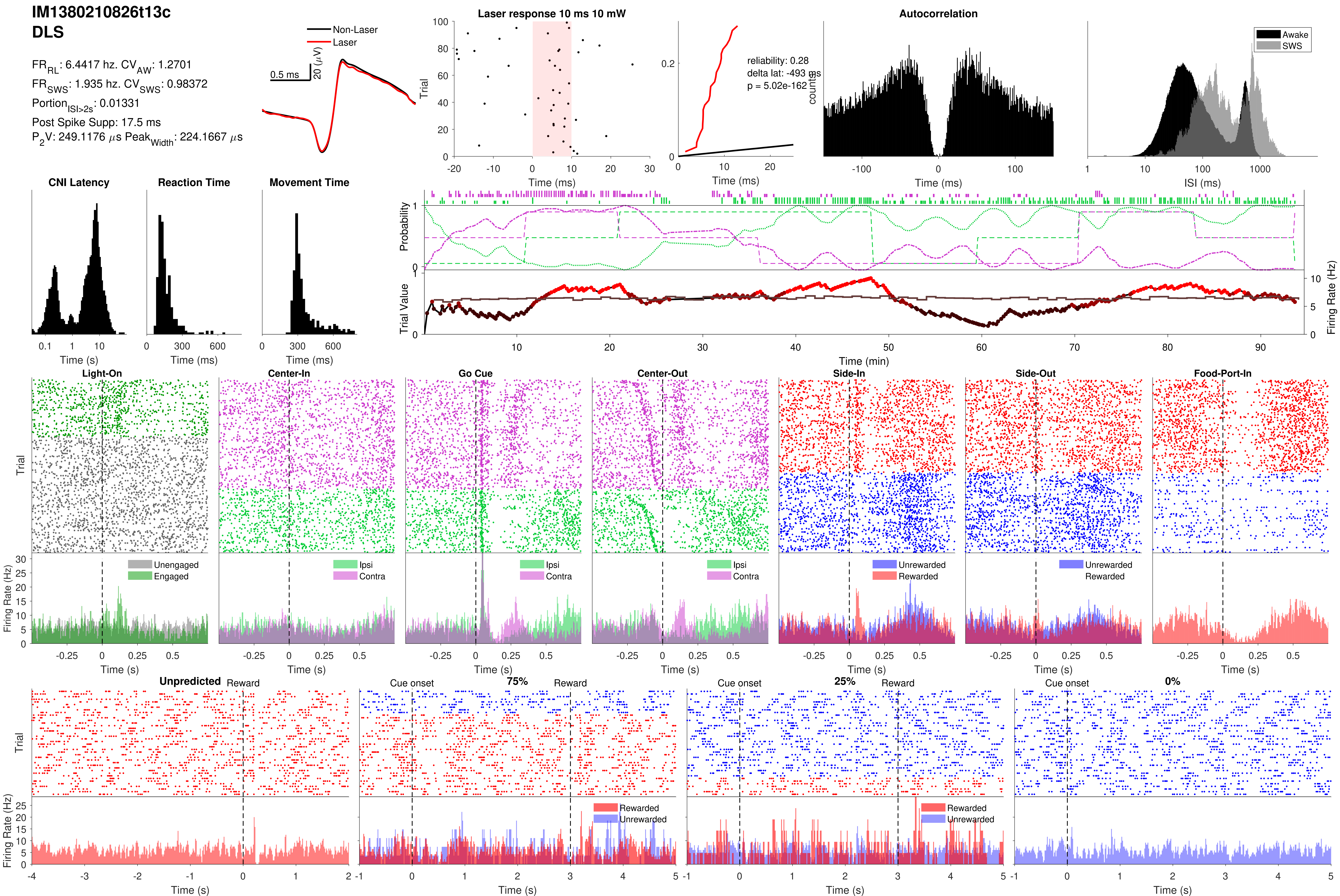
IM1380210826t13c
DLS
FR_RL: 6.4417 hz. CV_AW: 1.2701
FR_SWS: 1.935 hz. CV_SWS: 0.98372
Portion_ISI>2s: 0.01331
Post Spike Supp: 17.5 ms
P_2V: 249.1176 μs Peak_Width: 224.1667 μs
Non-Laser
Laser
0.5 ms
20 (μV)
Laser response 10 ms 10 mW
Trial
Time (ms)
reliability: 0.28
delta lat: -493 ms
p = 5.02e-162
counts
Autocorrelation
Awake
SWS
ISI (ms)
CNI Latency
Reaction Time
Movement Time
Time (s)
Time (ms)
Probability
Trial Value
Firing Rate (Hz)
Time (min)
Light-On
Center-In
Go Cue
Center-Out
Side-In
Side-Out
Food-Port-In
Unengaged
Engaged
Ipsi
Contra
Unrewarded
Rewarded
Unpredicted
Reward
Cue onset
75%
25%
0%
Time (s)

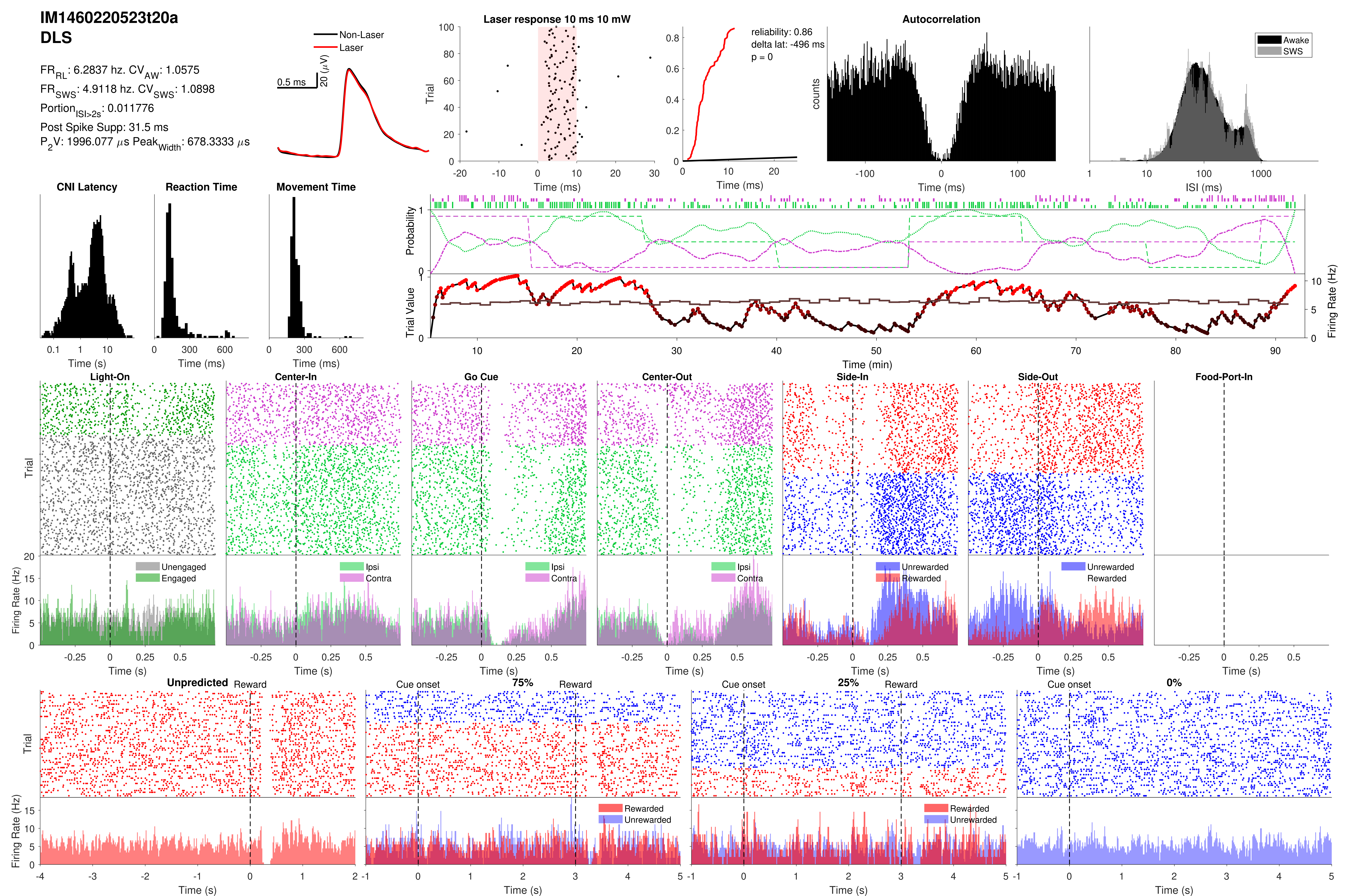

IM1460220523t20a
DLS
FR$_{RL}$: 6.2837 hz. CV$_{AW}$: 1.0575
FR$_{SWS}$: 4.9118 hz. CV$_{SWS}$: 1.0898
Portion$_{ISI>2s}$: 0.011776
Post Spike Supp: 31.5 ms
P$_2$V: 1996.077 $\mu$s Peak$_{Width}$: 678.3333 $\mu$s
Non-Laser
Laser
0.5 ms
20 ($\mu$V)
Laser response 10 ms 10 mW
Trial
Time (ms)
reliability: 0.86
delta lat: -496 ms
p = 0
Time (ms)
Autocorrelation
counts
Time (ms)
Awake
SWS
ISI (ms)
CNI Latency
Time (s)
Reaction Time
Time (ms)
Movement Time
Time (ms)
Probability
Trial Value
Firing Rate (Hz)
Time (min)
Light-On
Center-In
Go Cue
Center-Out
Side-In
Side-Out
Food-Port-In
Trial
Firing Rate (Hz)
Unengaged
Engaged
Ipsi
Contra
Unrewarded
Rewarded
Time (s)
Unpredicted
Reward
Cue onset
75%
25%
0%
Rewarded
Unrewarded
Time (s)

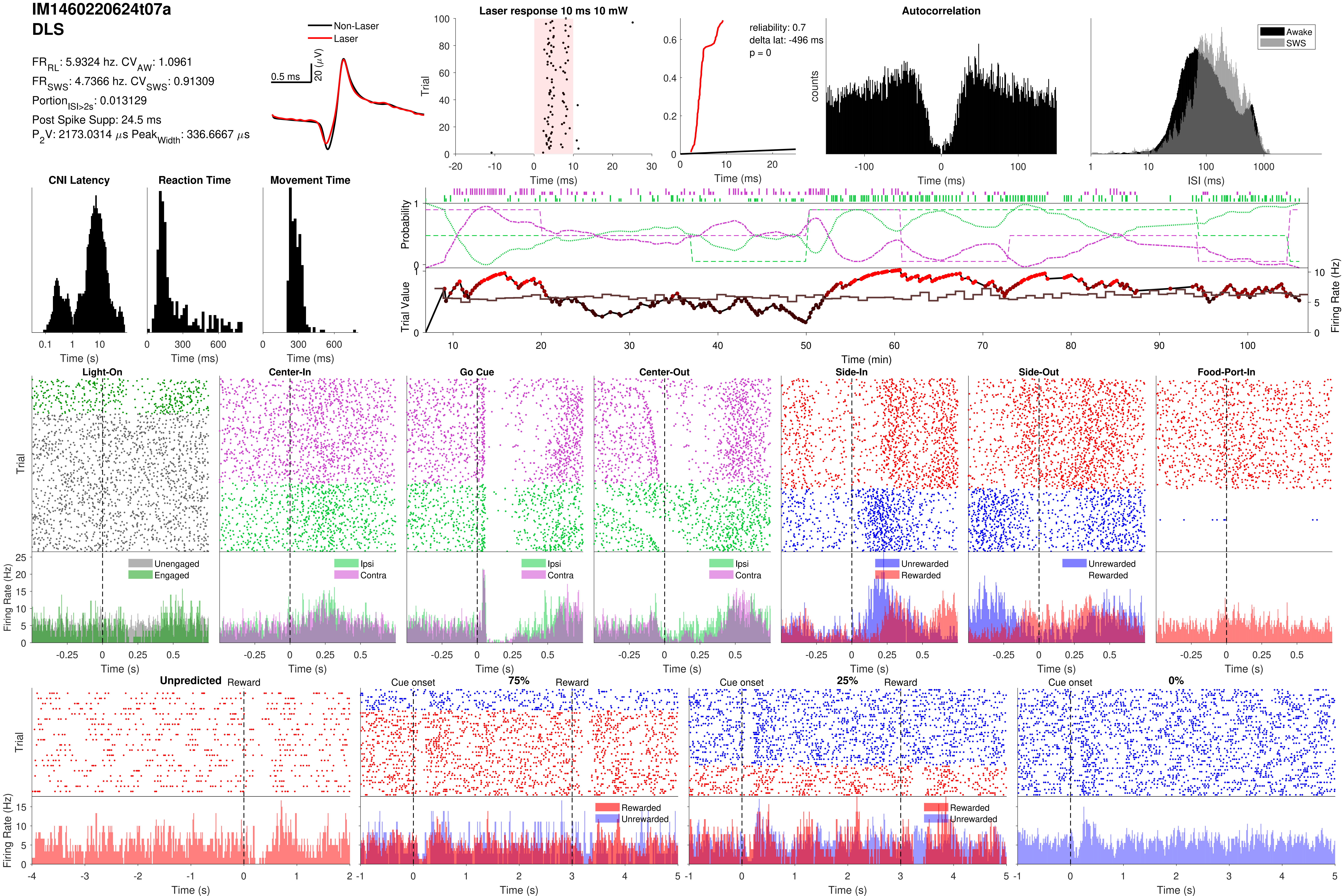

IM1460220624t07a
DLS
FR_RL: 5.9324 hz. CV_AW: 1.0961
FR_SWS: 4.7366 hz. CV_SWS: 0.91309
Portion_ISI>2s: 0.013129
Post Spike Supp: 24.5 ms
P2V: 2173.0314 µs Peak_Width: 336.6667 µs
Non-Laser
Laser
0.5 ms
20 (µV)
Laser response 10 ms 10 mW
Trial
Time (ms)
reliability: 0.7
delta lat: -496 ms
p = 0
Time (ms)
Autocorrelation
counts
Time (ms)
Awake
SWS
ISI (ms)
CNI Latency
Time (s)
Reaction Time
Time (ms)
Movement Time
Time (ms)
Probability
Trial Value
Firing Rate (Hz)
Time (min)
Light-On
Center-In
Go Cue
Center-Out
Side-In
Side-Out
Food-Port-In
Trial
Firing Rate (Hz)
Unengaged
Engaged
Ipsi
Contra
Unrewarded
Rewarded
Time (s)
Unpredicted
Reward
Cue onset
75%
25%
0%
Rewarded
Unrewarded
Trial
Firing Rate (Hz)
Time (s)

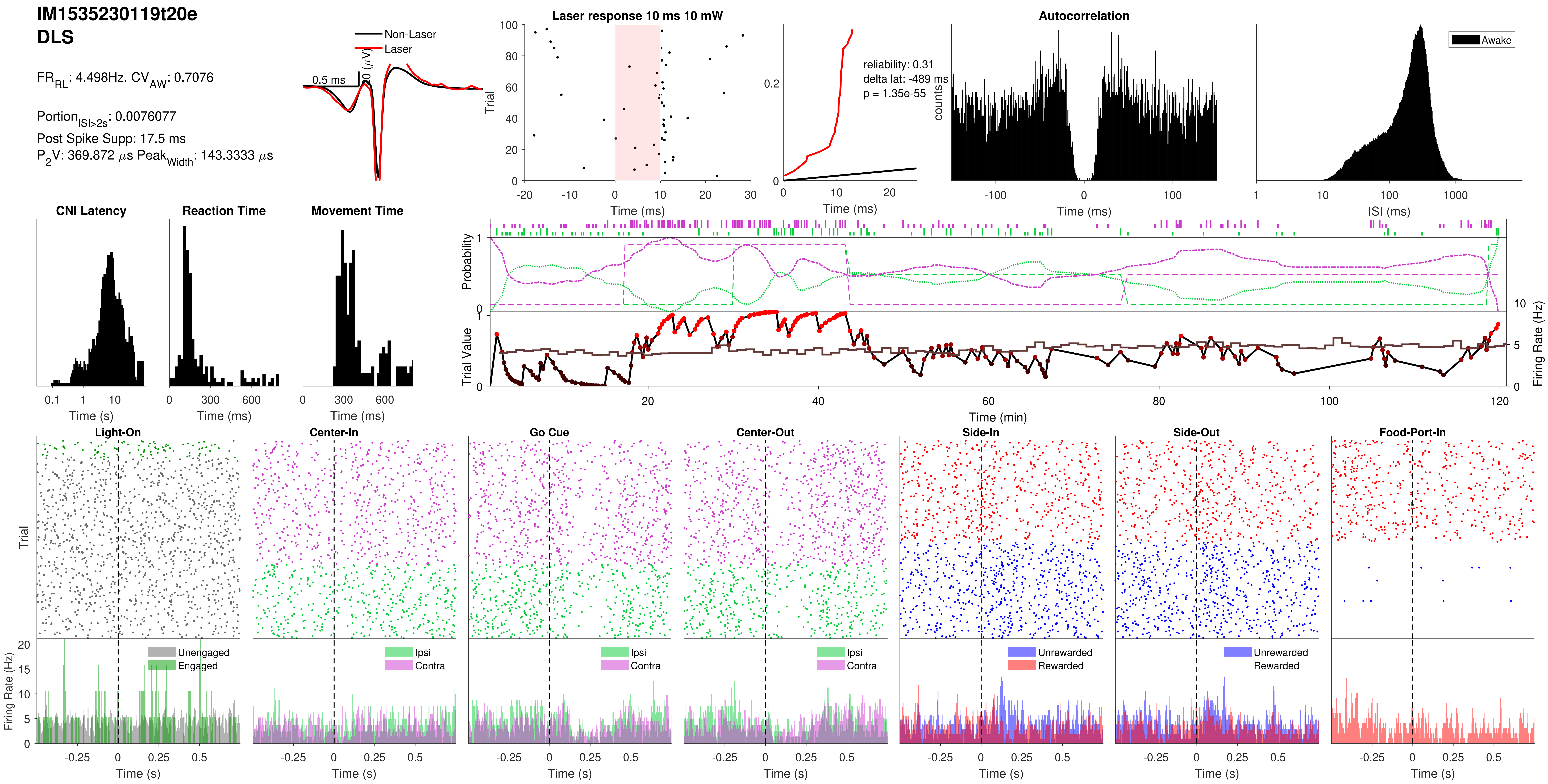
IM1535230119t20e
DLS
FR_RL: 4.498Hz. CV_AW: 0.7076
Portion_ISI>2s: 0.0076077
Post Spike Supp: 17.5 ms
P_2V: 369.872 µs Peak_Width: 143.3333 µs
Non-Laser
Laser
0.5 ms
0 (µV)
Laser response 10 ms 10 mW
Trial
Time (ms)
reliability: 0.31
delta lat: -489 ms
p = 1.35e-55
Time (ms)
Autocorrelation
counts
Time (ms)
Awake
ISI (ms)
CNI Latency
Time (s)
Reaction Time
Time (ms)
Movement Time
Time (ms)
Probability
Trial Value
Firing Rate (Hz)
Time (min)
Light-On
Center-In
Go Cue
Center-Out
Side-In
Side-Out
Food-Port-In
Trial
Firing Rate (Hz)
Unengaged
Engaged
Ipsi
Contra
Unrewarded
Rewarded
Time (s)

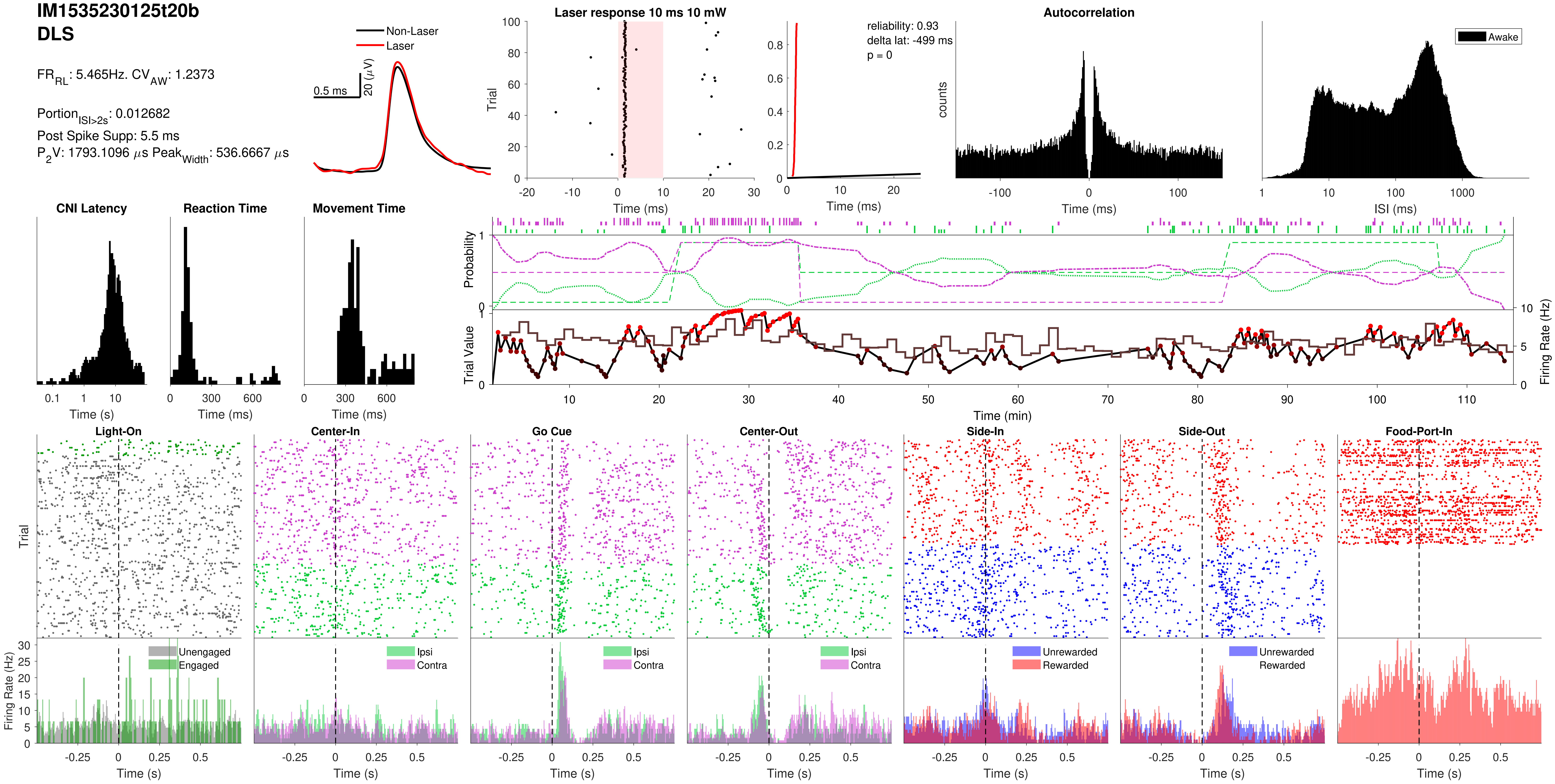

IM1535230125t20b
DLS
FR_RL: 5.465Hz. CV_AW: 1.2373
Portion_ISI>2s: 0.012682
Post Spike Supp: 5.5 ms
P2V: 1793.1096 μs Peak_Width: 536.6667 μs
Non-Laser
Laser
0.5 ms
20 (μV)
Laser response 10 ms 10 mW
Trial
Time (ms)
reliability: 0.93
delta lat: -499 ms
p = 0
Time (ms)
Autocorrelation
counts
Time (ms)
Awake
ISI (ms)
CNI Latency
Time (s)
Reaction Time
Time (ms)
Movement Time
Time (ms)
Probability
Trial Value
Firing Rate (Hz)
Time (min)
Light-On
Center-In
Go Cue
Center-Out
Side-In
Side-Out
Food-Port-In
Trial
Firing Rate (Hz)
Unengaged
Engaged
Ipsi
Contra
Unrewarded
Rewarded
Time (s)

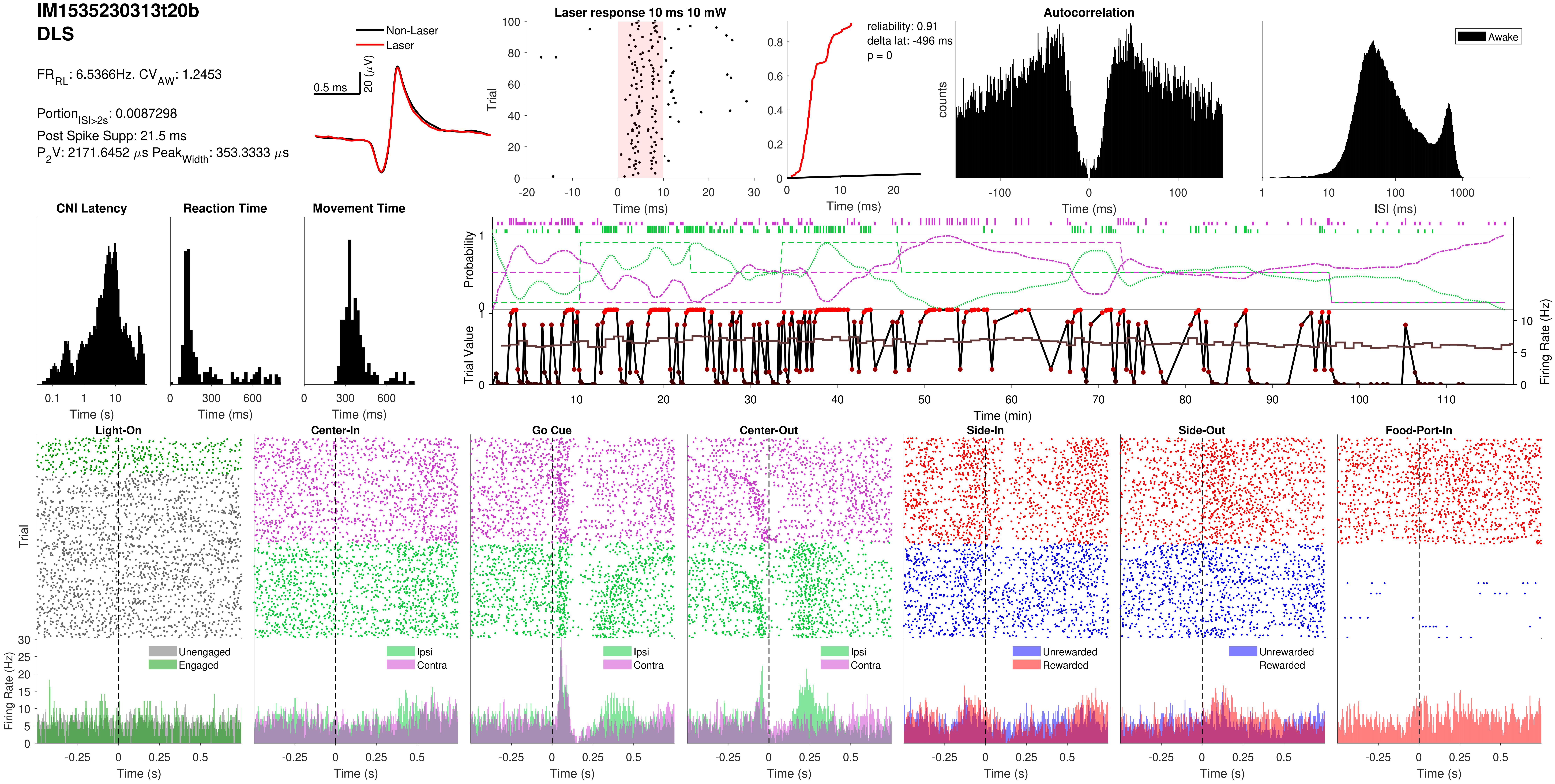

IM1535230313t20b
DLS
FR_RL: 6.5366Hz. CV_AW: 1.2453
Portion_ISI>2s: 0.0087298
Post Spike Supp: 21.5 ms
P_2V: 2171.6452 µs Peak_Width: 353.3333 µs
Non-Laser
Laser
0.5 ms
20 (µV)
Laser response 10 ms 10 mW
Trial
Time (ms)
reliability: 0.91
delta lat: -496 ms
p = 0
Autocorrelation
counts
Awake
ISI (ms)
CNI Latency
Reaction Time
Movement Time
Time (s)
Probability
Trial Value
Firing Rate (Hz)
Time (min)
Light-On
Center-In
Go Cue
Center-Out
Side-In
Side-Out
Food-Port-In
Unengaged
Engaged
Ipsi
Contra
Unrewarded
Rewarded

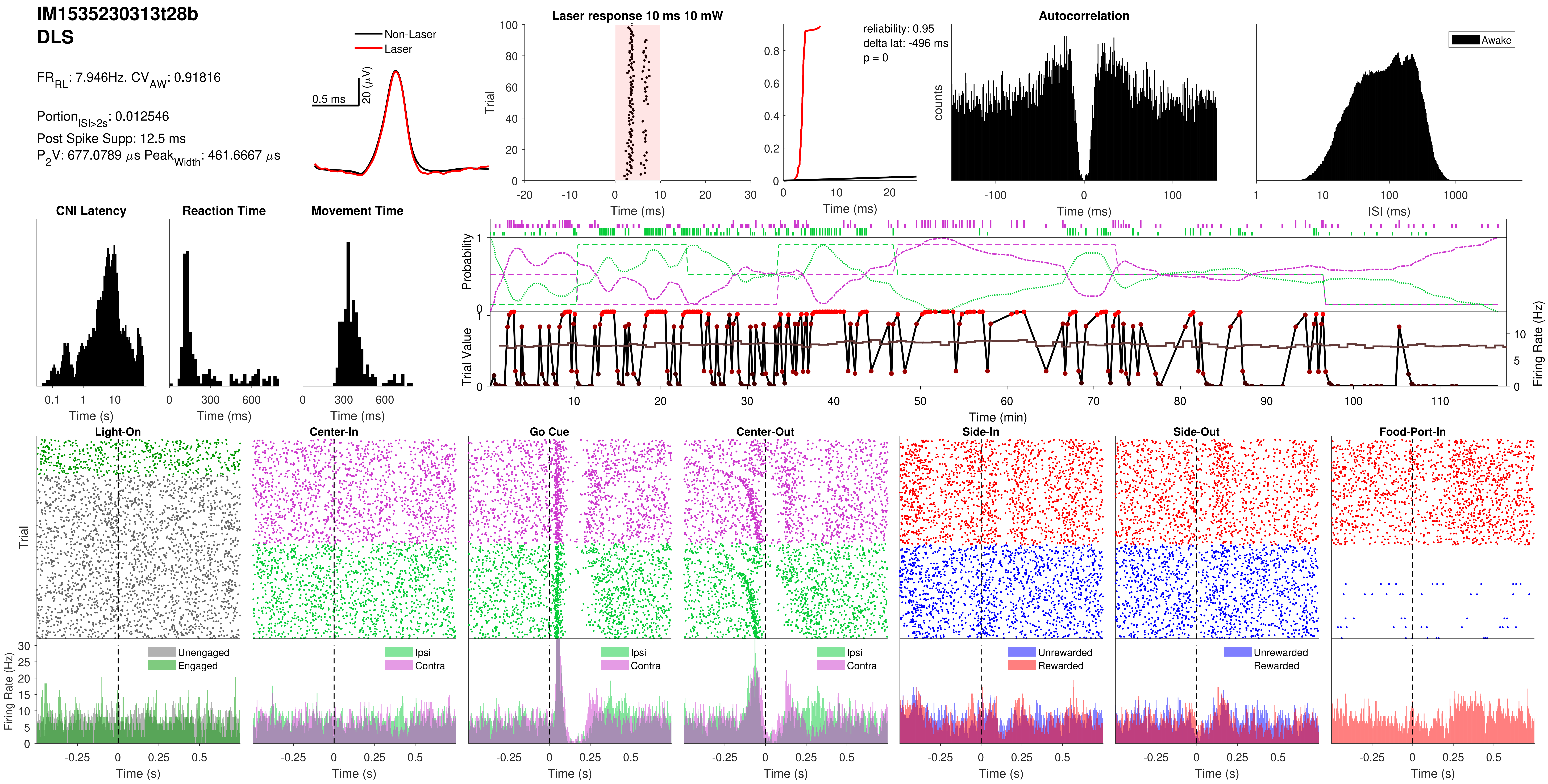
IM1535230313t28b
DLS
FR_RL: 7.946Hz. CV_AW: 0.91816
Portion_ISI>2s: 0.012546
Post Spike Supp: 12.5 ms
P_2V: 677.0789 µs Peak_Width: 461.6667 µs
Non-Laser
Laser
0.5 ms
20 (µV)
Laser response 10 ms 10 mW
Trial
Time (ms)
reliability: 0.95
delta lat: -496 ms
p = 0
Time (ms)
Autocorrelation
counts
Time (ms)
Awake
ISI (ms)
CNI Latency
Time (s)
Reaction Time
Time (ms)
Movement Time
Time (ms)
Probability
Trial Value
Firing Rate (Hz)
Time (min)
Light-On
Center-In
Go Cue
Center-Out
Side-In
Side-Out
Food-Port-In
Trial
Firing Rate (Hz)
Unengaged
Engaged
Ipsi
Contra
Ipsi
Contra
Ipsi
Contra
Unrewarded
Rewarded
Unrewarded
Rewarded
Time (s)
Time (s)
Time (s)
Time (s)
Time (s)
Time (s)
Time (s)

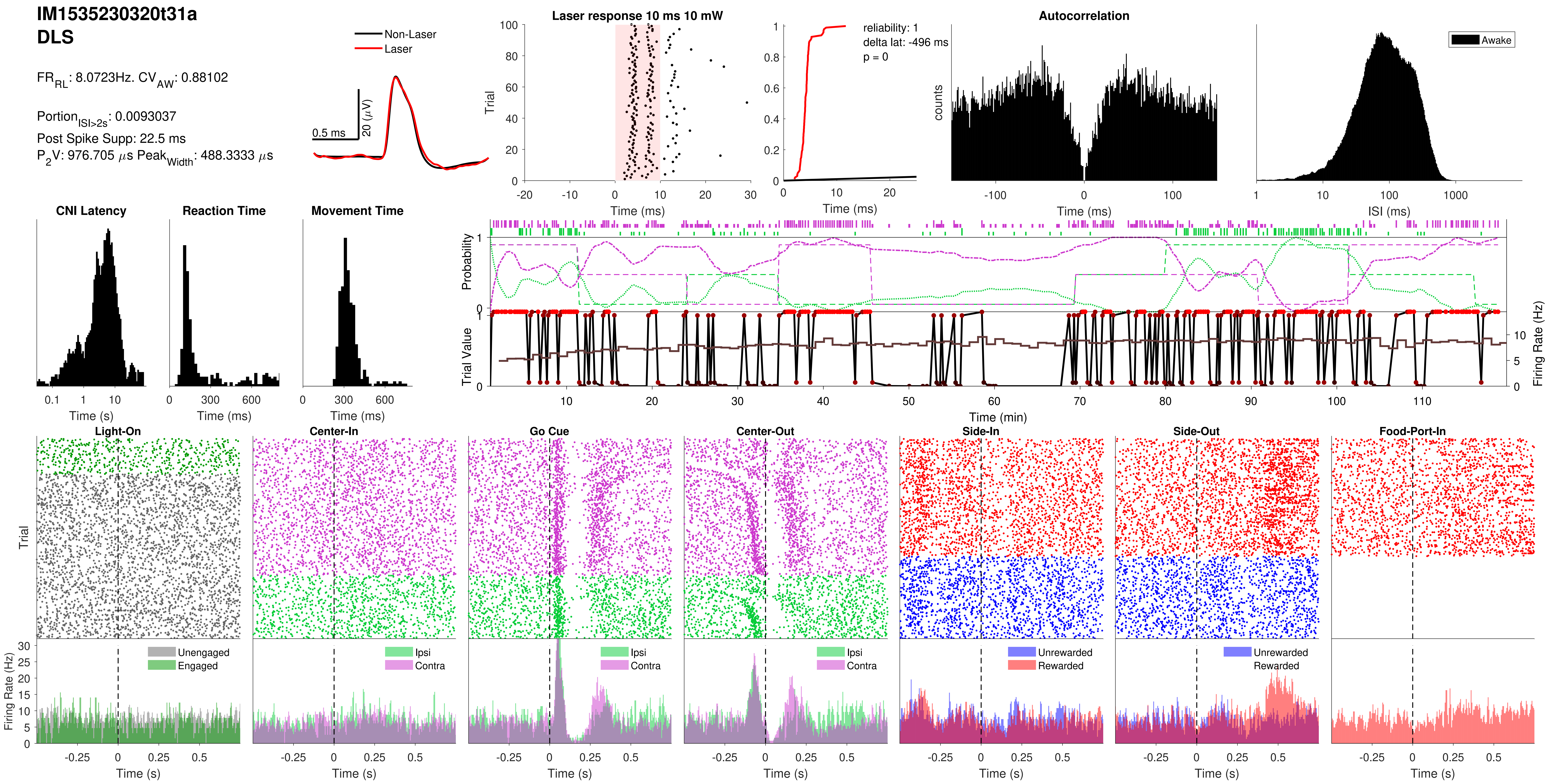
IM1535230320t31a
DLS
FR_RL: 8.0723Hz. CV_AW: 0.88102
Portion_ISI>2s: 0.0093037
Post Spike Supp: 22.5 ms
P_2V: 976.705 µs Peak_Width: 488.3333 µs
Non-Laser
Laser
0.5 ms
20 (µV)
Laser response 10 ms 10 mW
Trial
Time (ms)
reliability: 1
delta lat: -496 ms
p = 0
Autocorrelation
counts
Awake
ISI (ms)
CNI Latency
Time (s)
Reaction Time
Time (ms)
Movement Time
Probability
Trial Value
Firing Rate (Hz)
Time (min)
Light-On
Center-In
Go Cue
Center-Out
Side-In
Side-Out
Food-Port-In
Unengaged
Engaged
Ipsi
Contra
Unrewarded
Rewarded
Time (s)

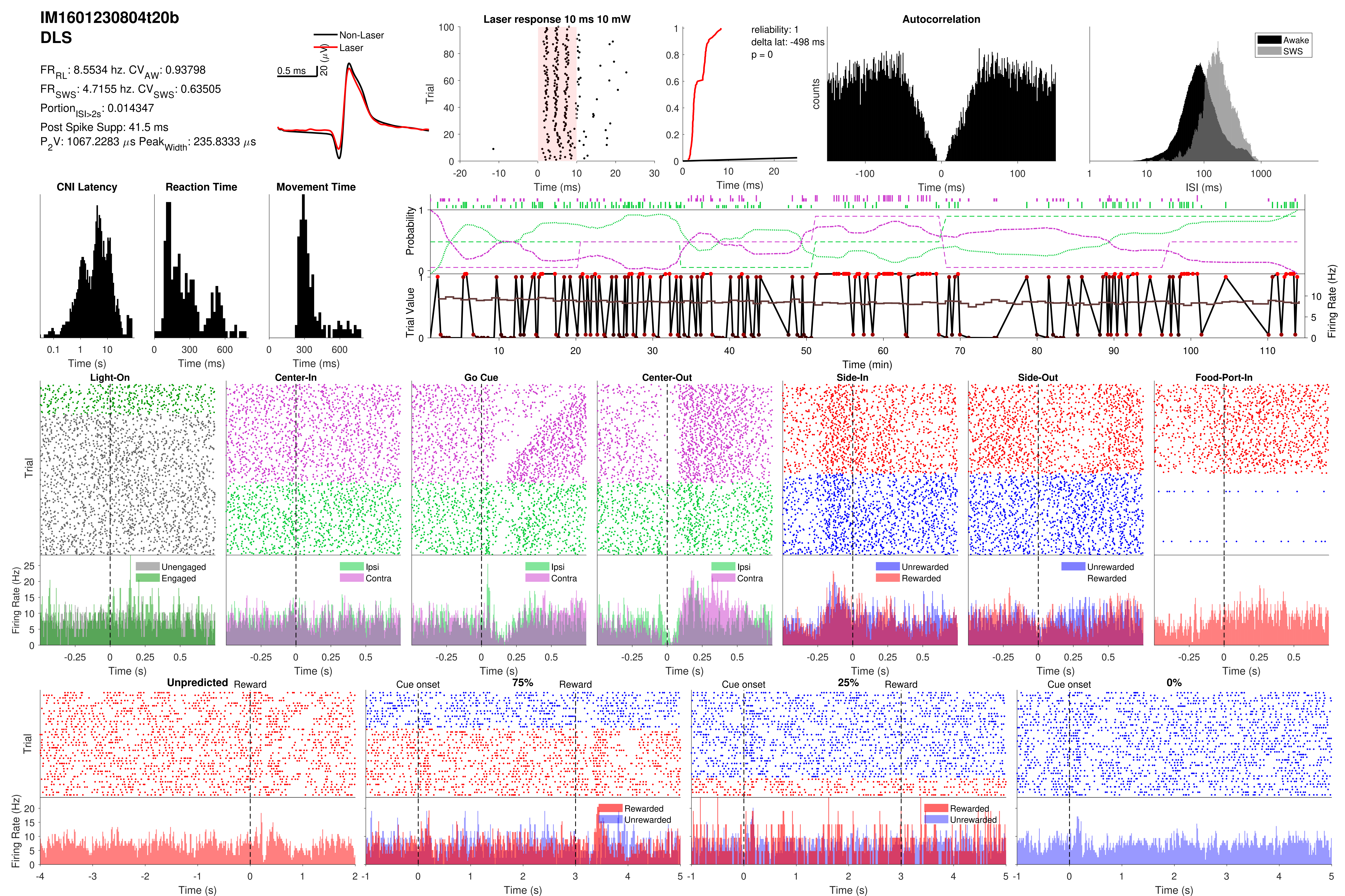
IM1601230804t20b
DLS
FR_RL: 8.5534 hz. CV_AW: 0.93798
FR_SWS: 4.7155 hz. CV_SWS: 0.63505
Portion_ISI>2s: 0.014347
Post Spike Supp: 41.5 ms
P2V: 1067.2283 µs Peak_Width: 235.8333 µs
Non-Laser
Laser
0.5 ms
20 (µV)
Laser response 10 ms 10 mW
reliability: 1
delta lat: -498 ms
p = 0
Autocorrelation
Awake
SWS
CNI Latency
Reaction Time
Movement Time
Probability
Trial Value
Firing Rate (Hz)
Time (min)
Light-On
Center-In
Go Cue
Center-Out
Side-In
Side-Out
Food-Port-In
Unengaged
Engaged
Ipsi
Contra
Unrewarded
Rewarded
Unpredicted
Reward
Cue onset
75%
25%
0%
Trial
Time (s)
Time (ms)
ISI (ms)
counts

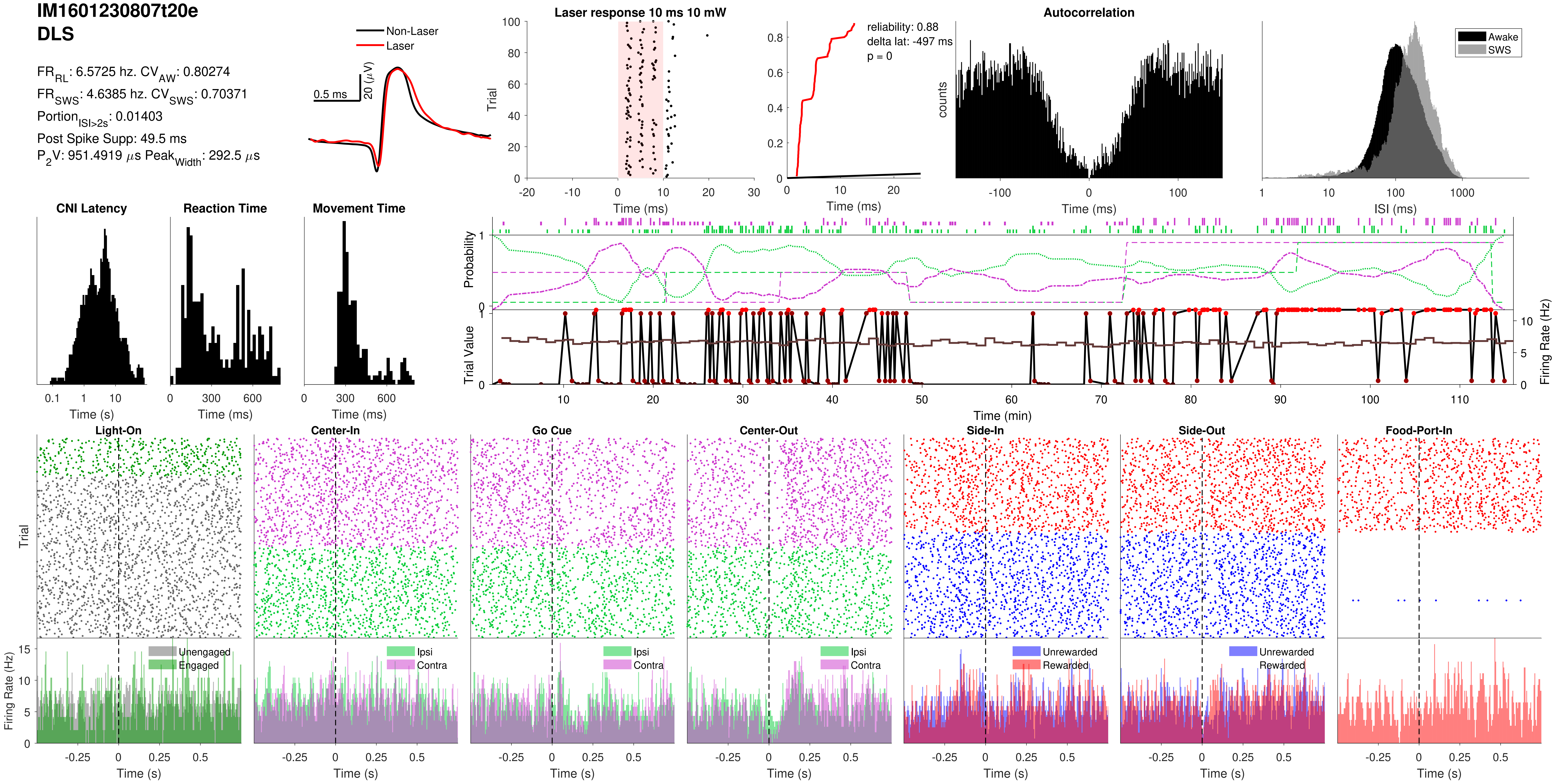

IM1601230807t20e
DLS
$FR_{RL}$: 6.5725 hz. $CV_{AW}$: 0.80274
$FR_{SWS}$: 4.6385 hz. $CV_{SWS}$: 0.70371
$Portion_{ISI>2s}$: 0.01403
Post Spike Supp: 49.5 ms
$P_2V$: 951.4919 $\mu$s $Peak_{Width}$: 292.5 $\mu$s
Non-Laser
Laser
0.5 ms
20 ($\mu$V)
Laser response 10 ms 10 mW
Trial
Time (ms)
reliability: 0.88
delta lat: -497 ms
p = 0
Autocorrelation
counts
Awake
SWS
ISI (ms)
CNI Latency
Reaction Time
Movement Time
Time (s)
Time (ms)
Probability
Trial Value
Firing Rate (Hz)
Time (min)
Light-On
Center-In
Go Cue
Center-Out
Side-In
Side-Out
Food-Port-In
Unengaged
Engaged
Ipsi
Contra
Unrewarded
Rewarded

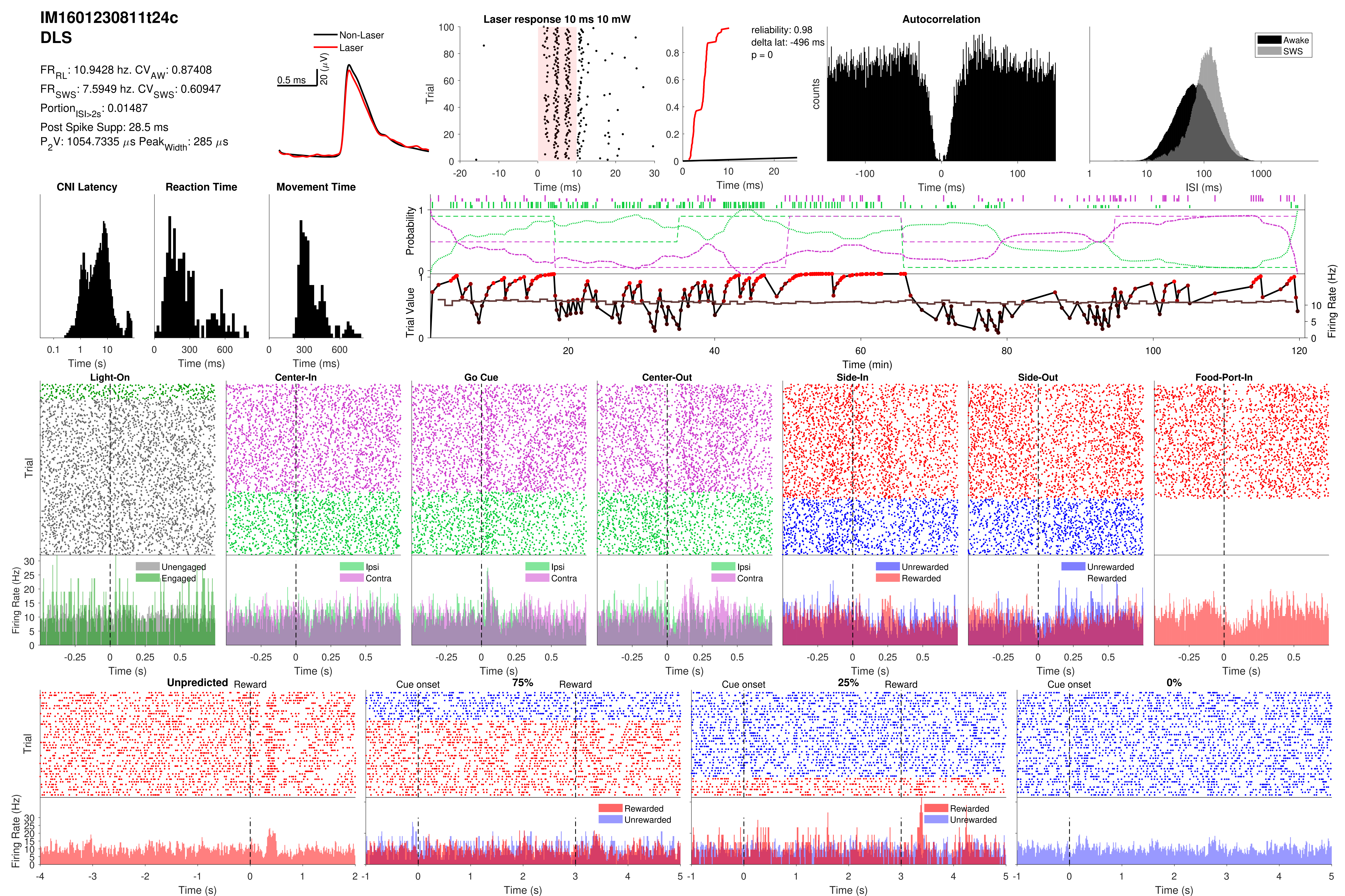
IM1601230811t24c
DLS
FR_RL: 10.9428 hz. CV_AW: 0.87408
FR_SWS: 7.5949 hz. CV_SWS: 0.60947
Portion_ISI>2s: 0.01487
Post Spike Supp: 28.5 ms
P_2V: 1054.7335 µs Peak_Width: 285 µs
Non-Laser
Laser
0.5 ms
20 (µV)
Laser response 10 ms 10 mW
Trial
Time (ms)
reliability: 0.98
delta lat: -496 ms
p = 0
Time (ms)
Autocorrelation
counts
Time (ms)
Awake
SWS
ISI (ms)
CNI Latency
Time (s)
Reaction Time
Time (ms)
Movement Time
Time (ms)
Probability
Trial Value
Firing Rate (Hz)
Time (min)
Light-On
Center-In
Go Cue
Center-Out
Side-In
Side-Out
Food-Port-In
Trial
Firing Rate (Hz)
Unengaged
Engaged
Ipsi
Contra
Unrewarded
Rewarded
Time (s)
Unpredicted
Reward
Cue onset
75%
25%
0%
Trial
Firing Rate (Hz)
Rewarded
Unrewarded
Time (s)

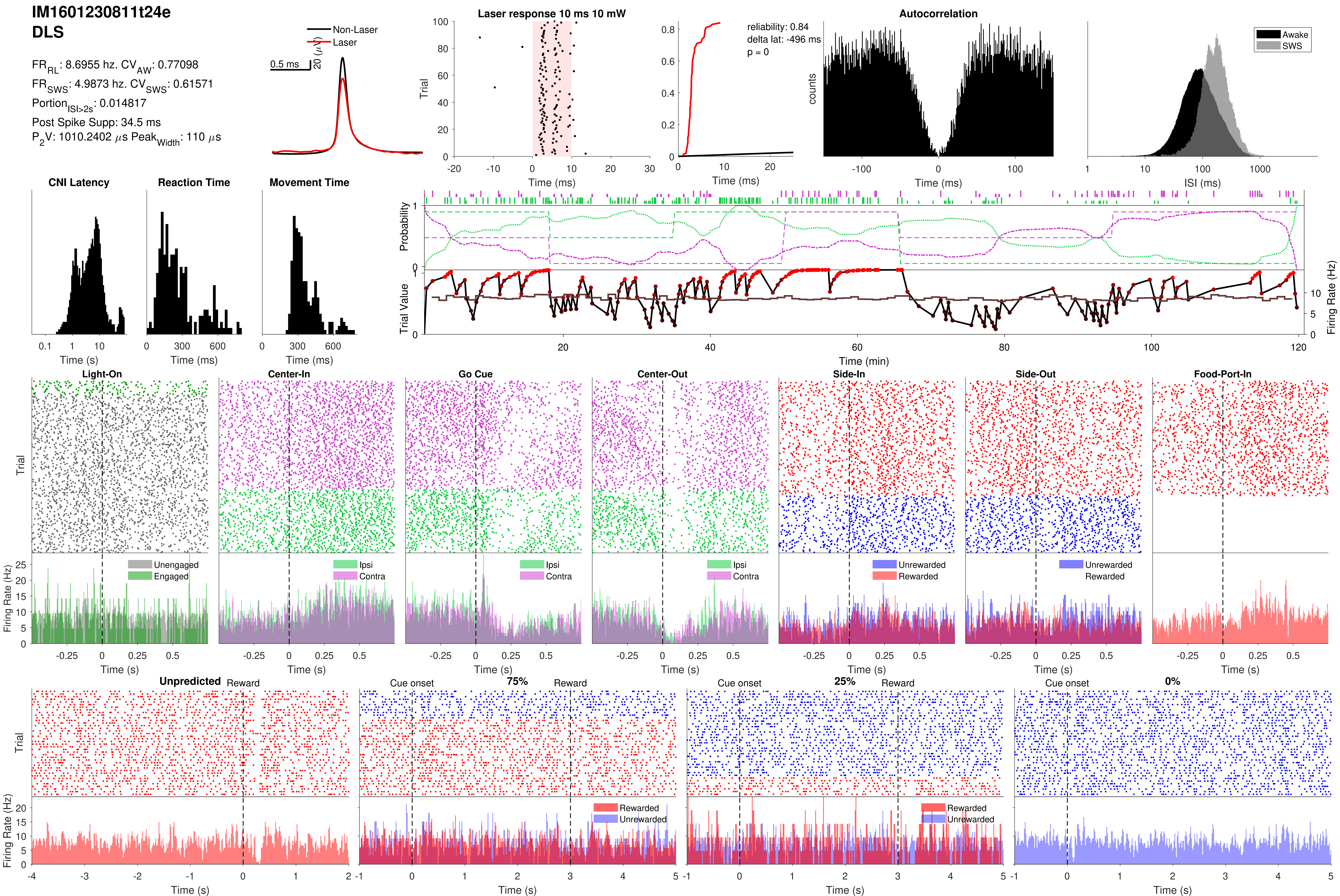

IM1601230811t24e
DLS
FR_RL: 8.6955 hz. CV_AW: 0.77098
FR_SWS: 4.9873 hz. CV_SWS: 0.61571
Portion_ISI>2s: 0.014817
Post Spike Supp: 34.5 ms
P_2V: 1010.2402 μs Peak_Width: 110 μs
Non-Laser
Laser
0.5 ms
20 (μV)
Laser response 10 ms 10 mW
reliability: 0.84
delta lat: -496 ms
p = 0
Autocorrelation
Awake
SWS
CNI Latency
Reaction Time
Movement Time
Probability
Trial Value
Firing Rate (Hz)
Time (min)
Light-On
Center-In
Go Cue
Center-Out
Side-In
Side-Out
Food-Port-In
Unengaged
Engaged
Ipsi
Contra
Unrewarded
Rewarded
Unpredicted
Reward
Cue onset
75%
25%
0%
Trial
Time (ms)
Time (s)
ISI (ms)
counts

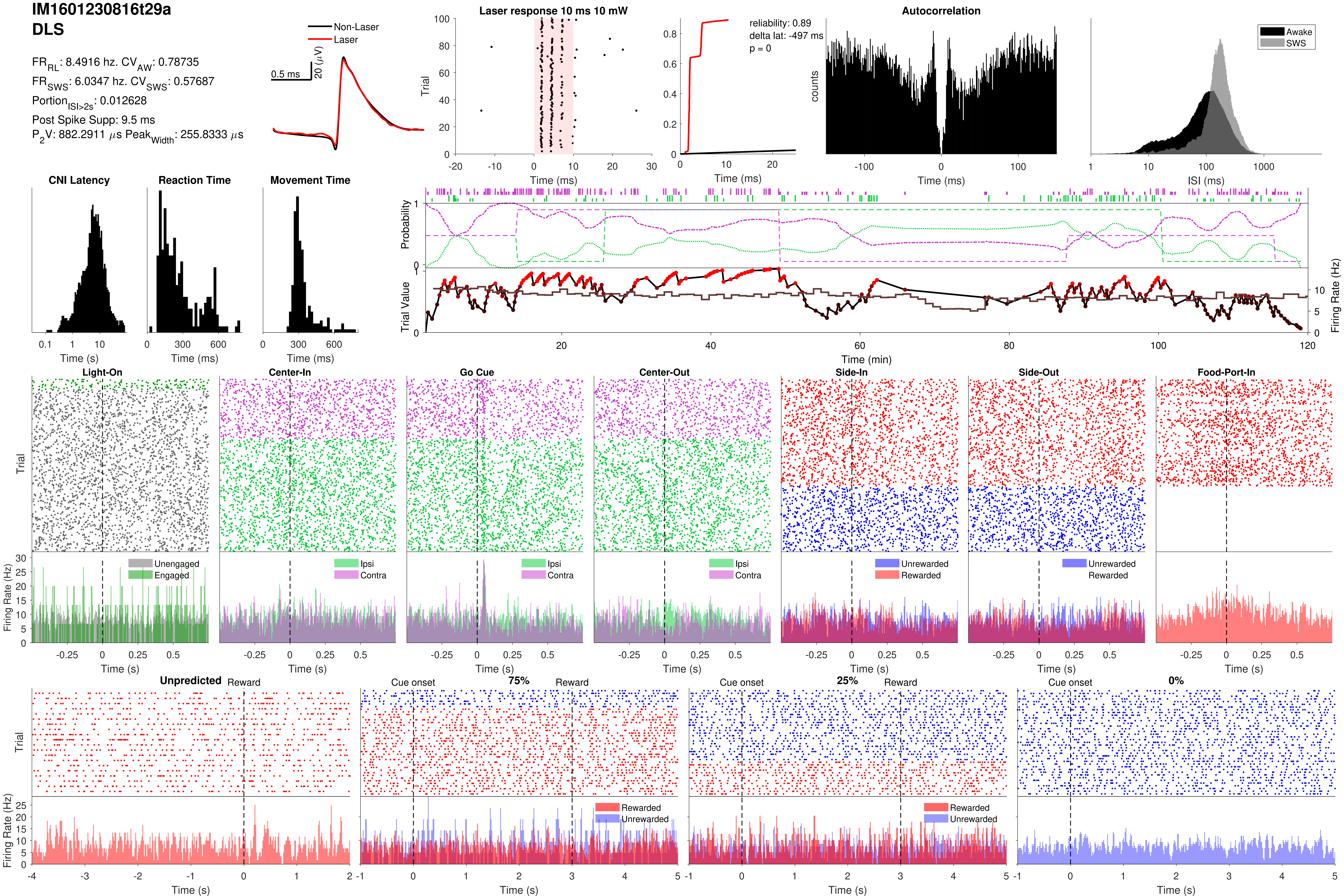

IM1601230816t29a
DLS
FR_RL: 8.4916 hz. CV_AW: 0.78735
FR_SWS: 6.0347 hz. CV_SWS: 0.57687
Portion_ISI>2s: 0.012628
Post Spike Supp: 9.5 ms
P2V: 882.2911 μs Peak_Width: 255.8333 μs
Non-Laser
Laser
0.5 ms
20 (μV)
Laser response 10 ms 10 mW
Trial
Time (ms)
reliability: 0.89
delta lat: -497 ms
p = 0
Autocorrelation
counts
Awake
SWS
ISI (ms)
CNI Latency
Reaction Time
Movement Time
Time (s)
Time (ms)
Probability
Trial Value
Firing Rate (Hz)
Time (min)
Light-On
Center-In
Go Cue
Center-Out
Side-In
Side-Out
Food-Port-In
Unengaged
Engaged
Ipsi
Contra
Unrewarded
Rewarded
Unpredicted
Reward
Cue onset
75%
25%
0%
Rewarded
Unrewarded
Time (s)

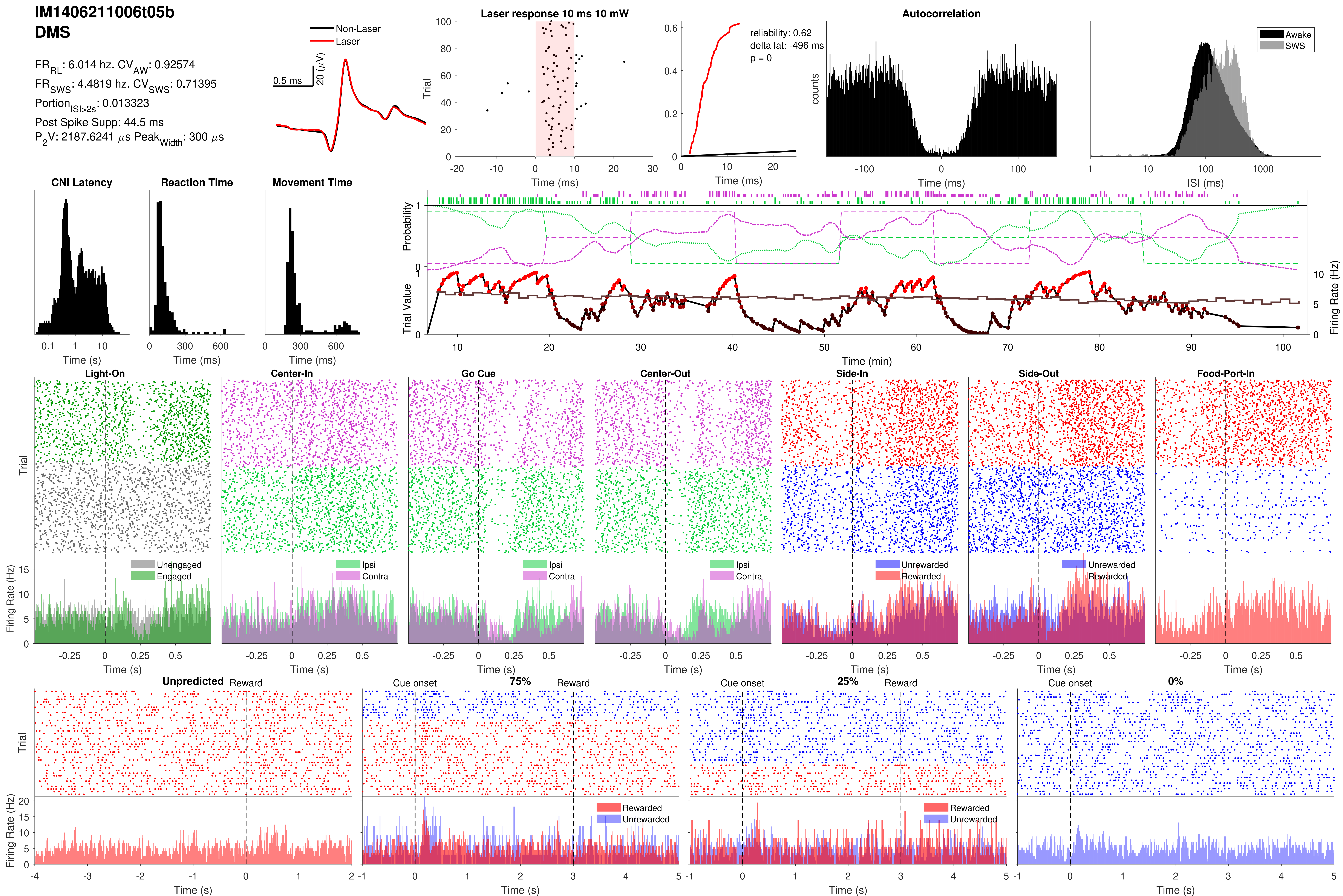

IM1406211006t05b
DMS
FR_RL: 6.014 hz. CV_AW: 0.92574
FR_SWS: 4.4819 hz. CV_SWS: 0.71395
Portion_ISI>2s: 0.013323
Post Spike Supp: 44.5 ms
P_2V: 2187.6241 µs Peak_Width: 300 µs
Non-Laser
Laser
0.5 ms
20 (µV)
Laser response 10 ms 10 mW
Trial
Time (ms)
reliability: 0.62
delta lat: -496 ms
p = 0
Time (ms)
Autocorrelation
counts
Time (ms)
Awake
SWS
ISI (ms)
CNI Latency
Time (s)
Reaction Time
Time (ms)
Movement Time
Time (ms)
Probability
Trial Value
Firing Rate (Hz)
Time (min)
Light-On
Center-In
Go Cue
Center-Out
Side-In
Side-Out
Food-Port-In
Trial
Firing Rate (Hz)
Unengaged
Engaged
Ipsi
Contra
Unrewarded
Rewarded
Time (s)
Unpredicted
Reward
Cue onset
75%
25%
0%
Rewarded
Unrewarded
Time (s)

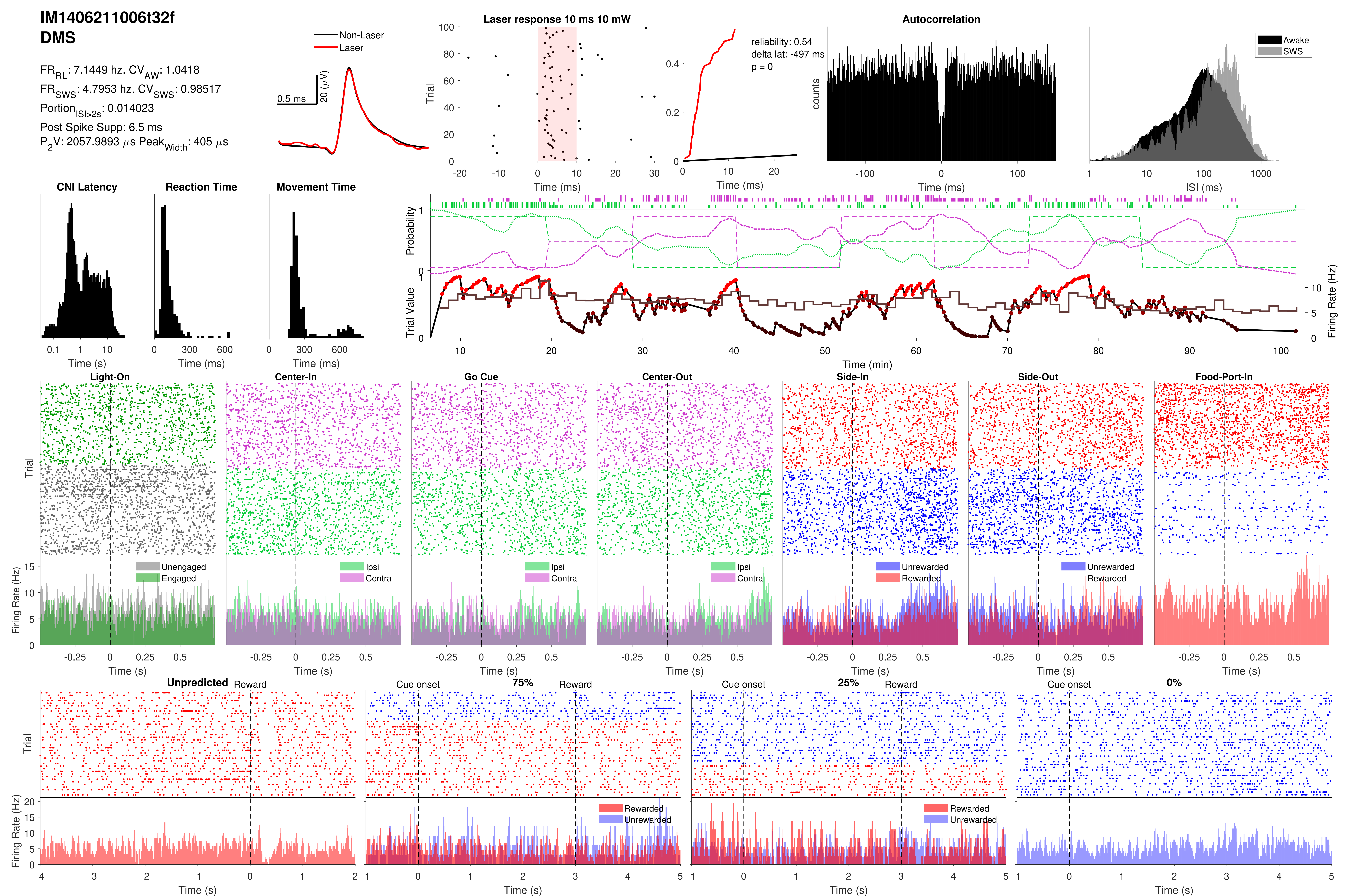

IM1406211006t32f
DMS
FR_RL: 7.1449 hz. CV_AW: 1.0418
FR_SWS: 4.7953 hz. CV_SWS: 0.98517
Portion_ISI>2s: 0.014023
Post Spike Supp: 6.5 ms
P_2V: 2057.9893 μs Peak_Width: 405 μs
Non-Laser
Laser
0.5 ms
20 (μV)
Laser response 10 ms 10 mW
Trial
Time (ms)
reliability: 0.54
delta lat: -497 ms
p = 0
Autocorrelation
counts
Awake
SWS
ISI (ms)
CNI Latency
Reaction Time
Movement Time
Time (s)
Time (ms)
Probability
Trial Value
Firing Rate (Hz)
Time (min)
Light-On
Center-In
Go Cue
Center-Out
Side-In
Side-Out
Food-Port-In
Unengaged
Engaged
Ipsi
Contra
Unrewarded
Rewarded
Unpredicted
Reward
Cue onset
75%
25%
0%

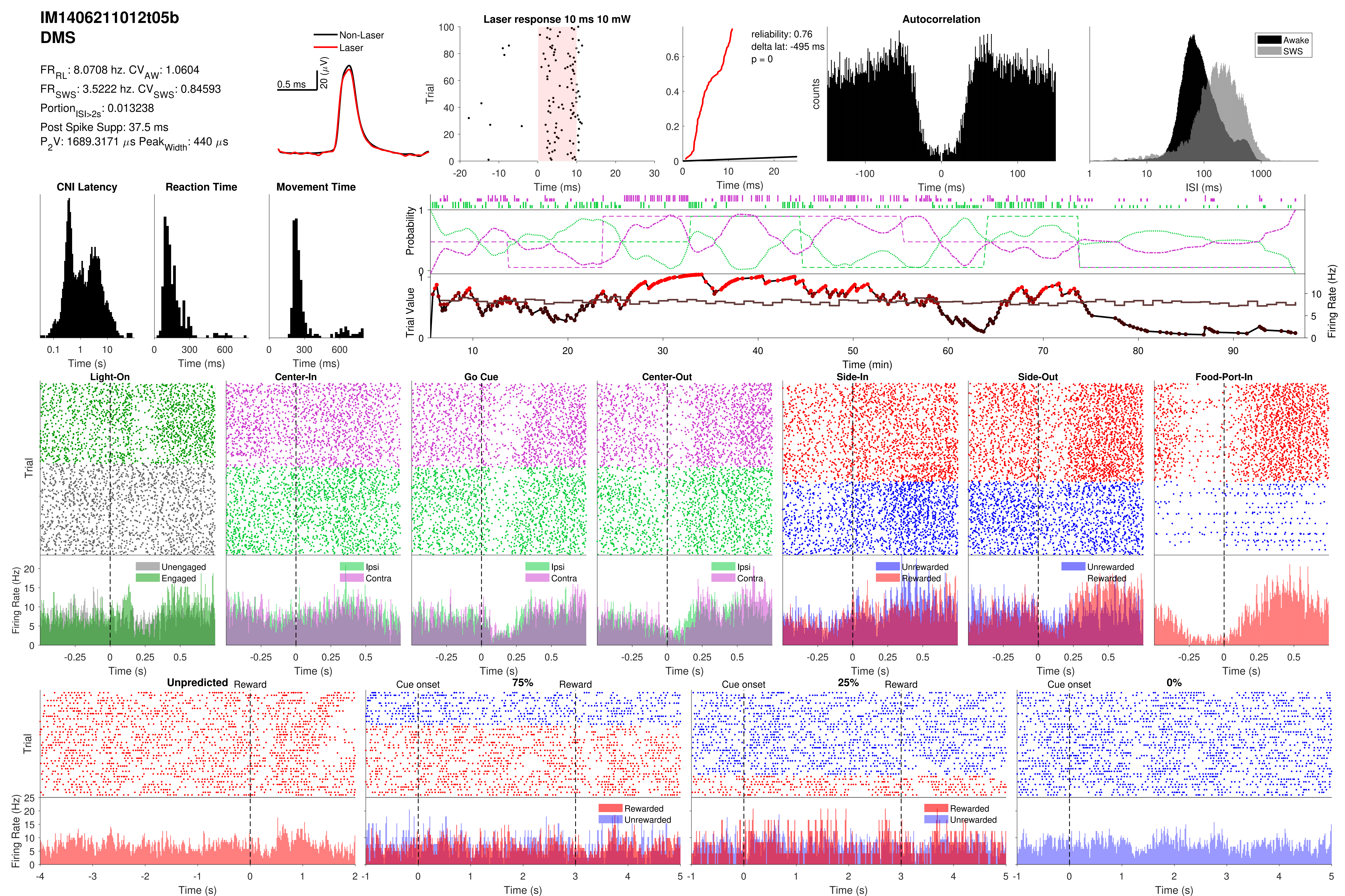
IM1406211012t05b
DMS
FR_RL: 8.0708 hz. CV_AW: 1.0604
FR_SWS: 3.5222 hz. CV_SWS: 0.84593
Portion_ISI>2s: 0.013238
Post Spike Supp: 37.5 ms
P_2V: 1689.3171 μs Peak_Width: 440 μs
Non-Laser
Laser
0.5 ms
20 (μV)
Laser response 10 ms 10 mW
Trial
Time (ms)
reliability: 0.76
delta lat: -495 ms
p = 0
Time (ms)
Autocorrelation
counts
Time (ms)
Awake
SWS
ISI (ms)
CNI Latency
Time (s)
Reaction Time
Time (ms)
Movement Time
Time (ms)
Probability
Trial Value
Firing Rate (Hz)
Time (min)
Light-On
Center-In
Go Cue
Center-Out
Side-In
Side-Out
Food-Port-In
Trial
Firing Rate (Hz)
Unengaged
Engaged
Ipsi
Contra
Unrewarded
Rewarded
Time (s)
Unpredicted
Reward
Cue onset
75%
25%
0%
Rewarded
Unrewarded
Trial
Firing Rate (Hz)
Time (s)

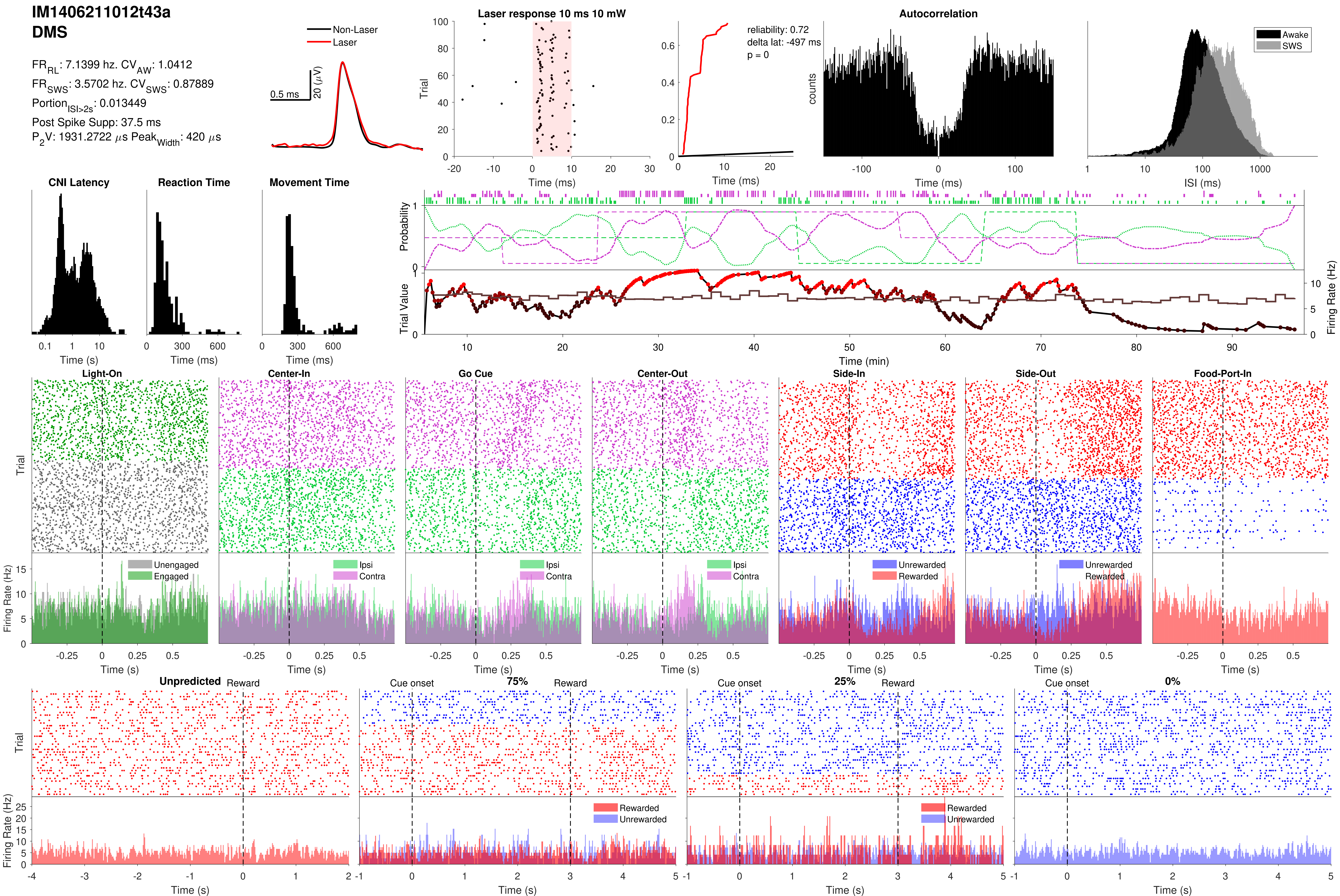
IM1406211012t43a
DMS
$FR_{RL}$: 7.1399 hz. $CV_{AW}$: 1.0412
$FR_{SWS}$: 3.5702 hz. $CV_{SWS}$: 0.87889
$Portion_{ISI>2s}$: 0.013449
Post Spike Supp: 37.5 ms
$P_2V$: 1931.2722 $\mu$s $Peak_{Width}$: 420 $\mu$s
Non-Laser
Laser
0.5 ms
20 ($\mu$V)
Laser response 10 ms 10 mW
Trial
Time (ms)
reliability: 0.72
delta lat: -497 ms
p = 0
Time (ms)
Autocorrelation
counts
Time (ms)
Awake
SWS
ISI (ms)
CNI Latency
Time (s)
Reaction Time
Time (ms)
Movement Time
Time (ms)
Probability
Trial Value
Firing Rate (Hz)
Time (min)
Light-On
Center-In
Go Cue
Center-Out
Side-In
Side-Out
Food-Port-In
Trial
Firing Rate (Hz)
Unengaged
Engaged
Ipsi
Contra
Ipsi
Contra
Ipsi
Contra
Unrewarded
Rewarded
Unrewarded
Rewarded
Time (s)
Unpredicted
Reward
Cue onset
75%
Reward
Cue onset
25%
Reward
Cue onset
0%
Trial
Firing Rate (Hz)
Rewarded
Unrewarded
Rewarded
Unrewarded
Time (s)

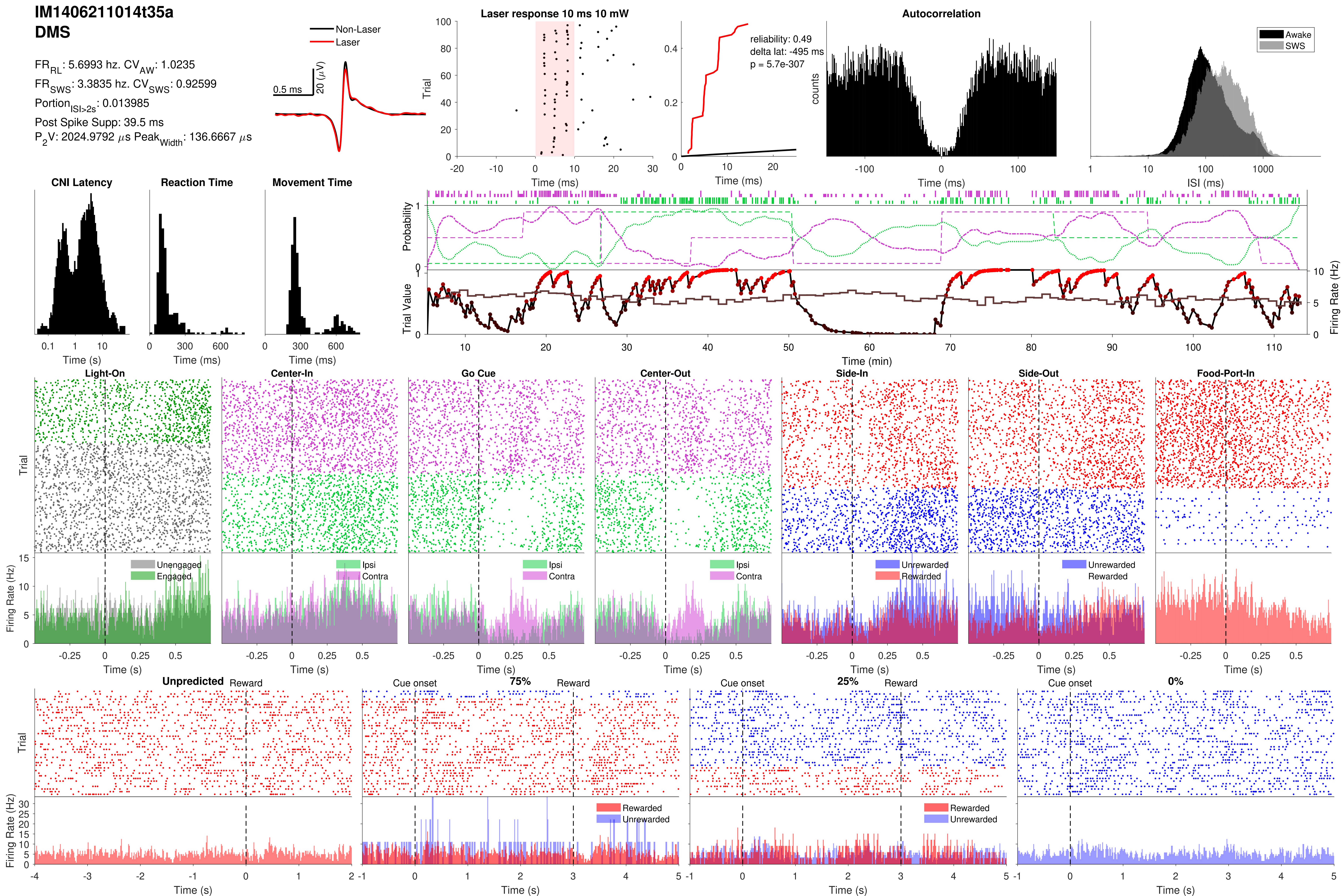
IM1406211014t35a
DMS
FR_RL: 5.6993 hz. CV_AW: 1.0235
FR_SWS: 3.3835 hz. CV_SWS: 0.92599
Portion_ISI>2s: 0.013985
Post Spike Supp: 39.5 ms
P_2V: 2024.9792 µs Peak_Width: 136.6667 µs
Non-Laser
Laser
0.5 ms
20 (µV)
Laser response 10 ms 10 mW
Trial
Time (ms)
reliability: 0.49
delta lat: -495 ms
p = 5.7e-307
Time (ms)
Autocorrelation
counts
Time (ms)
Awake
SWS
ISI (ms)
CNI Latency
Time (s)
Reaction Time
Time (ms)
Movement Time
Time (ms)
Probability
Trial Value
Firing Rate (Hz)
Time (min)
Light-On
Center-In
Go Cue
Center-Out
Side-In
Side-Out
Food-Port-In
Trial
Firing Rate (Hz)
Unengaged
Engaged
Ipsi
Contra
Unrewarded
Rewarded
Time (s)
Unpredicted
Reward
Cue onset
75%
25%
0%
Rewarded
Unrewarded
Time (s)

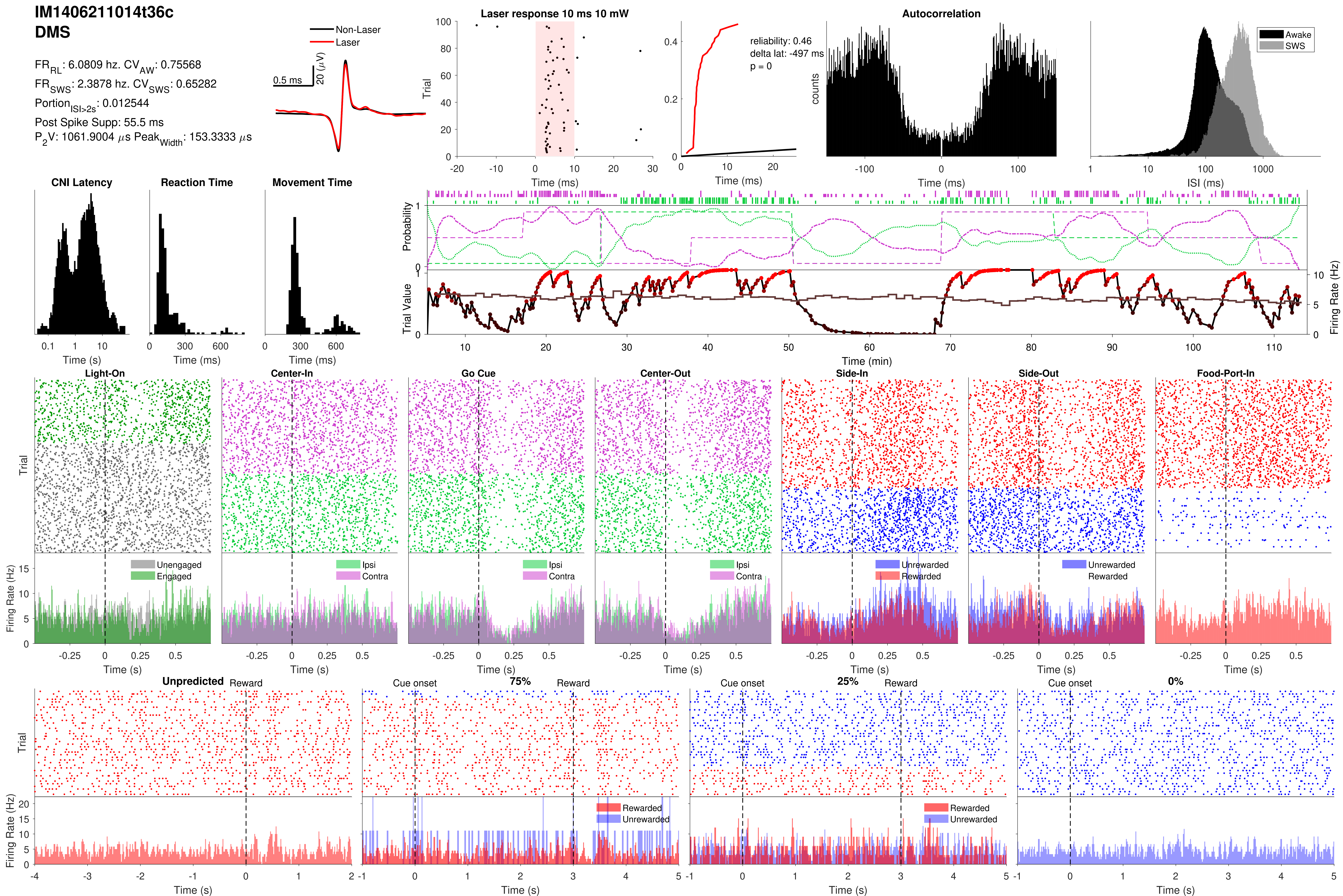
IM1406211014t36c
DMS
FR_RL: 6.0809 hz. CV_AW: 0.75568
FR_SWS: 2.3878 hz. CV_SWS: 0.65282
Portion_ISI>2s: 0.012544
Post Spike Supp: 55.5 ms
P_2V: 1061.9004 µs Peak_Width: 153.3333 µs
Non-Laser
Laser
0.5 ms
20 (µV)
Laser response 10 ms 10 mW
Trial
Time (ms)
reliability: 0.46
delta lat: -497 ms
p = 0
Time (ms)
Autocorrelation
counts
Time (ms)
Awake
SWS
ISI (ms)
CNI Latency
Time (s)
Reaction Time
Time (ms)
Movement Time
Time (ms)
Probability
Trial Value
Firing Rate (Hz)
Time (min)
Light-On
Center-In
Go Cue
Center-Out
Side-In
Side-Out
Food-Port-In
Trial
Firing Rate (Hz)
Unengaged
Engaged
Ipsi
Contra
Unrewarded
Rewarded
Time (s)
Unpredicted
Reward
Cue onset
75%
25%
0%
Rewarded
Unrewarded
Time (s)

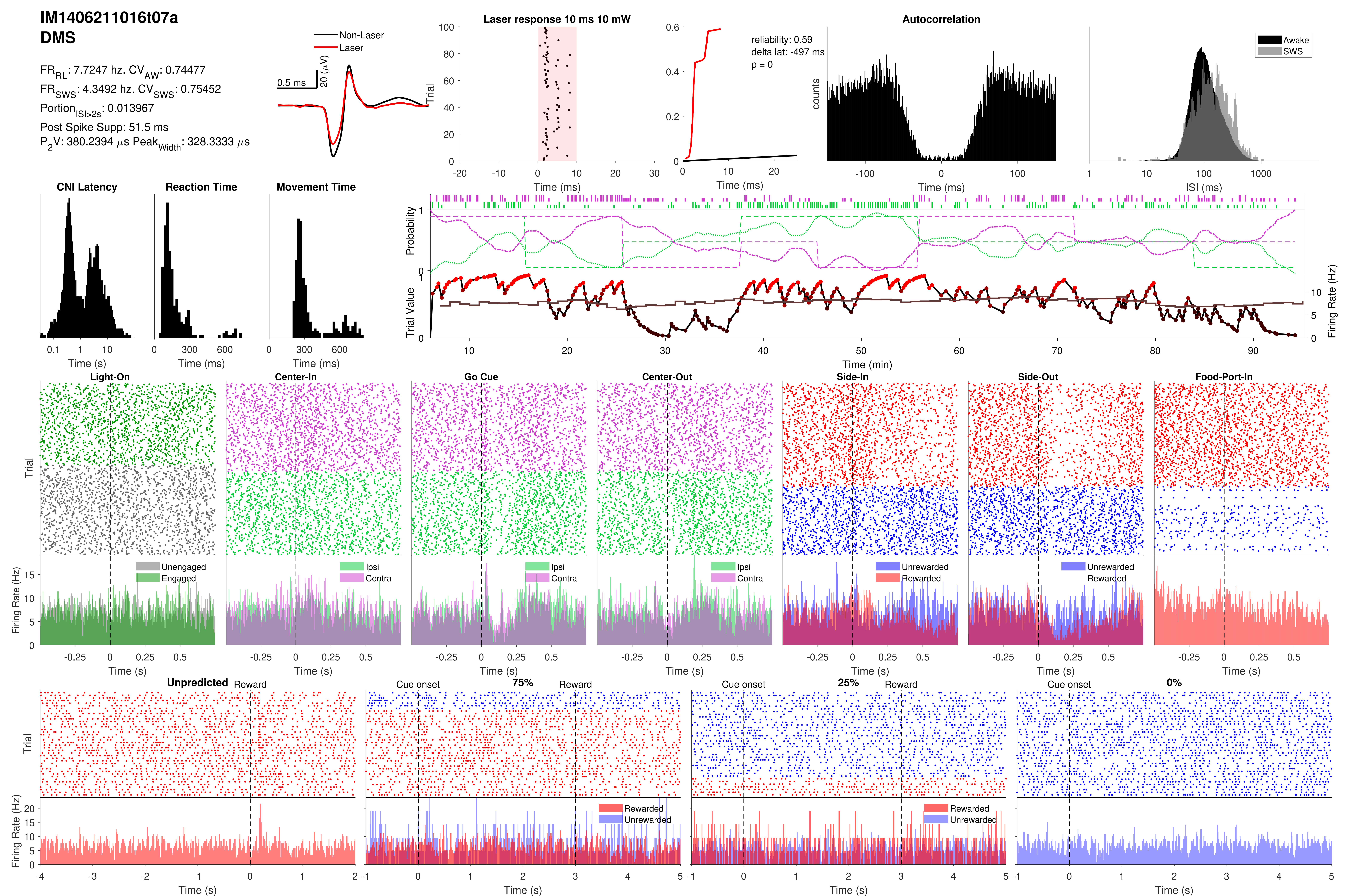
IM1406211016t07a
DMS
FR_RL: 7.7247 hz. CV_AW: 0.74477
FR_SWS: 4.3492 hz. CV_SWS: 0.75452
Portion_ISI>2s: 0.013967
Post Spike Supp: 51.5 ms
P_2V: 380.2394 µs Peak_Width: 328.3333 µs
Non-Laser
Laser
0.5 ms
20 (µV)
Laser response 10 ms 10 mW
Trial
Time (ms)
reliability: 0.59
delta lat: -497 ms
p = 0
Time (ms)
Autocorrelation
counts
Time (ms)
Awake
SWS
ISI (ms)
CNI Latency
Time (s)
Reaction Time
Time (ms)
Movement Time
Time (ms)
Probability
Trial Value
Firing Rate (Hz)
Time (min)
Light-On
Center-In
Go Cue
Center-Out
Side-In
Side-Out
Food-Port-In
Trial
Unengaged
Engaged
Ipsi
Contra
Unrewarded
Rewarded
Firing Rate (Hz)
Time (s)
Unpredicted
Reward
Cue onset
75%
25%
0%
Rewarded
Unrewarded
Trial
Firing Rate (Hz)
Time (s)

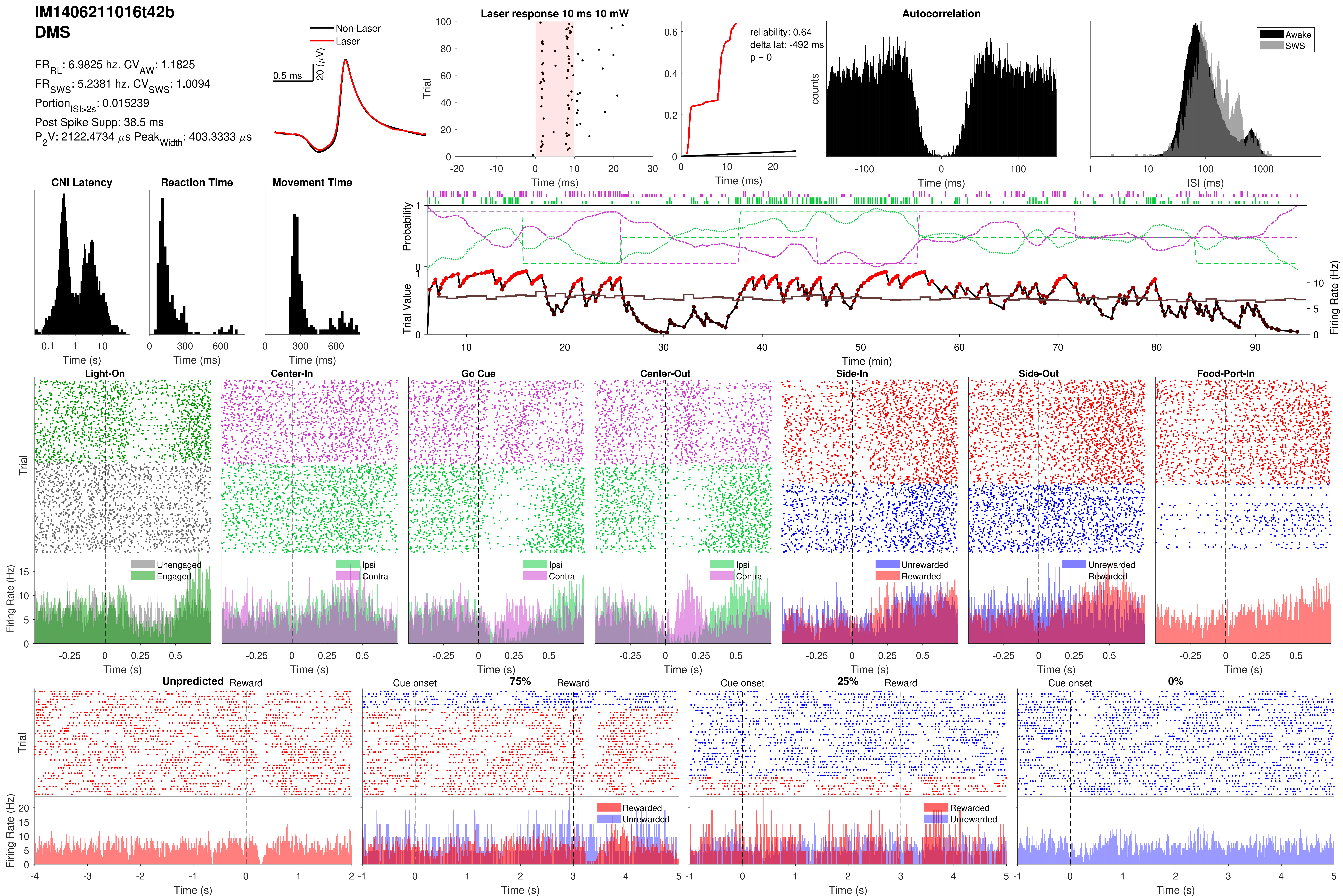

IM1406211016t42b
DMS
FR_RL: 6.9825 hz. CV_AW: 1.1825
FR_SWS: 5.2381 hz. CV_SWS: 1.0094
Portion_ISI>2s: 0.015239
Post Spike Supp: 38.5 ms
P_2V: 2122.4734 µs Peak_Width: 403.3333 µs
Non-Laser
Laser
0.5 ms
20 (µV)
Laser response 10 ms 10 mW
Trial
Time (ms)
reliability: 0.64
delta lat: -492 ms
p = 0
Time (ms)
Autocorrelation
counts
Time (ms)
Awake
SWS
ISI (ms)
CNI Latency
Time (s)
Reaction Time
Time (ms)
Movement Time
Time (ms)
Probability
Trial Value
Firing Rate (Hz)
Time (min)
Light-On
Center-In
Go Cue
Center-Out
Side-In
Side-Out
Food-Port-In
Trial
Firing Rate (Hz)
Unengaged
Engaged
Ipsi
Contra
Unrewarded
Rewarded
Time (s)
Unpredicted
Reward
Cue onset
75%
25%
0%
Rewarded
Unrewarded
Time (s)

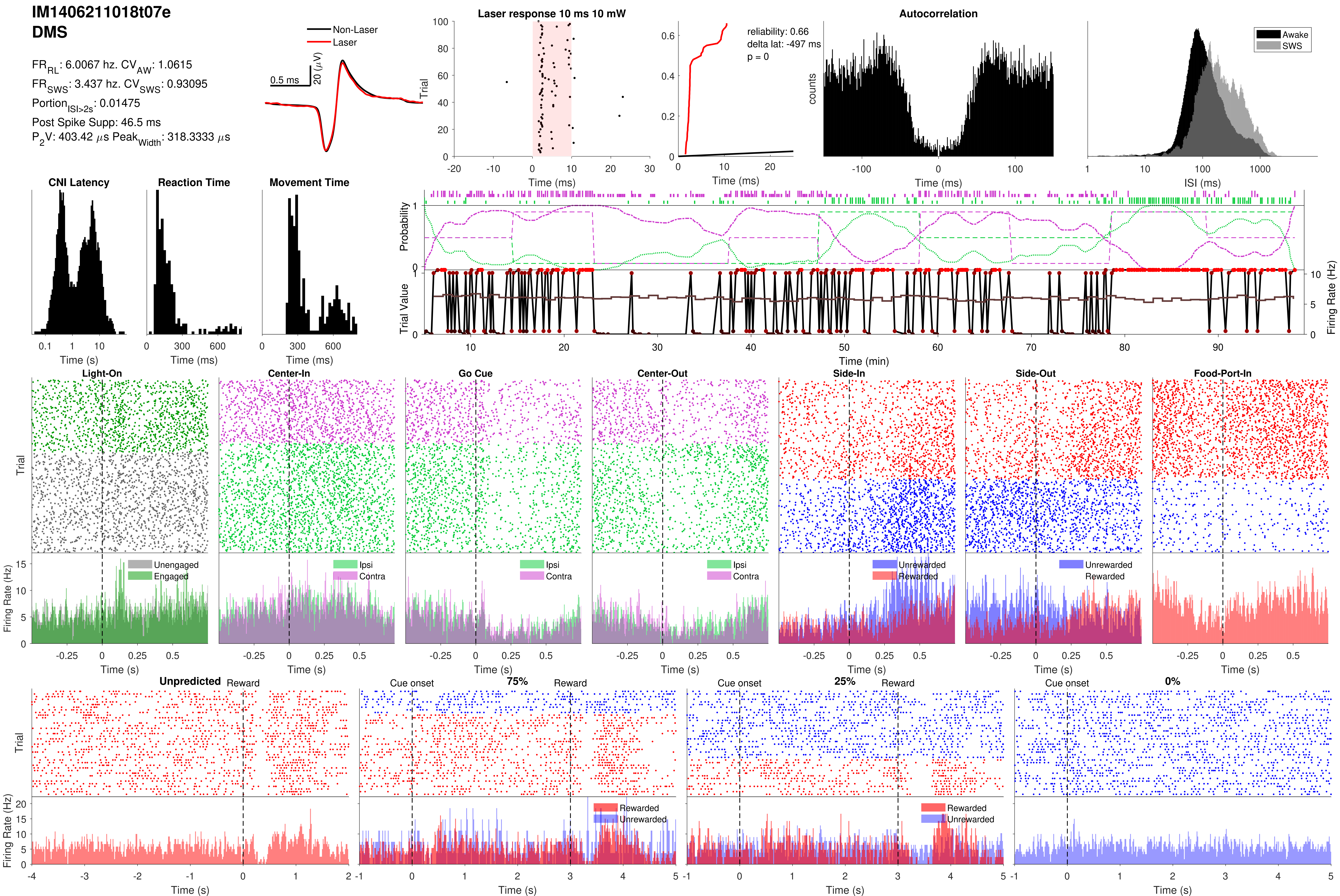
IM1406211018t07e
DMS
FR_RL: 6.0067 hz. CV_AW: 1.0615
FR_SWS: 3.437 hz. CV_SWS: 0.93095
Portion_ISI>2s: 0.01475
Post Spike Supp: 46.5 ms
P_2V: 403.42 μs Peak_Width: 318.3333 μs
Non-Laser
Laser
0.5 ms
20 (μV)
Laser response 10 ms 10 mW
Trial
Time (ms)
reliability: 0.66
delta lat: -497 ms
p = 0
Time (ms)
Autocorrelation
counts
Time (ms)
Awake
SWS
ISI (ms)
CNI Latency
Time (s)
Reaction Time
Time (ms)
Movement Time
Time (ms)
Probability
Trial Value
Firing Rate (Hz)
Time (min)
Light-On
Center-In
Go Cue
Center-Out
Side-In
Side-Out
Food-Port-In
Trial
Firing Rate (Hz)
Unengaged
Engaged
Ipsi
Contra
Unrewarded
Rewarded
Time (s)
Unpredicted
Reward
Cue onset
75%
25%
0%
Rewarded
Unrewarded
Trial
Firing Rate (Hz)
Time (s)

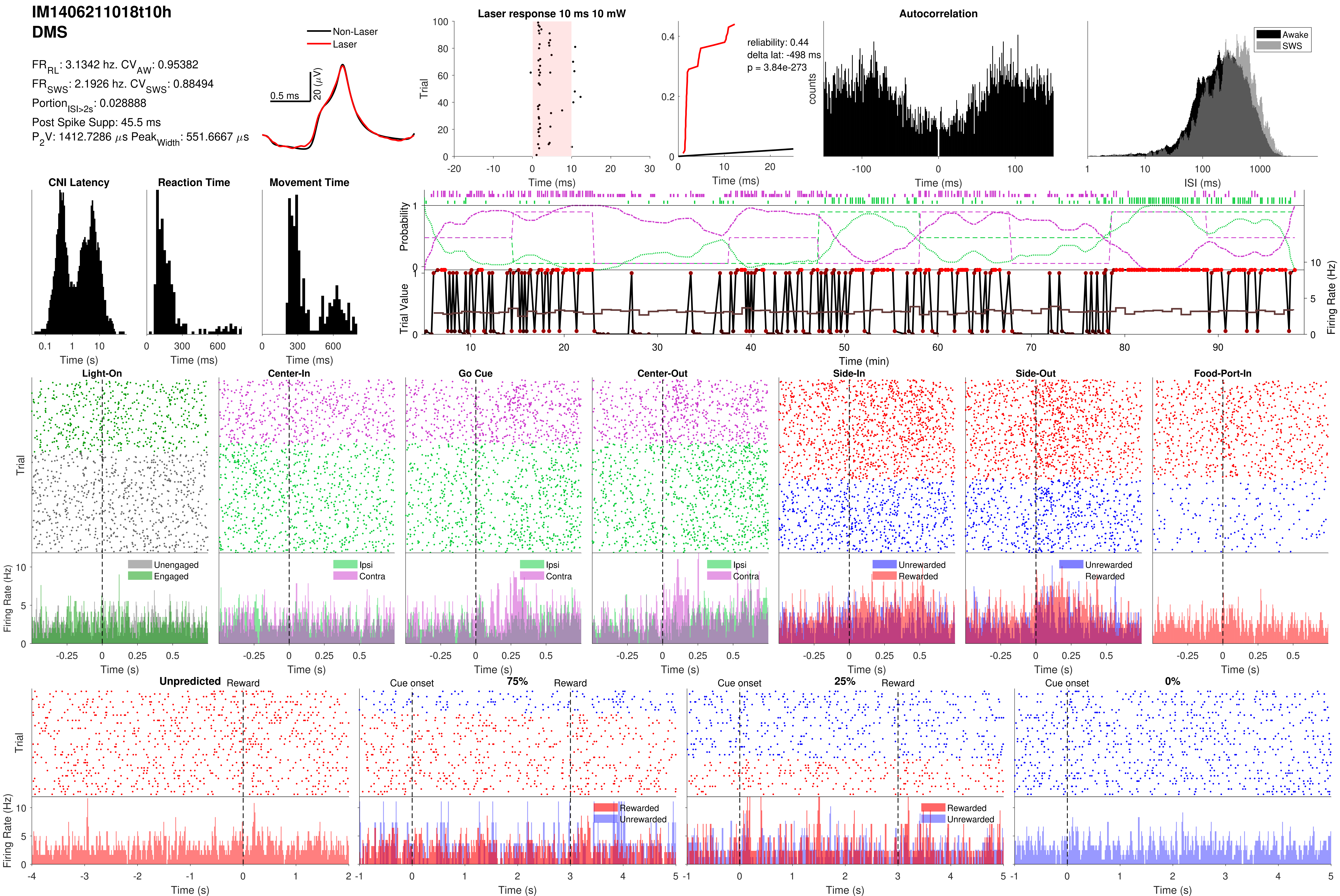

IM1406211018t10h
DMS
FR_RL: 3.1342 hz. CV_AW: 0.95382
FR_SWS: 2.1926 hz. CV_SWS: 0.88494
Portion_ISI>2s: 0.028888
Post Spike Supp: 45.5 ms
P_2V: 1412.7286 μs Peak_Width: 551.6667 μs
Non-Laser
Laser
0.5 ms
20 (μV)
Laser response 10 ms 10 mW
Trial
Time (ms)
reliability: 0.44
delta lat: -498 ms
p = 3.84e-273
Time (ms)
Autocorrelation
counts
Time (ms)
Awake
SWS
ISI (ms)
CNI Latency
Time (s)
Reaction Time
Time (ms)
Movement Time
Time (ms)
Probability
Trial Value
Firing Rate (Hz)
Time (min)
Light-On
Center-In
Go Cue
Center-Out
Side-In
Side-Out
Food-Port-In
Trial
Unengaged
Engaged
Ipsi
Contra
Unrewarded
Rewarded
Firing Rate (Hz)
Time (s)
Unpredicted
Reward
Cue onset
75%
25%
0%
Rewarded
Unrewarded
Firing Rate (Hz)
Time (s)

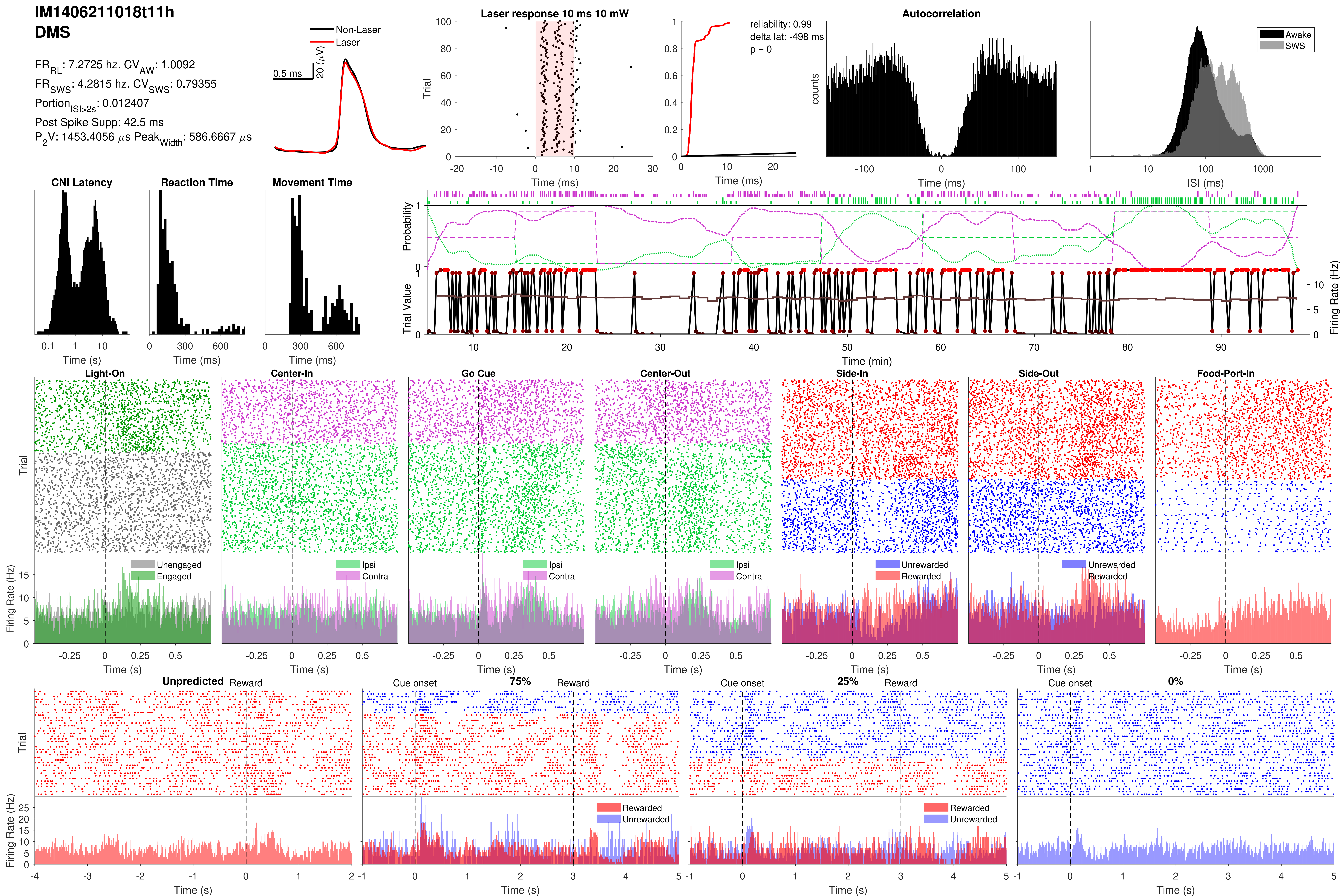

IM1406211018t11h
DMS
FR_RL: 7.2725 hz. CV_AW: 1.0092
FR_SWS: 4.2815 hz. CV_SWS: 0.79355
Portion_ISI>2s: 0.012407
Post Spike Supp: 42.5 ms
P_2V: 1453.4056 μs Peak_Width: 586.6667 μs
Non-Laser
Laser
0.5 ms
20 (μV)
Laser response 10 ms 10 mW
Trial
Time (ms)
reliability: 0.99
delta lat: -498 ms
p = 0
Time (ms)
Autocorrelation
counts
Time (ms)
Awake
SWS
ISI (ms)
CNI Latency
Time (s)
Reaction Time
Time (ms)
Movement Time
Time (ms)
Probability
Trial Value
Firing Rate (Hz)
Time (min)
Light-On
Center-In
Go Cue
Center-Out
Side-In
Side-Out
Food-Port-In
Trial
Firing Rate (Hz)
Unengaged
Engaged
Ipsi
Contra
Unrewarded
Rewarded
Time (s)
Unpredicted
Reward
Cue onset
75%
25%
0%
Rewarded
Unrewarded
Time (s)

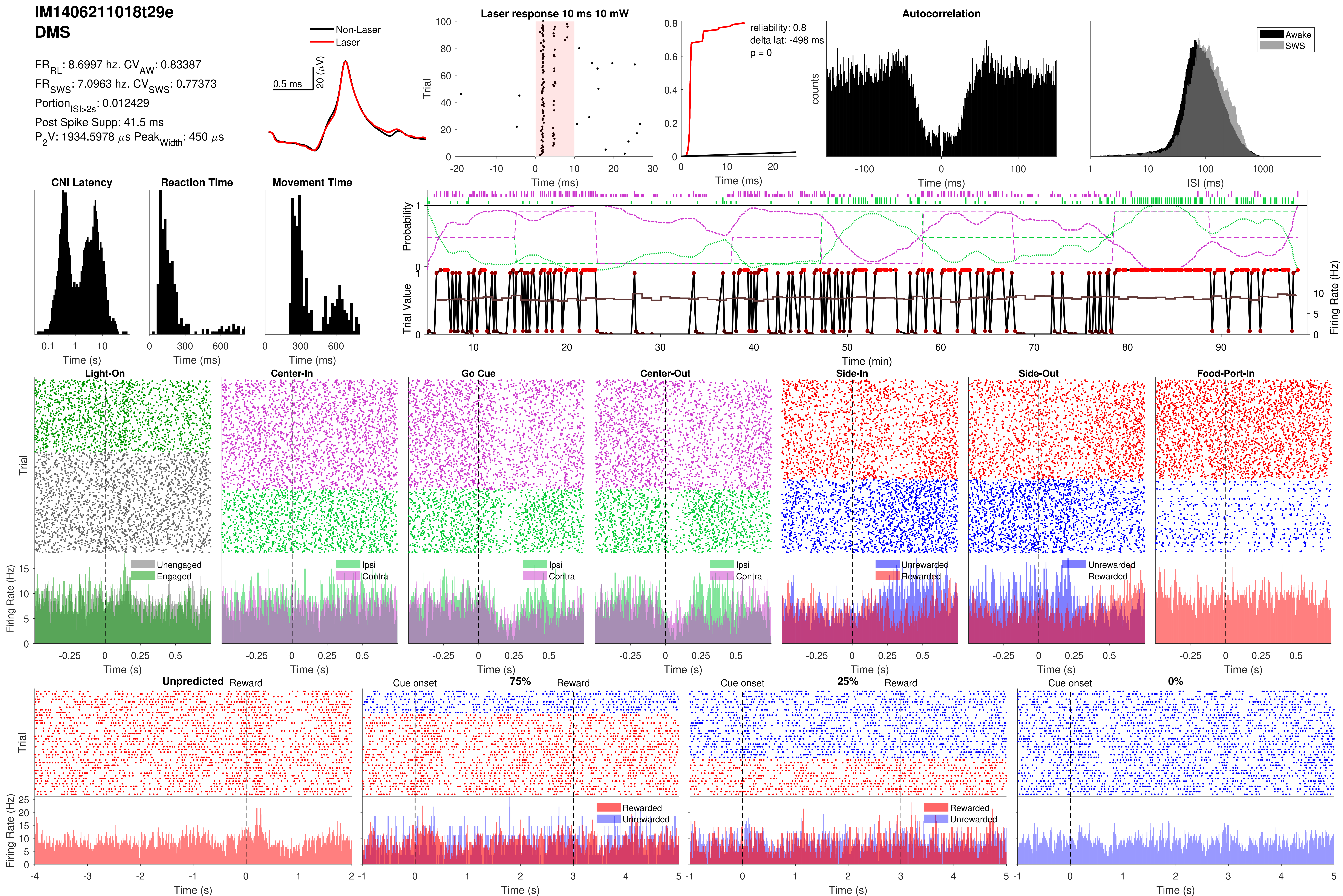
IM1406211018t29e
DMS
FR_RL: 8.6997 hz. CV_AW: 0.83387
FR_SWS: 7.0963 hz. CV_SWS: 0.77373
Portion_ISI>2s: 0.012429
Post Spike Supp: 41.5 ms
P_2V: 1934.5978 μs Peak_Width: 450 μs
Non-Laser
Laser
0.5 ms
20 (μV)
Laser response 10 ms 10 mW
Trial
Time (ms)
reliability: 0.8
delta lat: -498 ms
p = 0
Time (ms)
Autocorrelation
counts
Time (ms)
Awake
SWS
ISI (ms)
CNI Latency
Time (s)
Reaction Time
Time (ms)
Movement Time
Time (ms)
Probability
Trial Value
Firing Rate (Hz)
Time (min)
Light-On
Center-In
Go Cue
Center-Out
Side-In
Side-Out
Food-Port-In
Trial
Firing Rate (Hz)
Unengaged
Engaged
Ipsi
Contra
Unrewarded
Rewarded
Time (s)
Unpredicted
Reward
Cue onset
75%
25%
0%
Rewarded
Unrewarded
Time (s)

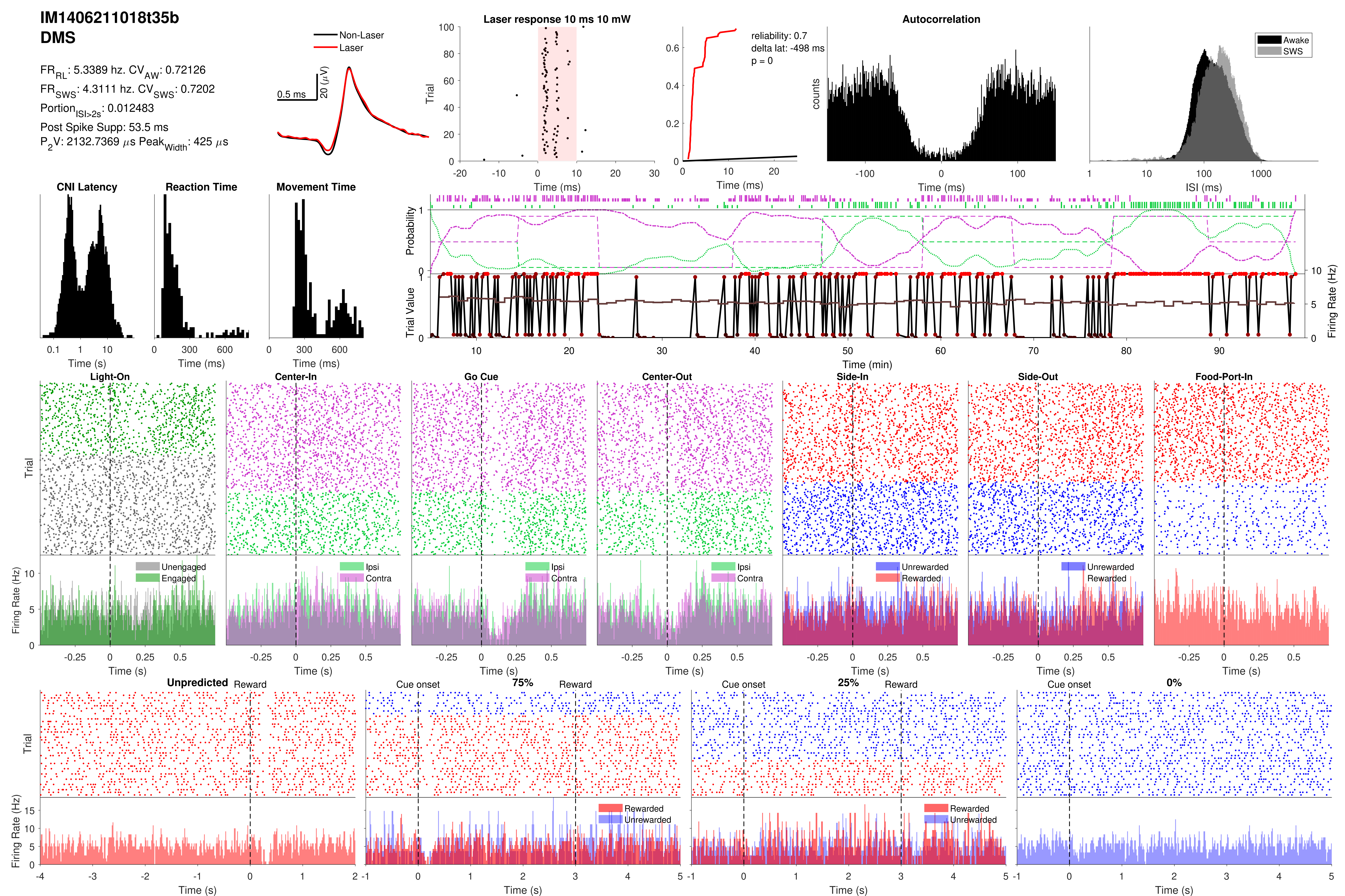

IM1406211018t35b
DMS
FR_RL: 5.3389 hz. CV_AW: 0.72126
FR_SWS: 4.3111 hz. CV_SWS: 0.7202
Portion_ISI>2s: 0.012483
Post Spike Supp: 53.5 ms
P_2V: 2132.7369 μs Peak_Width: 425 μs
Non-Laser
Laser
0.5 ms
20 (μV)
Laser response 10 ms 10 mW
Trial
Time (ms)
reliability: 0.7
delta lat: -498 ms
p = 0
Time (ms)
Autocorrelation
counts
Time (ms)
Awake
SWS
ISI (ms)
CNI Latency
Time (s)
Reaction Time
Time (ms)
Movement Time
Time (ms)
Probability
Trial Value
Firing Rate (Hz)
Time (min)
Light-On
Center-In
Go Cue
Center-Out
Side-In
Side-Out
Food-Port-In
Trial
Firing Rate (Hz)
Unengaged
Engaged
Ipsi
Contra
Unrewarded
Rewarded
Time (s)
Unpredicted
Reward
Cue onset
75%
25%
0%
Rewarded
Unrewarded
Time (s)

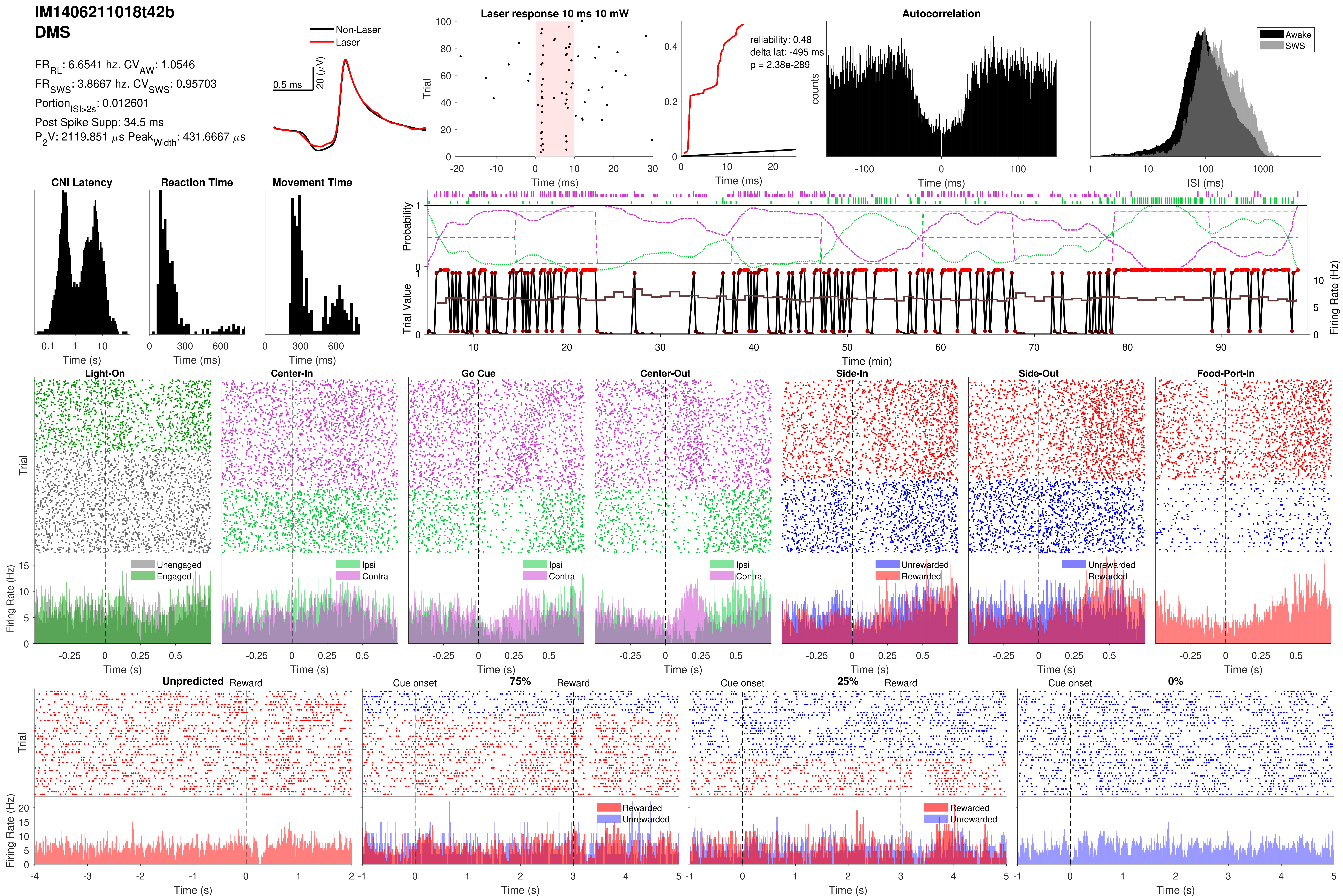
IM1406211018t42b
DMS
FR_RL: 6.6541 hz. CV_AW: 1.0546
FR_SWS: 3.8667 hz. CV_SWS: 0.95703
Portion_ISI>2s: 0.012601
Post Spike Supp: 34.5 ms
P_2V: 2119.851 μs Peak_Width: 431.6667 μs
Non-Laser
Laser
0.5 ms
20 (μV)
Laser response 10 ms 10 mW
Trial
Time (ms)
reliability: 0.48
delta lat: -495 ms
p = 2.38e-289
Time (ms)
Autocorrelation
counts
Time (ms)
Awake
SWS
ISI (ms)
CNI Latency
Time (s)
Reaction Time
Time (ms)
Movement Time
Time (ms)
Probability
Trial Value
Firing Rate (Hz)
Time (min)
Light-On
Center-In
Go Cue
Center-Out
Side-In
Side-Out
Food-Port-In
Trial
Unengaged
Engaged
Ipsi
Contra
Unrewarded
Rewarded
Firing Rate (Hz)
Time (s)
Unpredicted
Reward
Cue onset
75%
25%
0%
Trial
Rewarded
Unrewarded
Firing Rate (Hz)
Time (s)

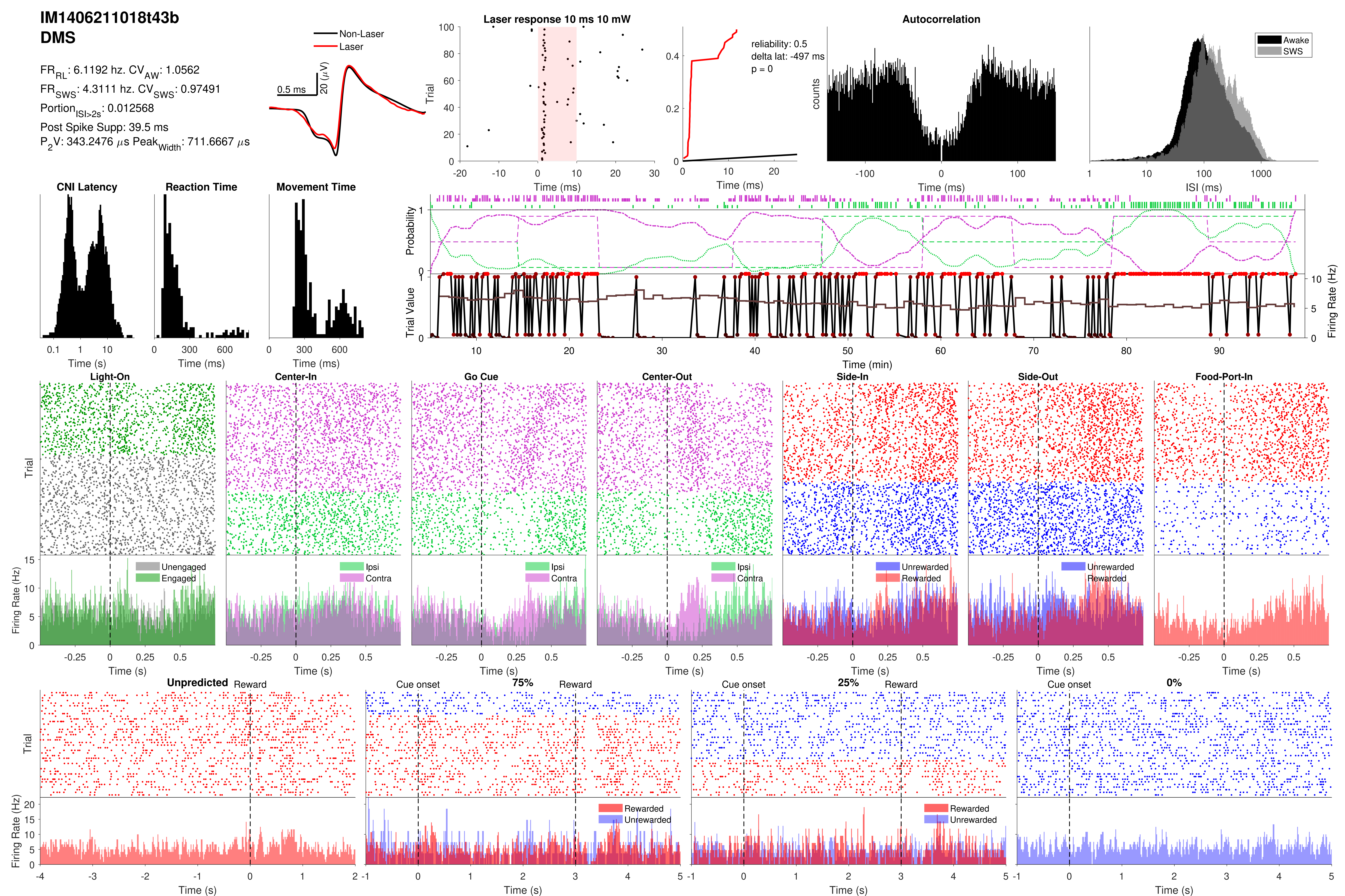
IM1406211018t43b
DMS
FR_RL: 6.1192 hz. CV_AW: 1.0562
FR_SWS: 4.3111 hz. CV_SWS: 0.97491
Portion_ISI>2s: 0.012568
Post Spike Supp: 39.5 ms
P_2V: 343.2476 μs Peak_Width: 711.6667 μs
Non-Laser
Laser
0.5 ms
20 (μV)
Laser response 10 ms 10 mW
Trial
Time (ms)
reliability: 0.5
delta lat: -497 ms
p = 0
Time (ms)
Autocorrelation
counts
Time (ms)
Awake
SWS
ISI (ms)
CNI Latency
Time (s)
Reaction Time
Time (ms)
Movement Time
Time (ms)
Probability
Trial Value
Firing Rate (Hz)
Time (min)
Light-On
Center-In
Go Cue
Center-Out
Side-In
Side-Out
Food-Port-In
Trial
Firing Rate (Hz)
Unengaged
Engaged
Ipsi
Contra
Unrewarded
Rewarded
Time (s)
Unpredicted
Reward
Cue onset
75%
25%
0%
Rewarded
Unrewarded
Time (s)

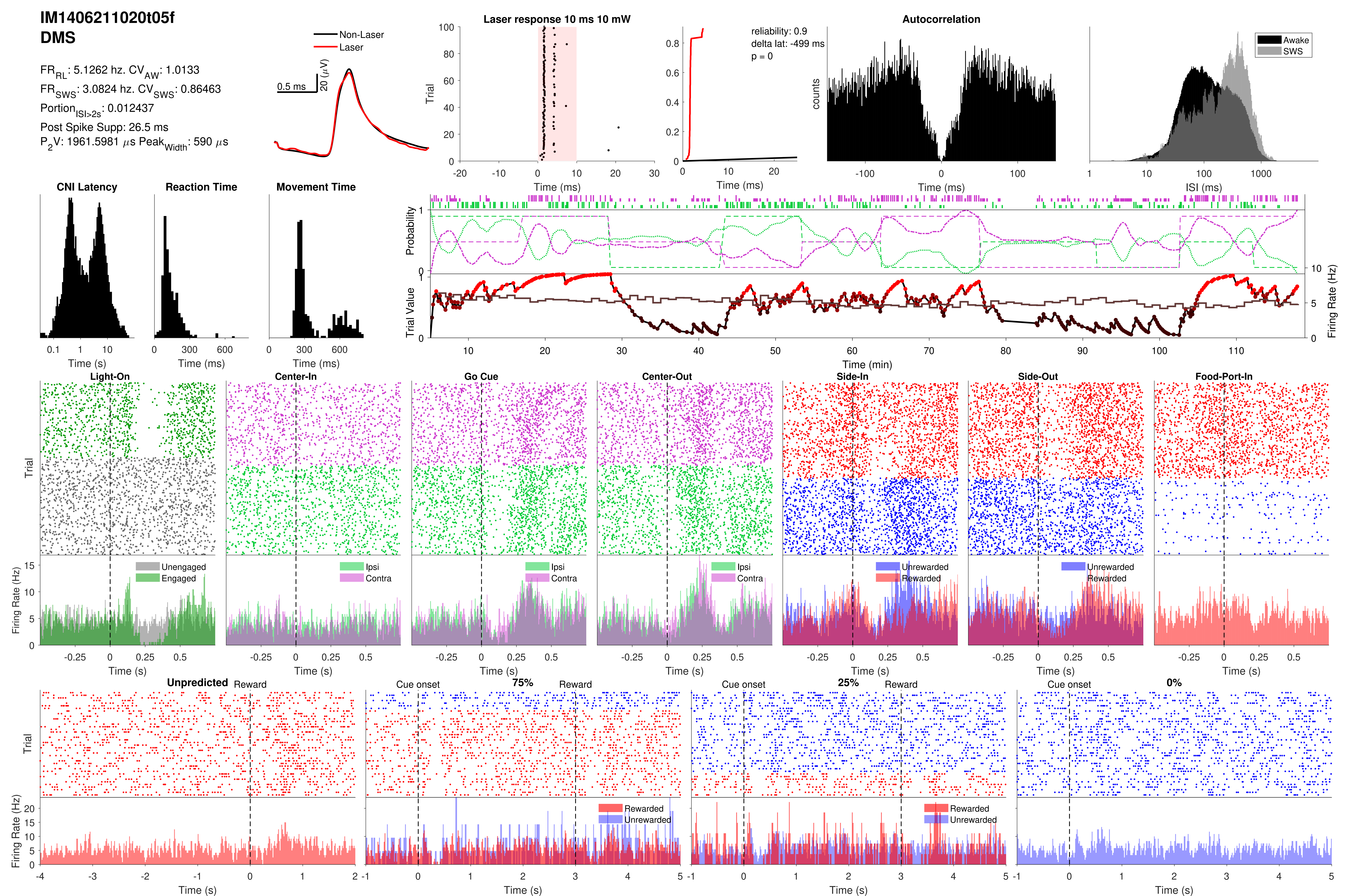

IM1406211020t05f
DMS
FR_RL: 5.1262 hz. CV_AW: 1.0133
FR_SWS: 3.0824 hz. CV_SWS: 0.86463
Portion_ISI>2s: 0.012437
Post Spike Supp: 26.5 ms
P_2V: 1961.5981 μs Peak_Width: 590 μs
Non-Laser
Laser
0.5 ms
20 (μV)
Laser response 10 ms 10 mW
Trial
Time (ms)
reliability: 0.9
delta lat: -499 ms
p = 0
Time (ms)
Autocorrelation
counts
Time (ms)
Awake
SWS
ISI (ms)
CNI Latency
Time (s)
Reaction Time
Time (ms)
Movement Time
Time (ms)
Probability
Trial Value
Firing Rate (Hz)
Time (min)
Light-On
Center-In
Go Cue
Center-Out
Side-In
Side-Out
Food-Port-In
Trial
Unengaged
Engaged
Ipsi
Contra
Unrewarded
Rewarded
Firing Rate (Hz)
Time (s)
Unpredicted
Reward
Cue onset
75%
25%
0%
Rewarded
Unrewarded

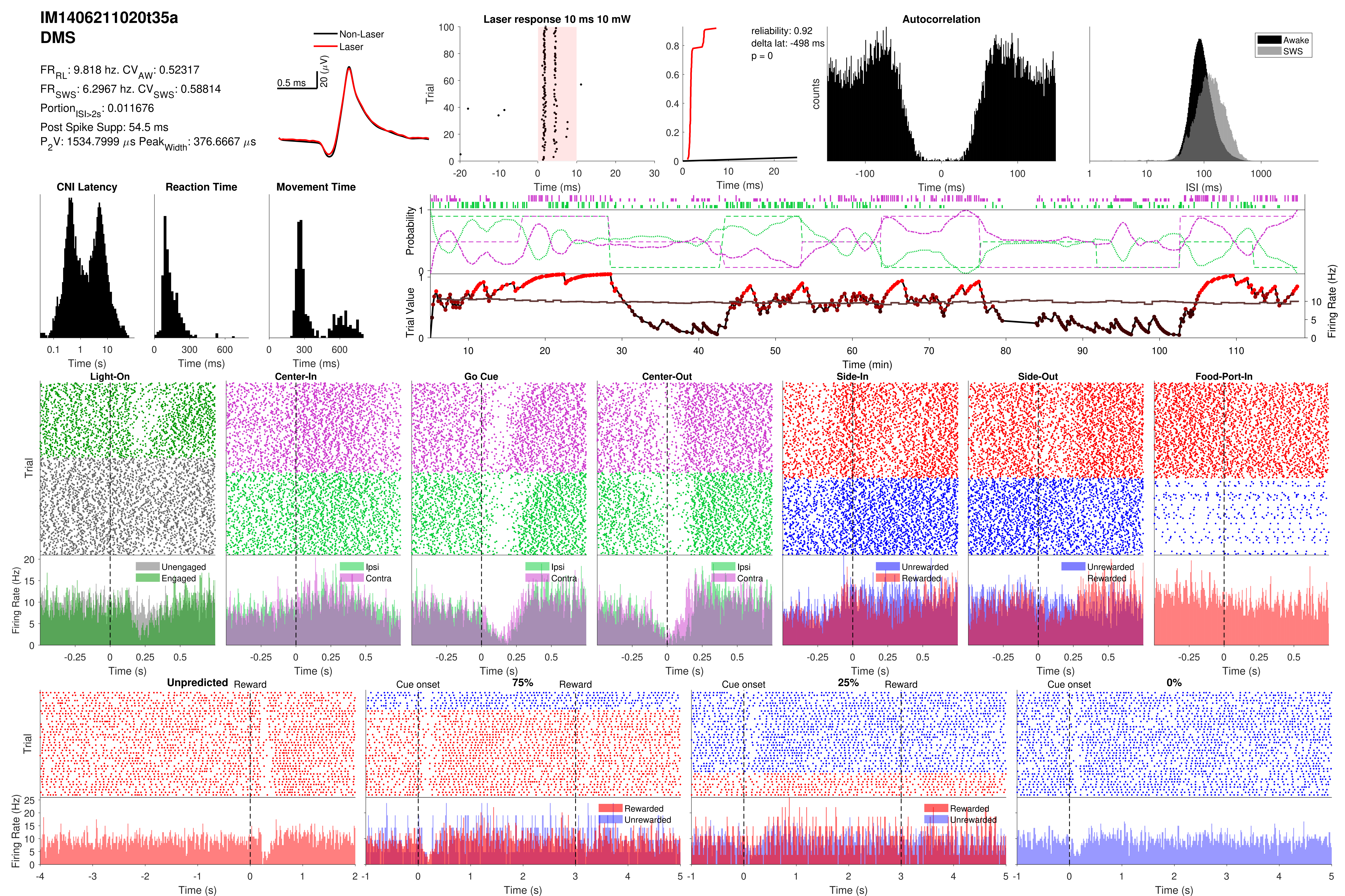
IM1406211020t35a
DMS
FR_RL: 9.818 hz. CV_AW: 0.52317
FR_SWS: 6.2967 hz. CV_SWS: 0.58814
Portion_ISI>2s: 0.011676
Post Spike Supp: 54.5 ms
P_2V: 1534.7999 μs Peak_Width: 376.6667 μs
Non-Laser
Laser
0.5 ms
20 (μV)
Laser response 10 ms 10 mW
Trial
Time (ms)
reliability: 0.92
delta lat: -498 ms
p = 0
Autocorrelation
counts
Awake
SWS
ISI (ms)
CNI Latency
Reaction Time
Movement Time
Time (s)
Probability
Trial Value
Firing Rate (Hz)
Time (min)
Light-On
Center-In
Go Cue
Center-Out
Side-In
Side-Out
Food-Port-In
Unengaged
Engaged
Ipsi
Contra
Unrewarded
Rewarded
Unpredicted
Reward
Cue onset
75%
25%
0%

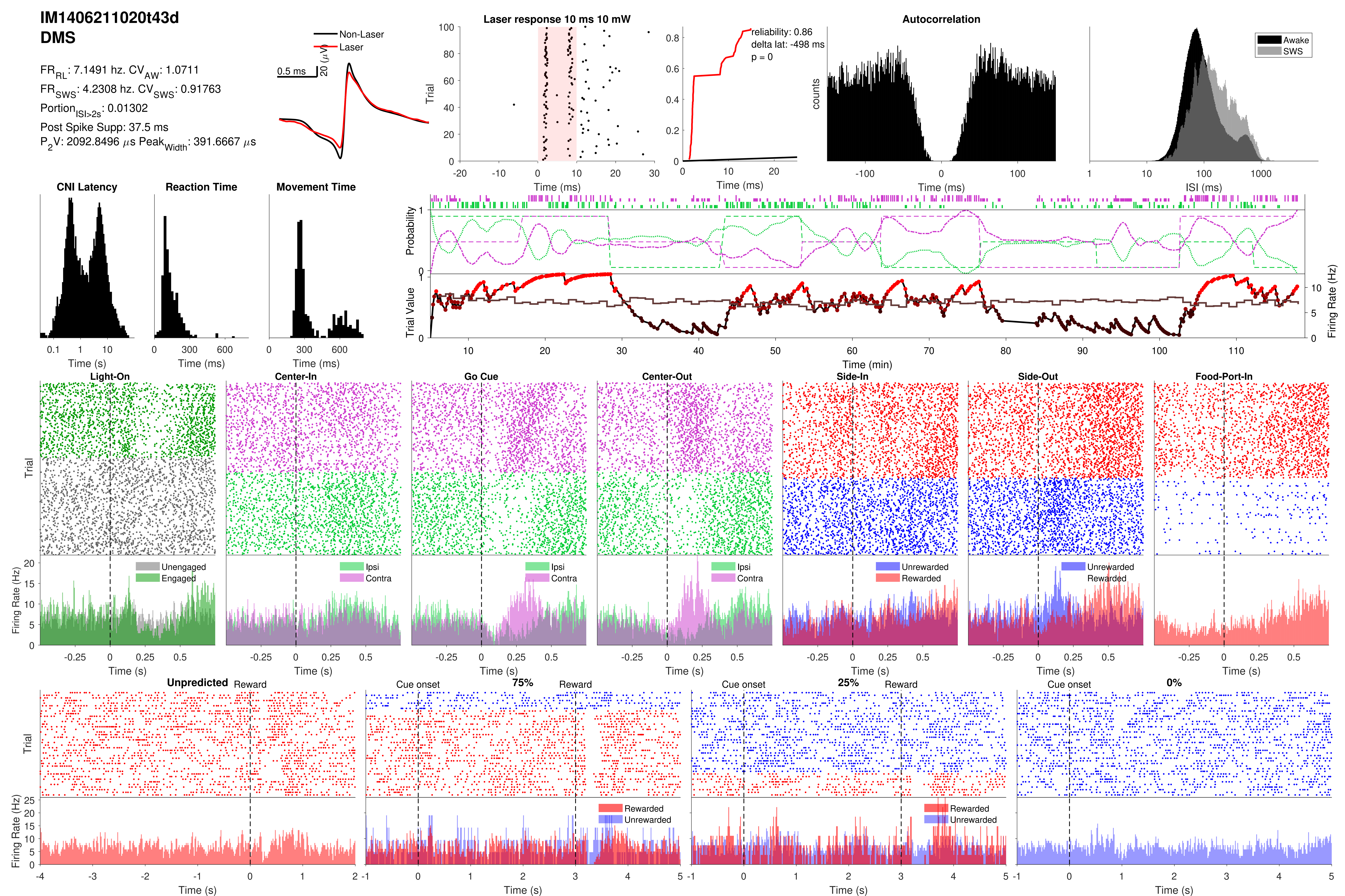

IM1406211020t43d
DMS
FR_RL: 7.1491 hz. CV_AW: 1.0711
FR_SWS: 4.2308 hz. CV_SWS: 0.91763
Portion_ISI>2s: 0.01302
Post Spike Supp: 37.5 ms
P_2V: 2092.8496 µs Peak_Width: 391.6667 µs
Non-Laser
Laser
0.5 ms
20 (µV)
Laser response 10 ms 10 mW
Trial
Time (ms)
reliability: 0.86
delta lat: -498 ms
p = 0
Autocorrelation
counts
Awake
SWS
ISI (ms)
CNI Latency
Reaction Time
Movement Time
Time (s)
Probability
Trial Value
Firing Rate (Hz)
Time (min)
Light-On
Center-In
Go Cue
Center-Out
Side-In
Side-Out
Food-Port-In
Unengaged
Engaged
Ipsi
Contra
Unrewarded
Rewarded
Unpredicted
Reward
Cue onset
75%
25%
0%

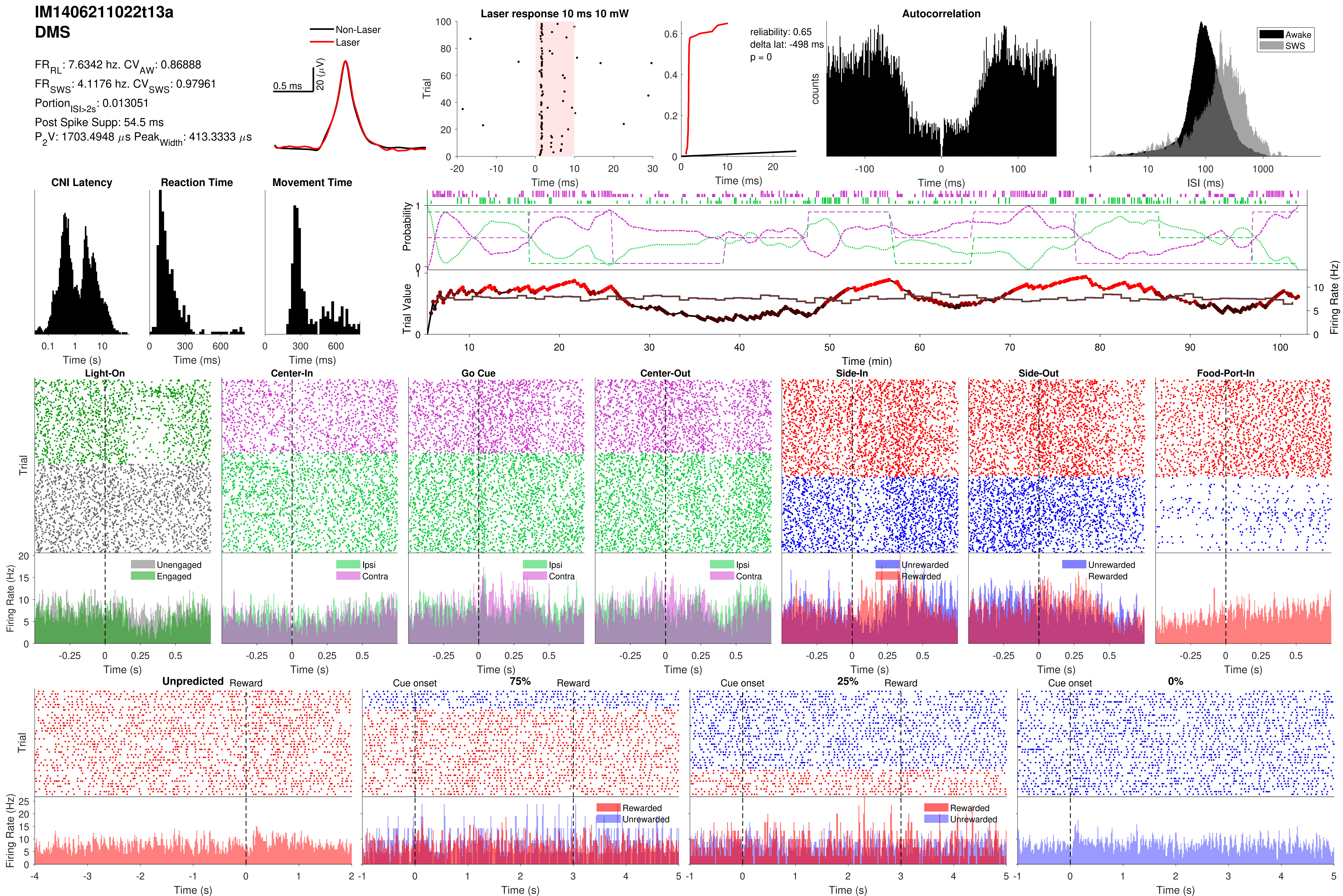

IM1406211022t13a
DMS
FR_RL: 7.6342 hz. CV_AW: 0.86888
FR_SWS: 4.1176 hz. CV_SWS: 0.97961
Portion_ISI>2s: 0.013051
Post Spike Supp: 54.5 ms
P_2V: 1703.4948 µs Peak_Width: 413.3333 µs
Non-Laser
Laser
0.5 ms
20 (µV)
Laser response 10 ms 10 mW
Trial
Time (ms)
reliability: 0.65
delta lat: -498 ms
p = 0
Time (ms)
Autocorrelation
counts
Time (ms)
Awake
SWS
ISI (ms)
CNI Latency
Time (s)
Reaction Time
Time (ms)
Movement Time
Time (ms)
Probability
Trial Value
Firing Rate (Hz)
Time (min)
Light-On
Center-In
Go Cue
Center-Out
Side-In
Side-Out
Food-Port-In
Trial
Firing Rate (Hz)
Unengaged
Engaged
Ipsi
Contra
Unrewarded
Rewarded
Time (s)
Unpredicted
Reward
Cue onset
75%
25%
0%
Trial
Firing Rate (Hz)
Rewarded
Unrewarded
Time (s)

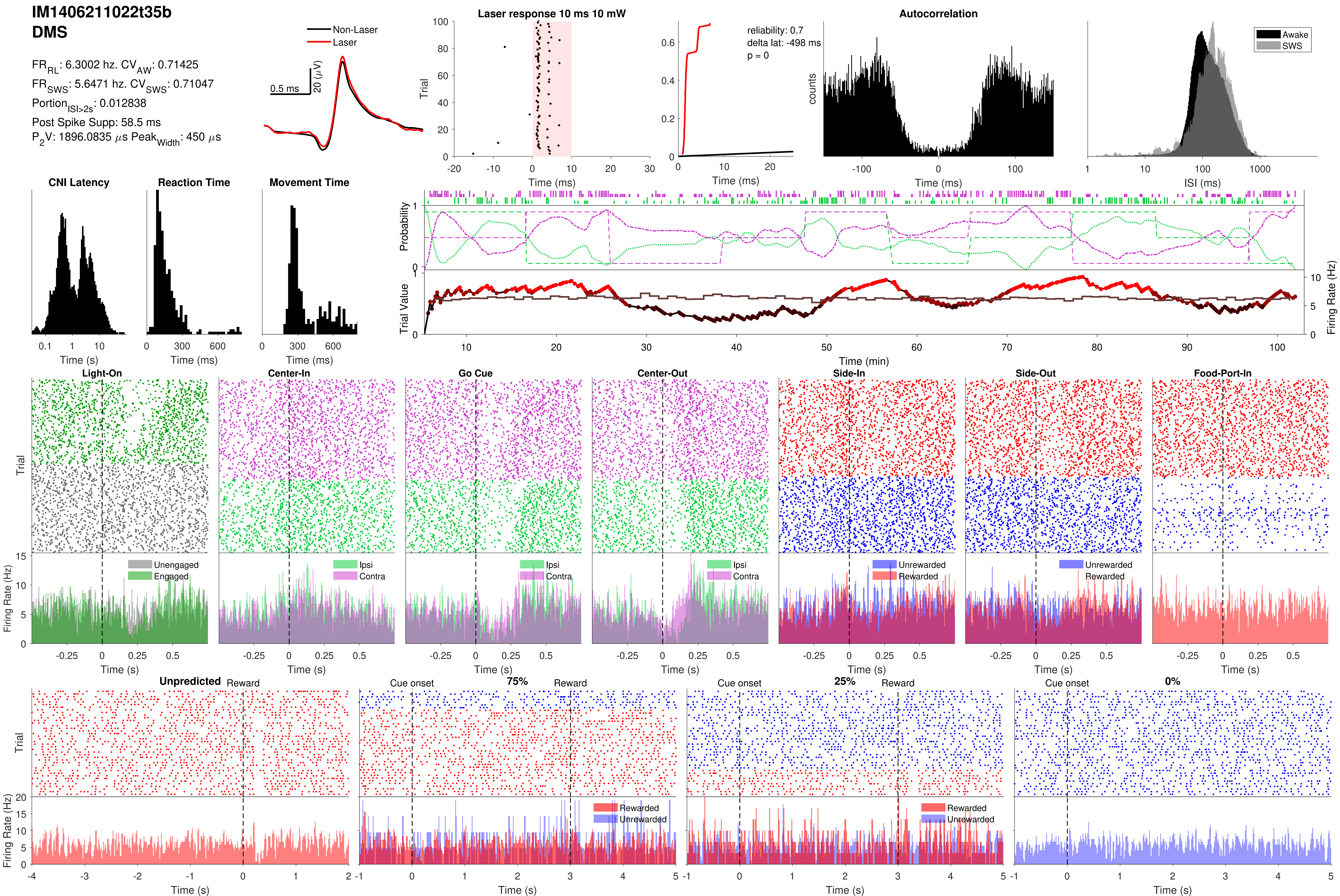

IM1406211022t35b
DMS
FR_RL: 6.3002 hz. CV_AW: 0.71425
FR_SWS: 5.6471 hz. CV_SWS: 0.71047
Portion_ISI>2s: 0.012838
Post Spike Supp: 58.5 ms
P_2V: 1896.0835 µs Peak_Width: 450 µs
Non-Laser
Laser
0.5 ms
20 (µV)
Laser response 10 ms 10 mW
Trial
Time (ms)
reliability: 0.7
delta lat: -498 ms
p = 0
Time (ms)
Autocorrelation
counts
Time (ms)
Awake
SWS
ISI (ms)
CNI Latency
Time (s)
Reaction Time
Time (ms)
Movement Time
Time (ms)
Probability
Trial Value
Firing Rate (Hz)
Time (min)
Light-On
Center-In
Go Cue
Center-Out
Side-In
Side-Out
Food-Port-In
Trial
Firing Rate (Hz)
Unengaged
Engaged
Ipsi
Contra
Unrewarded
Rewarded
Time (s)
Unpredicted
Reward
Cue onset
75%
25%
0%
Rewarded
Unrewarded
Time (s)

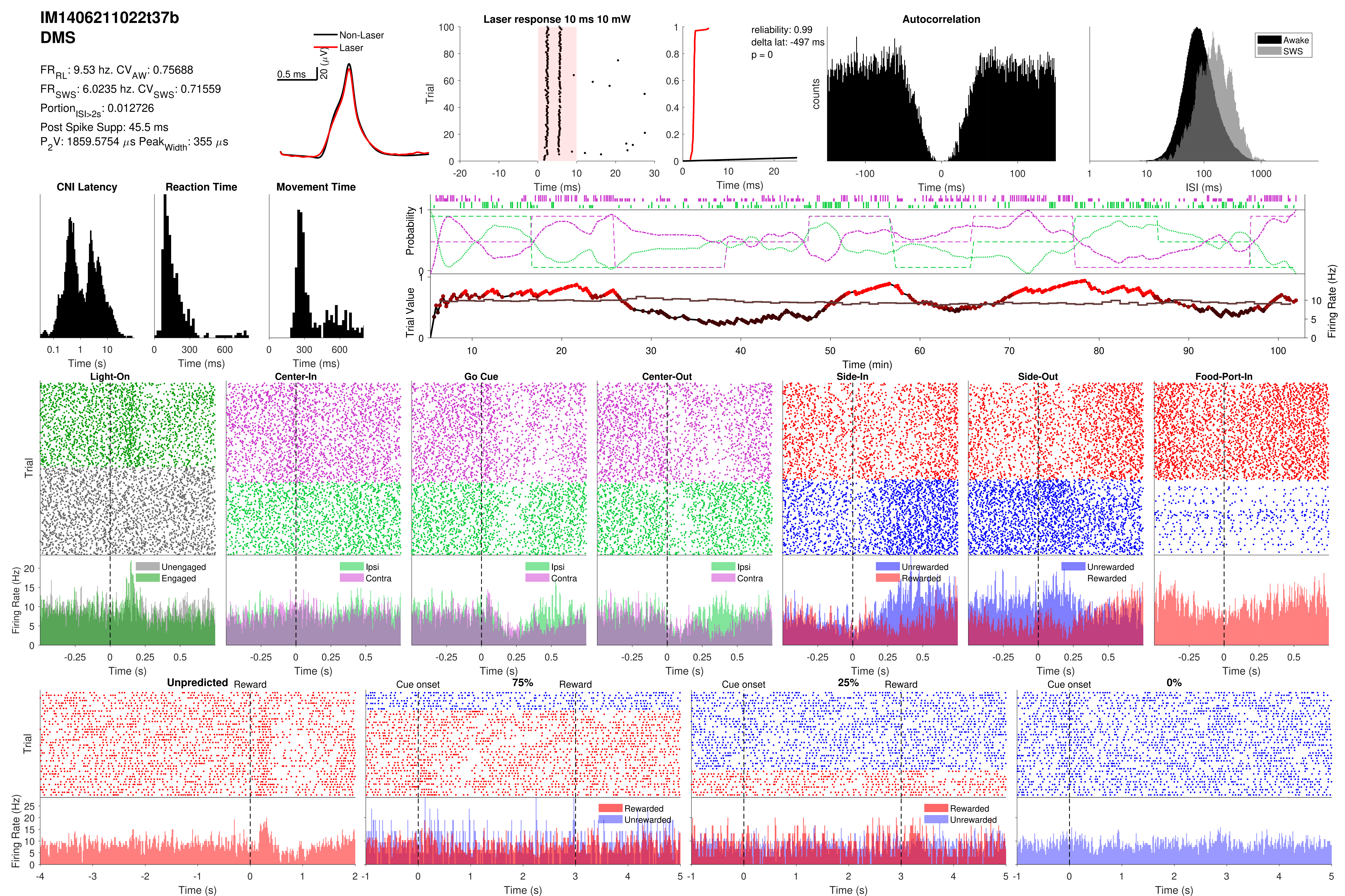
IM1406211022t37b
DMS
FR_RL: 9.53 hz. CV_AW: 0.75688
FR_SWS: 6.0235 hz. CV_SWS: 0.71559
Portion_ISI>2s: 0.012726
Post Spike Supp: 45.5 ms
P_2V: 1859.5754 μs Peak_Width: 355 μs
Non-Laser
Laser
0.5 ms
20 (μV)
Laser response 10 ms 10 mW
Trial
Time (ms)
reliability: 0.99
delta lat: -497 ms
p = 0
Time (ms)
Autocorrelation
counts
Time (ms)
Awake
SWS
ISI (ms)
CNI Latency
Time (s)
Reaction Time
Time (ms)
Movement Time
Time (ms)
Probability
Trial Value
Firing Rate (Hz)
Time (min)
Light-On
Center-In
Go Cue
Center-Out
Side-In
Side-Out
Food-Port-In
Trial
Firing Rate (Hz)
Unengaged
Engaged
Ipsi
Contra
Unrewarded
Rewarded
Time (s)
Unpredicted
Reward
Cue onset
75%
25%
0%
Rewarded
Unrewarded
Time (s)

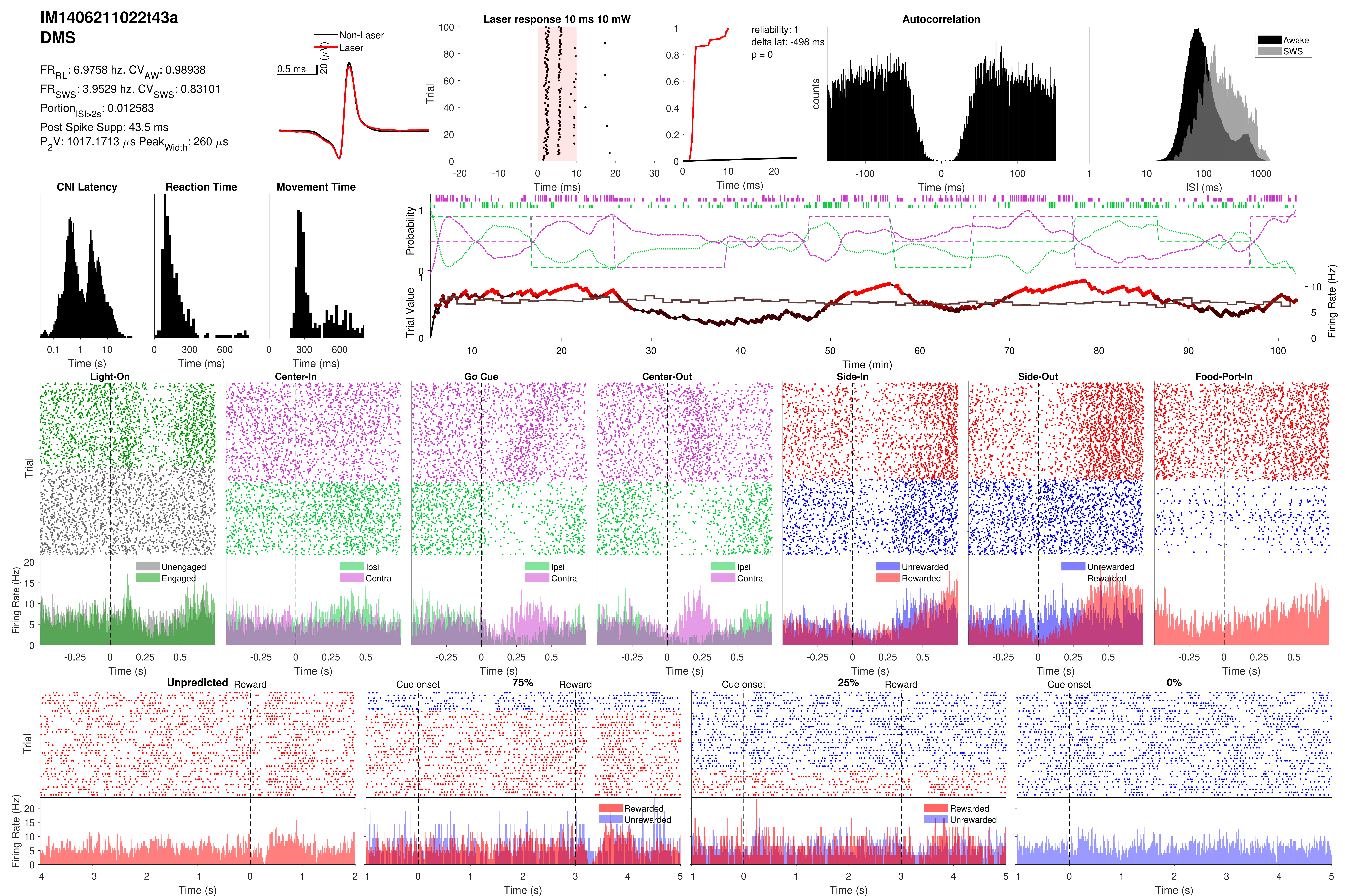
IM1406211022t43a
DMS
FR_RL: 6.9758 hz. CV_AW: 0.98938
FR_SWS: 3.9529 hz. CV_SWS: 0.83101
Portion_ISI>2s: 0.012583
Post Spike Supp: 43.5 ms
P_2V: 1017.1713 µs Peak_Width: 260 µs
Non-Laser
Laser
0.5 ms
20 (µV)
Laser response 10 ms 10 mW
Trial
Time (ms)
reliability: 1
delta lat: -498 ms
p = 0
Time (ms)
Autocorrelation
counts
Time (ms)
Awake
SWS
ISI (ms)
CNI Latency
Time (s)
Reaction Time
Time (ms)
Movement Time
Time (ms)
Probability
Trial Value
Firing Rate (Hz)
Time (min)
Light-On
Center-In
Go Cue
Center-Out
Side-In
Side-Out
Food-Port-In
Trial
Firing Rate (Hz)
Unengaged
Engaged
Ipsi
Contra
Unrewarded
Rewarded
Time (s)
Unpredicted
Reward
Cue onset
75%
25%
0%
Rewarded
Unrewarded
Time (s)

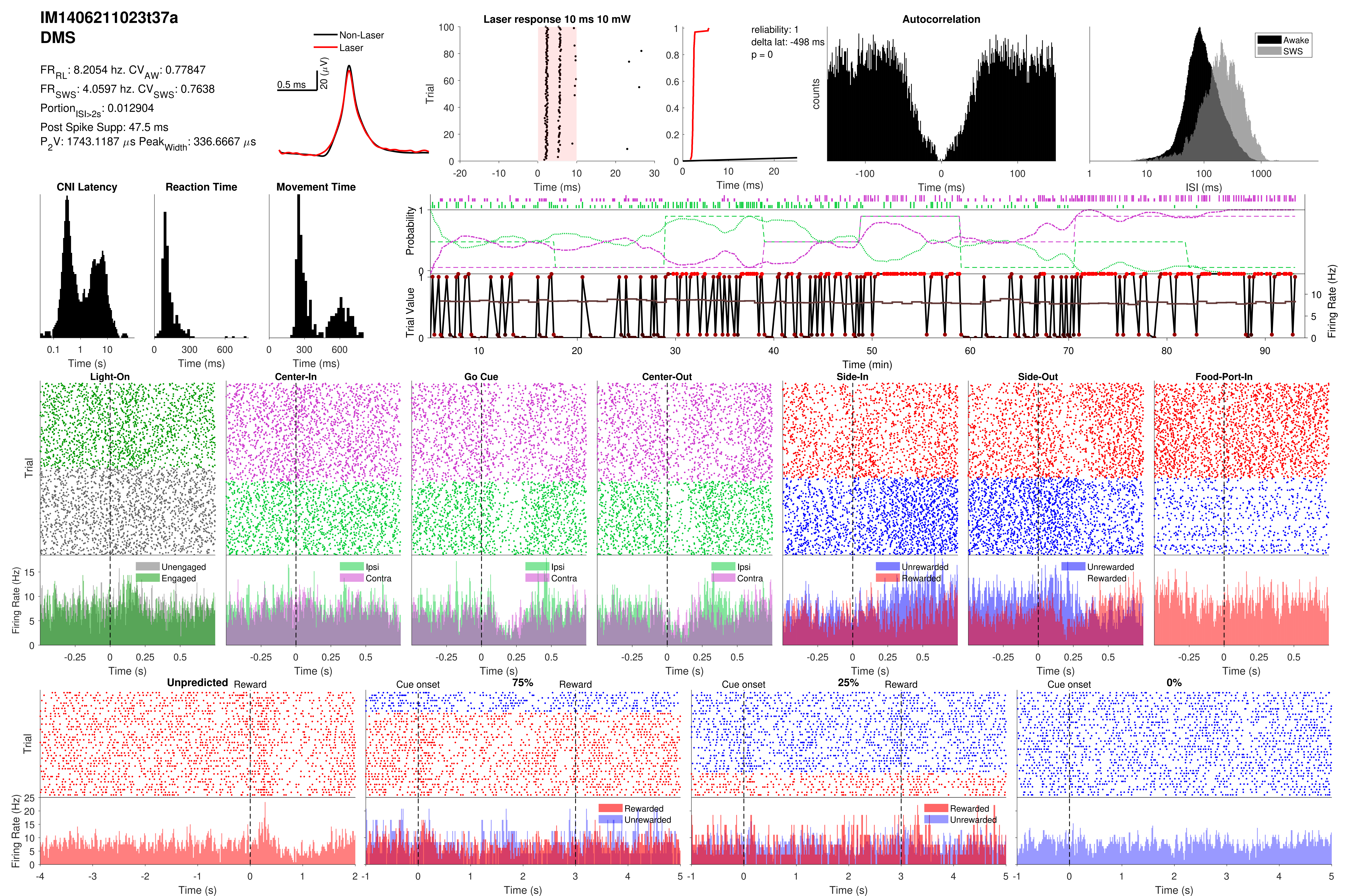

IM1406211023t37a
DMS
FR_RL: 8.2054 hz. CV_AW: 0.77847
FR_SWS: 4.0597 hz. CV_SWS: 0.7638
Portion_ISI>2s: 0.012904
Post Spike Supp: 47.5 ms
P2V: 1743.1187 µs Peak_Width: 336.6667 µs
Non-Laser
Laser
0.5 ms
20 (µV)
Laser response 10 ms 10 mW
Trial
Time (ms)
reliability: 1
delta lat: -498 ms
p = 0
Time (ms)
Autocorrelation
counts
Time (ms)
Awake
SWS
ISI (ms)
CNI Latency
Time (s)
Reaction Time
Time (ms)
Movement Time
Time (ms)
Probability
Trial Value
Firing Rate (Hz)
Time (min)
Light-On
Center-In
Go Cue
Center-Out
Side-In
Side-Out
Food-Port-In
Trial
Firing Rate (Hz)
Unengaged
Engaged
Ipsi
Contra
Ipsi
Contra
Ipsi
Contra
Unrewarded
Rewarded
Unrewarded
Rewarded
Time (s)
Unpredicted
Reward
Cue onset
75%
Reward
Cue onset
25%
Reward
Cue onset
0%
Trial
Firing Rate (Hz)
Rewarded
Unrewarded
Rewarded
Unrewarded
Time (s)

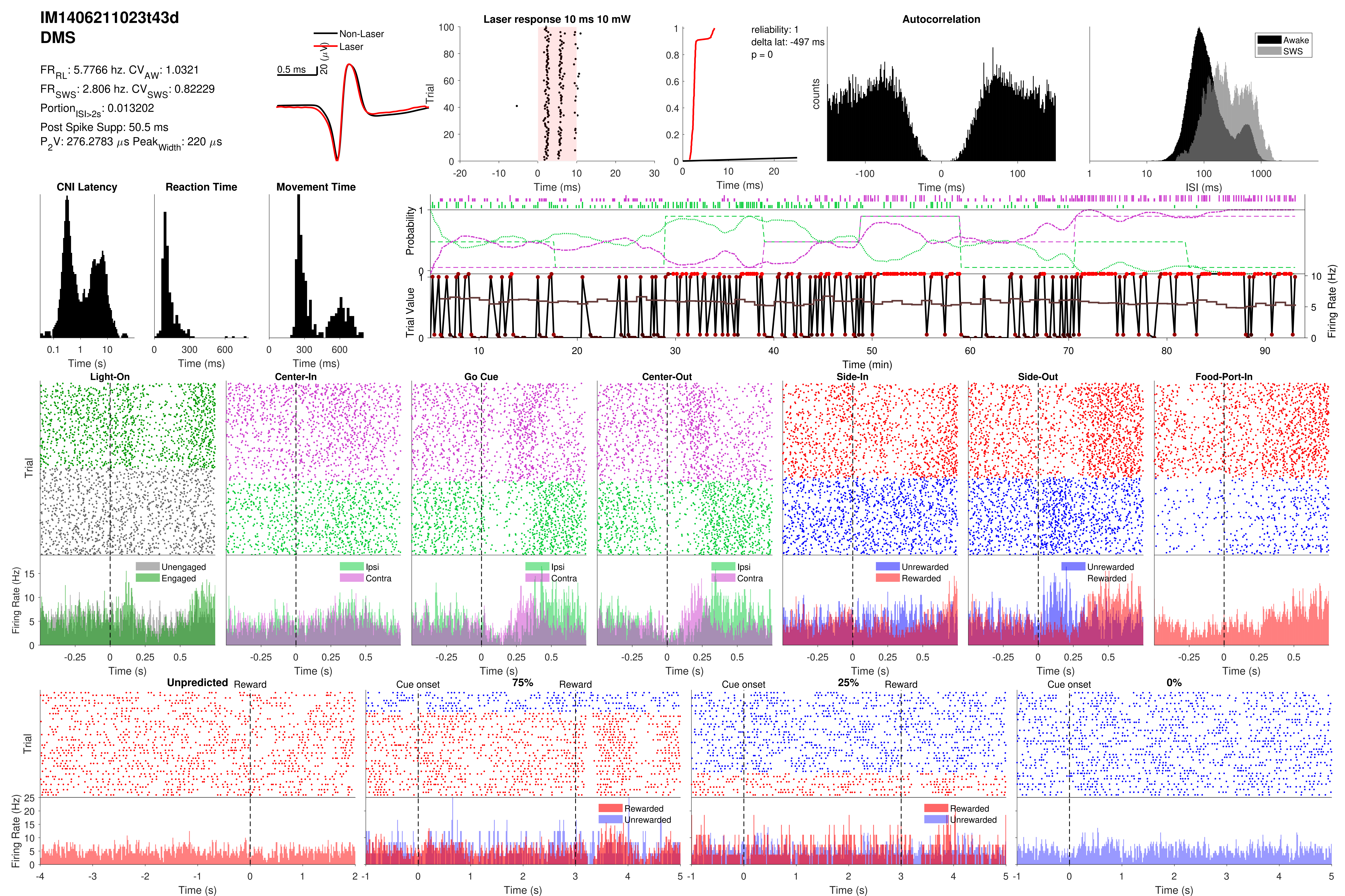
IM1406211023t43d
DMS
FR_RL: 5.7766 hz. CV_AW: 1.0321
FR_SWS: 2.806 hz. CV_SWS: 0.82229
Portion_ISI>2s: 0.013202
Post Spike Supp: 50.5 ms
P_2V: 276.2783 µs Peak_Width: 220 µs
Non-Laser
Laser
0.5 ms
20 (µV)
Laser response 10 ms 10 mW
reliability: 1
delta lat: -497 ms
p = 0
Autocorrelation
Awake
SWS
CNI Latency
Reaction Time
Movement Time
Probability
Trial Value
Firing Rate (Hz)
Time (min)
Light-On
Center-In
Go Cue
Center-Out
Side-In
Side-Out
Food-Port-In
Unengaged
Engaged
Ipsi
Contra
Unrewarded
Rewarded
Unpredicted
Reward
Cue onset
75%
25%
0%
Trial
counts
Time (ms)
Time (s)
ISI (ms)

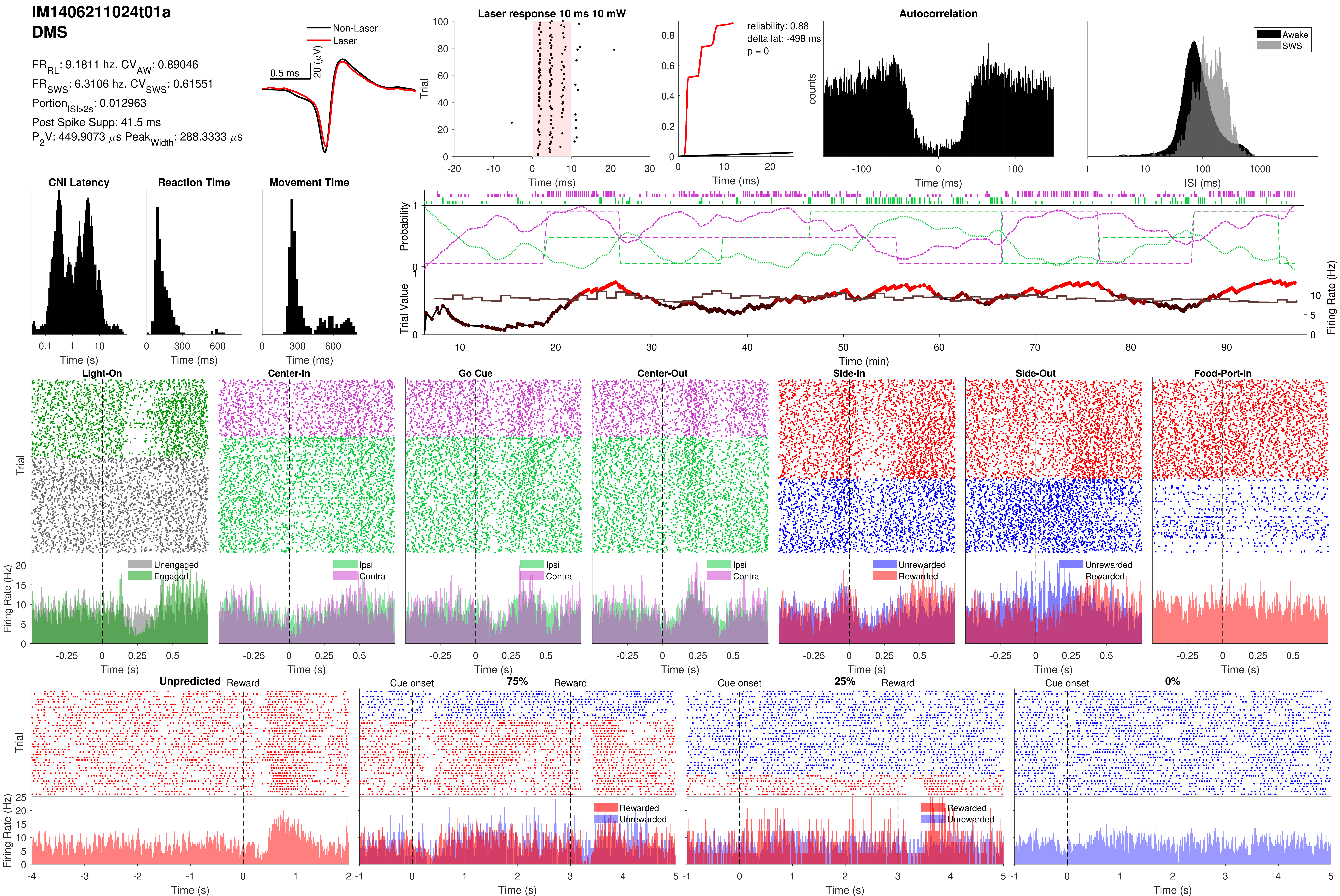
IM1406211024t01a
DMS
FR_RL: 9.1811 hz. CV_AW: 0.89046
FR_SWS: 6.3106 hz. CV_SWS: 0.61551
Portion_ISI>2s: 0.012963
Post Spike Supp: 41.5 ms
P_2V: 449.9073 μs Peak_Width: 288.3333 μs
Non-Laser
Laser
0.5 ms
20 (μV)
Laser response 10 ms 10 mW
Trial
Time (ms)
reliability: 0.88
delta lat: -498 ms
p = 0
Time (ms)
Autocorrelation
counts
Time (ms)
Awake
SWS
ISI (ms)
CNI Latency
Time (s)
Reaction Time
Time (ms)
Movement Time
Time (ms)
Probability
Trial Value
Firing Rate (Hz)
Time (min)
Light-On
Center-In
Go Cue
Center-Out
Side-In
Side-Out
Food-Port-In
Trial
Firing Rate (Hz)
Unengaged
Engaged
Ipsi
Contra
Unrewarded
Rewarded
Time (s)
Unpredicted
Reward
Cue onset
75%
25%
0%
Rewarded
Unrewarded
Time (s)

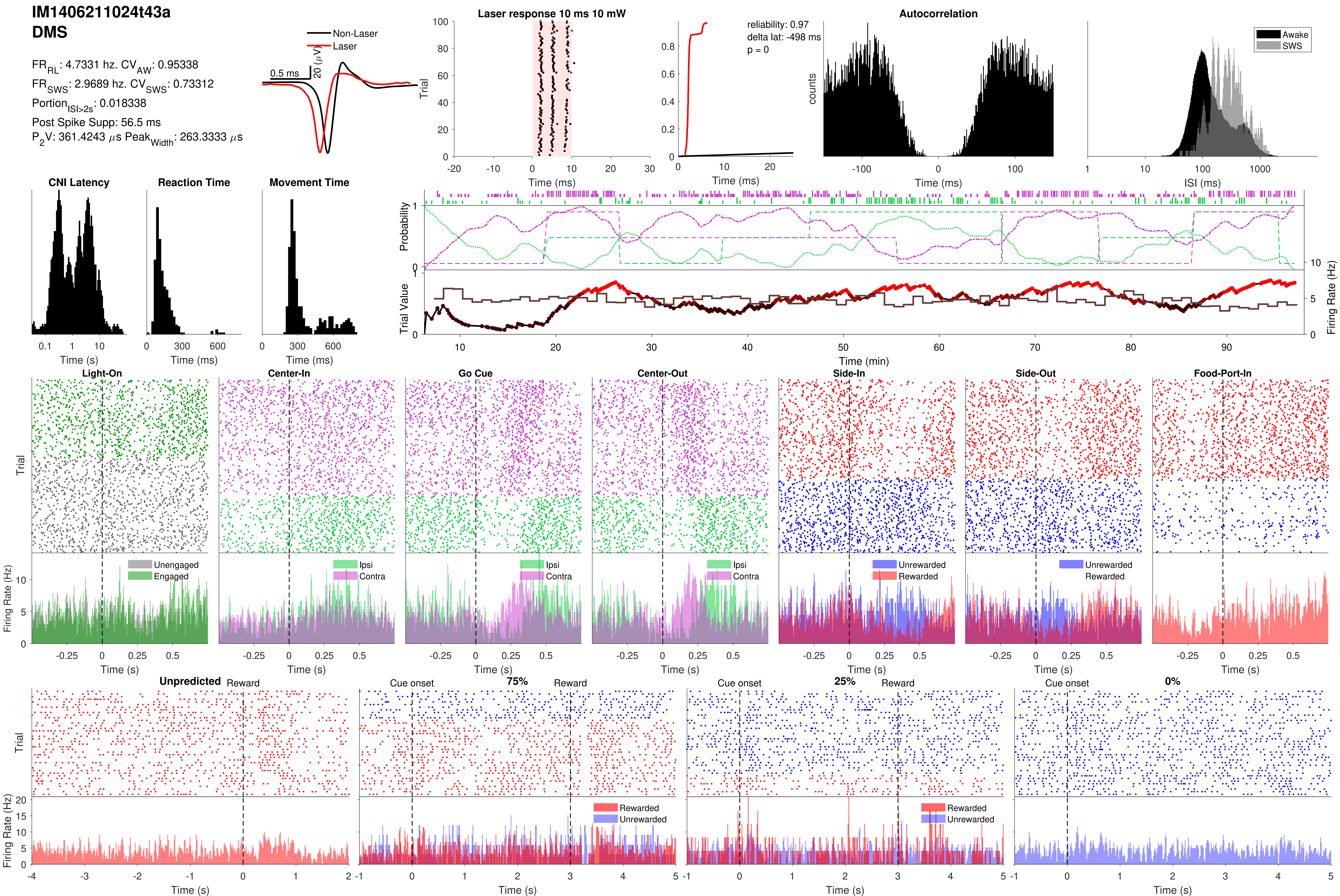
IM1406211024t43a
DMS
FR_RL: 4.7331 hz. CV_AW: 0.95338
FR_SWS: 2.9689 hz. CV_SWS: 0.73312
Portion_ISI>2s: 0.018338
Post Spike Supp: 56.5 ms
P_2V: 361.4243 μs Peak_Width: 263.3333 μs
Non-Laser
Laser
0.5 ms
20 (μV)
Laser response 10 ms 10 mW
Trial
Time (ms)
reliability: 0.97
delta lat: -498 ms
p = 0
Time (ms)
Autocorrelation
counts
Time (ms)
Awake
SWS
ISI (ms)
CNI Latency
Time (s)
Reaction Time
Time (ms)
Movement Time
Time (ms)
Probability
Trial Value
Firing Rate (Hz)
Time (min)
Light-On
Center-In
Go Cue
Center-Out
Side-In
Side-Out
Food-Port-In
Trial
Unengaged
Engaged
Ipsi
Contra
Unrewarded
Rewarded
Firing Rate (Hz)
Time (s)
Unpredicted
Reward
Cue onset
75%
25%
0%
Rewarded
Unrewarded
Trial
Firing Rate (Hz)
Time (s)

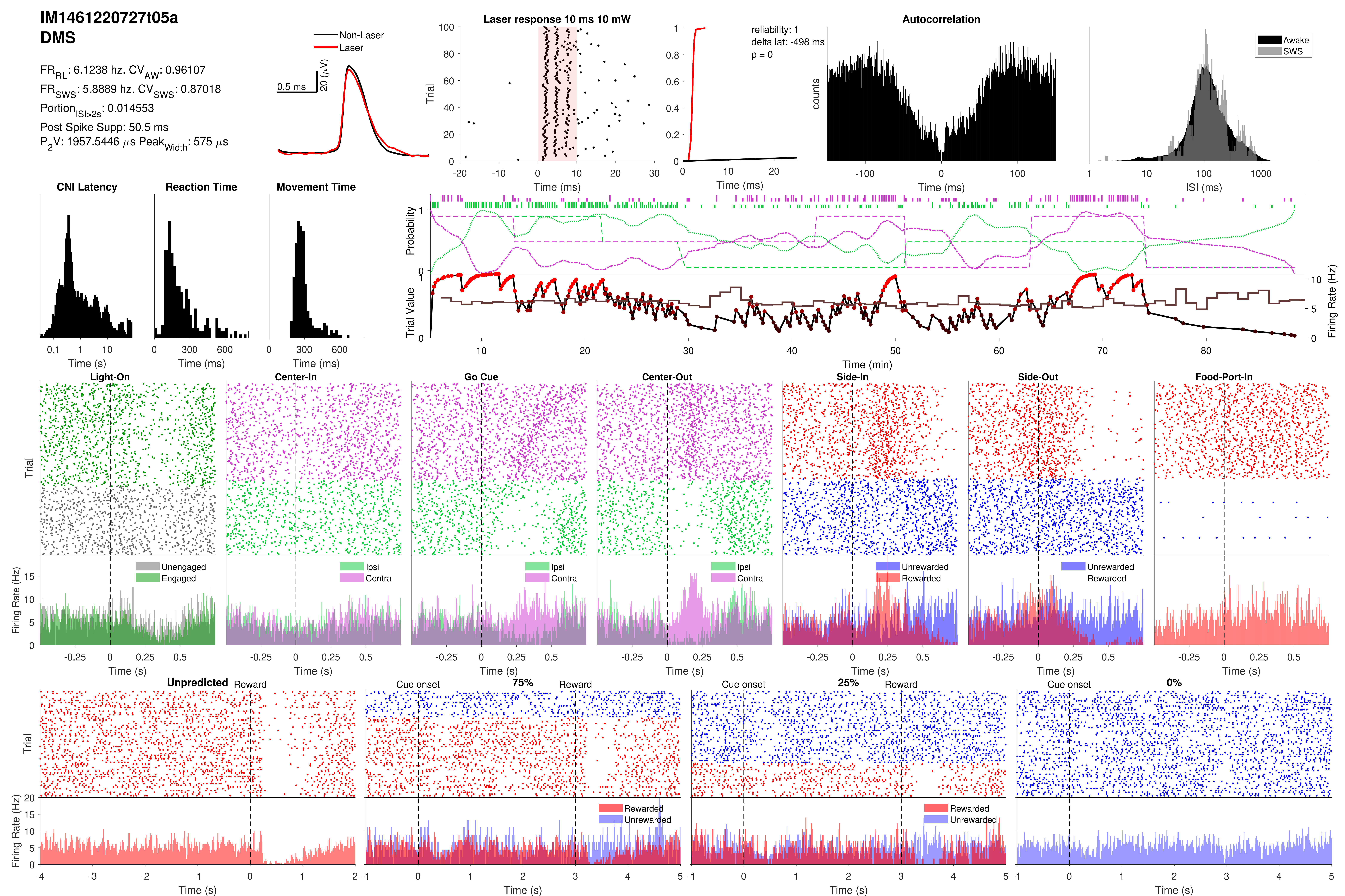

IM1461220727t05a
DMS
FR_RL: 6.1238 hz. CV_AW: 0.96107
FR_SWS: 5.8889 hz. CV_SWS: 0.87018
Portion_ISI>2s: 0.014553
Post Spike Supp: 50.5 ms
P_2V: 1957.5446 µs Peak_Width: 575 µs
Non-Laser
Laser
0.5 ms
20 (µV)
Laser response 10 ms 10 mW
Trial
Time (ms)
reliability: 1
delta lat: -498 ms
p = 0
Time (ms)
Autocorrelation
counts
Time (ms)
Awake
SWS
ISI (ms)
CNI Latency
Time (s)
Reaction Time
Time (ms)
Movement Time
Time (ms)
Probability
Trial Value
Firing Rate (Hz)
Time (min)
Light-On
Center-In
Go Cue
Center-Out
Side-In
Side-Out
Food-Port-In
Trial
Unengaged
Engaged
Ipsi
Contra
Unrewarded
Rewarded
Firing Rate (Hz)
Time (s)
Unpredicted
Reward
Cue onset
75%
25%
0%
Rewarded
Unrewarded
Trial
Firing Rate (Hz)
Time (s)

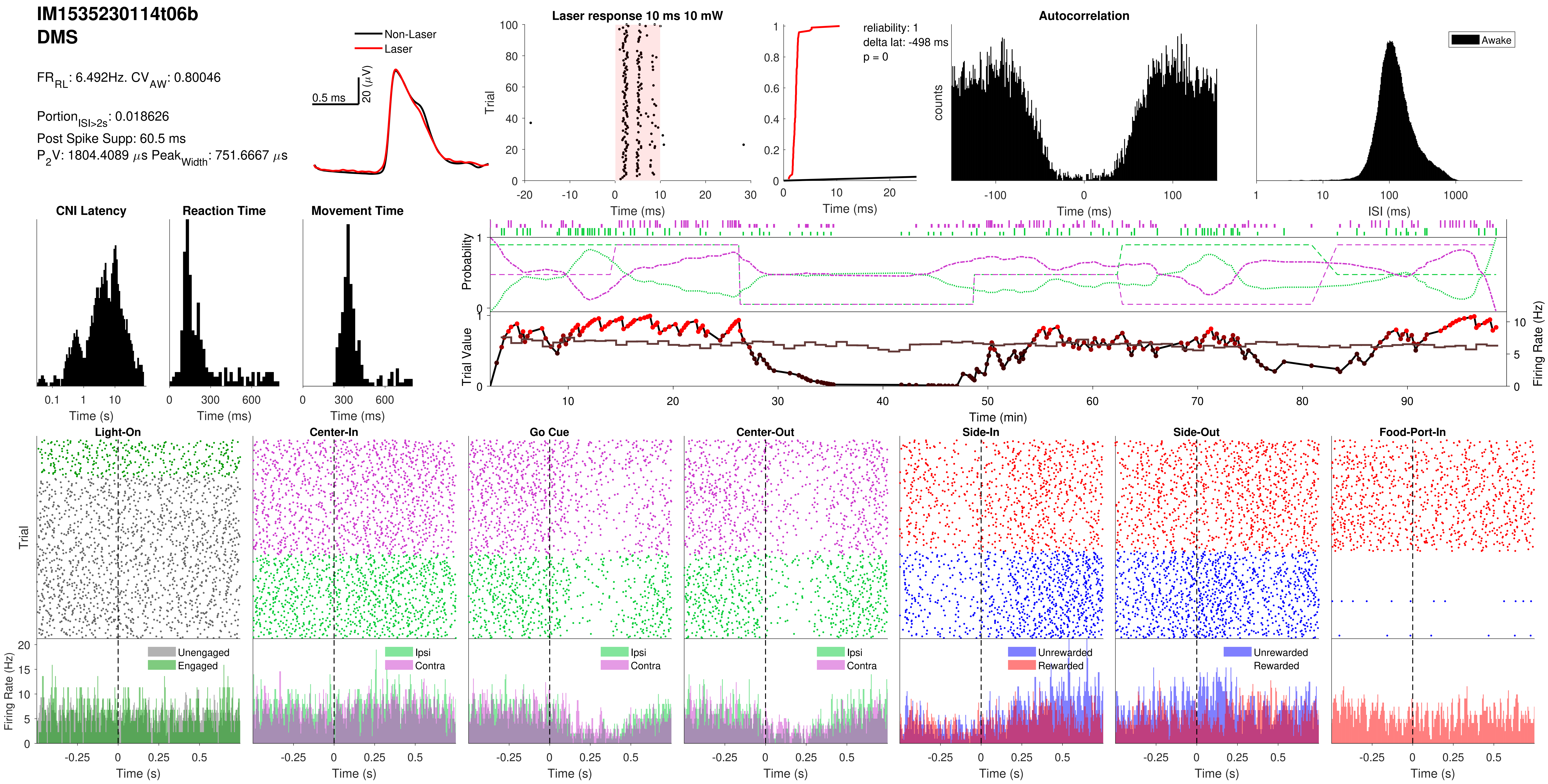

IM1535230114t06b
DMS
FR_RL: 6.492Hz. CV_AW: 0.80046
Portion_ISI>2s: 0.018626
Post Spike Supp: 60.5 ms
P_2V: 1804.4089 μs Peak_Width: 751.6667 μs
Non-Laser
Laser
0.5 ms
20 (μV)
Laser response 10 ms 10 mW
Trial
Time (ms)
reliability: 1
delta lat: -498 ms
p = 0
Time (ms)
Autocorrelation
counts
Time (ms)
Awake
ISI (ms)
CNI Latency
Time (s)
Reaction Time
Time (ms)
Movement Time
Time (ms)
Probability
Trial Value
Firing Rate (Hz)
Time (min)
Light-On
Center-In
Go Cue
Center-Out
Side-In
Side-Out
Food-Port-In
Trial
Firing Rate (Hz)
Unengaged
Engaged
Ipsi
Contra
Unrewarded
Rewarded
Time (s)

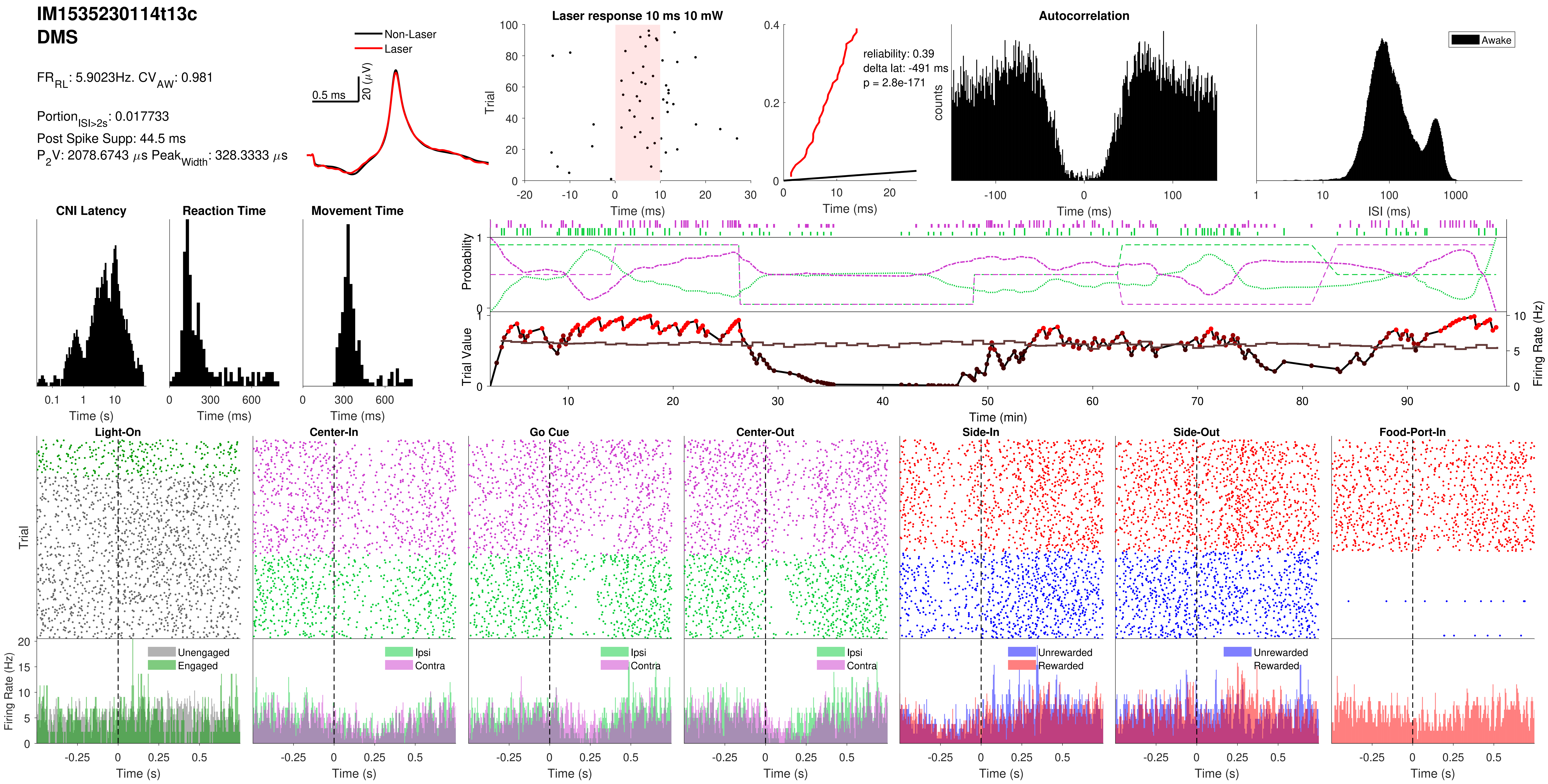

IM1535230114t13c
DMS
FR_RL: 5.9023Hz. CV_AW: 0.981
Portion_ISI>2s: 0.017733
Post Spike Supp: 44.5 ms
P_2V: 2078.6743 μs Peak_Width: 328.3333 μs
Non-Laser
Laser
0.5 ms
20 (μV)
Laser response 10 ms 10 mW
Trial
Time (ms)
reliability: 0.39
delta lat: -491 ms
p = 2.8e-171
Autocorrelation
counts
Awake
ISI (ms)
CNI Latency
Reaction Time
Movement Time
Time (s)
Probability
Trial Value
Firing Rate (Hz)
Time (min)
Light-On
Center-In
Go Cue
Center-Out
Side-In
Side-Out
Food-Port-In
Unengaged
Engaged
Ipsi
Contra
Unrewarded
Rewarded

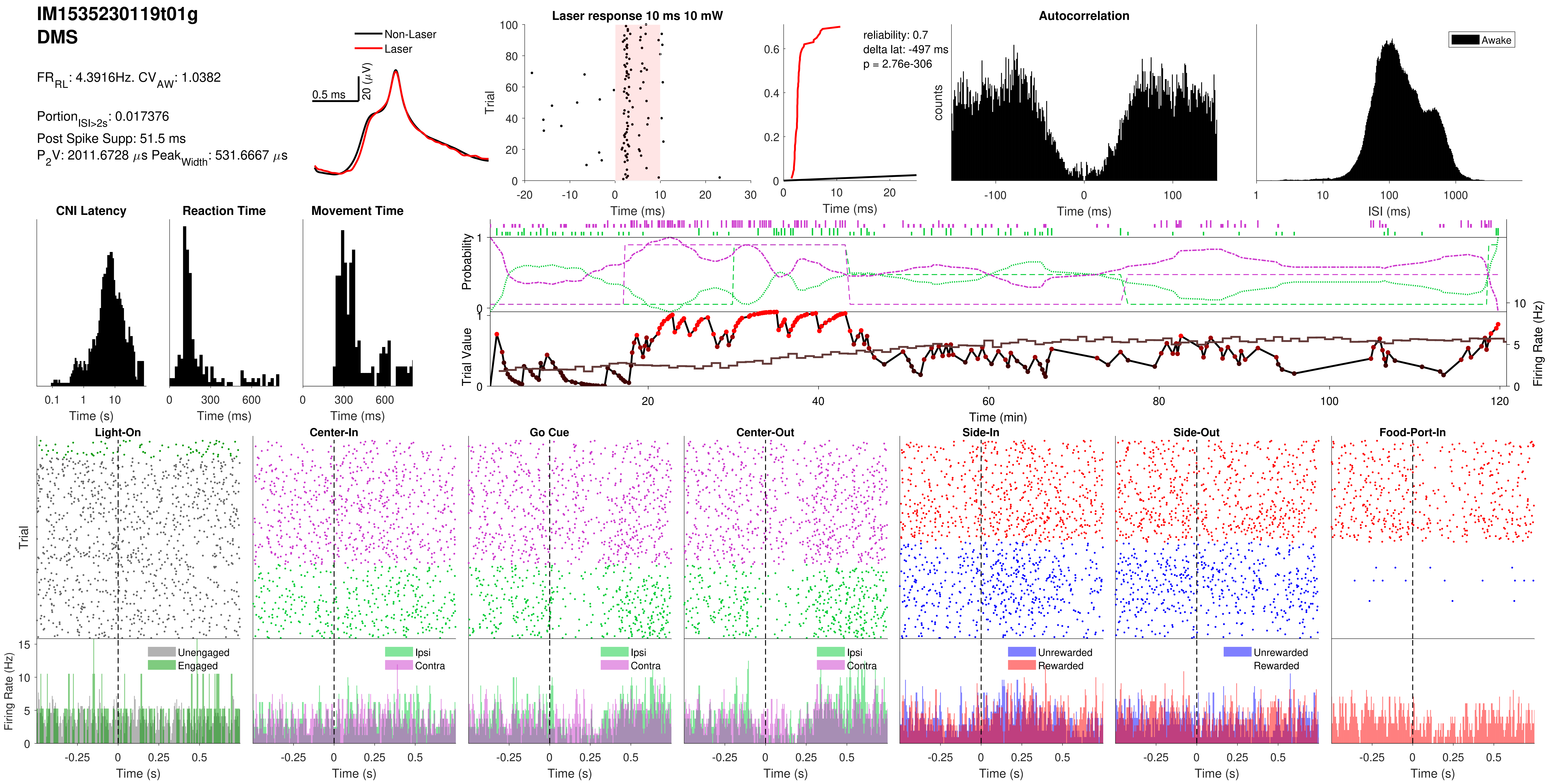
IM1535230119t01g
DMS
FR_RL: 4.3916Hz. CV_AW: 1.0382
Portion_ISI>2s: 0.017376
Post Spike Supp: 51.5 ms
P_2V: 2011.6728 μs Peak_Width: 531.6667 μs
Non-Laser
Laser
0.5 ms
20 (μV)
Laser response 10 ms 10 mW
Trial
Time (ms)
reliability: 0.7
delta lat: -497 ms
p = 2.76e-306
Autocorrelation
counts
Awake
ISI (ms)
CNI Latency
Reaction Time
Movement Time
Time (s)
Probability
Trial Value
Firing Rate (Hz)
Time (min)
Light-On
Center-In
Go Cue
Center-Out
Side-In
Side-Out
Food-Port-In
Unengaged
Engaged
Ipsi
Contra
Unrewarded
Rewarded
Firing Rate (Hz)

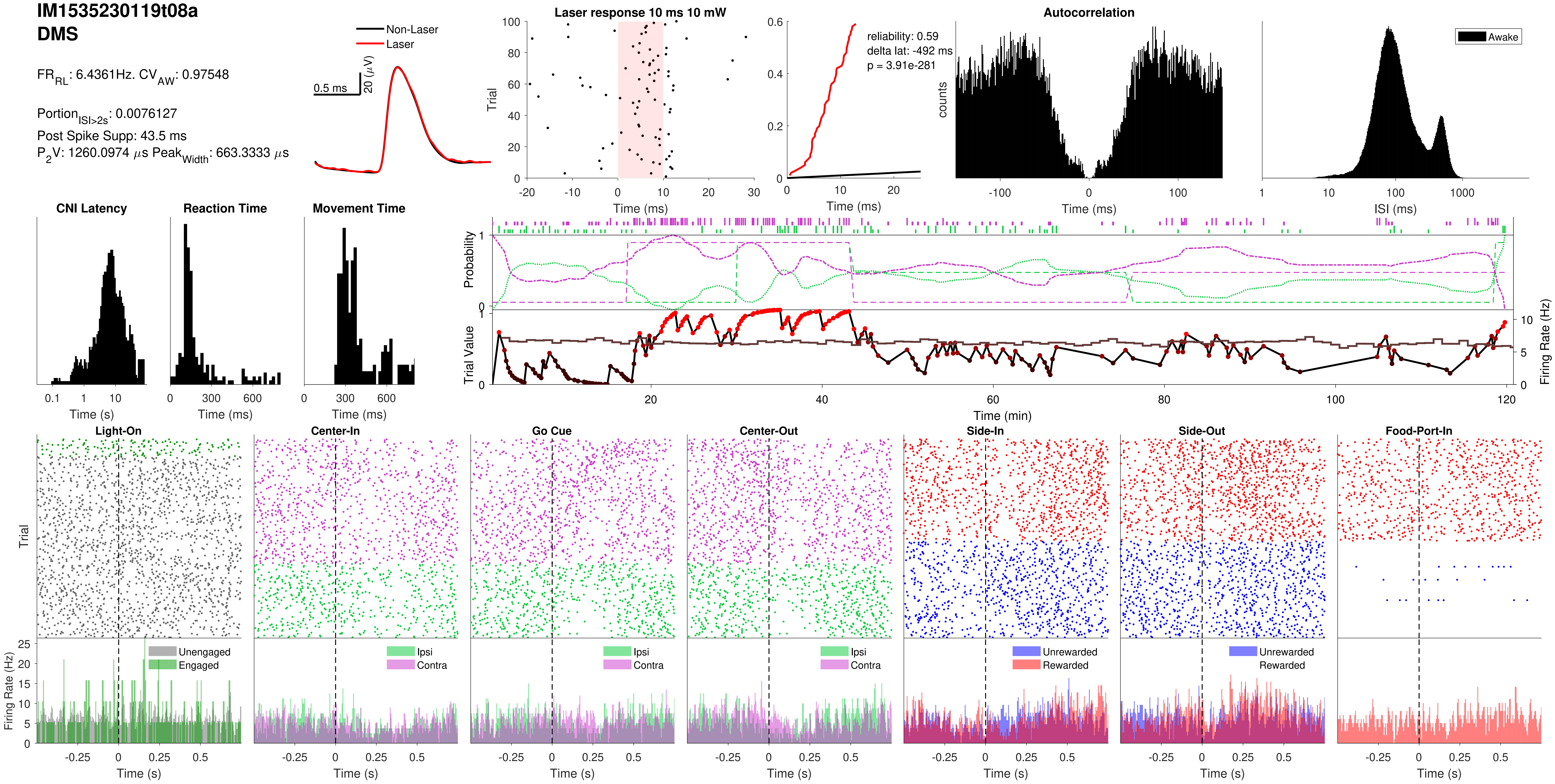
IM1535230119t08a
DMS
FR_RL: 6.4361Hz. CV_AW: 0.97548
Portion_ISI>2s: 0.0076127
Post Spike Supp: 43.5 ms
P2V: 1260.0974 μs Peak_Width: 663.3333 μs
Non-Laser
Laser
0.5 ms
20 (μV)
Laser response 10 ms 10 mW
Trial
Time (ms)
reliability: 0.59
delta lat: -492 ms
p = 3.91e-281
Time (ms)
Autocorrelation
counts
Time (ms)
Awake
ISI (ms)
CNI Latency
Time (s)
Reaction Time
Time (ms)
Movement Time
Time (ms)
Probability
Trial Value
Firing Rate (Hz)
Time (min)
Light-On
Center-In
Go Cue
Center-Out
Side-In
Side-Out
Food-Port-In
Trial
Firing Rate (Hz)
Unengaged
Engaged
Ipsi
Contra
Unrewarded
Rewarded
Time (s)

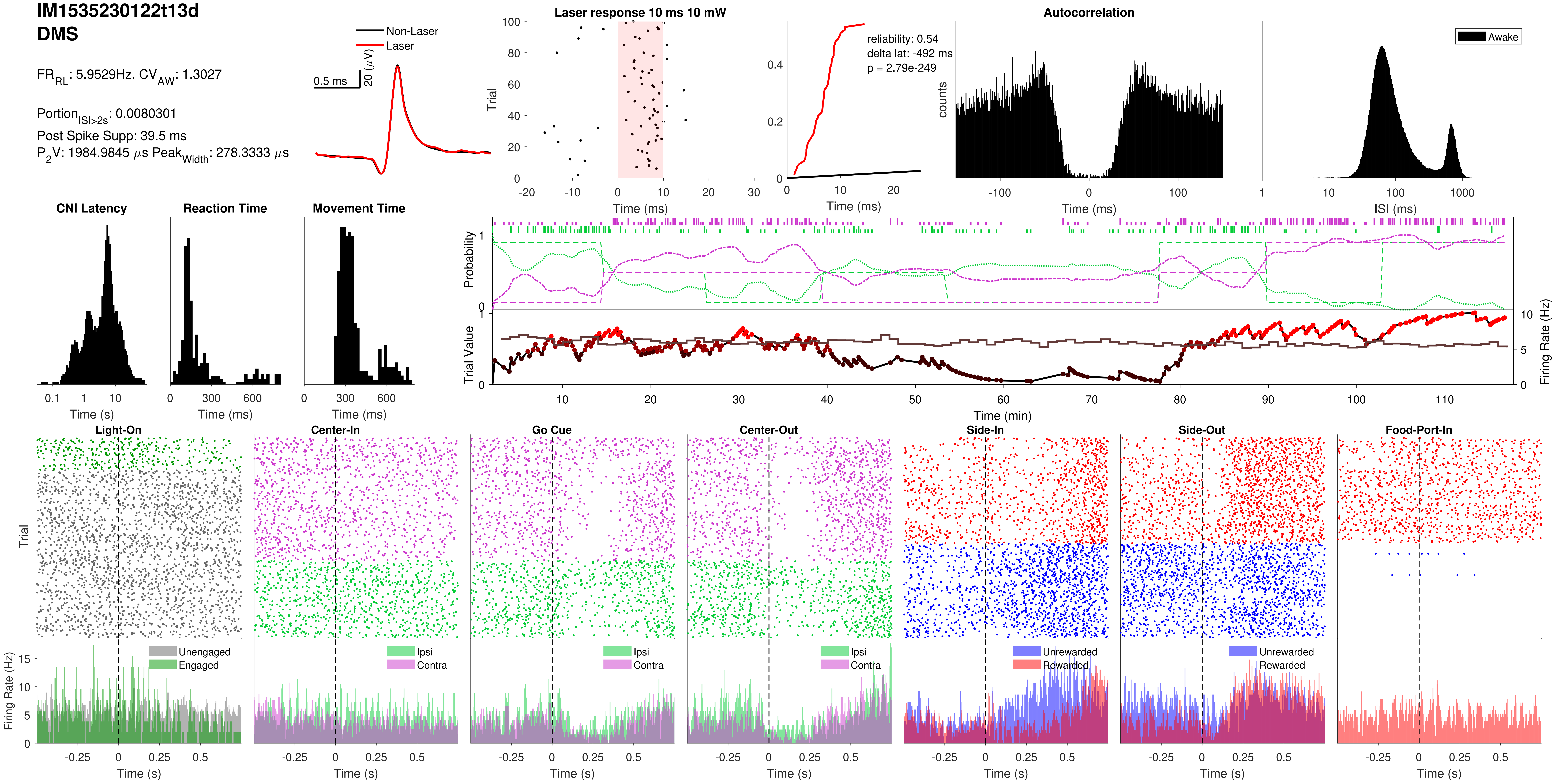
IM1535230122t13d
DMS
FR_RL: 5.9529Hz. CV_AW: 1.3027
Portion_ISI>2s: 0.0080301
Post Spike Supp: 39.5 ms
P_2V: 1984.9845 μs Peak_Width: 278.3333 μs
Non-Laser
Laser
0.5 ms
20 (μV)
Laser response 10 ms 10 mW
Trial
Time (ms)
reliability: 0.54
delta lat: -492 ms
p = 2.79e-249
Time (ms)
Autocorrelation
counts
Time (ms)
Awake
ISI (ms)
CNI Latency
Time (s)
Reaction Time
Time (ms)
Movement Time
Time (ms)
Probability
Trial Value
Firing Rate (Hz)
Time (min)
Light-On
Center-In
Go Cue
Center-Out
Side-In
Side-Out
Food-Port-In
Trial
Firing Rate (Hz)
Unengaged
Engaged
Ipsi
Contra
Unrewarded
Rewarded
Time (s)

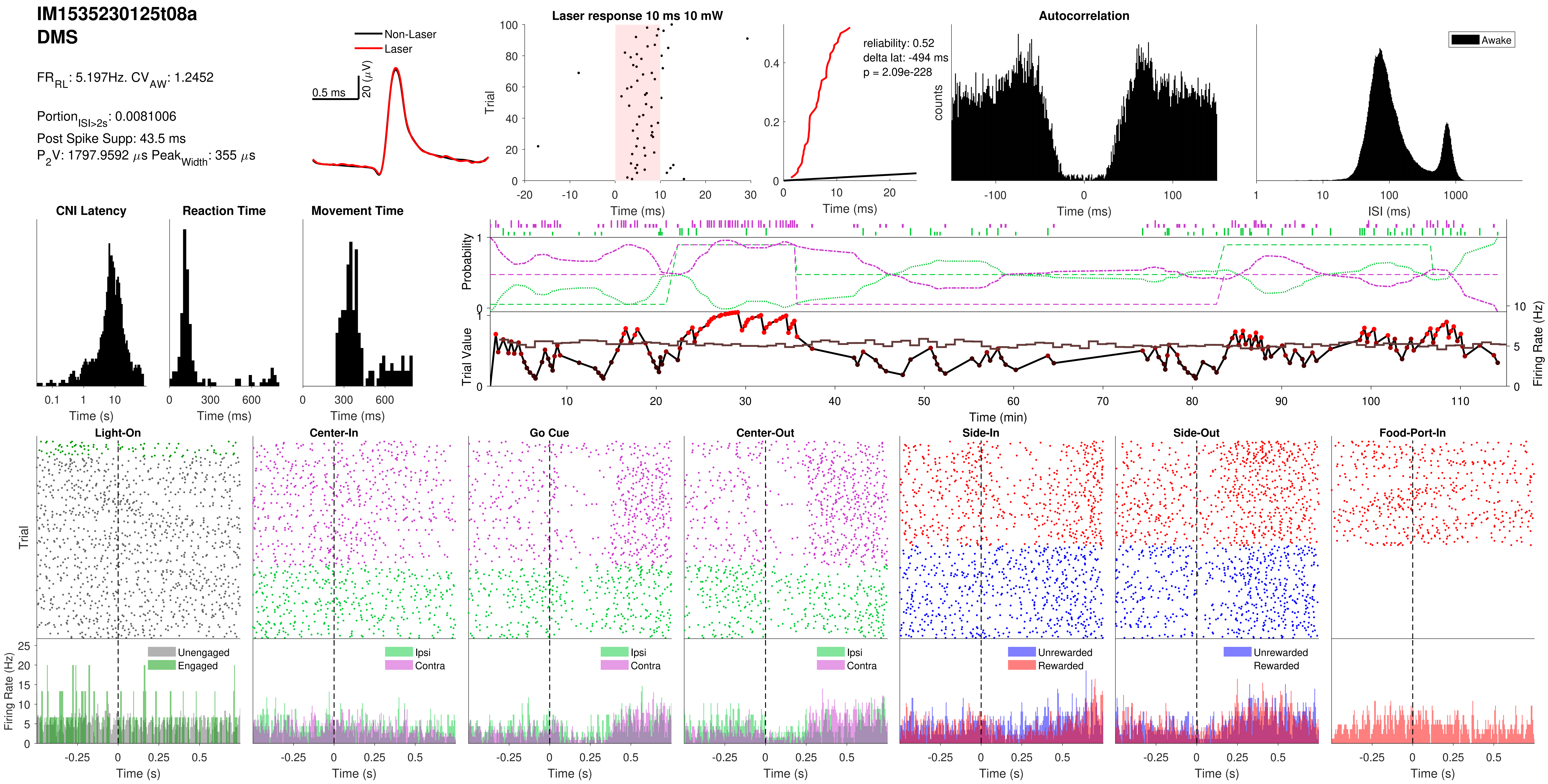
IM1535230125t08a
DMS
$FR_{RL}$: 5.197Hz. $CV_{AW}$: 1.2452
$Portion_{ISI>2s}$: 0.0081006
Post Spike Supp: 43.5 ms
$P_2V$: 1797.9592 $\mu$s $Peak_{Width}$: 355 $\mu$s
Non-Laser
Laser
0.5 ms
20 ($\mu$V)
Laser response 10 ms 10 mW
Trial
Time (ms)
reliability: 0.52
delta lat: -494 ms
p = 2.09e-228
Time (ms)
Autocorrelation
counts
Time (ms)
Awake
ISI (ms)
CNI Latency
Time (s)
Reaction Time
Time (ms)
Movement Time
Time (ms)
Probability
Trial Value
Firing Rate (Hz)
Time (min)
Light-On
Center-In
Go Cue
Center-Out
Side-In
Side-Out
Food-Port-In
Trial
Firing Rate (Hz)
Unengaged
Engaged
Ipsi
Contra
Unrewarded
Rewarded
Time (s)

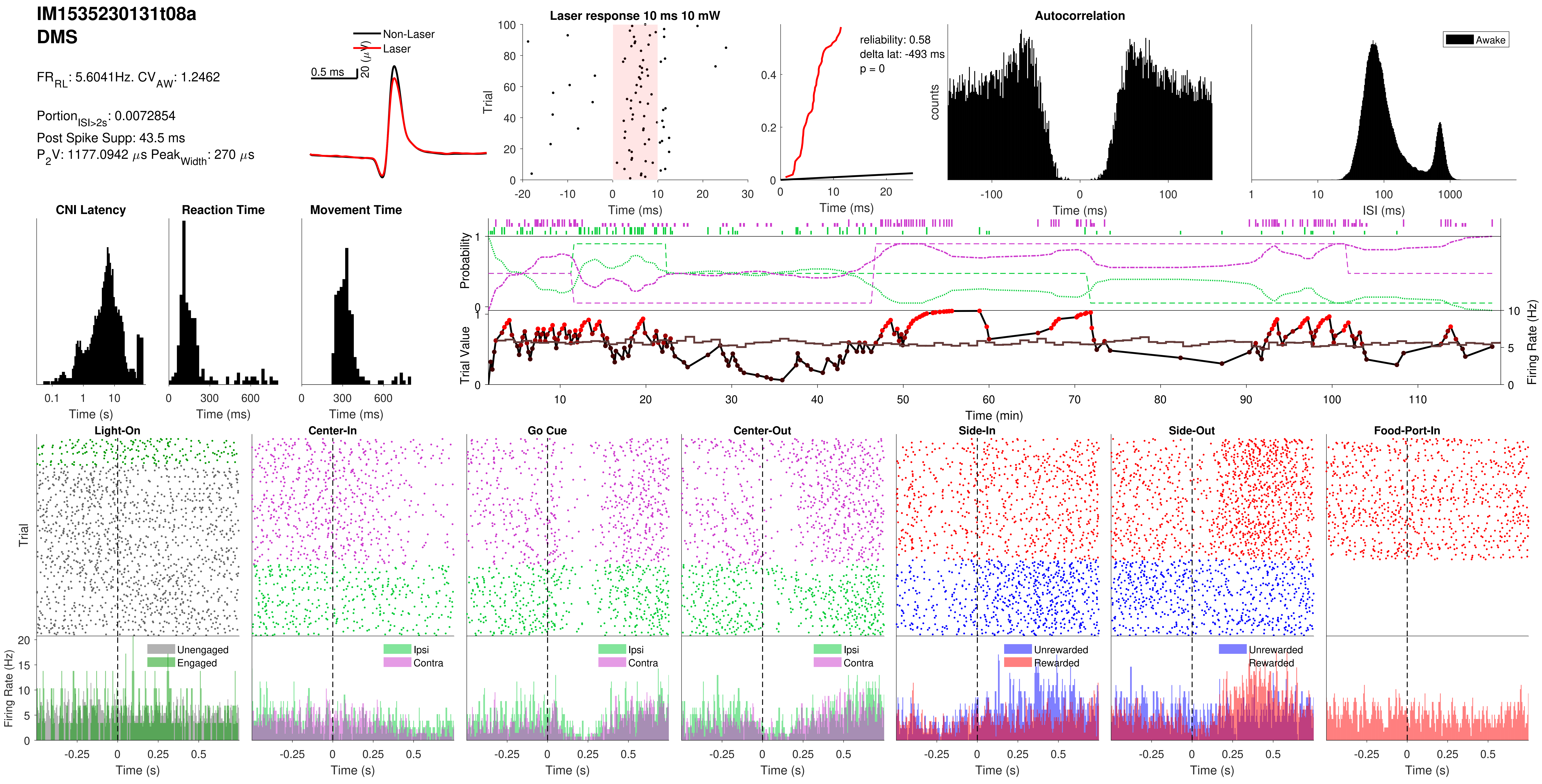
IM1535230131t08a
DMS
FR_RL: 5.6041Hz. CV_AW: 1.2462
Portion_ISI>2s: 0.0072854
Post Spike Supp: 43.5 ms
P_2V: 1177.0942 µs Peak_Width: 270 µs
Non-Laser
Laser
0.5 ms
20 (µV)
Laser response 10 ms 10 mW
Trial
Time (ms)
reliability: 0.58
delta lat: -493 ms
p = 0
Time (ms)
Autocorrelation
counts
Time (ms)
Awake
ISI (ms)
CNI Latency
Time (s)
Reaction Time
Time (ms)
Movement Time
Time (ms)
Probability
Trial Value
Firing Rate (Hz)
Time (min)
Light-On
Center-In
Go Cue
Center-Out
Side-In
Side-Out
Food-Port-In
Trial
Firing Rate (Hz)
Unengaged
Engaged
Ipsi
Contra
Unrewarded
Rewarded
Time (s)

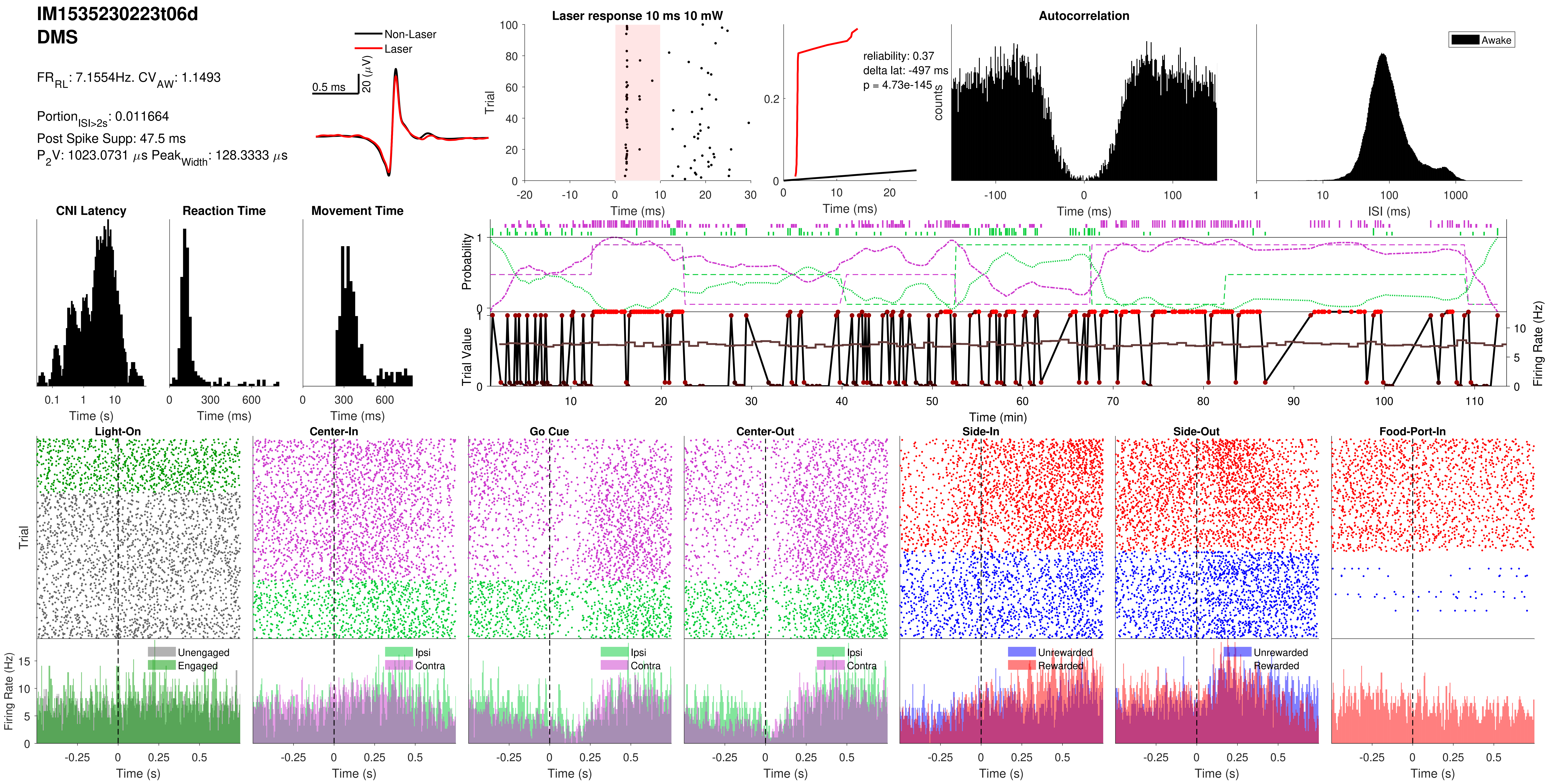
IM1535230223t06d
DMS
FR_RL: 7.1554Hz. CV_AW: 1.1493
Portion_ISI>2s: 0.011664
Post Spike Supp: 47.5 ms
P_2V: 1023.0731 µs Peak_Width: 128.3333 µs
Non-Laser
Laser
0.5 ms
20 (µV)
Laser response 10 ms 10 mW
Trial
Time (ms)
reliability: 0.37
delta lat: -497 ms
p = 4.73e-145
Time (ms)
Autocorrelation
counts
Time (ms)
Awake
ISI (ms)
CNI Latency
Time (s)
Reaction Time
Time (ms)
Movement Time
Time (ms)
Probability
Trial Value
Firing Rate (Hz)
Time (min)
Light-On
Center-In
Go Cue
Center-Out
Side-In
Side-Out
Food-Port-In
Trial
Unengaged
Engaged
Ipsi
Contra
Unrewarded
Rewarded
Firing Rate (Hz)
Time (s)

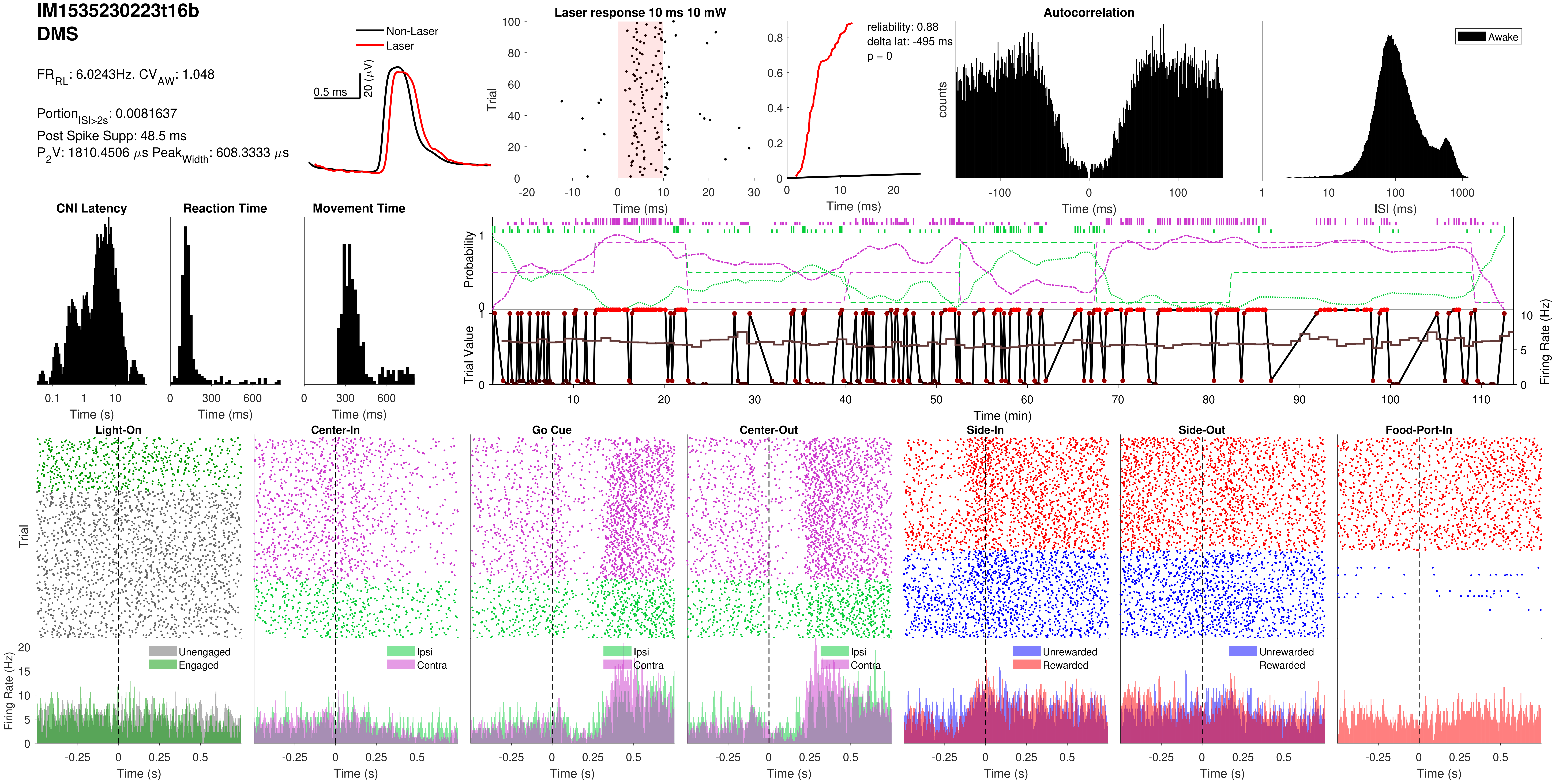
IM1535230223t16b
DMS
FR_RL: 6.0243Hz. CV_AW: 1.048
Portion_ISI>2s: 0.0081637
Post Spike Supp: 48.5 ms
P2V: 1810.4506 μs Peak_Width: 608.3333 μs
Non-Laser
Laser
0.5 ms
20 (μV)
Laser response 10 ms 10 mW
Trial
Time (ms)
reliability: 0.88
delta lat: -495 ms
p = 0
Time (ms)
Autocorrelation
counts
Time (ms)
Awake
ISI (ms)
CNI Latency
Time (s)
Reaction Time
Time (ms)
Movement Time
Time (ms)
Probability
Trial Value
Firing Rate (Hz)
Time (min)
Light-On
Center-In
Go Cue
Center-Out
Side-In
Side-Out
Food-Port-In
Trial
Unengaged
Engaged
Ipsi
Contra
Ipsi
Contra
Ipsi
Contra
Unrewarded
Rewarded
Unrewarded
Rewarded
Firing Rate (Hz)
Time (s)

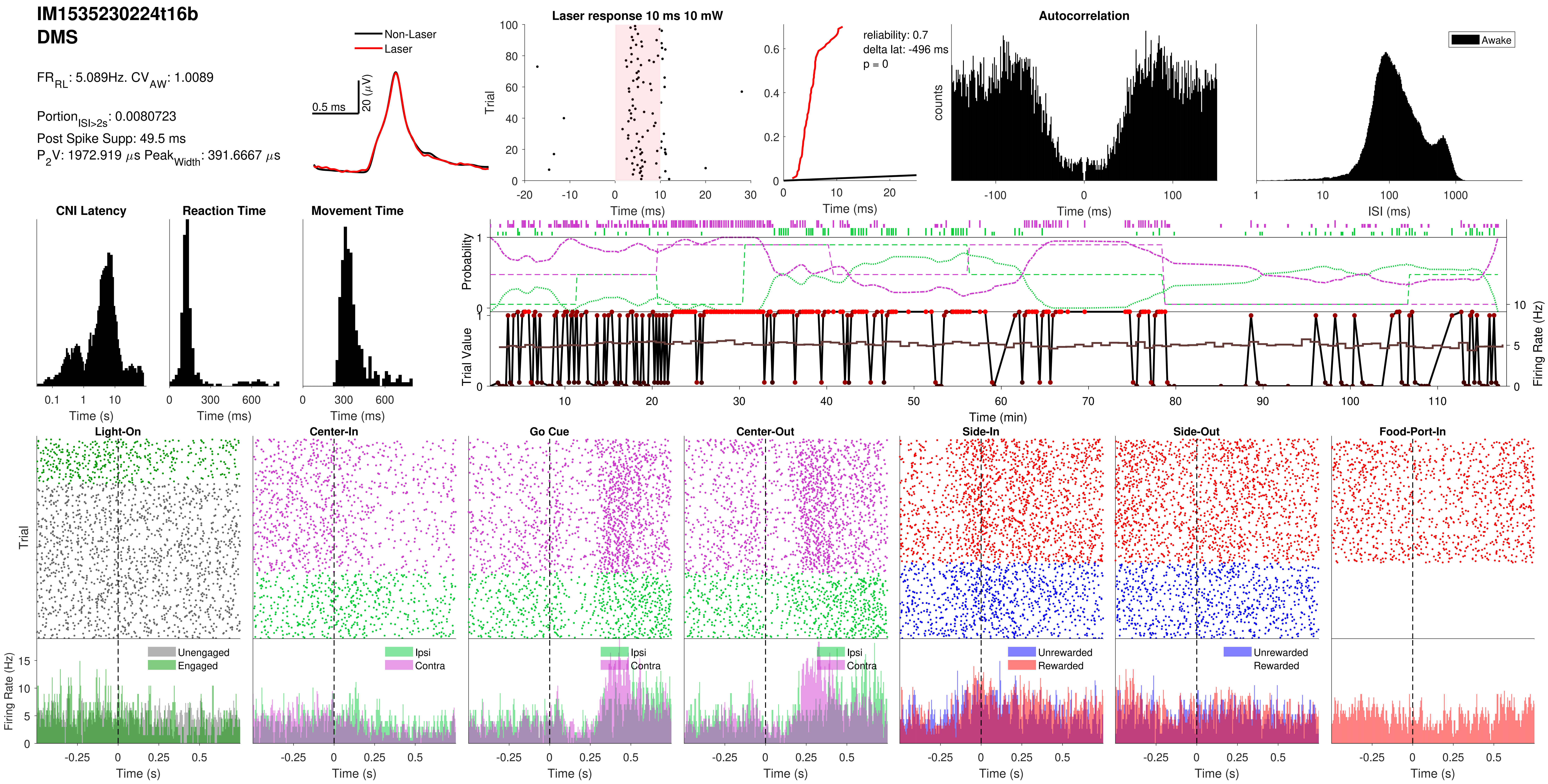
IM1535230224t16b
DMS
FR_RL: 5.089Hz. CV_AW: 1.0089
Portion_ISI>2s: 0.0080723
Post Spike Supp: 49.5 ms
P_2V: 1972.919 μs Peak_Width: 391.6667 μs
Non-Laser
Laser
0.5 ms
20 (μV)
Laser response 10 ms 10 mW
Trial
Time (ms)
reliability: 0.7
delta lat: -496 ms
p = 0
Time (ms)
Autocorrelation
counts
Time (ms)
Awake
ISI (ms)
CNI Latency
Time (s)
Reaction Time
Time (ms)
Movement Time
Time (ms)
Probability
Trial Value
Firing Rate (Hz)
Time (min)
Light-On
Center-In
Go Cue
Center-Out
Side-In
Side-Out
Food-Port-In
Trial
Firing Rate (Hz)
Unengaged
Engaged
Ipsi
Contra
Unrewarded
Rewarded
Time (s)

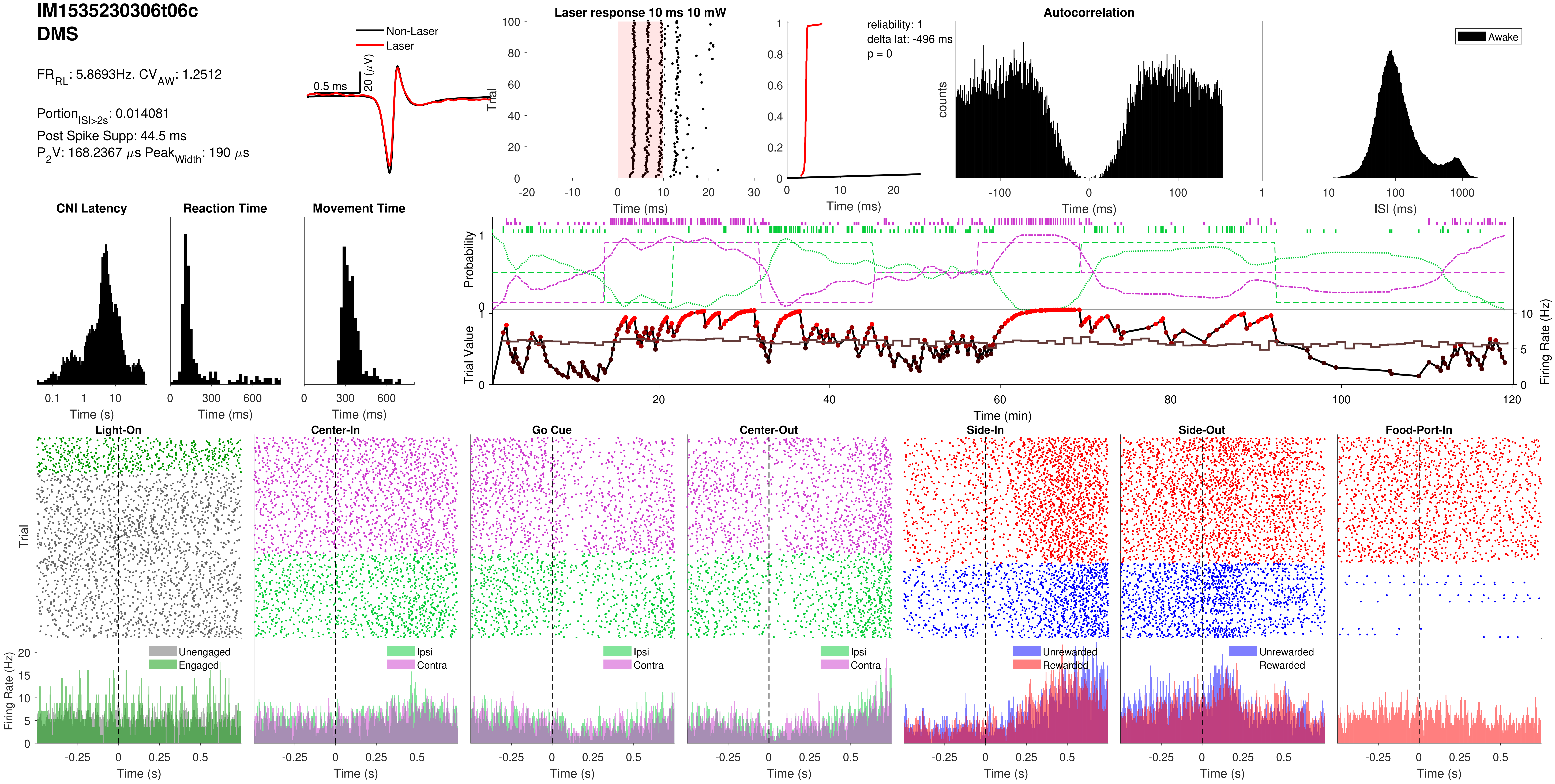

IM1535230306t06c
DMS
FR_RL: 5.8693Hz. CV_AW: 1.2512
Portion_ISI>2s: 0.014081
Post Spike Supp: 44.5 ms
P_2V: 168.2367 μs Peak_Width: 190 μs
Non-Laser
Laser
0.5 ms
20 (μV)
Laser response 10 ms 10 mW
Trial
Time (ms)
reliability: 1
delta lat: -496 ms
p = 0
Autocorrelation
counts
Awake
ISI (ms)
CNI Latency
Reaction Time
Movement Time
Time (s)
Time (ms)
Probability
Trial Value
Firing Rate (Hz)
Time (min)
Light-On
Center-In
Go Cue
Center-Out
Side-In
Side-Out
Food-Port-In
Unengaged
Engaged
Ipsi
Contra
Unrewarded
Rewarded
Time (s)

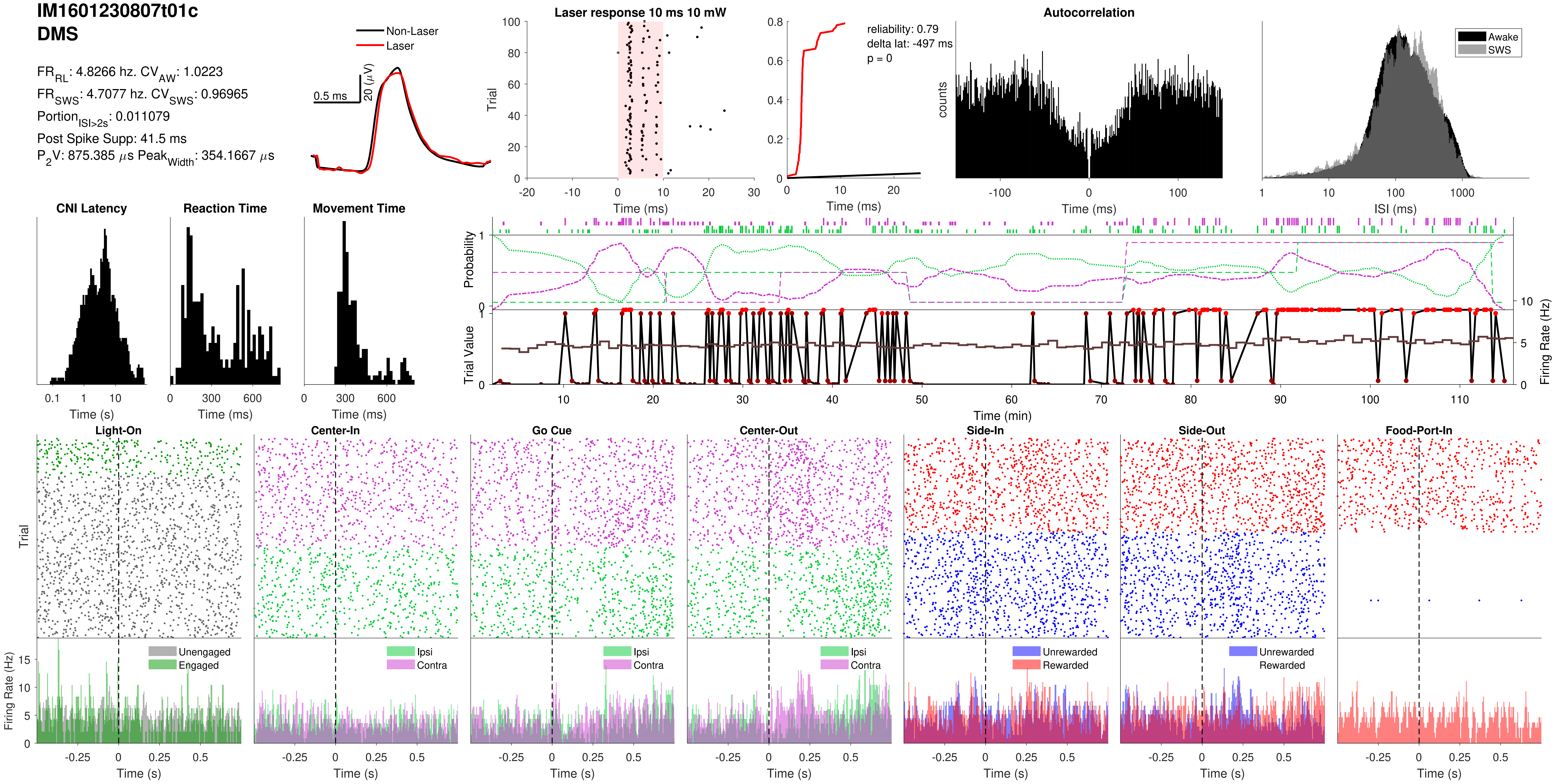

IM1601230807t01c
DMS
FR_RL: 4.8266 hz. CV_AW: 1.0223
FR_SWS: 4.7077 hz. CV_SWS: 0.96965
Portion_ISI>2s: 0.011079
Post Spike Supp: 41.5 ms
P_2V: 875.385 μs Peak_Width: 354.1667 μs
Non-Laser
Laser
0.5 ms
20 (μV)
Laser response 10 ms 10 mW
Trial
Time (ms)
reliability: 0.79
delta lat: -497 ms
p = 0
Time (ms)
Autocorrelation
counts
Time (ms)
Awake
SWS
ISI (ms)
CNI Latency
Time (s)
Reaction Time
Time (ms)
Movement Time
Time (ms)
Probability
Trial Value
Firing Rate (Hz)
Time (min)
Light-On
Center-In
Go Cue
Center-Out
Side-In
Side-Out
Food-Port-In
Trial
Firing Rate (Hz)
Unengaged
Engaged
Ipsi
Contra
Unrewarded
Rewarded
Time (s)

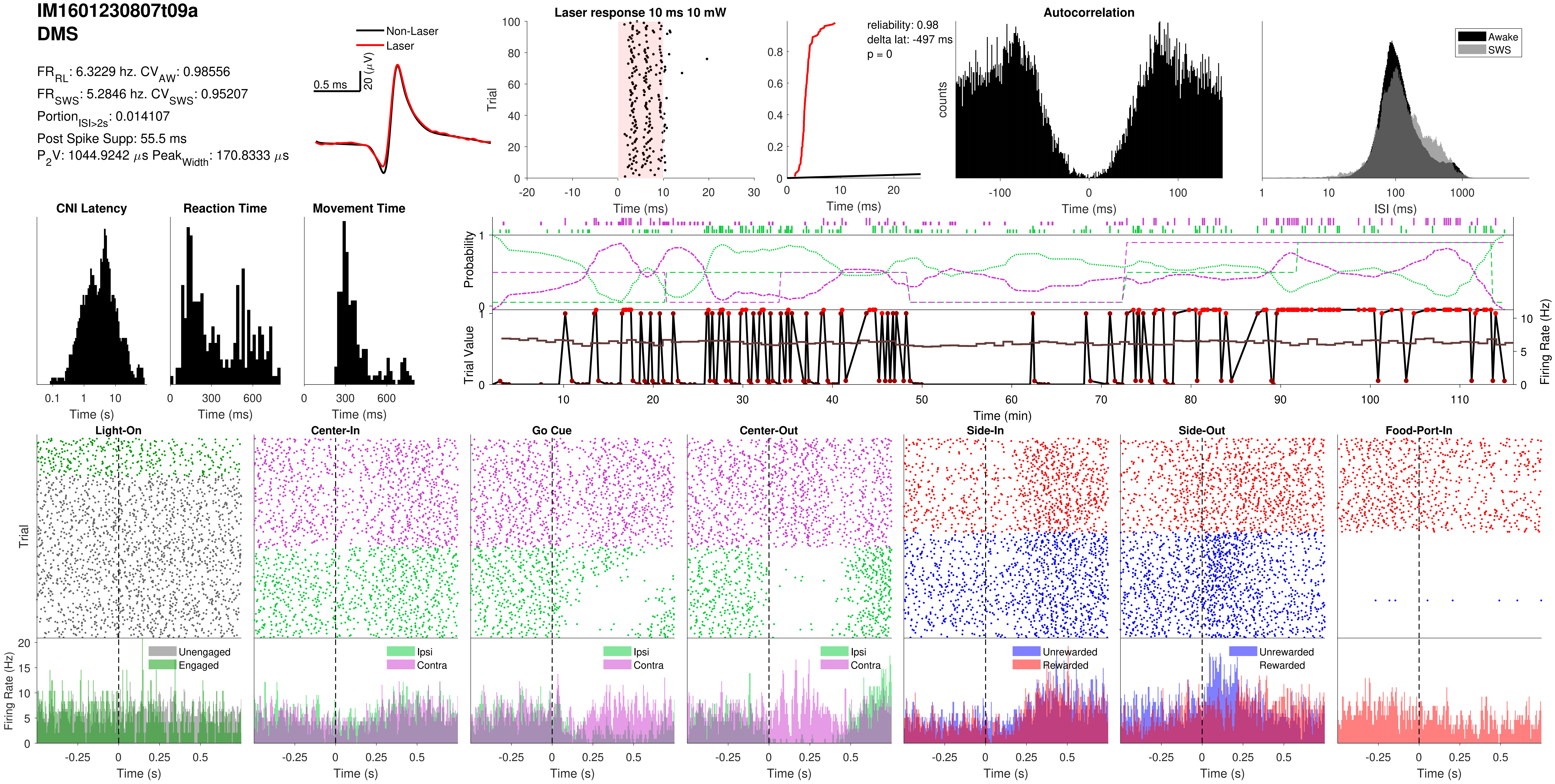
IM1601230807t09a
DMS
FR_RL: 6.3229 hz. CV_AW: 0.98556
FR_SWS: 5.2846 hz. CV_SWS: 0.95207
Portion_ISI>2s: 0.014107
Post Spike Supp: 55.5 ms
P_2V: 1044.9242 µs Peak_Width: 170.8333 µs
Non-Laser
Laser
0.5 ms
20 (µV)
Laser response 10 ms 10 mW
Trial
Time (ms)
reliability: 0.98
delta lat: -497 ms
p = 0
Time (ms)
Autocorrelation
counts
Time (ms)
Awake
SWS
ISI (ms)
CNI Latency
Time (s)
Reaction Time
Time (ms)
Movement Time
Time (ms)
Probability
Trial Value
Firing Rate (Hz)
Time (min)
Light-On
Center-In
Go Cue
Center-Out
Side-In
Side-Out
Food-Port-In
Trial
Firing Rate (Hz)
Unengaged
Engaged
Ipsi
Contra
Unrewarded
Rewarded
Time (s)

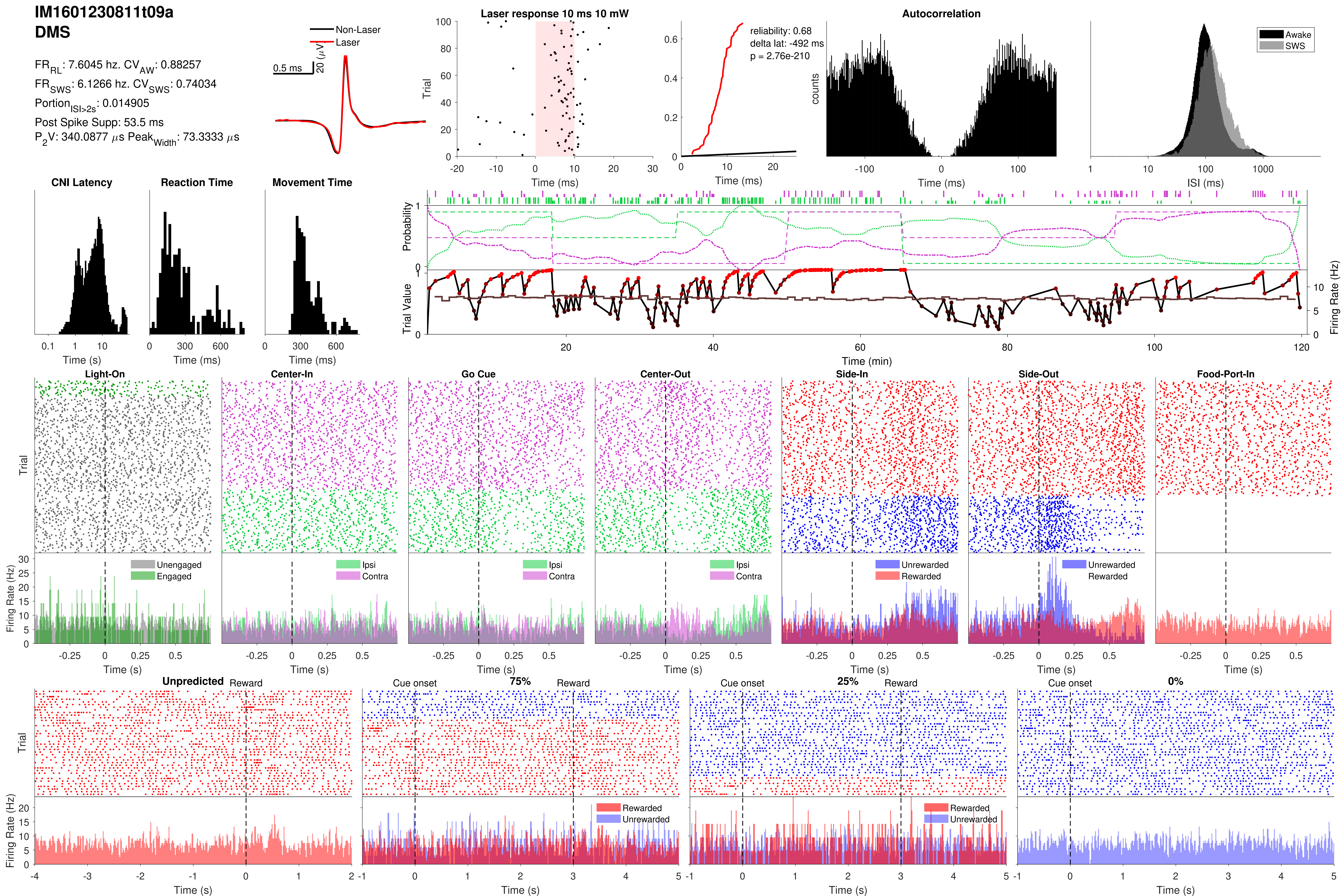
IM1601230811t09a
DMS
FR_RL: 7.6045 hz. CV_AW: 0.88257
FR_SWS: 6.1266 hz. CV_SWS: 0.74034
Portion_ISI>2s: 0.014905
Post Spike Supp: 53.5 ms
P_2V: 340.0877 μs Peak_Width: 73.3333 μs
Non-Laser
Laser
0.5 ms
20 (μV)
Laser response 10 ms 10 mW
Trial
Time (ms)
reliability: 0.68
delta lat: -492 ms
p = 2.76e-210
Time (ms)
Autocorrelation
counts
Time (ms)
Awake
SWS
ISI (ms)
CNI Latency
Time (s)
Reaction Time
Time (ms)
Movement Time
Time (ms)
Probability
Trial Value
Firing Rate (Hz)
Time (min)
Light-On
Center-In
Go Cue
Center-Out
Side-In
Side-Out
Food-Port-In
Trial
Firing Rate (Hz)
Unengaged
Engaged
Ipsi
Contra
Unrewarded
Rewarded
Time (s)
Unpredicted
Reward
Cue onset
75%
25%
0%
Rewarded
Unrewarded
Trial
Firing Rate (Hz)
Time (s)

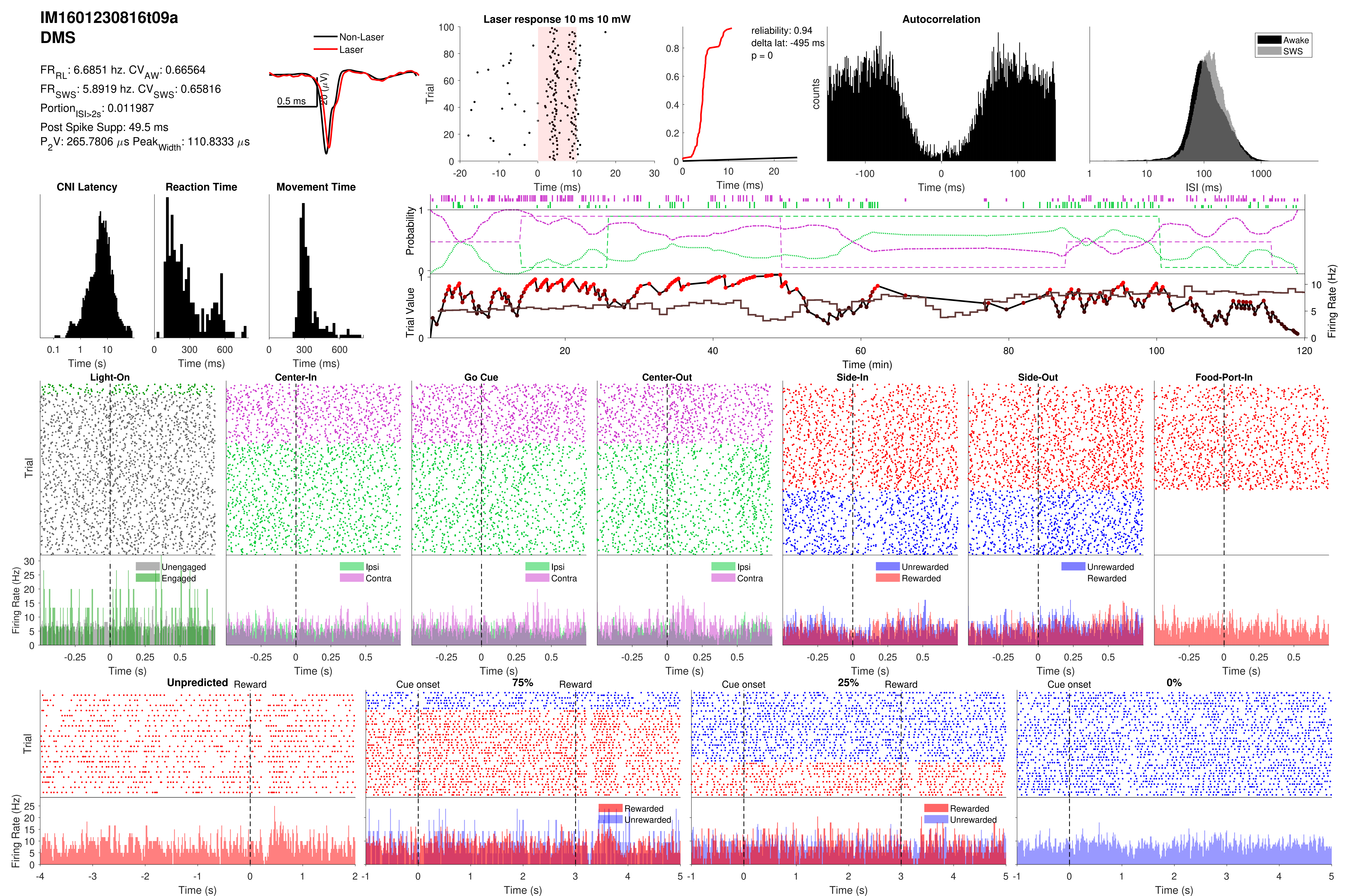
IM1601230816t09a
DMS
FR_RL: 6.6851 hz. CV_AW: 0.66564
FR_SWS: 5.8919 hz. CV_SWS: 0.65816
Portion_ISI>2s: 0.011987
Post Spike Supp: 49.5 ms
P_2V: 265.7806 µs Peak_Width: 110.8333 µs
Non-Laser
Laser
0.5 ms
20 (µV)
Laser response 10 ms 10 mW
reliability: 0.94
delta lat: -495 ms
p = 0
Autocorrelation
Awake
SWS
counts
ISI (ms)
CNI Latency
Reaction Time
Movement Time
Probability
Trial Value
Firing Rate (Hz)
Time (min)
Light-On
Center-In
Go Cue
Center-Out
Side-In
Side-Out
Food-Port-In
Unengaged
Engaged
Ipsi
Contra
Unrewarded
Rewarded
Trial
Time (s)
Time (ms)
Unpredicted
Reward
Cue onset
75%
25%
0%

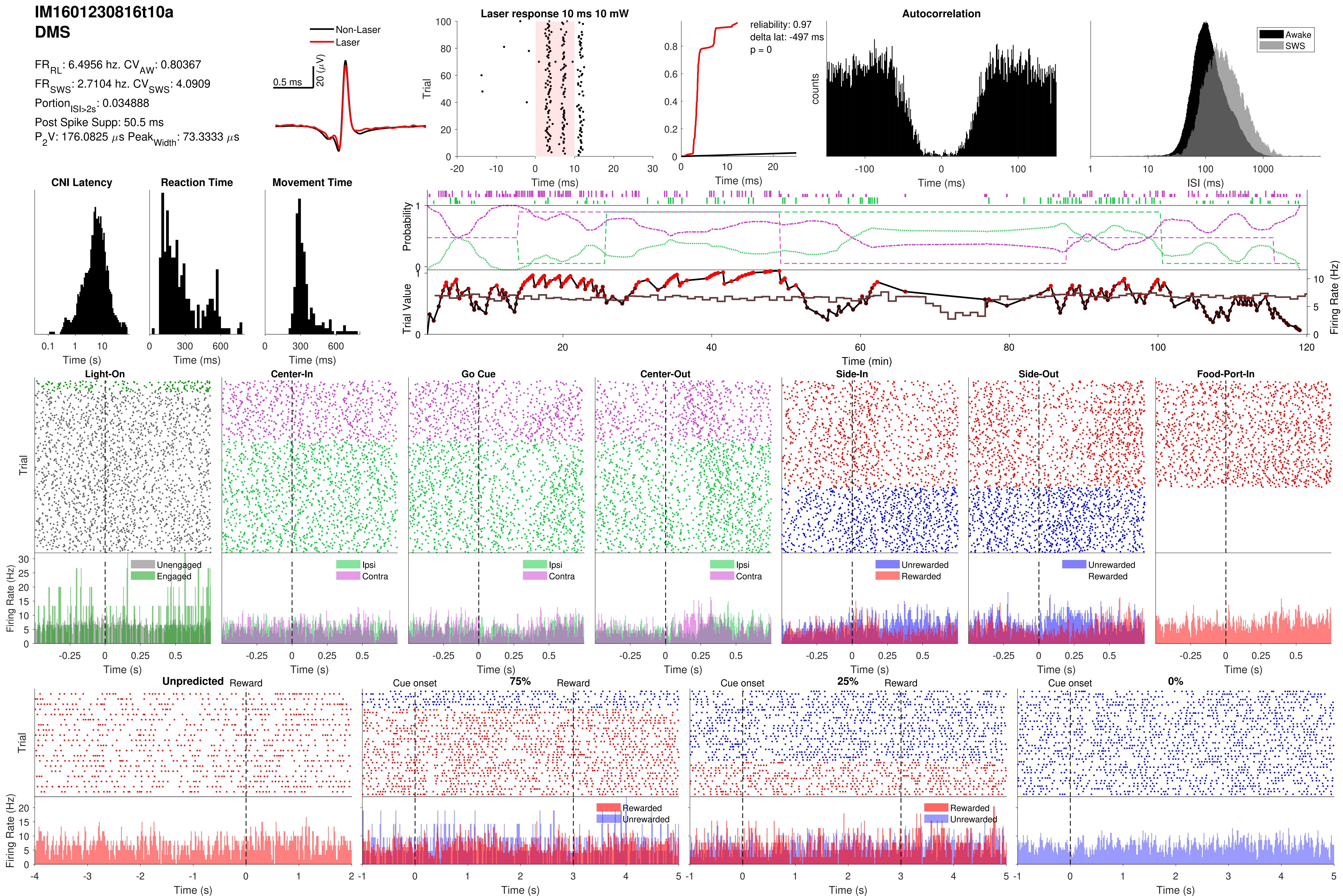

IM1601230816t10a
DMS
FR_RL: 6.4956 hz. CV_AW: 0.80367
FR_SWS: 2.7104 hz. CV_SWS: 4.0909
Portion_ISI>2s: 0.034888
Post Spike Supp: 50.5 ms
P2V: 176.0825 μs Peak_Width: 73.3333 μs
Non-Laser
Laser
0.5 ms
20 (μV)
Laser response 10 ms 10 mW
Trial
Time (ms)
reliability: 0.97
delta lat: -497 ms
p = 0
Autocorrelation
counts
Awake
SWS
ISI (ms)
CNI Latency
Reaction Time
Movement Time
Time (s)
Time (ms)
Probability
Trial Value
Firing Rate (Hz)
Time (min)
Light-On
Center-In
Go Cue
Center-Out
Side-In
Side-Out
Food-Port-In
Unengaged
Engaged
Ipsi
Contra
Unrewarded
Rewarded
Unpredicted
Reward
Cue onset
75%
25%
0%

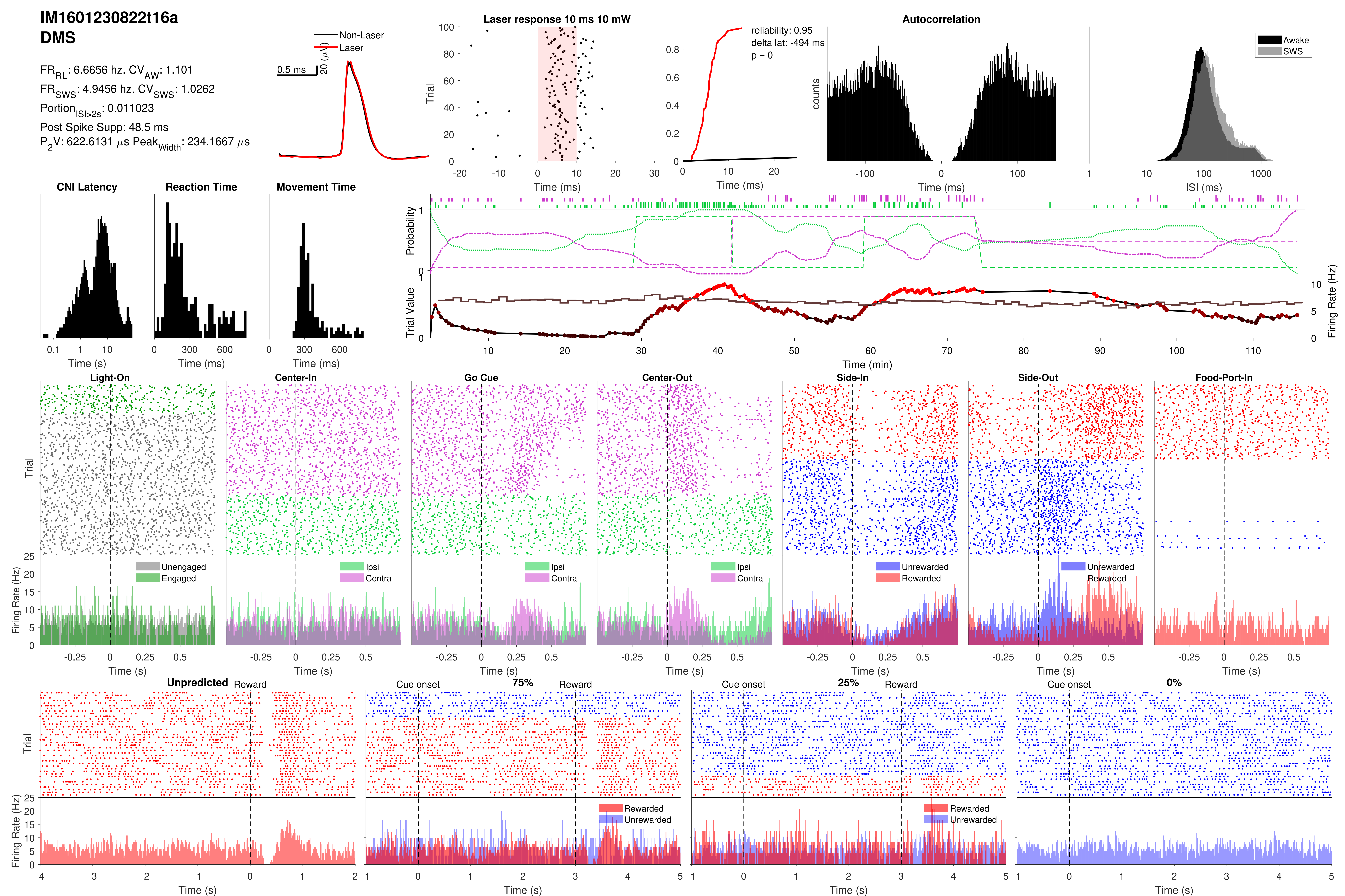

IM1601230822t16a
DMS
FR_RL: 6.6656 hz. CV_AW: 1.101
FR_SWS: 4.9456 hz. CV_SWS: 1.0262
Portion_ISI>2s: 0.011023
Post Spike Supp: 48.5 ms
P_2V: 622.6131 µs Peak_Width: 234.1667 µs
Non-Laser
Laser
0.5 ms
20 (µV)
Laser response 10 ms 10 mW
Trial
Time (ms)
reliability: 0.95
delta lat: -494 ms
p = 0
Autocorrelation
counts
Awake
SWS
ISI (ms)
CNI Latency
Reaction Time
Movement Time
Time (s)
Time (ms)
Probability
Trial Value
Firing Rate (Hz)
Time (min)
Light-On
Center-In
Go Cue
Center-Out
Side-In
Side-Out
Food-Port-In
Unengaged
Engaged
Ipsi
Contra
Unrewarded
Rewarded
Unpredicted
Reward
Cue onset
75%
25%
0%

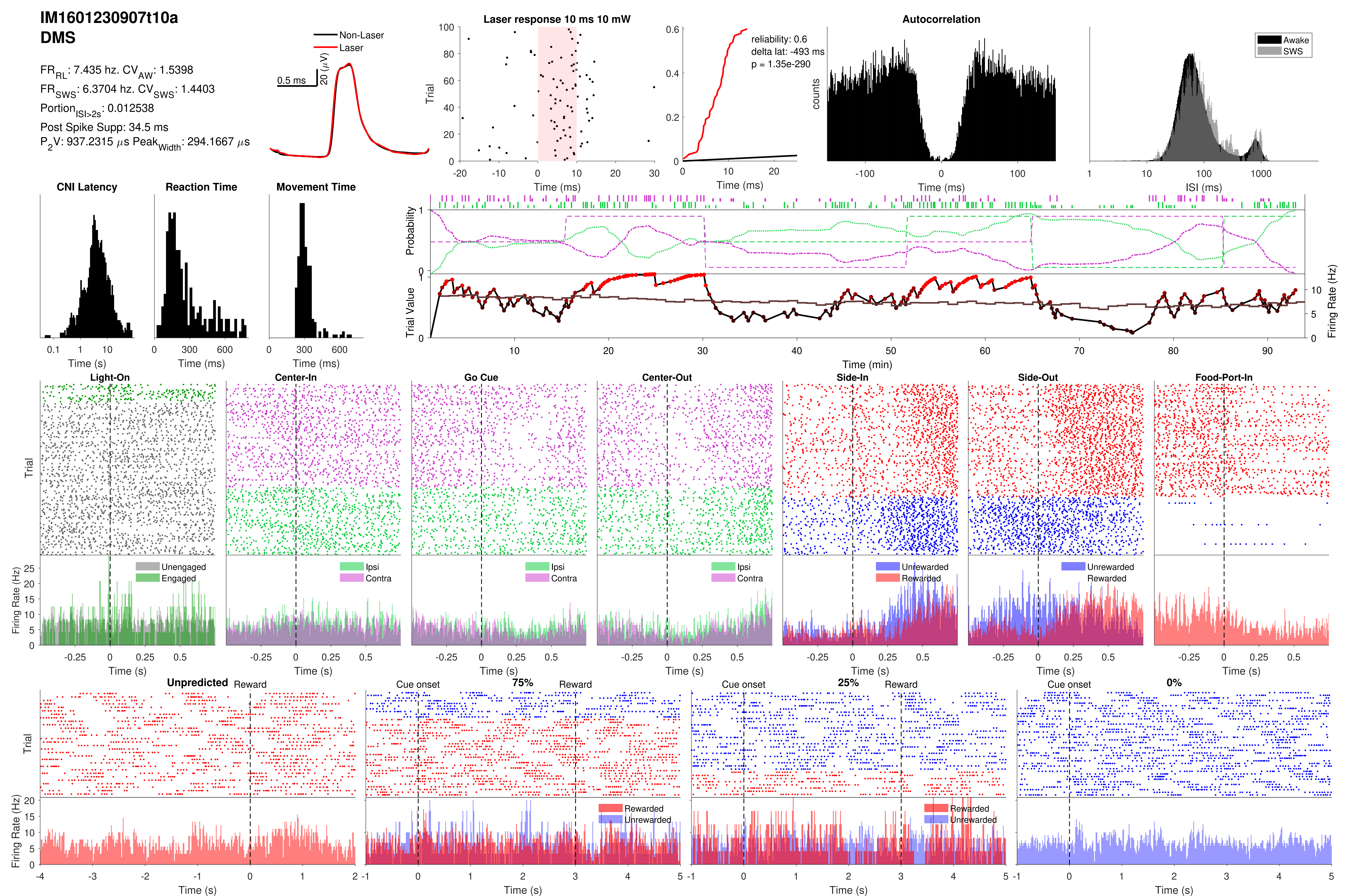
IM1601230907t10a
DMS
FR_RL: 7.435 hz. CV_AW: 1.5398
FR_SWS: 6.3704 hz. CV_SWS: 1.4403
Portion_ISI>2s: 0.012538
Post Spike Supp: 34.5 ms
P_2V: 937.2315 μs Peak_Width: 294.1667 μs
Non-Laser
Laser
0.5 ms
20 (μV)
Laser response 10 ms 10 mW
Trial
Time (ms)
reliability: 0.6
delta lat: -493 ms
p = 1.35e-290
Time (ms)
Autocorrelation
counts
Time (ms)
Awake
SWS
ISI (ms)
CNI Latency
Time (s)
Reaction Time
Time (ms)
Movement Time
Time (ms)
Probability
Trial Value
Firing Rate (Hz)
Time (min)
Light-On
Center-In
Go Cue
Center-Out
Side-In
Side-Out
Food-Port-In
Trial
Firing Rate (Hz)
Unengaged
Engaged
Ipsi
Contra
Unrewarded
Rewarded
Time (s)
Unpredicted
Reward
Cue onset
75%
25%
0%
Rewarded
Unrewarded
Time (s)

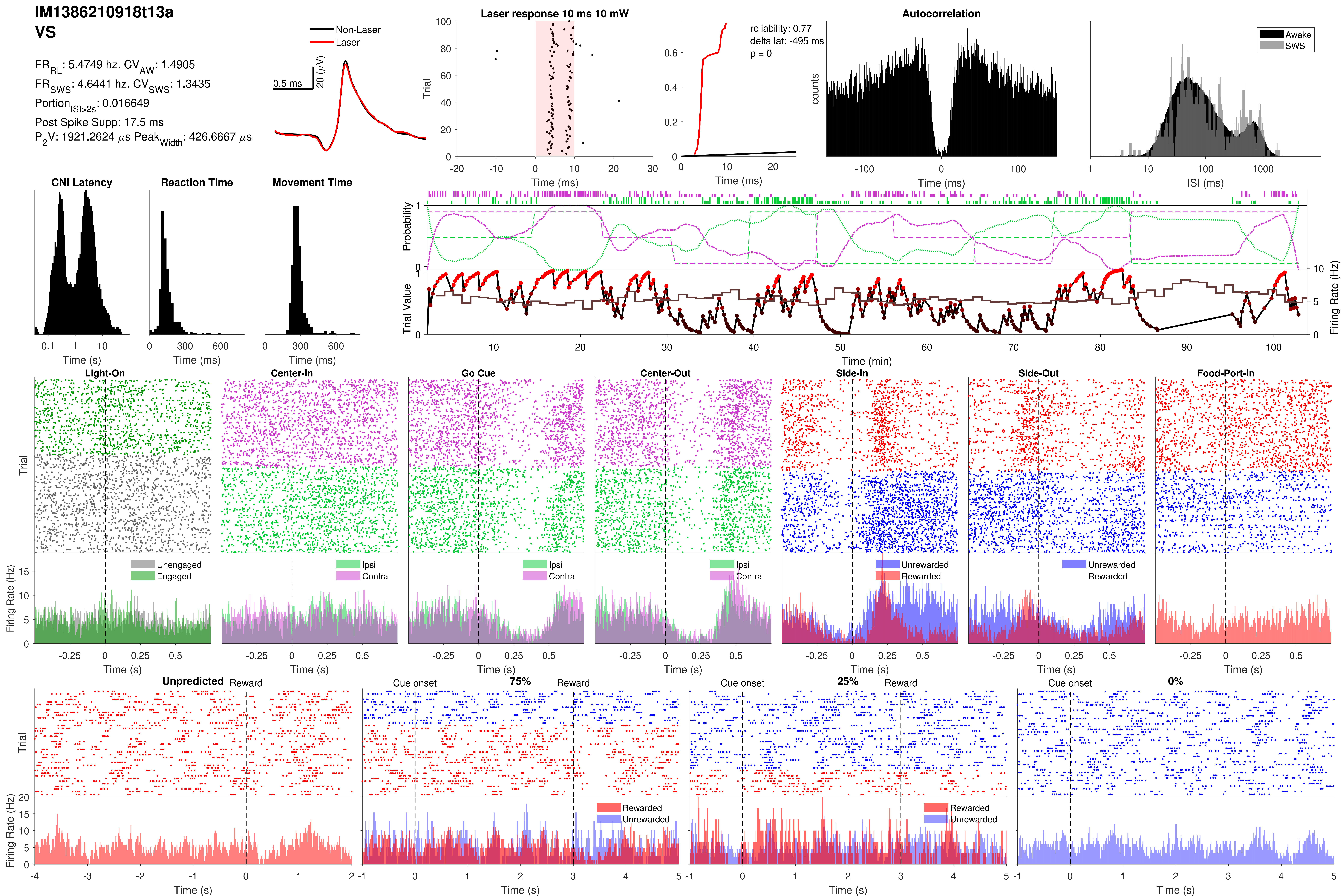

IM1386210918t13a
VS
FR_RL: 5.4749 hz. CV_AW: 1.4905
FR_SWS: 4.6441 hz. CV_SWS: 1.3435
Portion_ISI>2s: 0.016649
Post Spike Supp: 17.5 ms
P_2V: 1921.2624 μs Peak_Width: 426.6667 μs
Non-Laser
Laser
0.5 ms
20 (μV)
Laser response 10 ms 10 mW
Trial
Time (ms)
reliability: 0.77
delta lat: -495 ms
p = 0
Time (ms)
Autocorrelation
counts
Time (ms)
Awake
SWS
ISI (ms)
CNI Latency
Time (s)
Reaction Time
Time (ms)
Movement Time
Time (ms)
Probability
Trial Value
Firing Rate (Hz)
Time (min)
Light-On
Center-In
Go Cue
Center-Out
Side-In
Side-Out
Food-Port-In
Trial
Firing Rate (Hz)
Unengaged
Engaged
Ipsi
Contra
Unrewarded
Rewarded
Time (s)
Unpredicted
Reward
Cue onset
75%
25%
0%
Rewarded
Unrewarded
Trial
Firing Rate (Hz)
Time (s)

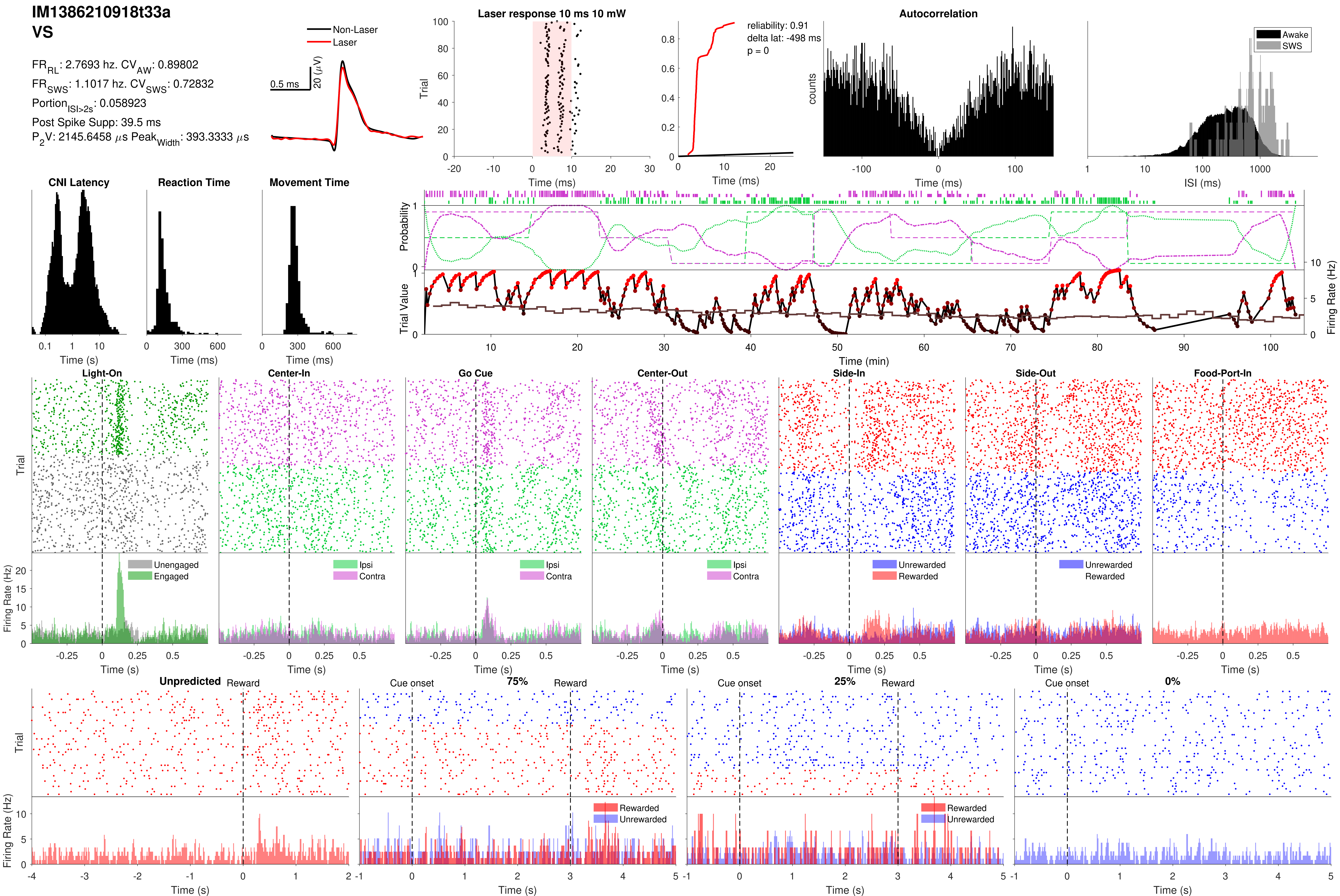
IM1386210918t33a
VS
Non-Laser
Laser
FRRL: 2.7693 hz. CVAW: 0.89802
FRSWS: 1.1017 hz. CVSWS: 0.72832
PortionISI>2s: 0.058923
Post Spike Supp: 39.5 ms
P2V: 2145.6458 µs PeakWidth: 393.3333 µs
0.5 ms
20 (µV)
Laser response 10 ms 10 mW
Trial
Time (ms)
reliability: 0.91
delta lat: -498 ms
p = 0
Time (ms)
Autocorrelation
counts
Time (ms)
Awake
SWS
ISI (ms)
CNI Latency
Time (s)
Reaction Time
Time (ms)
Movement Time
Time (ms)
Probability
Trial Value
Firing Rate (Hz)
Time (min)
Light-On
Center-In
Go Cue
Center-Out
Side-In
Side-Out
Food-Port-In
Trial
Firing Rate (Hz)
Unengaged
Engaged
Ipsi
Contra
Unrewarded
Rewarded
Time (s)
Unpredicted
Reward
Cue onset
75%
25%
0%
Trial
Firing Rate (Hz)
Rewarded
Unrewarded
Time (s)

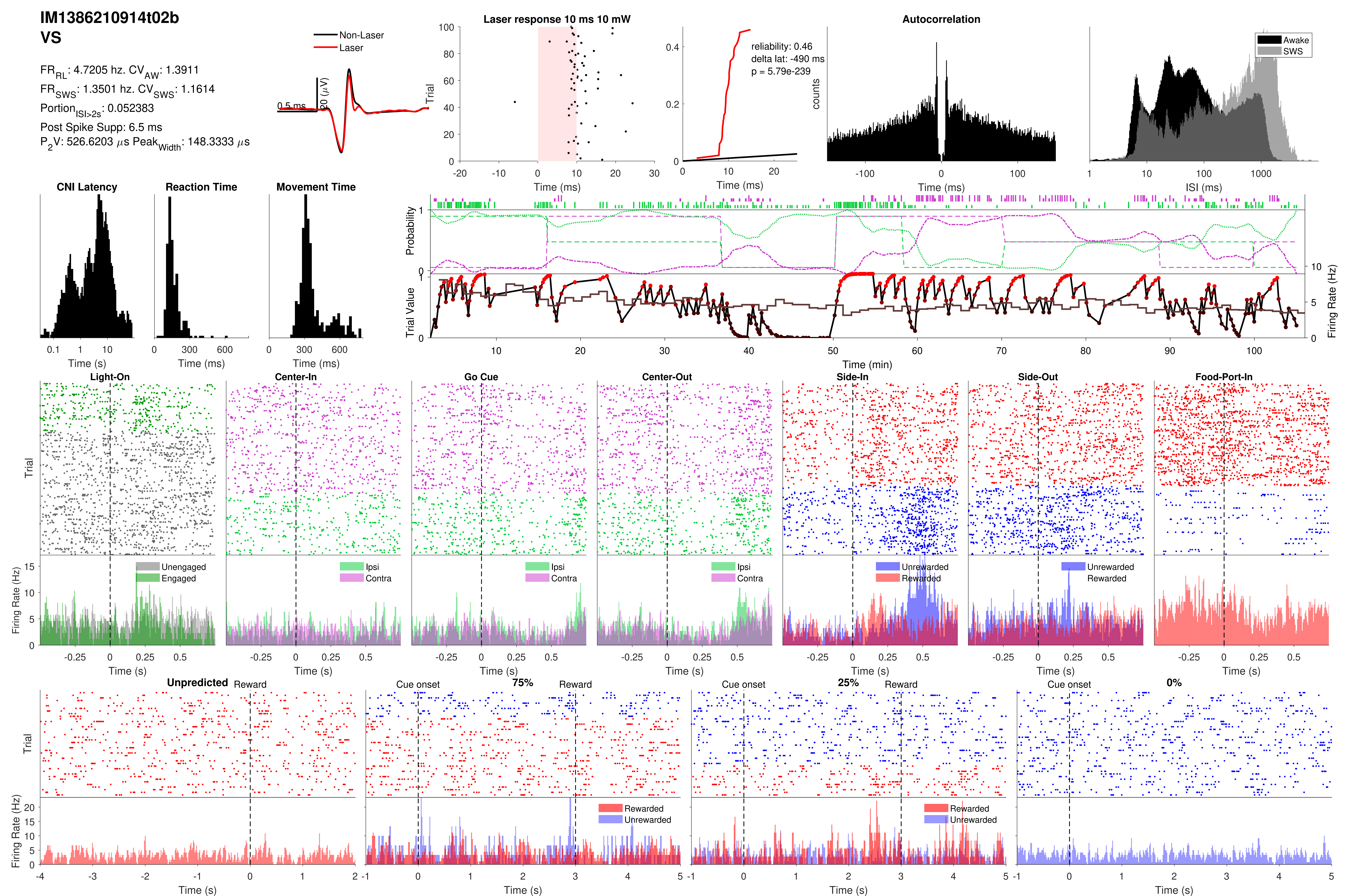

IM1386210914t02b
VS
FR_RL: 4.7205 hz. CV_AW: 1.3911
FR_SWS: 1.3501 hz. CV_SWS: 1.1614
Portion_ISI>2s: 0.052383
Post Spike Supp: 6.5 ms
P_2V: 526.6203 µs Peak_Width: 148.3333 µs
Non-Laser
Laser
0.5 ms
20 (µV)
Laser response 10 ms 10 mW
Trial
Time (ms)
reliability: 0.46
delta lat: -490 ms
p = 5.79e-239
Time (ms)
Autocorrelation
counts
Time (ms)
Awake
SWS
ISI (ms)
CNI Latency
Time (s)
Reaction Time
Time (ms)
Movement Time
Time (ms)
Probability
Trial Value
Firing Rate (Hz)
Time (min)
Light-On
Center-In
Go Cue
Center-Out
Side-In
Side-Out
Food-Port-In
Trial
Firing Rate (Hz)
Unengaged
Engaged
Ipsi
Contra
Unrewarded
Rewarded
Time (s)
Unpredicted
Reward
Cue onset
75%
25%
0%
Rewarded
Unrewarded
Trial
Firing Rate (Hz)
Time (s)

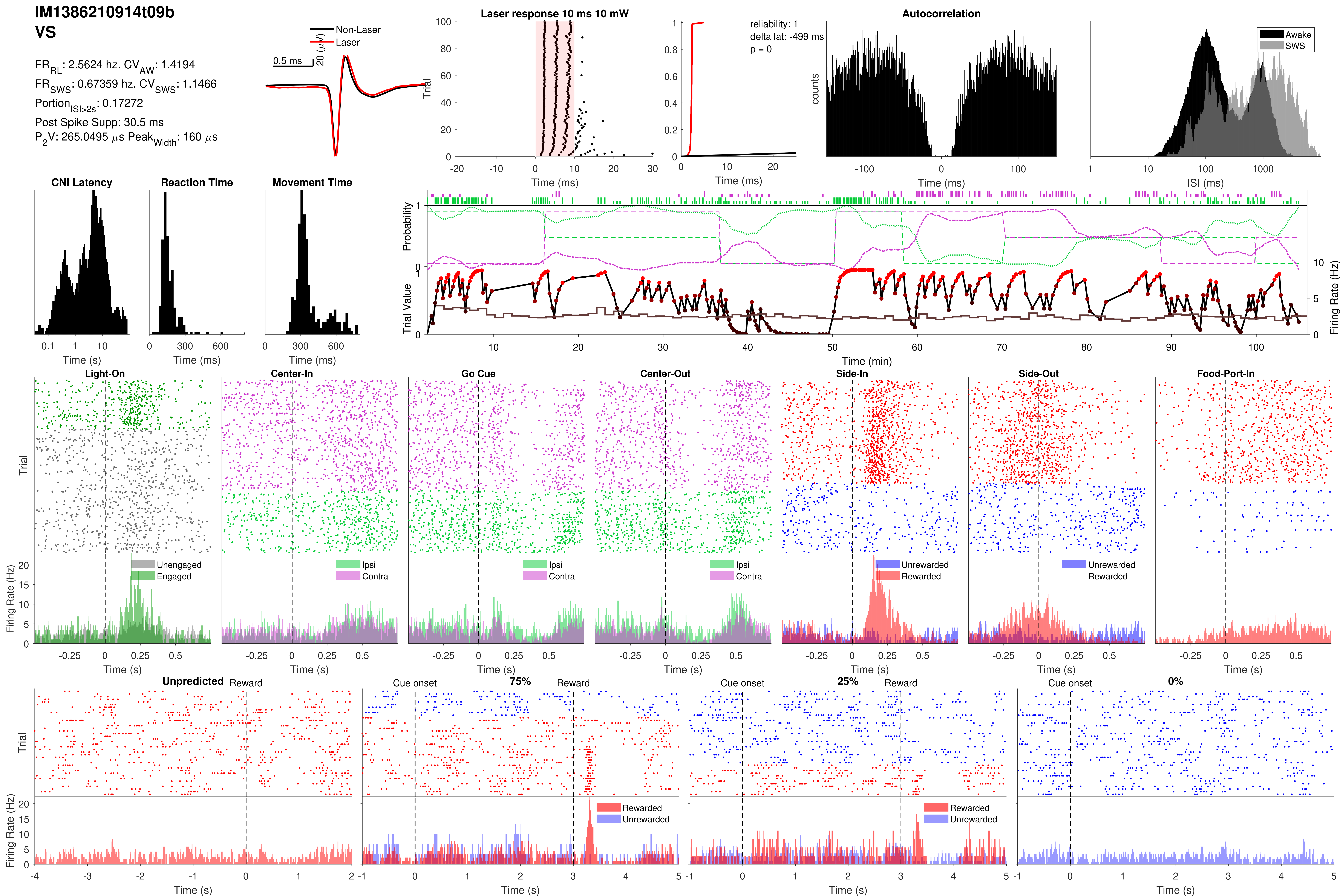
IM1386210914t09b
VS
FR_RL: 2.5624 hz. CV_AW: 1.4194
FR_SWS: 0.67359 hz. CV_SWS: 1.1466
Portion_ISI>2s: 0.17272
Post Spike Supp: 30.5 ms
P2V: 265.0495 μs Peak_Width: 160 μs
Non-Laser
Laser
0.5 ms
20 (μV)
Laser response 10 ms 10 mW
Trial
Time (ms)
reliability: 1
delta lat: -499 ms
p = 0
Time (ms)
Autocorrelation
counts
Time (ms)
Awake
SWS
ISI (ms)
CNI Latency
Time (s)
Reaction Time
Time (ms)
Movement Time
Time (ms)
Probability
Trial Value
Firing Rate (Hz)
Time (min)
Light-On
Center-In
Go Cue
Center-Out
Side-In
Side-Out
Food-Port-In
Trial
Firing Rate (Hz)
Unengaged
Engaged
Ipsi
Contra
Unrewarded
Rewarded
Time (s)
Unpredicted
Reward
Cue onset
75%
25%
0%
Trial
Firing Rate (Hz)
Rewarded
Unrewarded
Time (s)

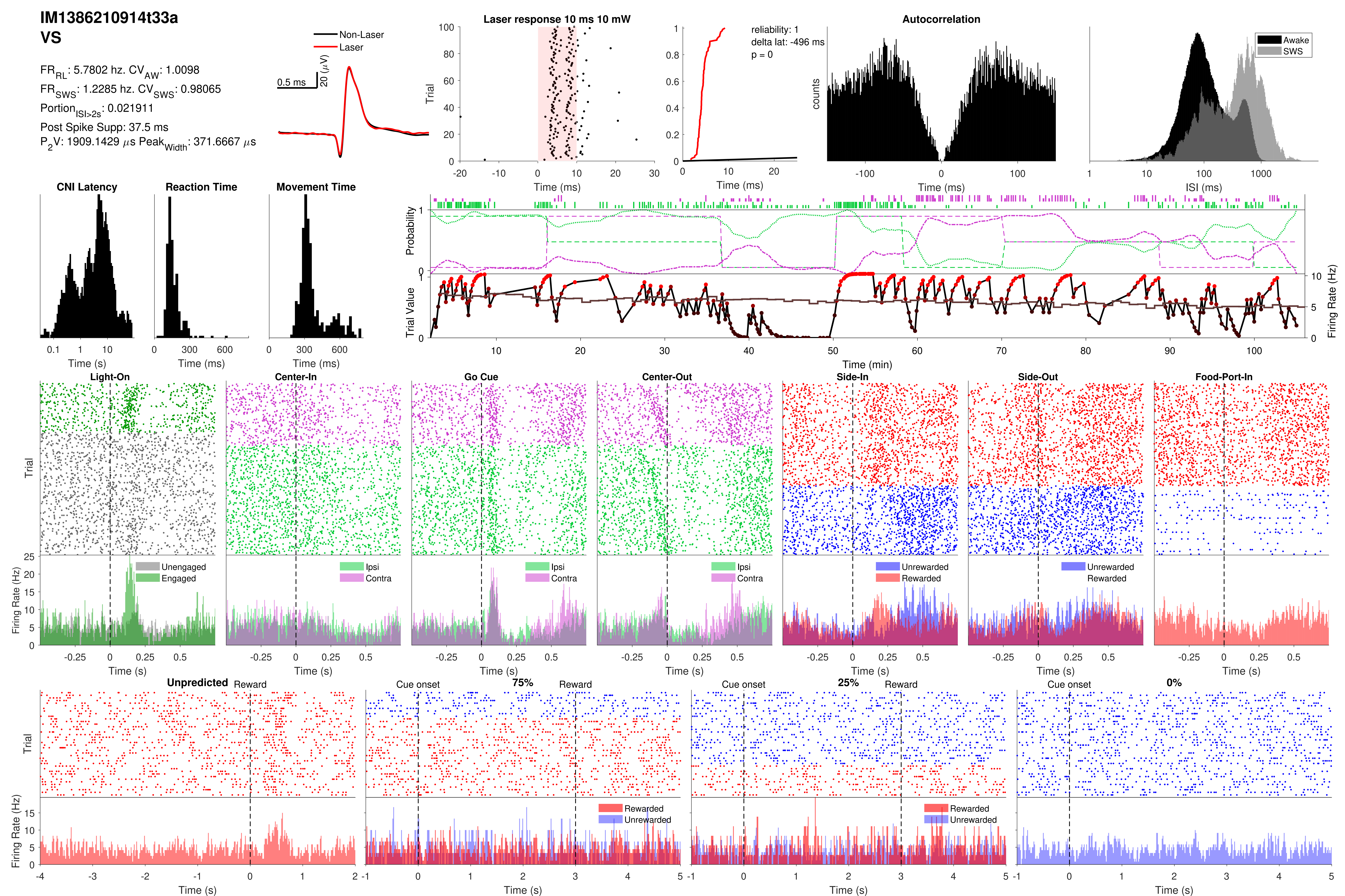
IM1386210914t33a
VS
FR_RL: 5.7802 hz. CV_AW: 1.0098
FR_SWS: 1.2285 hz. CV_SWS: 0.98065
Portion_ISI>2s: 0.021911
Post Spike Supp: 37.5 ms
P_2V: 1909.1429 µs Peak_Width: 371.6667 µs
Non-Laser
Laser
0.5 ms
20 (µV)
Laser response 10 ms 10 mW
Trial
Time (ms)
reliability: 1
delta lat: -496 ms
p = 0
Time (ms)
Autocorrelation
counts
Time (ms)
Awake
SWS
ISI (ms)
CNI Latency
Time (s)
Reaction Time
Time (ms)
Movement Time
Time (ms)
Probability
Trial Value
Firing Rate (Hz)
Time (min)
Light-On
Center-In
Go Cue
Center-Out
Side-In
Side-Out
Food-Port-In
Trial
Firing Rate (Hz)
Unengaged
Engaged
Ipsi
Contra
Unrewarded
Rewarded
Time (s)
Unpredicted
Reward
Cue onset
75%
25%
0%
Trial
Firing Rate (Hz)
Rewarded
Unrewarded
Time (s)

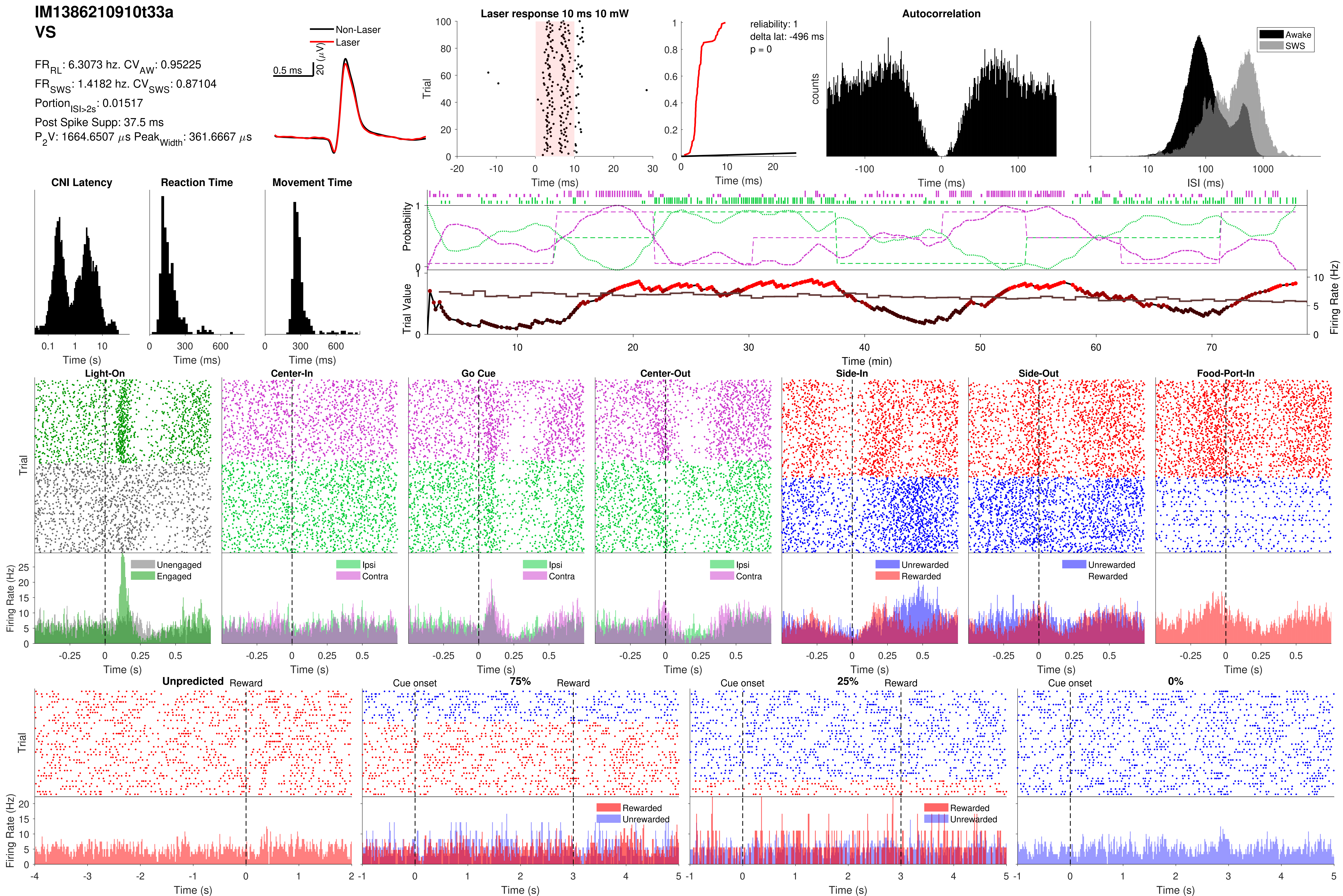

IM1386210910t33a
VS
FR_RL: 6.3073 hz. CV_AW: 0.95225
FR_SWS: 1.4182 hz. CV_SWS: 0.87104
Portion_ISI>2s: 0.01517
Post Spike Supp: 37.5 ms
P_2V: 1664.6507 µs Peak_Width: 361.6667 µs
Non-Laser
Laser
0.5 ms
20 (µV)
Laser response 10 ms 10 mW
Trial
Time (ms)
reliability: 1
delta lat: -496 ms
p = 0
Time (ms)
Autocorrelation
counts
Time (ms)
Awake
SWS
ISI (ms)
CNI Latency
Time (s)
Reaction Time
Time (ms)
Movement Time
Time (ms)
Probability
Trial Value
Firing Rate (Hz)
Time (min)
Light-On
Center-In
Go Cue
Center-Out
Side-In
Side-Out
Food-Port-In
Trial
Unengaged
Engaged
Ipsi
Contra
Ipsi
Contra
Ipsi
Contra
Unrewarded
Rewarded
Unrewarded
Rewarded
Firing Rate (Hz)
Time (s)
Unpredicted
Reward
Cue onset
75%
Reward
Cue onset
25%
Reward
Cue onset
0%
Trial
Rewarded
Unrewarded
Rewarded
Unrewarded
Firing Rate (Hz)
Time (s)

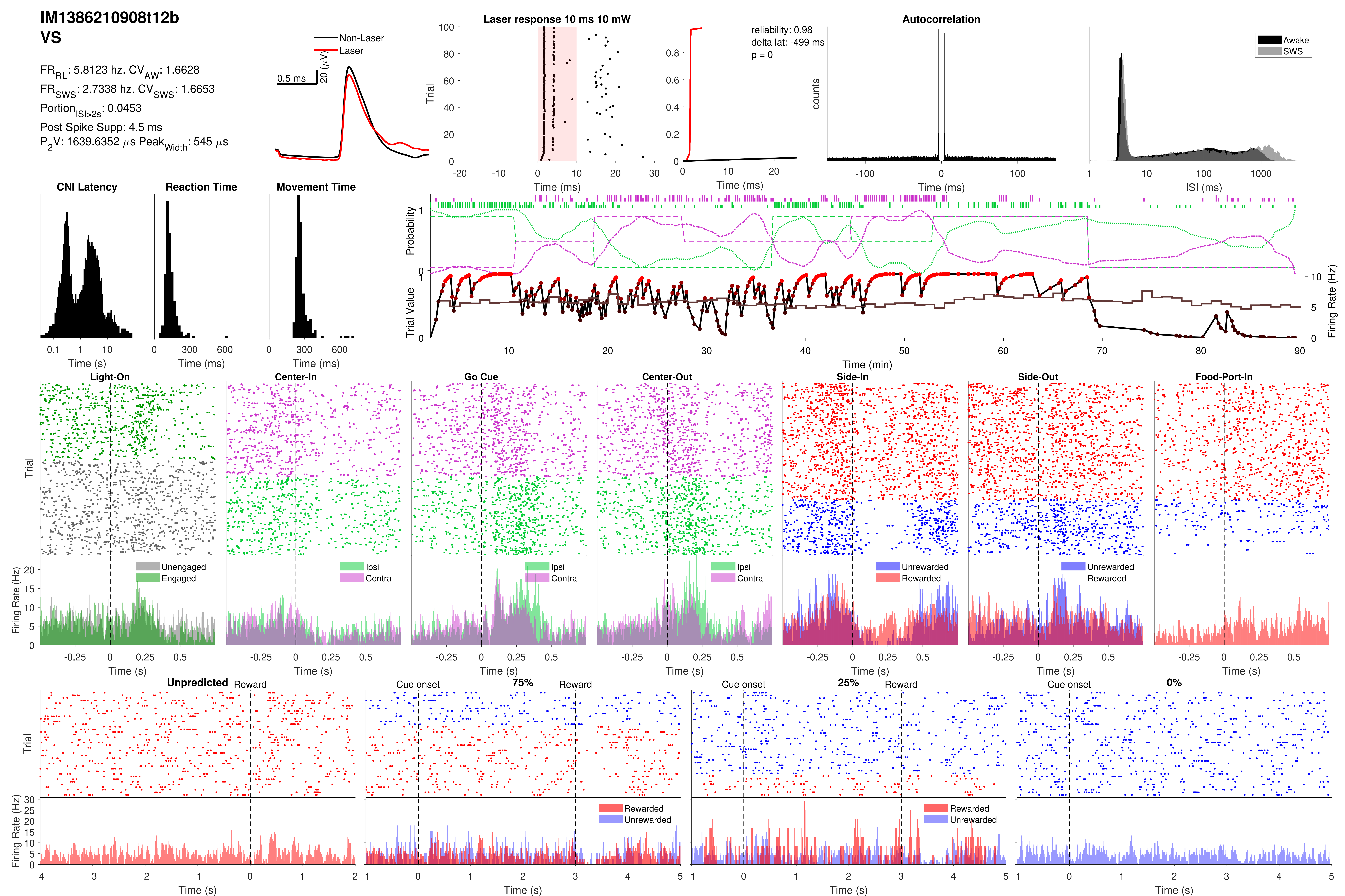

IM1386210908t12b
VS
FR_RL: 5.8123 hz. CV_AW: 1.6628
FR_SWS: 2.7338 hz. CV_SWS: 1.6653
Portion_ISI>2s: 0.0453
Post Spike Supp: 4.5 ms
P_2V: 1639.6352 µs Peak_Width: 545 µs
Non-Laser
Laser
0.5 ms
20 (µV)
Laser response 10 ms 10 mW
Trial
Time (ms)
reliability: 0.98
delta lat: -499 ms
p = 0
Time (ms)
Autocorrelation
counts
Time (ms)
Awake
SWS
ISI (ms)
CNI Latency
Time (s)
Reaction Time
Time (ms)
Movement Time
Time (ms)
Probability
Trial Value
Firing Rate (Hz)
Time (min)
Light-On
Center-In
Go Cue
Center-Out
Side-In
Side-Out
Food-Port-In
Trial
Firing Rate (Hz)
Unengaged
Engaged
Ipsi
Contra
Unrewarded
Rewarded
Time (s)
Unpredicted
Reward
Cue onset
75%
25%
0%
Rewarded
Unrewarded
Trial
Firing Rate (Hz)
Time (s)

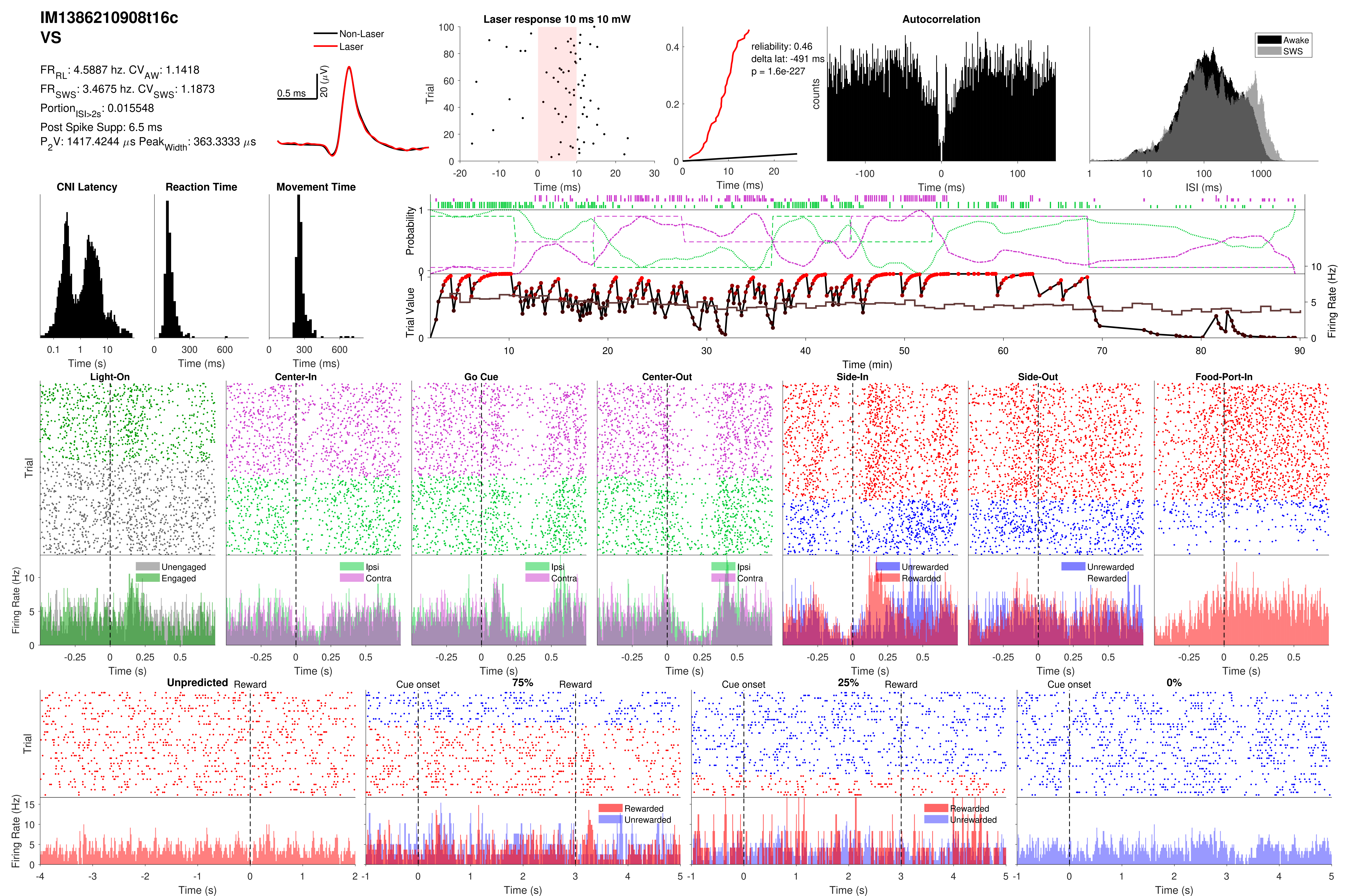

IM1386210908t16c
VS
FR_RL: 4.5887 hz. CV_AW: 1.1418
FR_SWS: 3.4675 hz. CV_SWS: 1.1873
Portion_ISI>2s: 0.015548
Post Spike Supp: 6.5 ms
P_2V: 1417.4244 µs Peak_Width: 363.3333 µs
Non-Laser
Laser
0.5 ms
20 (µV)
Laser response 10 ms 10 mW
Trial
Time (ms)
reliability: 0.46
delta lat: -491 ms
p = 1.6e-227
Autocorrelation
counts
Awake
SWS
ISI (ms)
CNI Latency
Reaction Time
Movement Time
Time (s)
Time (ms)
Probability
Trial Value
Firing Rate (Hz)
Time (min)
Light-On
Center-In
Go Cue
Center-Out
Side-In
Side-Out
Food-Port-In
Unengaged
Engaged
Ipsi
Contra
Unrewarded
Rewarded
Unpredicted
Reward
Cue onset
75%
25%
0%

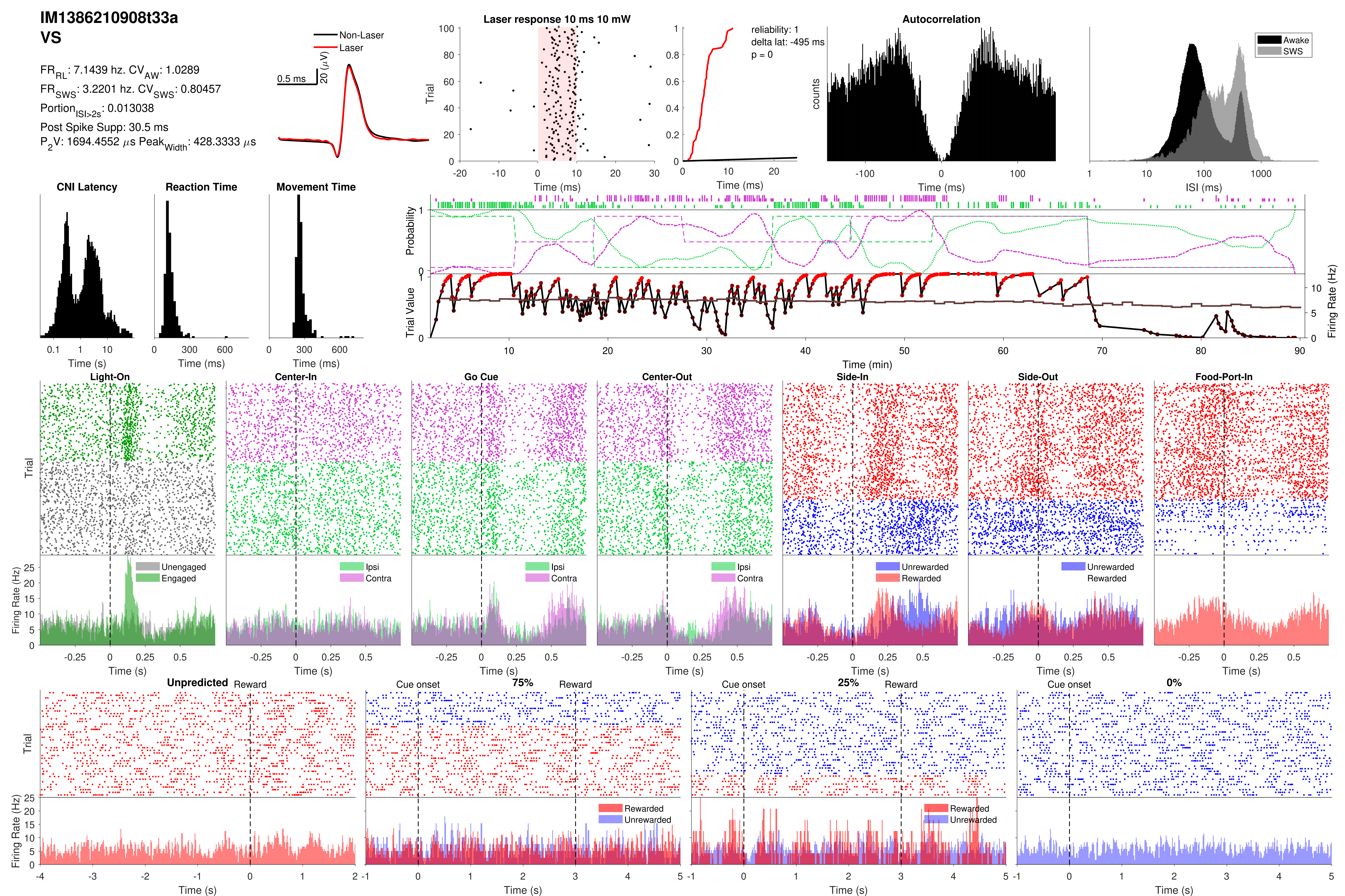
IM1386210908t33a
VS
FR_RL: 7.1439 hz. CV_AW: 1.0289
FR_SWS: 3.2201 hz. CV_SWS: 0.80457
Portion_ISI>2s: 0.013038
Post Spike Supp: 30.5 ms
P_2V: 1694.4552 μs Peak_Width: 428.3333 μs
Non-Laser
Laser
0.5 ms
20 (μV)
Laser response 10 ms 10 mW
Trial
Time (ms)
reliability: 1
delta lat: -495 ms
p = 0
Time (ms)
Autocorrelation
counts
Time (ms)
Awake
SWS
ISI (ms)
CNI Latency
Time (s)
Reaction Time
Time (ms)
Movement Time
Time (ms)
Probability
Trial Value
Firing Rate (Hz)
Time (min)
Light-On
Center-In
Go Cue
Center-Out
Side-In
Side-Out
Food-Port-In
Trial
Firing Rate (Hz)
Unengaged
Engaged
Ipsi
Contra
Unrewarded
Rewarded
Time (s)
Unpredicted
Reward
Cue onset
75%
25%
0%
Rewarded
Unrewarded
Time (s)

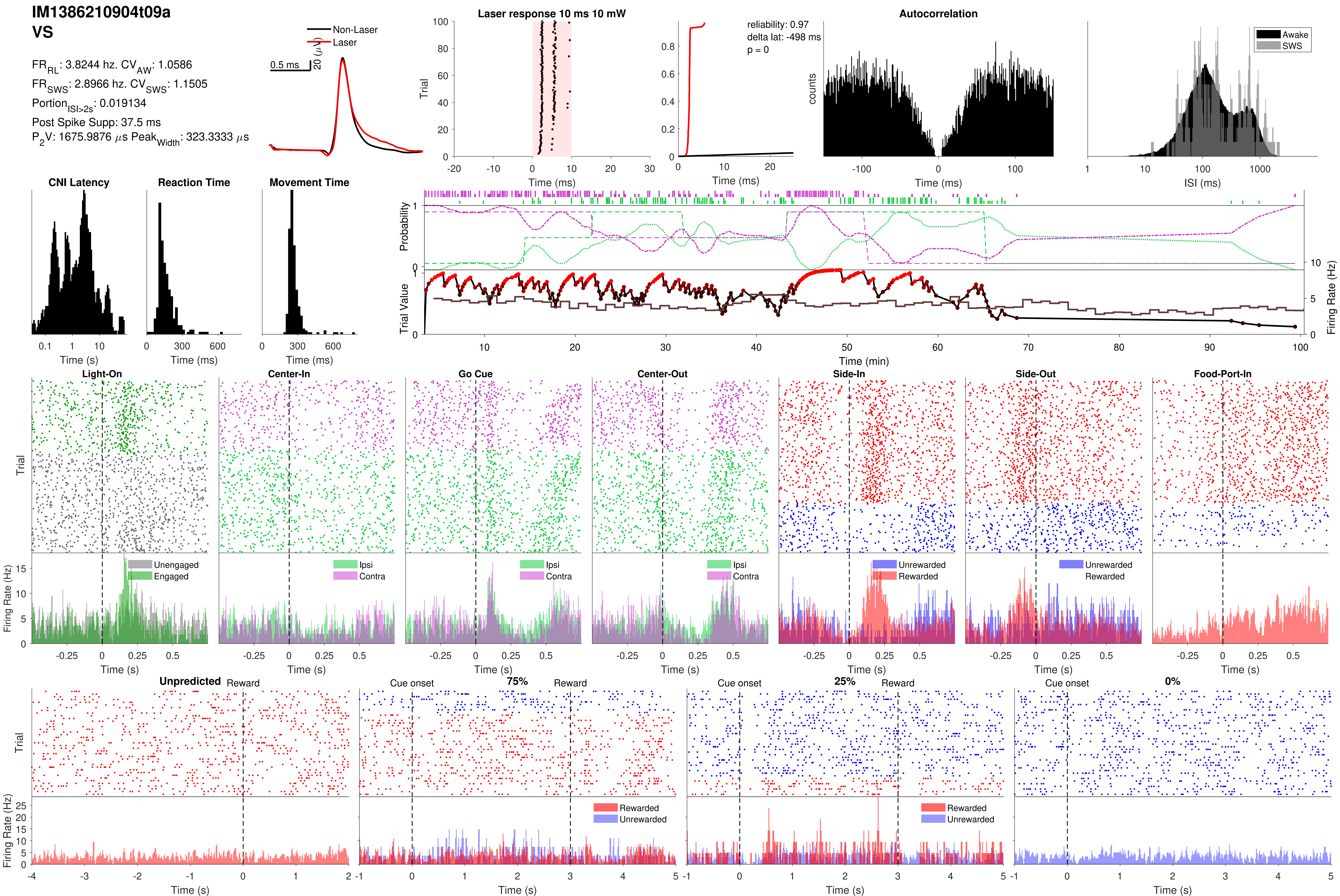
IM1386210904t09a
VS
FR_RL: 3.8244 hz. CV_AW: 1.0586
FR_SWS: 2.8966 hz. CV_SWS: 1.1505
Portion_ISI>2s: 0.019134
Post Spike Supp: 37.5 ms
P_2V: 1675.9876 µs Peak_Width: 323.3333 µs
Non-Laser
Laser
0.5 ms
20 (µV)
Laser response 10 ms 10 mW
Trial
Time (ms)
reliability: 0.97
delta lat: -498 ms
p = 0
Time (ms)
Autocorrelation
counts
Time (ms)
Awake
SWS
ISI (ms)
CNI Latency
Time (s)
Reaction Time
Time (ms)
Movement Time
Time (ms)
Probability
Trial Value
Firing Rate (Hz)
Time (min)
Light-On
Center-In
Go Cue
Center-Out
Side-In
Side-Out
Food-Port-In
Trial
Firing Rate (Hz)
Unengaged
Engaged
Ipsi
Contra
Unrewarded
Rewarded
Time (s)
Unpredicted
Reward
Cue onset
75%
25%
0%
Trial
Firing Rate (Hz)
Rewarded
Unrewarded
Time (s)

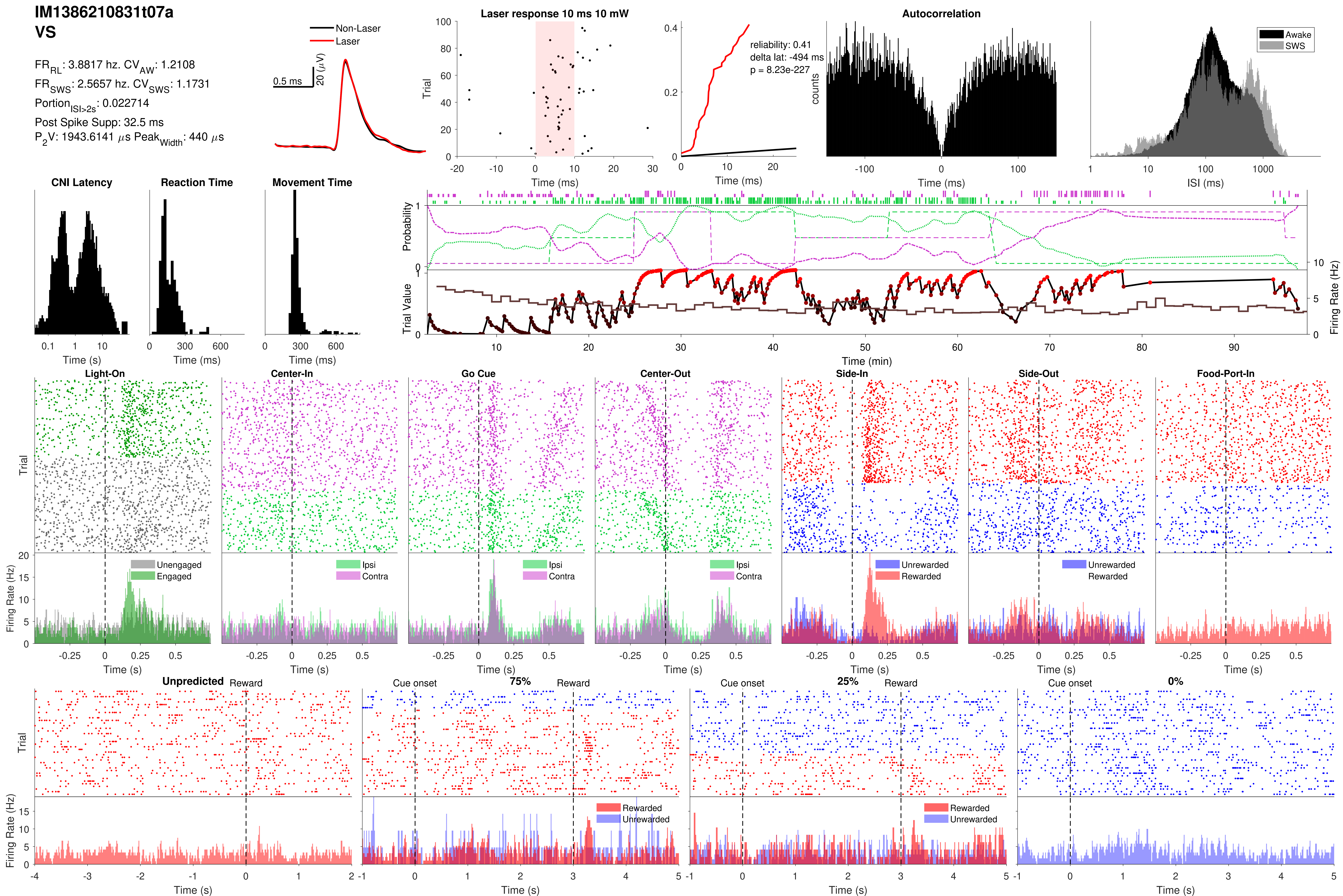
IM1386210831t07a
VS
FR_RL: 3.8817 hz. CV_AW: 1.2108
FR_SWS: 2.5657 hz. CV_SWS: 1.1731
Portion_ISI>2s: 0.022714
Post Spike Supp: 32.5 ms
P_2V: 1943.6141 µs Peak_Width: 440 µs
Non-Laser
Laser
0.5 ms
20 (µV)
Laser response 10 ms 10 mW
Trial
Time (ms)
reliability: 0.41
delta lat: -494 ms
p = 8.23e-227
Time (ms)
Autocorrelation
counts
Time (ms)
Awake
SWS
ISI (ms)
CNI Latency
Time (s)
Reaction Time
Time (ms)
Movement Time
Time (ms)
Probability
Trial Value
Firing Rate (Hz)
Time (min)
Light-On
Center-In
Go Cue
Center-Out
Side-In
Side-Out
Food-Port-In
Trial
Firing Rate (Hz)
Unengaged
Engaged
Ipsi
Contra
Unrewarded
Rewarded
Time (s)
Unpredicted
Reward
Cue onset
75%
25%
0%
Rewarded
Unrewarded
Trial
Firing Rate (Hz)
Time (s)

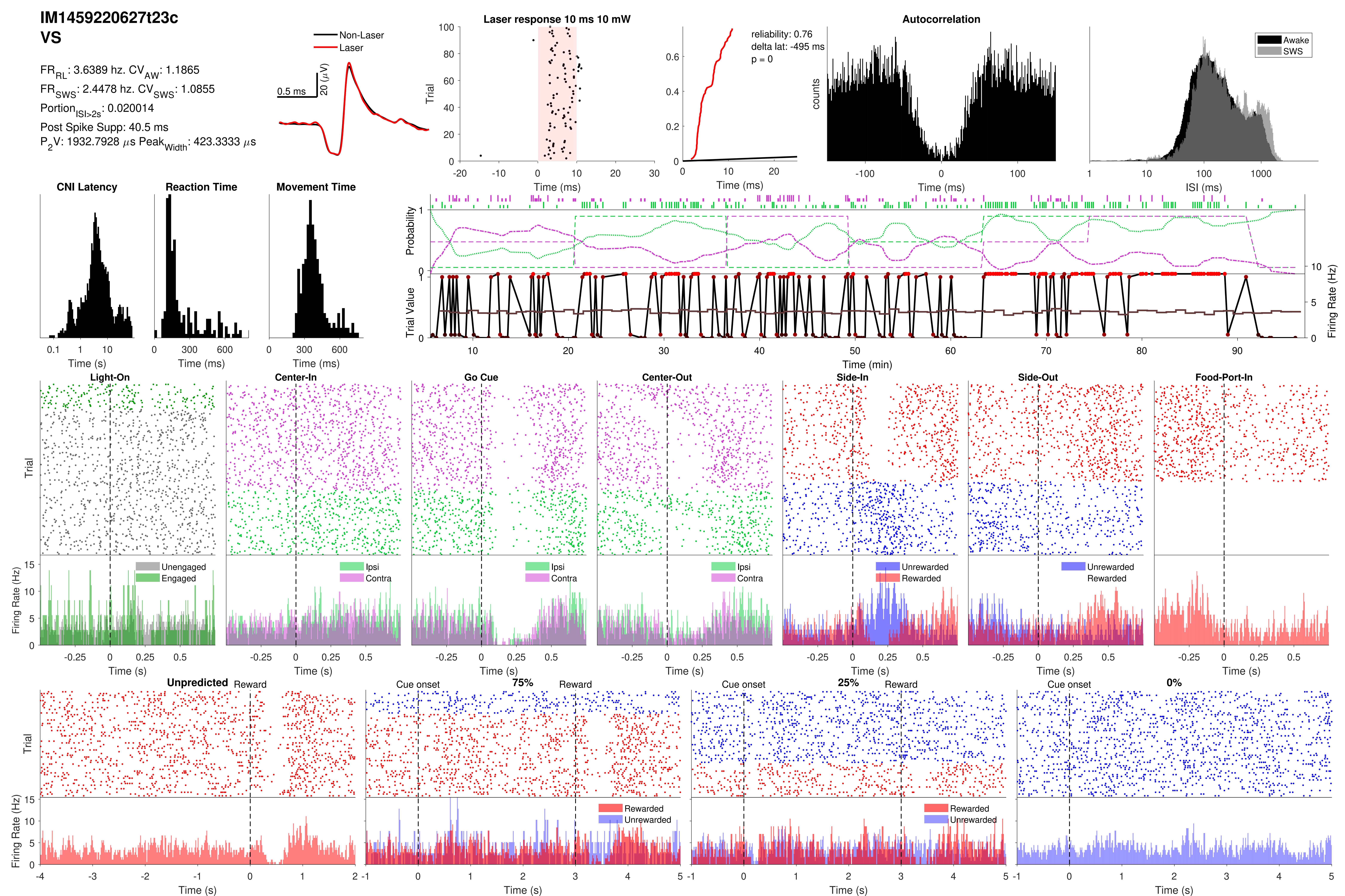

IM1459220627t23c
VS
FR_RL: 3.6389 hz. CV_AW: 1.1865
FR_SWS: 2.4478 hz. CV_SWS: 1.0855
Portion_ISI>2s: 0.020014
Post Spike Supp: 40.5 ms
P_2V: 1932.7928 µs Peak_Width: 423.3333 µs
Non-Laser
Laser
0.5 ms
20 (µV)
Laser response 10 ms 10 mW
Trial
Time (ms)
reliability: 0.76
delta lat: -495 ms
p = 0
Autocorrelation
counts
Awake
SWS
ISI (ms)
CNI Latency
Reaction Time
Movement Time
Time (s)
Probability
Trial Value
Firing Rate (Hz)
Time (min)
Light-On
Center-In
Go Cue
Center-Out
Side-In
Side-Out
Food-Port-In
Unengaged
Engaged
Ipsi
Contra
Unwarded
Rewarded
Unpredicted
Reward
Cue onset
75%
25%
0%
Rewarded
Unrewarded

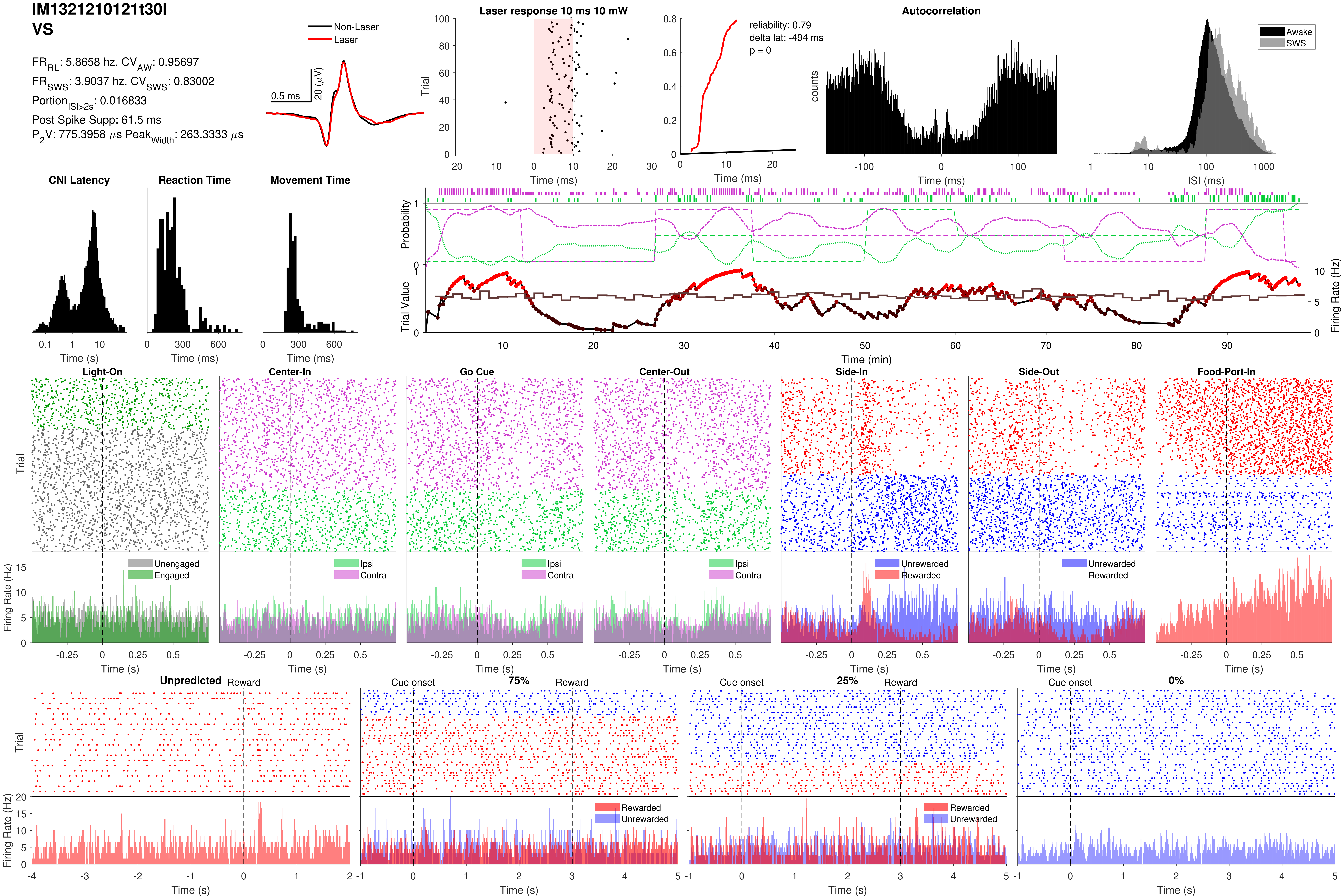
IM1321210121t30l
VS
FR_RL: 5.8658 hz. CV_AW: 0.95697
FR_SWS: 3.9037 hz. CV_SWS: 0.83002
Portion_ISI>2s: 0.016833
Post Spike Supp: 61.5 ms
P_2V: 775.3958 μs Peak_Width: 263.3333 μs
Non-Laser
Laser
0.5 ms
20 (μV)
Laser response 10 ms 10 mW
Trial
Time (ms)
reliability: 0.79
delta lat: -494 ms
p = 0
Time (ms)
Autocorrelation
counts
Time (ms)
Awake
SWS
ISI (ms)
CNI Latency
Time (s)
Reaction Time
Time (ms)
Movement Time
Time (ms)
Probability
Trial Value
Firing Rate (Hz)
Time (min)
Light-On
Center-In
Go Cue
Center-Out
Side-In
Side-Out
Food-Port-In
Trial
Firing Rate (Hz)
Unengaged
Engaged
Ipsi
Contra
Unrewarded
Rewarded
Time (s)
Unpredicted
Reward
Cue onset
75%
25%
0%
Rewarded
Unrewarded
Time (s)

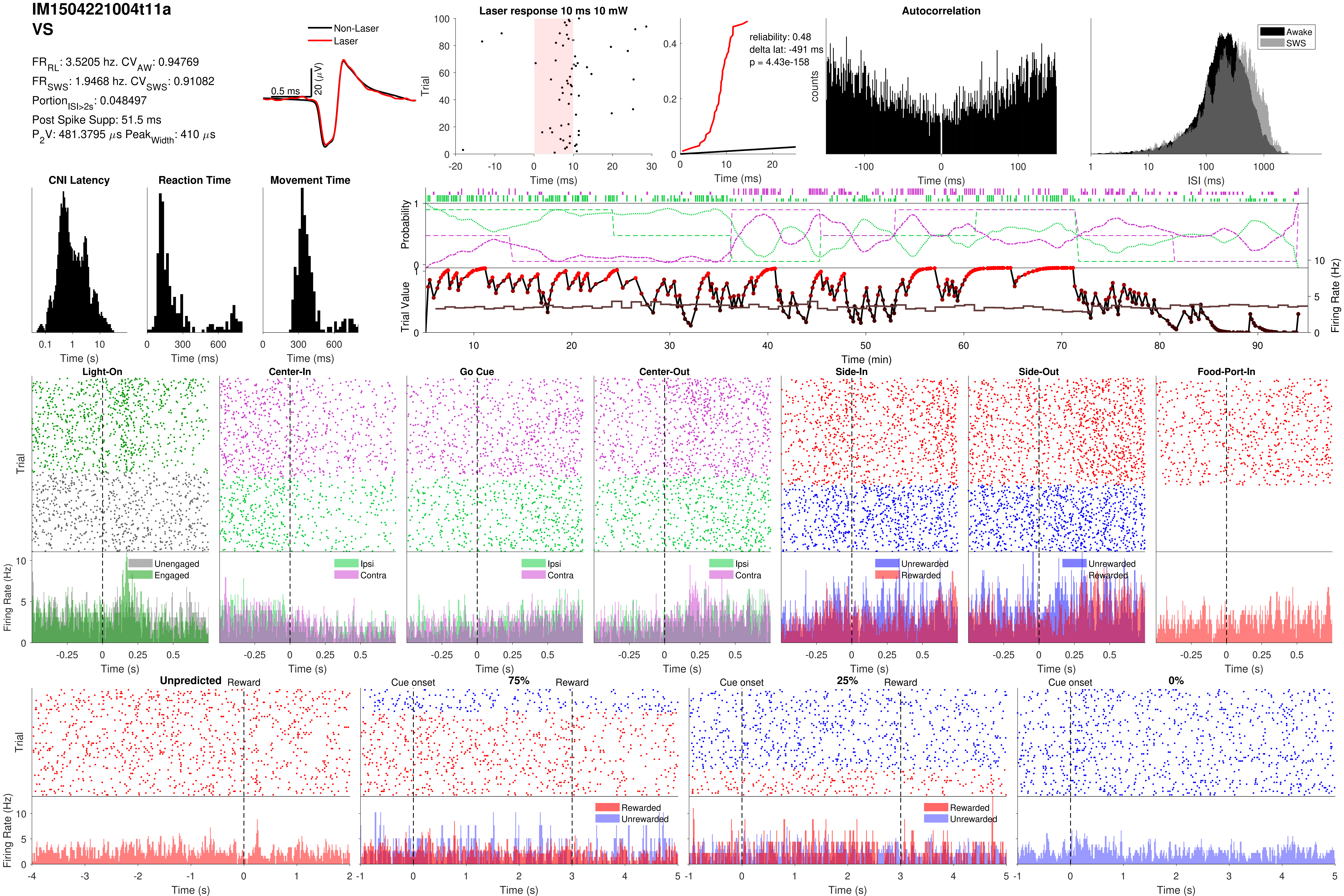
IM1504221004t11a
VS
FR_RL: 3.5205 hz. CV_AW: 0.94769
FR_SWS: 1.9468 hz. CV_SWS: 0.91082
Portion_ISI>2s: 0.048497
Post Spike Supp: 51.5 ms
P_2V: 481.3795 μs Peak_Width: 410 μs
Non-Laser
Laser
0.5 ms
20 (μV)
Laser response 10 ms 10 mW
reliability: 0.48
delta lat: -491 ms
p = 4.43e-158
Autocorrelation
Awake
SWS
Trial
Time (ms)
counts
ISI (ms)
CNI Latency
Reaction Time
Movement Time
Time (s)
Probability
Trial Value
Firing Rate (Hz)
Time (min)
Light-On
Center-In
Go Cue
Center-Out
Side-In
Side-Out
Food-Port-In
Unengaged
Engaged
Ipsi
Contra
Unrewarded
Rewarded
Unpredicted
Reward
Cue onset
75%
25%
0%

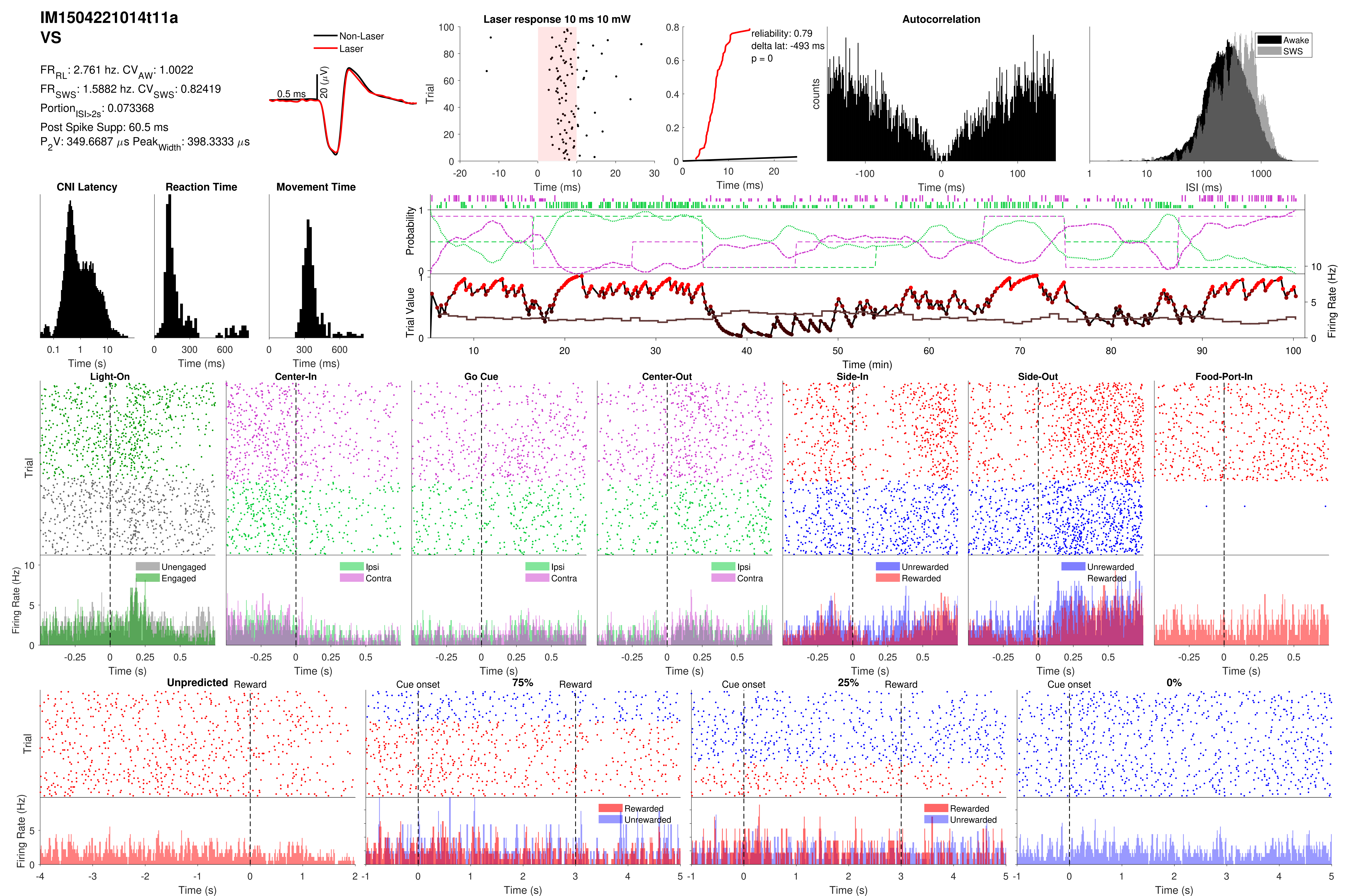
IM1504221014t11a
VS
FR_RL: 2.761 hz. CV_AW: 1.0022
FR_SWS: 1.5882 hz. CV_SWS: 0.82419
Portion_ISI>2s: 0.073368
Post Spike Supp: 60.5 ms
P_2V: 349.6687 μs Peak_Width: 398.3333 μs
Non-Laser
Laser
0.5 ms
20 (μV)
Laser response 10 ms 10 mW
Trial
Time (ms)
reliability: 0.79
delta lat: -493 ms
p = 0
Time (ms)
Autocorrelation
counts
Time (ms)
Awake
SWS
ISI (ms)
CNI Latency
Time (s)
Reaction Time
Time (ms)
Movement Time
Time (ms)
Probability
Trial Value
Firing Rate (Hz)
Time (min)
Light-On
Center-In
Go Cue
Center-Out
Side-In
Side-Out
Food-Port-In
Trial
Unengaged
Engaged
Ipsi
Contra
Ipsi
Contra
Ipsi
Contra
Unrewarded
Rewarded
Unrewarded
Rewarded
Firing Rate (Hz)
Time (s)
Unpredicted
Reward
Cue onset
75%
Reward
Cue onset
25%
Reward
Cue onset
0%
Trial
Rewarded
Unrewarded
Rewarded
Unrewarded
Firing Rate (Hz)
Time (s)

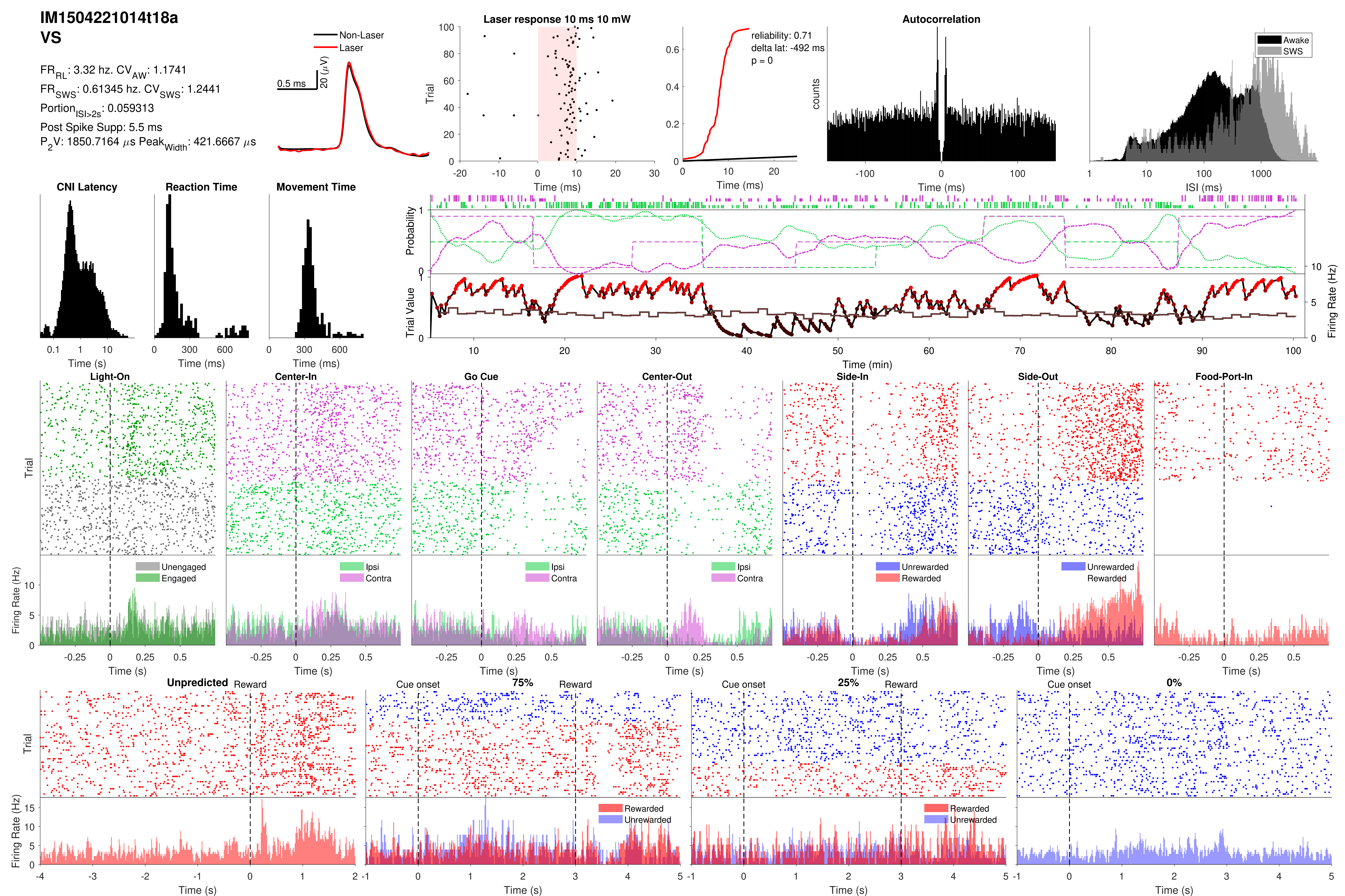
IM1504221014t18a
VS
FR_RL: 3.32 hz. CV_AW: 1.1741
FR_SWS: 0.61345 hz. CV_SWS: 1.2441
Portion_ISI>2s: 0.059313
Post Spike Supp: 5.5 ms
P_2V: 1850.7164 µs Peak_Width: 421.6667 µs
Non-Laser
Laser
0.5 ms
20 (µV)
Laser response 10 ms 10 mW
Trial
Time (ms)
reliability: 0.71
delta lat: -492 ms
p = 0
Time (ms)
Autocorrelation
counts
Time (ms)
Awake
SWS
ISI (ms)
CNI Latency
Time (s)
Reaction Time
Time (ms)
Movement Time
Time (ms)
Probability
Trial Value
Firing Rate (Hz)
Time (min)
Light-On
Center-In
Go Cue
Center-Out
Side-In
Side-Out
Food-Port-In
Trial
Unengaged
Engaged
Ipsi
Contra
Unrewarded
Rewarded
Firing Rate (Hz)
Time (s)
Unpredicted
Reward
Cue onset
75%
25%
0%
Trial
Rewarded
Unrewarded
Firing Rate (Hz)
Time (s)

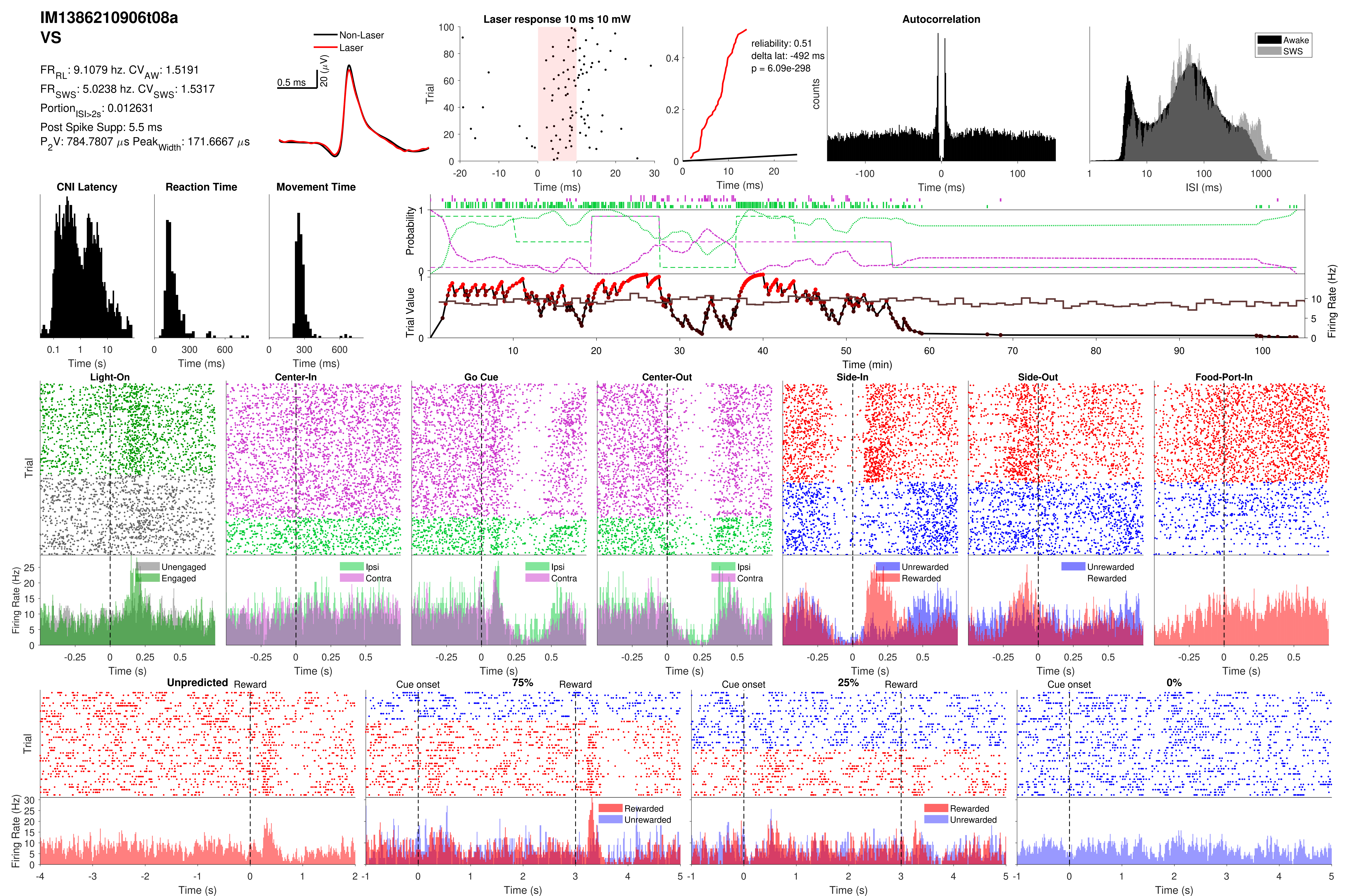

IM1386210906t08a
VS
FR_RL: 9.1079 hz. CV_AW: 1.5191
FR_SWS: 5.0238 hz. CV_SWS: 1.5317
Portion_ISI>2s: 0.012631
Post Spike Supp: 5.5 ms
P_2V: 784.7807 μs Peak_Width: 171.6667 μs
Non-Laser
Laser
0.5 ms
20 (μV)
Laser response 10 ms 10 mW
Trial
Time (ms)
reliability: 0.51
delta lat: -492 ms
p = 6.09e-298
Time (ms)
Autocorrelation
counts
Time (ms)
Awake
SWS
ISI (ms)
CNI Latency
Time (s)
Reaction Time
Time (ms)
Movement Time
Time (ms)
Probability
Trial Value
Firing Rate (Hz)
Time (min)
Light-On
Center-In
Go Cue
Center-Out
Side-In
Side-Out
Food-Port-In
Trial
Firing Rate (Hz)
Unengaged
Engaged
Ipsi
Contra
Unrewarded
Rewarded
Time (s)
Unpredicted
Reward
Cue onset
75%
25%
0%
Rewarded
Unrewarded
Time (s)

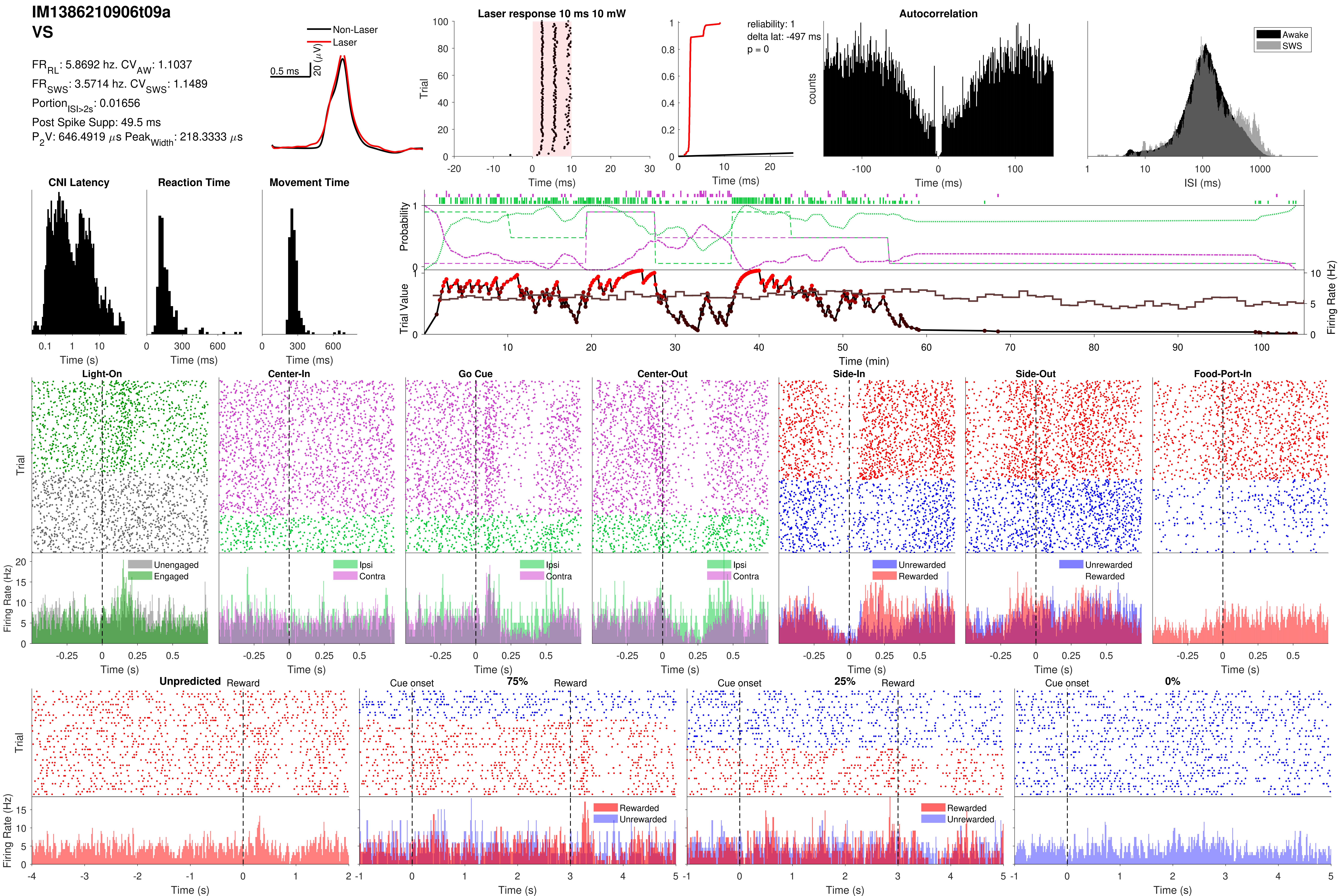

IM1386210906t09a
VS
FR_RL: 5.8692 hz. CV_AW: 1.1037
FR_SWS: 3.5714 hz. CV_SWS: 1.1489
Portion_ISI>2s: 0.01656
Post Spike Supp: 49.5 ms
P_2V: 646.4919 µs Peak_Width: 218.3333 µs
Non-Laser
Laser
0.5 ms
20 (µV)
Laser response 10 ms 10 mW
Trial
Time (ms)
reliability: 1
delta lat: -497 ms
p = 0
Autocorrelation
counts
Awake
SWS
ISI (ms)
CNI Latency
Reaction Time
Movement Time
Time (s)
Probability
Trial Value
Firing Rate (Hz)
Time (min)
Light-On
Center-In
Go Cue
Center-Out
Side-In
Side-Out
Food-Port-In
Unengaged
Engaged
Ipsi
Contra
Unrewarded
Rewarded
Unpredicted
Reward
Cue onset
75%
25%
0%

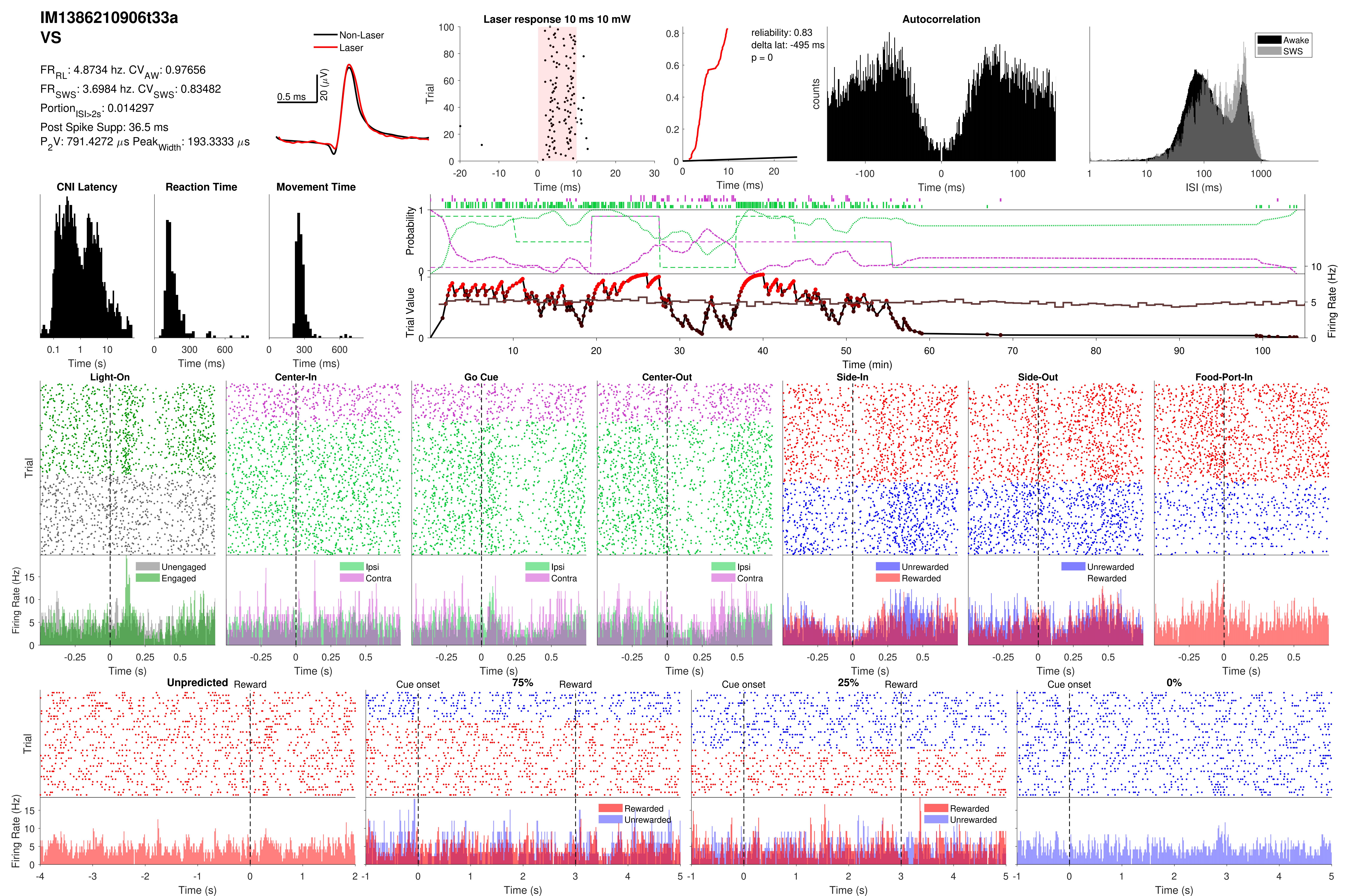
IM1386210906t33a
VS
FR_RL: 4.8734 hz. CV_AW: 0.97656
FR_SWS: 3.6984 hz. CV_SWS: 0.83482
Portion_ISI>2s: 0.014297
Post Spike Supp: 36.5 ms
P_2V: 791.4272 μs Peak_Width: 193.3333 μs
Non-Laser
Laser
0.5 ms
20 (μV)
Laser response 10 ms 10 mW
Trial
Time (ms)
reliability: 0.83
delta lat: -495 ms
p = 0
Time (ms)
Autocorrelation
counts
Time (ms)
Awake
SWS
ISI (ms)
CNI Latency
Time (s)
Reaction Time
Time (ms)
Movement Time
Time (ms)
Probability
Trial Value
Firing Rate (Hz)
Time (min)
Light-On
Center-In
Go Cue
Center-Out
Side-In
Side-Out
Food-Port-In
Trial
Firing Rate (Hz)
Unengaged
Engaged
Ipsi
Contra
Unrewarded
Rewarded
Time (s)
Unpredicted
Reward
Cue onset
75%
25%
0%
Rewarded
Unrewarded
Time (s)

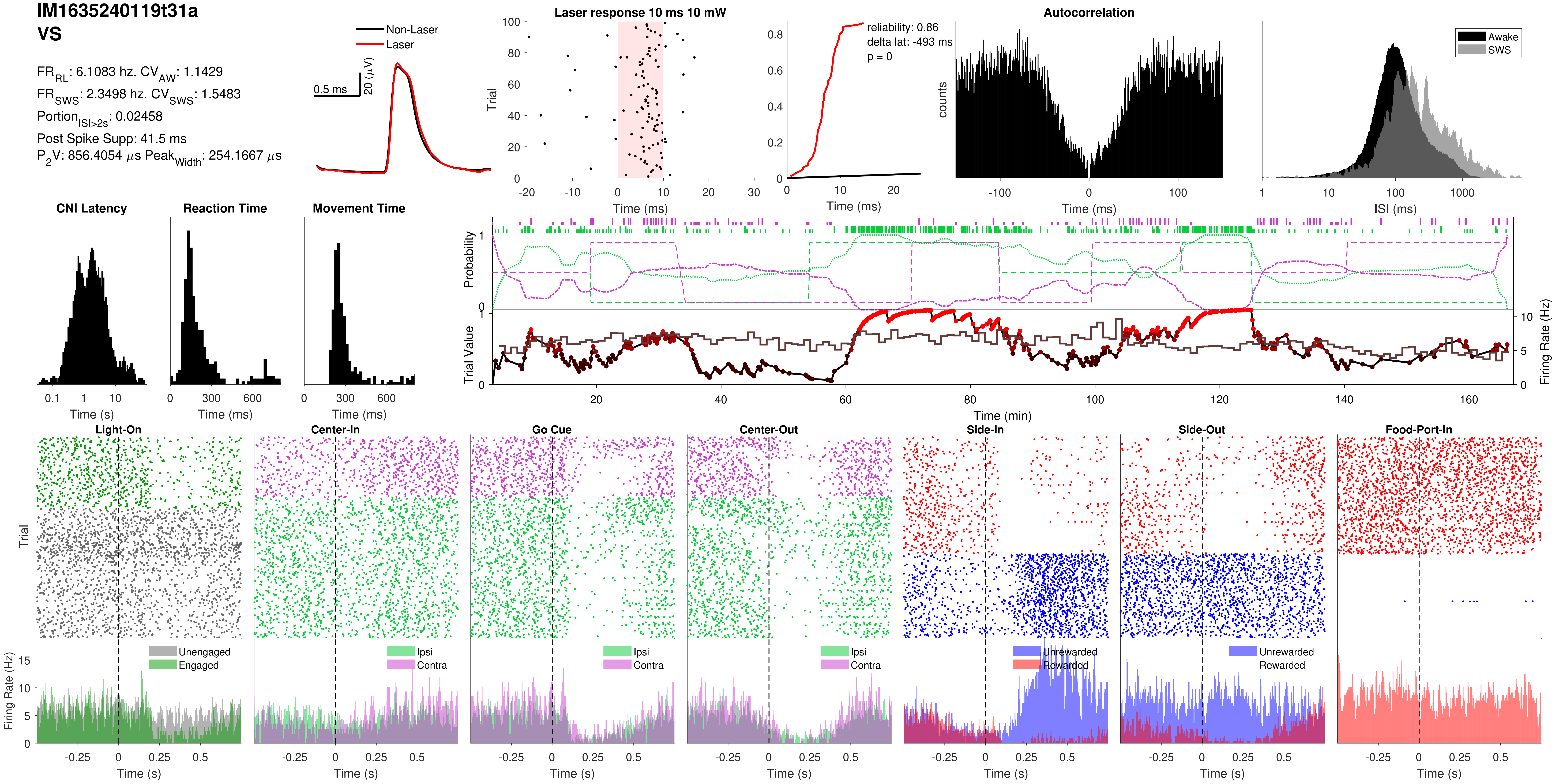
IM1635240119t31a
VS
FR_RL: 6.1083 hz. CV_AW: 1.1429
FR_SWS: 2.3498 hz. CV_SWS: 1.5483
Portion_ISI>2s: 0.02458
Post Spike Supp: 41.5 ms
P_2V: 856.4054 µs Peak_Width: 254.1667 µs
Non-Laser
Laser
0.5 ms
20 (µV)
Laser response 10 ms 10 mW
Trial
Time (ms)
reliability: 0.86
delta lat: -493 ms
p = 0
Autocorrelation
counts
Awake
SWS
ISI (ms)
CNI Latency
Reaction Time
Movement Time
Time (s)
Time (ms)
Probability
Trial Value
Firing Rate (Hz)
Time (min)
Light-On
Center-In
Go Cue
Center-Out
Side-In
Side-Out
Food-Port-In
Unengaged
Engaged
Ipsi
Contra
Unrewarded
Rewarded
Time (s)

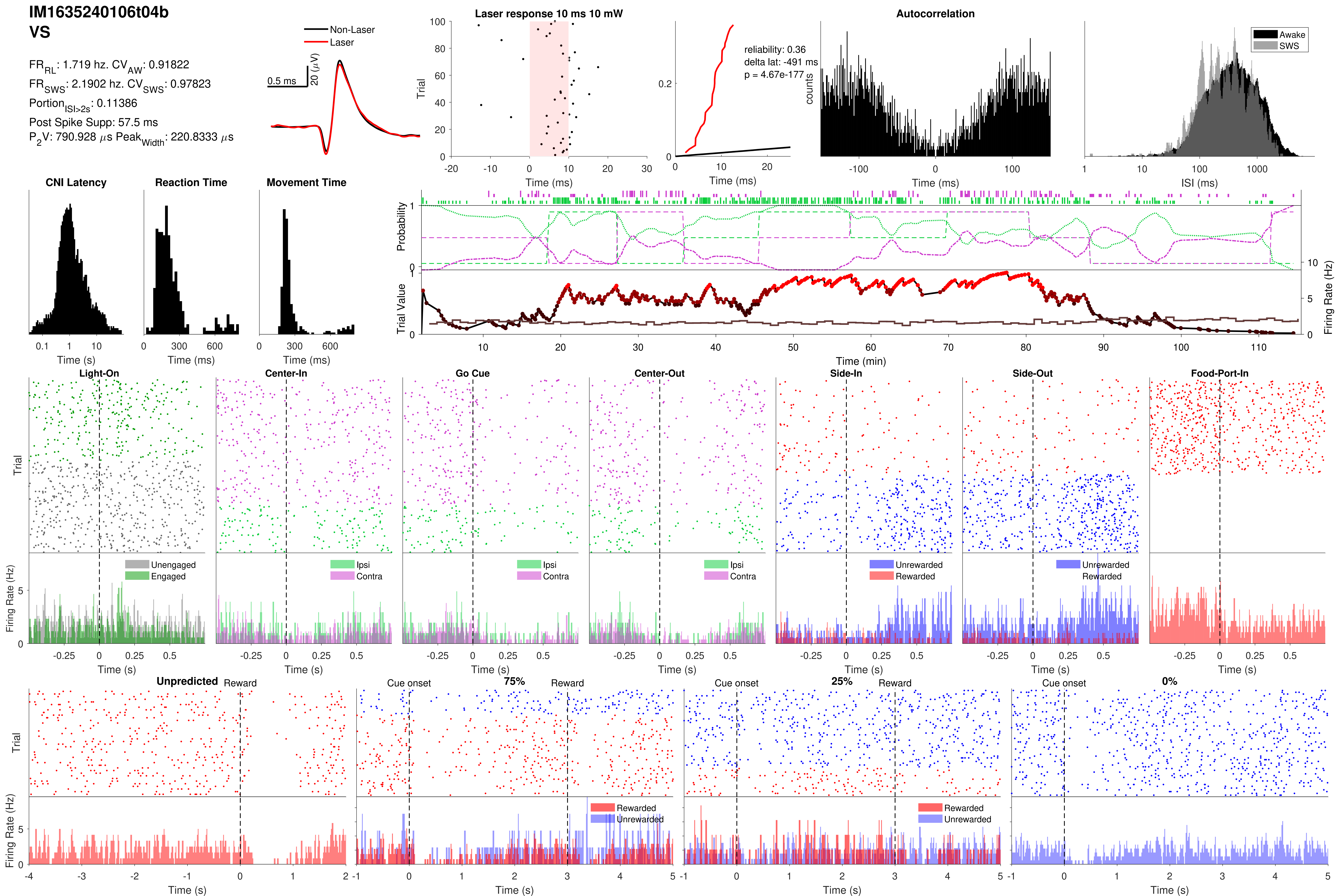
IM1635240106t04b
VS
FR_RL: 1.719 hz. CV_AW: 0.91822
FR_SWS: 2.1902 hz. CV_SWS: 0.97823
Portion_ISI>2s: 0.11386
Post Spike Supp: 57.5 ms
P_2V: 790.928 μs Peak_Width: 220.8333 μs
Non-Laser
Laser
0.5 ms
20 (μV)
Laser response 10 ms 10 mW
Trial
Time (ms)
reliability: 0.36
delta lat: -491 ms
p = 4.67e-177
Time (ms)
Autocorrelation
counts
Time (ms)
Awake
SWS
ISI (ms)
CNI Latency
Time (s)
Reaction Time
Time (ms)
Movement Time
Time (ms)
Probability
Trial Value
Firing Rate (Hz)
Time (min)
Light-On
Center-In
Go Cue
Center-Out
Side-In
Side-Out
Food-Port-In
Trial
Unengaged
Engaged
Ipsi
Contra
Unrewarded
Rewarded
Firing Rate (Hz)
Time (s)
Unpredicted
Reward
Cue onset
75%
25%
0%
Rewarded
Unrewarded
Trial
Firing Rate (Hz)
Time (s)

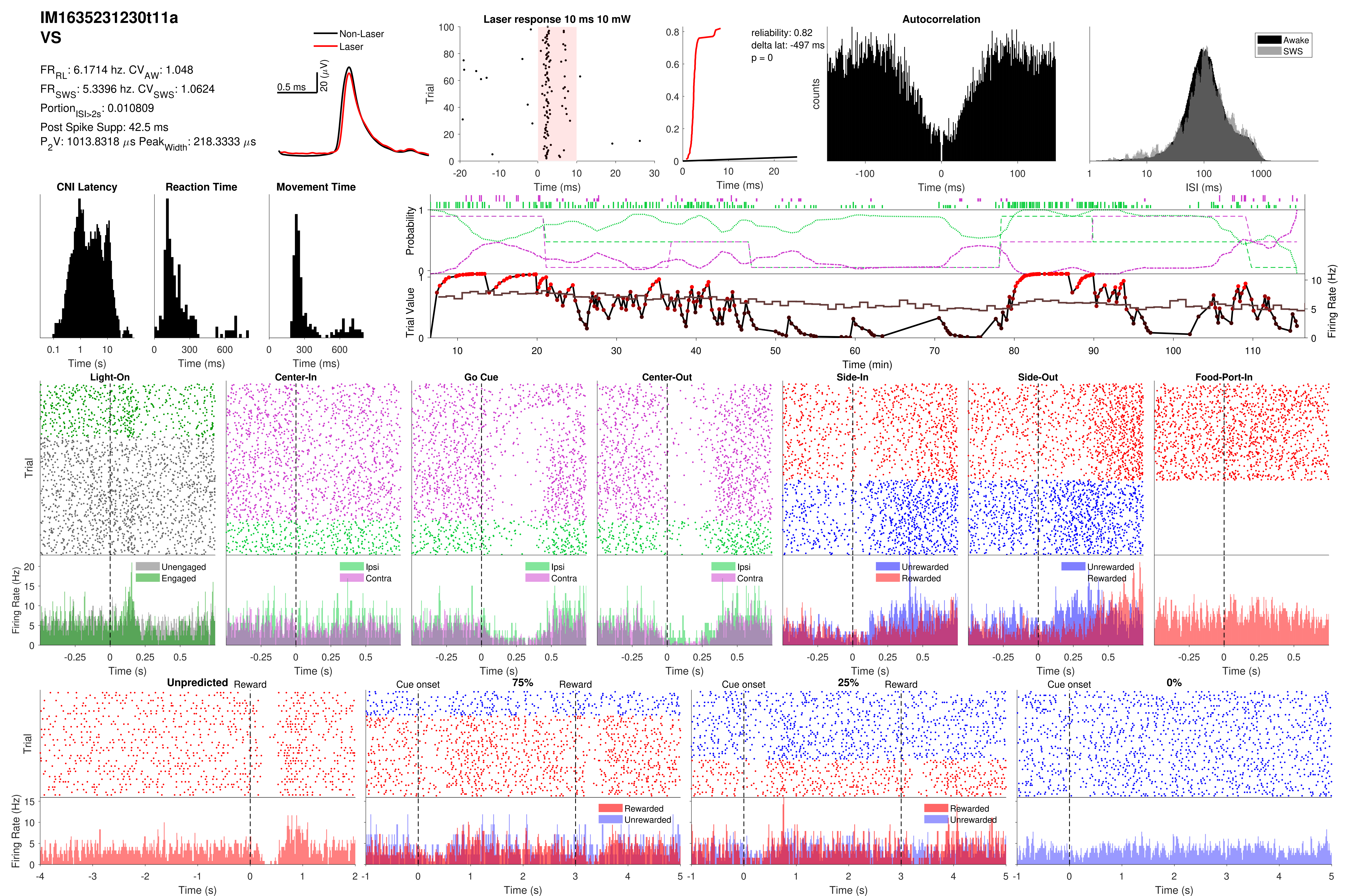
IM1635231230t11a
VS
FR_RL: 6.1714 hz. CV_AW: 1.048
FR_SWS: 5.3396 hz. CV_SWS: 1.0624
Portion_ISI>2s: 0.010809
Post Spike Supp: 42.5 ms
P_2V: 1013.8318 µs Peak_Width: 218.3333 µs
Non-Laser
Laser
0.5 ms
20 (µV)
Laser response 10 ms 10 mW
Trial
Time (ms)
reliability: 0.82
delta lat: -497 ms
p = 0
Time (ms)
Autocorrelation
counts
Time (ms)
Awake
SWS
ISI (ms)
CNI Latency
Time (s)
Reaction Time
Time (ms)
Movement Time
Time (ms)
Probability
Trial Value
Firing Rate (Hz)
Time (min)
Light-On
Center-In
Go Cue
Center-Out
Side-In
Side-Out
Food-Port-In
Trial
Firing Rate (Hz)
Unengaged
Engaged
Ipsi
Contra
Unrewarded
Rewarded
Time (s)
Unpredicted
Reward
Cue onset
75%
25%
0%
Rewarded
Unrewarded
Trial
Firing Rate (Hz)
Time (s)

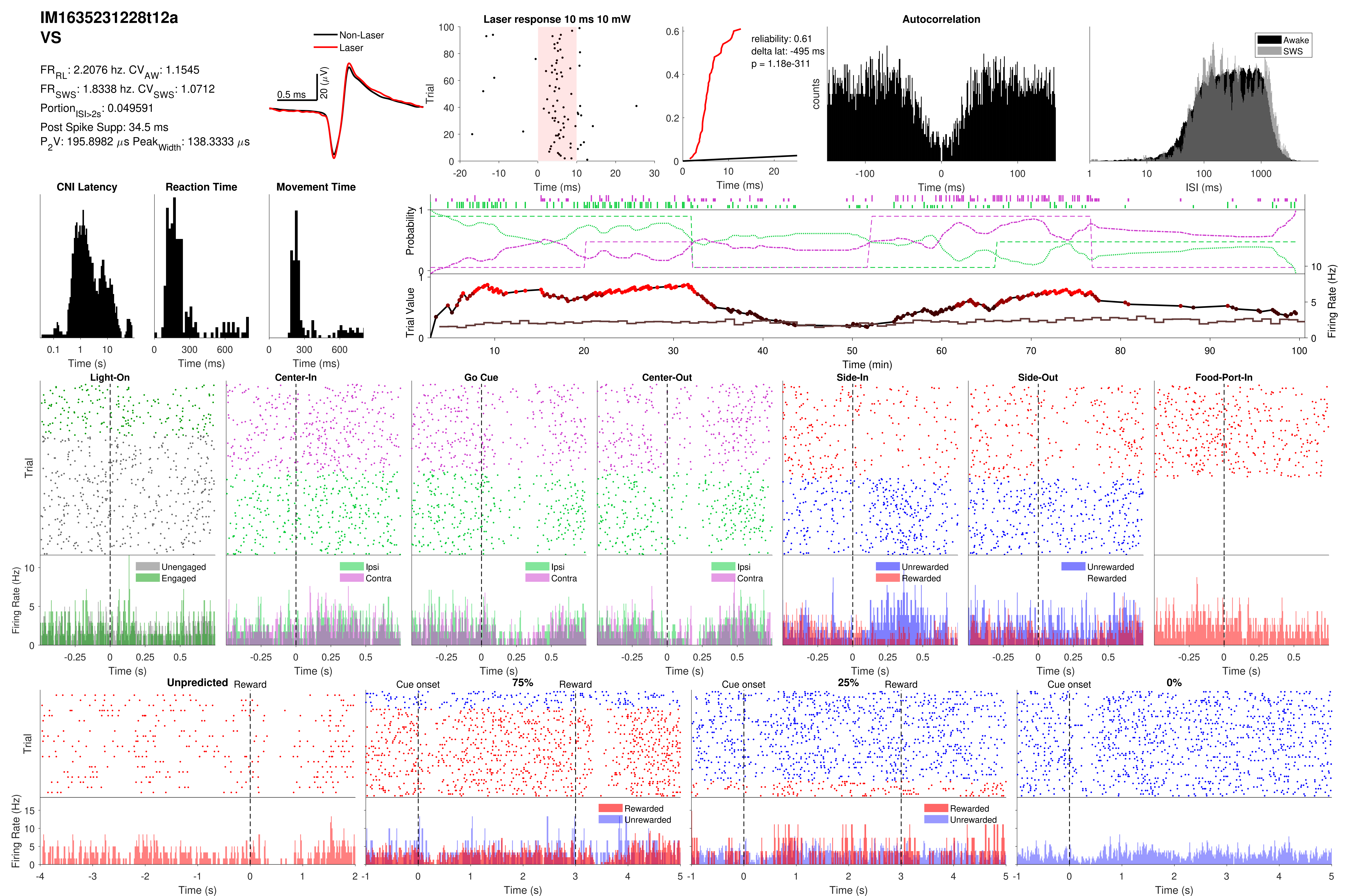

IM1635231228t12a
VS
FR_RL: 2.2076 hz. CV_AW: 1.1545
FR_SWS: 1.8338 hz. CV_SWS: 1.0712
Portion_ISI>2s: 0.049591
Post Spike Supp: 34.5 ms
P_2V: 195.8982 μs Peak_Width: 138.3333 μs
Non-Laser
Laser
0.5 ms
20 (μV)
Laser response 10 ms 10 mW
Trial
Time (ms)
reliability: 0.61
delta lat: -495 ms
p = 1.18e-311
Time (ms)
Autocorrelation
counts
Time (ms)
Awake
SWS
ISI (ms)
CNI Latency
Time (s)
Reaction Time
Time (ms)
Movement Time
Time (ms)
Probability
Trial Value
Firing Rate (Hz)
Time (min)
Light-On
Center-In
Go Cue
Center-Out
Side-In
Side-Out
Food-Port-In
Trial
Firing Rate (Hz)
Unengaged
Engaged
Ipsi
Contra
Unrewarded
Rewarded
Time (s)
Unpredicted
Reward
Cue onset
75%
25%
0%
Rewarded
Unrewarded
Trial
Firing Rate (Hz)
Time (s)
